# Pangenome analysis of *Clostridium scindens*: a collection of diverse bile acid and steroid metabolizing commensal gut bacterial strains

**DOI:** 10.1101/2024.09.06.610859

**Authors:** Kelly Y. Olivos-Caicedo, Francelys Fernandez, Steven L. Daniel, Karthik Anantharaman, Jason M. Ridlon, João M. P. Alves

## Abstract

*Clostridium scindens* is a commensal gut bacterium capable of forming the secondary bile acids deoxycholic acid and lithocholic acid from the primary bile acids cholic acid and chenodeoxycholic acid, respectively, as well as converting glucocorticoids to androgens. Historically, only two strains, *C. scindens* ATCC 35704 and *C. scindens* VPI 12708, have been characterized *in vitro* and *in vivo* to any significant extent. The formation of secondary bile acids is important in maintaining normal gastrointestinal function, in regulating the structure of the gut microbiome, in the etiology of such diseases such as cancers of the GI tract, and in the prevention of *Clostridium difficile* infection. We therefore wanted to determine the pangenome of 34 cultured strains of *C. scindens* and a set of 200 metagenome-assembled genomes (MAGs) to understand the variability among strains. The results indicate that the 34 strains of *C. scindens* have an open pangenome with 12,720 orthologous gene groups, and a core genome with 1,630 gene families, in addition to 7,051 and 4,039 gene families in the accessory and unique (i.e., strain-exclusive) genomes, respectively. The core genome contains 39% of the proteins with predicted metabolic function, and, in the unique genome, the function of storage and processing of information prevails, with 34% of the proteins being in that category. The pangenome profile including the MAGs also proved to be open. The presence of bile acid inducible (*bai*) and steroid-17,20-desmolase (*des*) genes was identified among groups of strains. The analysis reveals that *C. scindens* strains are distributed into two clades, indicating the possible onset of *C. scindens* separation into two species, confirmed by gene content, phylogenomic, and average nucleotide identity (ANI) analyses. This study provides insight into the structure and function of the *C. scindens* pangenome, offering a genetic foundation of significance for many aspects of research on the intestinal microbiota and bile acid metabolism.

## INTRODUCTION

The emulsification of dietary lipids in the aqueous milieu of the vertebrate small bowel represents a fundamental problem that was solved through the evolution of a complex pathway in the liver that converts cholesterol into detergents known as primary bile acids (Hofmann and Hagey 2014). The term “secondary bile acid” was coined in 1960, denoting microbial conversion products of host primary bile acids produced in the liver (Ridlon et al. 2023). That year, the same research group proposed a two-step mechanism for bile acid 7α-dehydroxylation that we have referred to as the Samuelsson-Bergstrӧm model (Ridlon et al. 2023). Yet, the cultivation and storage of anaerobic bacteria capable of bile acid 7α-dehydroxylation were enduring problems that slowed progress in the field (Ridlon et al. 2023). It wasn’t until the early 1980s that strains of *Clostridium scindens* (i.e., VPI 12708 and ATCC 35704) were characterized so that the Samuelsson-Bergström model could be adequately tested (Hylemon et al. 1980, White et al. 1980, Winter et al. 1984, Morris et al. 1985).

In 1991, a new, more complex model of bile acid 7α-dehydroxylation was proposed, which we have recently termed the Hylemon-Bjӧrkhem (HB) pathway that uniquely explains the formation of allo-secondary bile acids (Ridlon et al. 2023). A bile acid-inducible (*bai*) regulon in *C. scindens* VPI 12708 was cloned during the 1990s, and studies spanning the subsequent three decades have uncovered enzymes catalyzing each step of the HB pathway (Ridlon and Gaskins 2024). This biochemical pathway involved in 7α-dehydroxylation of bile acids is encoded by the *bai* operon which is restricted to a phylogenetic group of bacterial species belonging to the *Clostridium* clusters IV (*Ruminococcaceae*), XI (*Peptostreptococcaceae*), and XIVa (*Lachnospiraceae*), of which the most extensively studied species in *Lachnospiraceae* are *C. scindens* and *Clostridium hylemonae* (Devendran et al. 2019, Marion et al. 2019, Marion et al. 2020, Ridlon et al. 2020, Ridlon et al. 2023), and most recently *Faecalicatena contorta* (Jin et al. 2022).

There is much current interest in the role of bai genes and in the formation of hydrophobic secondary bile acids such as deoxycholic acid (DCA) and lithocholic acid (LCA) in gastrointestinal (GI) diseases (Ridlon and Gaskins 2024). For many decades, the enrichment of DCA and LCA associated with Western Diets high in animal protein and saturated fat were linked with increased risk for colorectal cancer (CRC) (O’Keefe 2016). Diet exchange studies demonstrate that high animal protein and fat diets drive the elevation of bile acid-metabolizing genes and functional activities such as increased bile salt hydrolysis (*bsh*) and bile acid 7α-dehydroxylation (*bai*) genes (David et al. 2014, O’Keefe et al. 2015). Indeed, there is compelling evidence for the co-carcinogenic role of hydrophobic secondary bile acids in the GI tract (Bernstein and Bernstein 2023). Meta-analysis of metagenomic studies reveal an enrichment of *bai* genes in CRC (Wirbel et al. 2019). The antimicrobial nature of hydrophobic secondary bile acids also results in lower gut microbial diversity (Islam et al. 2011), which is also associated with development of CRC (Ridlon et al. 2016). A recent study provided compelling evidence that disruption of *baiH* in *F. contorta* reduces dextran sodium sulfate (DSS)-induced colitis in mice (Jin et al. 2022)*. C. scindens* also appears to exacerbate bile acid diarrhea, at least in mice, through alteration in liver production of primary bile acids that inhibit ileal FGF15 production through decreased FXR activation (Zhao et al. 2020).

Current thinking about secondary bile acids has expanded in recent years as new evidence accumulates regarding the importance of maintaining moderate levels of DCA and LCA (O’Keefe 2016, Ocvirk and O’Keefe 2021). *Clostridium difficile* infection (Abt et al. 2016) and inflammatory bowel disease (Heinken et al. 2019) are associated with gut dysbiosis and low levels of fecal secondary bile acids and enrichment in primary conjugated bile acids. *C. scindens* remains one of limited taxa in phylum *Bacillota* capable of forming DCA and LCA in the vertebrate GI tract (Ridlon and Gaskins 2024). Studies with complex consortia of gut bacteria both *in vitro* and *in vivo* confirm that, despite their low abundance in the gut microbiome, the expression of functional *bai* genes by bile acid 7α-dehydroxylating bacteria is a necessary condition for generating DCA and LCA from primary bile acids (Jin et al. 2022, Wang et al. 2023). It has been suggested that the reestablishment of DCA levels in feces along with *C. scindens* is important in the treatment and prevention of *C. difficile* (re)infection (Buffie et al. 2015, Abt et al. 2016). This points to the potential for *C. scindens* or the *bai* pathway to have therapeutic benefit in certain acute or even chronic conditions in the GI tract (Wise and Cummings 2022). By contrast, controlling the output of primary bile acids and maintaining moderate levels of DCA and LCA through diets low in animal protein and fat and high in vegetables and fiber may be beneficial in other clinical contexts such as CRC (Ridlon et al. 2016).

However, our understanding of *C. scindens* is very limited to date with much of the *in vitro* and *in vivo* work focusing on *C. scindens* ATCC 35704 (Studer et al. 2016, Marion et al. 2020, Wang et al. 2023), the type strain, and *C. scindens* VPI 12708 (Ridlon et al. 2023), the first isolate shown to be capable of 7α-dehydroxylation. While assumed, it has not been determined whether the *bai* pathway is found in most (or all) strains of this species, and the extent of genomic diversity between strains of *C. scindens* is currently unknown. In selection of potentially therapeutic strains, it will be important to have baseline information relating to the genomic diversity of a gut bacterial species (Martín et al. 2023). Here, we present a current comprehensive genomic study of *C. scindens* strains from both cultured isolates and metagenome assembled genomes (MAGs). Our major findings are that (1) all strains identified as *C. scindens* possess the *bai* regulon; (2) isolates with >97% full length 16S rRNA gene identity separate into two groups with ∼4-5% difference in genomic sequence, suggesting that potentially two separate microbial species are present, or are in the process of speciating; and (3) while the core genome is closed, the pangenome remains open, indicating that additional strain diversity exists.

## RESULTS

### Assembly and genomic characteristics of C. scindens strains

Comparative analysis of the recently sequenced *C. scindens* strains and the genomes available at NCBI showed strain variation with respect to genome size. The genome lengths are between 3.2 and 4.6 Mbp, belonging to strains MSK.1.26 and DFI.1.130, respectively, and presenting 3,140 to 4,177 coding sequences (CDS). Percent GC composition showed little variation, ranging from 45.5 to 47.8%, with strains BL389WT3D and VE202-05 having the lowest and highest values, respectively. Genomic characteristics of the 34 *C. scindens* strains, such as the number of contigs and RNA types are shown in **Table 1**.

**Table 1.** Genomic characteristics of 34 *C. scindens* strains.

|  | Strain | N° contigs | Genome size (bp) | CDS | rRNA | tRNA | tmRNA | G+C% |
| --- | --- | --- | --- | --- | --- | --- | --- | --- |
| 1 | I10 | 1 | 3,435,295 | 3602 | 12 | 56 | 2 | 46.5 |
| 2 | JCM10419 | 1 | 3,940,699 | 3682 | 12 | 57 | 1 | 47.5 |
| 3 | JCM10420 | 1 | 4,020,045 | 3768 | 11 | 57 | 1 | 47.4 |
| 4 | JCM10422 | 2 | 3,501,106 | 3390 | 12 | 56 | 1 | 46.6 |
| 5 | JCM10423 | 1 | 4,306,053 | 4057 | 11 | 57 | 1 | 46.9 |
| 6 | MO32 | 1 | 3,929,075 | 3691 | 11 | 57 | 1 | 47.8 |
| 7 | NTI82 | 1 | 4,318,168 | 4137 | 11 | 59 | 1 | 47.1 |
| 8 | S076 | 1 | 4,290,604 | 4103 | 11 | 58 | 1 | 47.3 |
| 9 | S077 | 5 | 3,403,497 | 3386 | 12 | 57 | 2 | 46.5 |
| 10 | VPI12708* | 1 | 3,983,052 | 3716 | 12 | 56 | 1 | 47.7 |
| 11 | ATCC 35704* | 1 | 3,658,040 | 3656 | 12 | 58 | 2 | 46.3 |
| 12 | BL389WT3D* | 1 | 3,785,527 | 3655 | 12 | 58 | 2 | 45.5 |
| 13 | CE91-St59* | 1 | 3,608,085 | 3602 | 12 | 56 | 2 | 45.9 |
| 14 | CE91-St60* | 1 | 3,608,087 | 3608 | 12 | 56 | 2 | 45.9 |
| 15 | FDAARGOS_12<br>27* | 1 | 3,619,096 | 3603 | 12 | 57 | 2 | 46.3 |
| 16 | G10* | 1 | 3,315,593 | 3295 | 12 | 57 | 2 | 46.6 |
| 17 | Q4* | 1 | 3,941,835 | 3725 | 12 | 57 | 1 | 47.7 |
| 18 | AM05-22* | 56 | 3,330,149 | 3269 | 3 | 54 | 2 | 46.5 |
| 19 | AM07-30* | 50 | 3,331,670 | 3272 | 2 | 49 | 2 | 46.5 |
| 20 | DFI.1.130* | 797 | 4,565,863 | 4177 | 4 | 53 | 1 | 46.9 |
| 21 | DFI.1.161* | 160 | 4,325,545 | 4041 | 4 | 51 | 1 | 47.1 |
| 22 | DFI.1.162* | 125 | 4,396,306 | 4161 | 4 | 52 | 1 | 46.9 |
| 23 | DFI.1.217* | 96 | 4,389,871 | 4161 | 4 | 52 | 1 | 46.9 |
| 24 | DFI.1.234* | 107 | 4,316,625 | 4072 | 4 | 52 | 1 | 47.0 |
| 25 | DFI.1.60* | 197 | 4,309,437 | 4047 | 4 | 52 | 1 | 47.1 |
| 26 | DFI.4.63* | 195 | 4,167,375 | 3958 | 4 | 52 | 1 | 47.5 |
| 27 | GGCC_0168* | 256 | 3,417,088 | 3390 | 12 | 57 | 2 | 46.6 |
| 28 | MGYG-HGUT-013<br>03* | 41 | 3,622,605 | 3574 | 22 | 64 | 2 | 46.4 |
| 29 | MSK.1.16* | 93 | 3,230,100 | 3143 | 4 | 54 | 2 | 46.4 |
| 30 | MSK.1.26* | 93 | 3,227,969 | 3140 | 3 | 54 | 2 | 46.4 |
| 31 | MSK.5.24* | 21 | 4,072,709 | 3845 | 4 | 54 | 1 | 47.5 |
| 32 | NB2A-7-D5* | 39 | 4,182,602 | 3980 | 4 | 60 | 1 | 47.5 |
| 33 | SL.1.22* | 52 | 3,970,092 | 3706 | 3 | 55 | 1 | 47.5 |
| 34 | VE202-05* | 102 | 3,912,387 | 4523 | 2 | 48 | 0 | 47.8 |

\*: Genomes previously sequenced and available in GenBank.

The BUSCO tool was used to assess the completeness of the genome assembly of the 34 *C. scindens* strains using the Clostridia dataset with 247 orthologs as reference. All strains have a high quality, with almost complete sets of single-copy orthologous genes (between 245 and 247), except strain VE202-05 which showed a greater amount of fragmented and missing genes, 35 and 13 genes respectively (**Supplementary** Figure 1a). Only 6 strains (AM05-22, AM07-30, BL389WT3D, DFI.4.63, G10, and Q4) have 1 fragmented ortholog, while 15 strains have 1 missing ortholog each. Assembly completeness of the MAGs was assessed by the CheckM tool, giving maximum and minimum values of 99.42% and 50.78%, respectively. The results indicate the presence of markers or sets of single-copy genes in each genome analyzed. Data is presented in **Supplementary** Figure 1b **and Supplementary Table 1**.

### Pangenome of the 34 cultured strains of C. scindens

The Roary analysis identified a pangenome containing 12,720 gene families, distributed in the core genome, accessory genome, and unique or strain-specific genes. A total of 1,630 gene groups are in the core, representing almost 13% of the total pangenome, with 7,051 accessory groups (**Figure 1**). Furthermore, 4,039 strain-specific genes distributed among *C. scindens* strains are represented (**Figure 2**), where 19 strains have between 1 and 55 strain-specific genes, and 15 have more than 100 genes. The human-associated strain VE202-05 (Atarashi et al. 2013) has the highest number of unique genes with a total of 907 genes, followed by BL389WT3D with 420 genes, which might be due to its isolation from pig feces (Wylensek et al. 2020).

**Figure 1.**
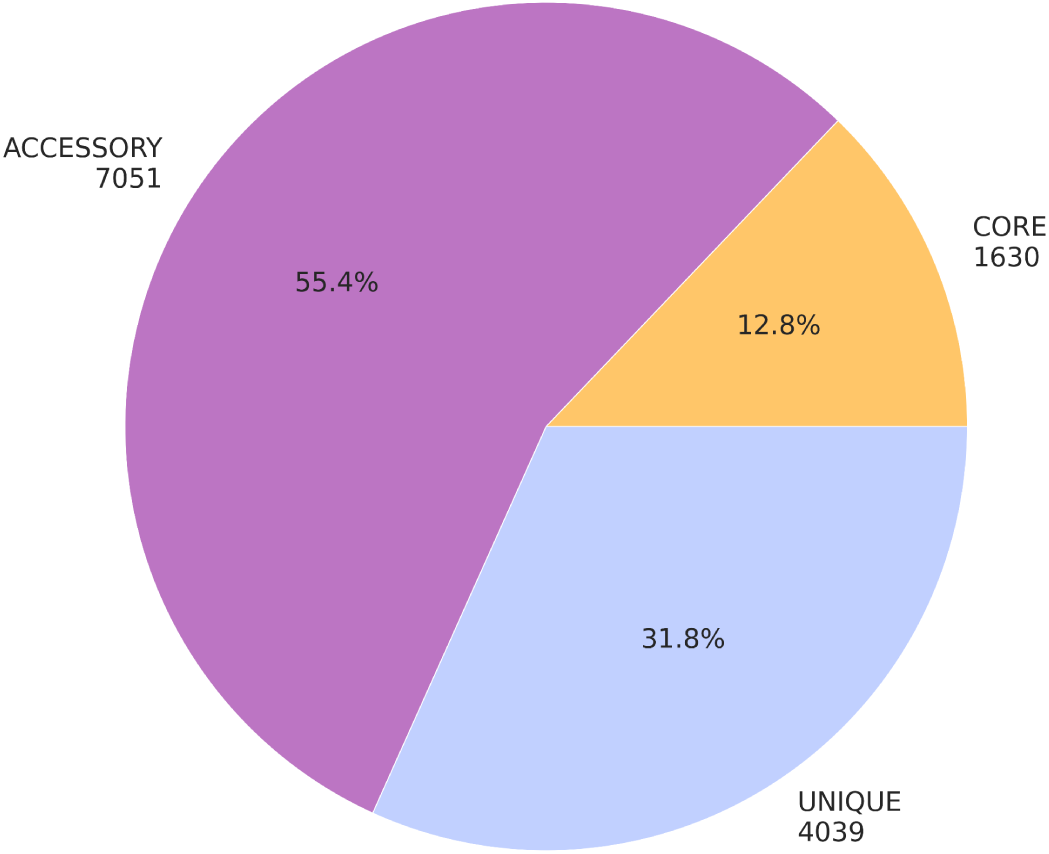
Distribution of gene families in the pangenome of 34 *C. scindens* strains. The graph shows the number and percentage of core, accessory and unique gene groups.

**Figure 2.**
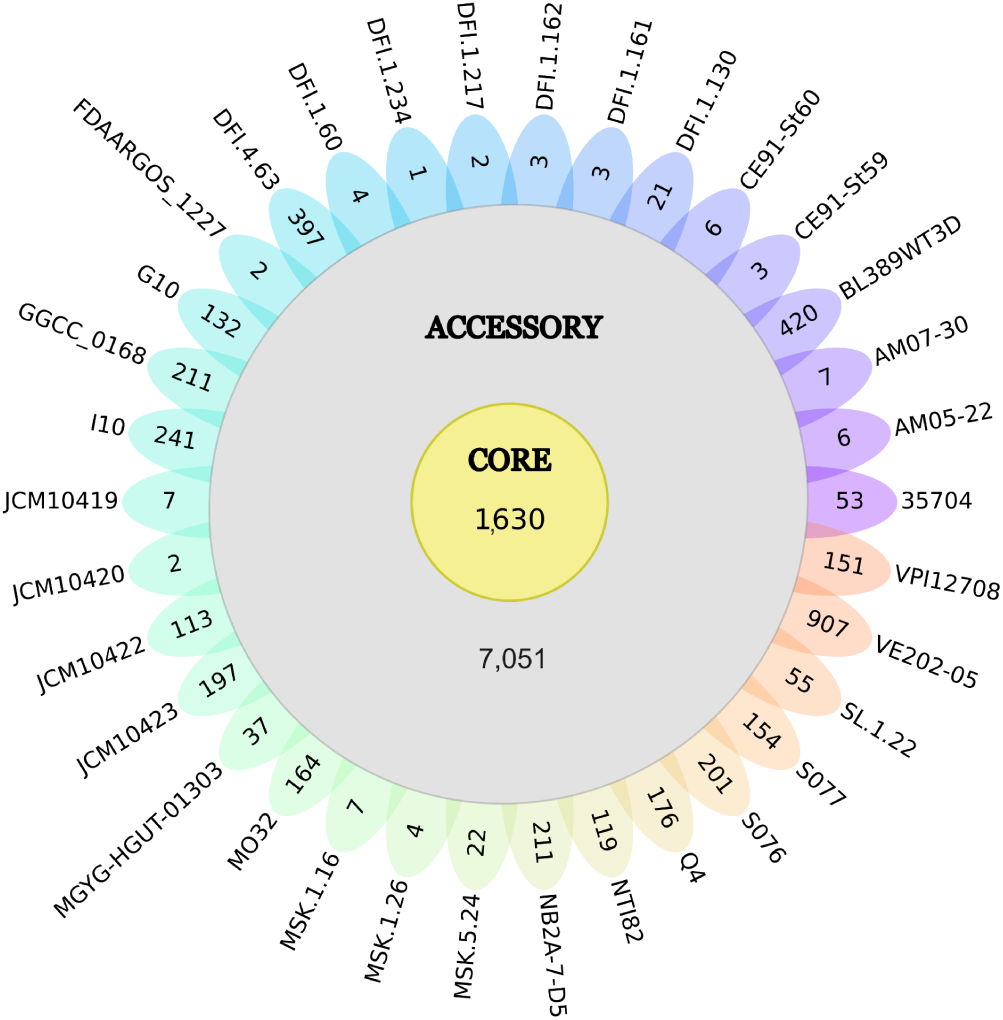
Genetic diversity of *C. scindens*. The flowerplot shows the size of the core genome, accessory genome and unique genes of the 34 C. scindens strains. The number of core gene clusters is represented by the yellow circle in the center of the flower (1,630), the total number of accessory genes is represented by the gray circle (7,051), and the genes unique to each strain are shown on each petal of the flower.

Analysis using a binomial mixture model as implemented in function binomixEstimate from the micropan package yielded an estimate of core genome size of between 1,356 (higher complexity model K=6) and 1,630 (lower complexity model, K=3), which is very close to the number the curve of the core element seems to have stabilized at with 34 genomes (**Figure 3a**). This indicates that the essential set of genes for *C. scindens* has been identified and should not change significantly with the sequencing of more strains. The identification of the core genome allows us to infer characteristics such as the bacteria’s lifestyle, as this group of genes can encode resistance to antibiotics and heavy metals, cell wall components, virulence, and metabolic genes, among other information. On the other hand, strain-unique genes, which do not bear similarity to closely related strains, confer biological individuality, host specificity, and pathogenesis (Bhardwaj and Somvanshi 2017).

**Figure 3.**
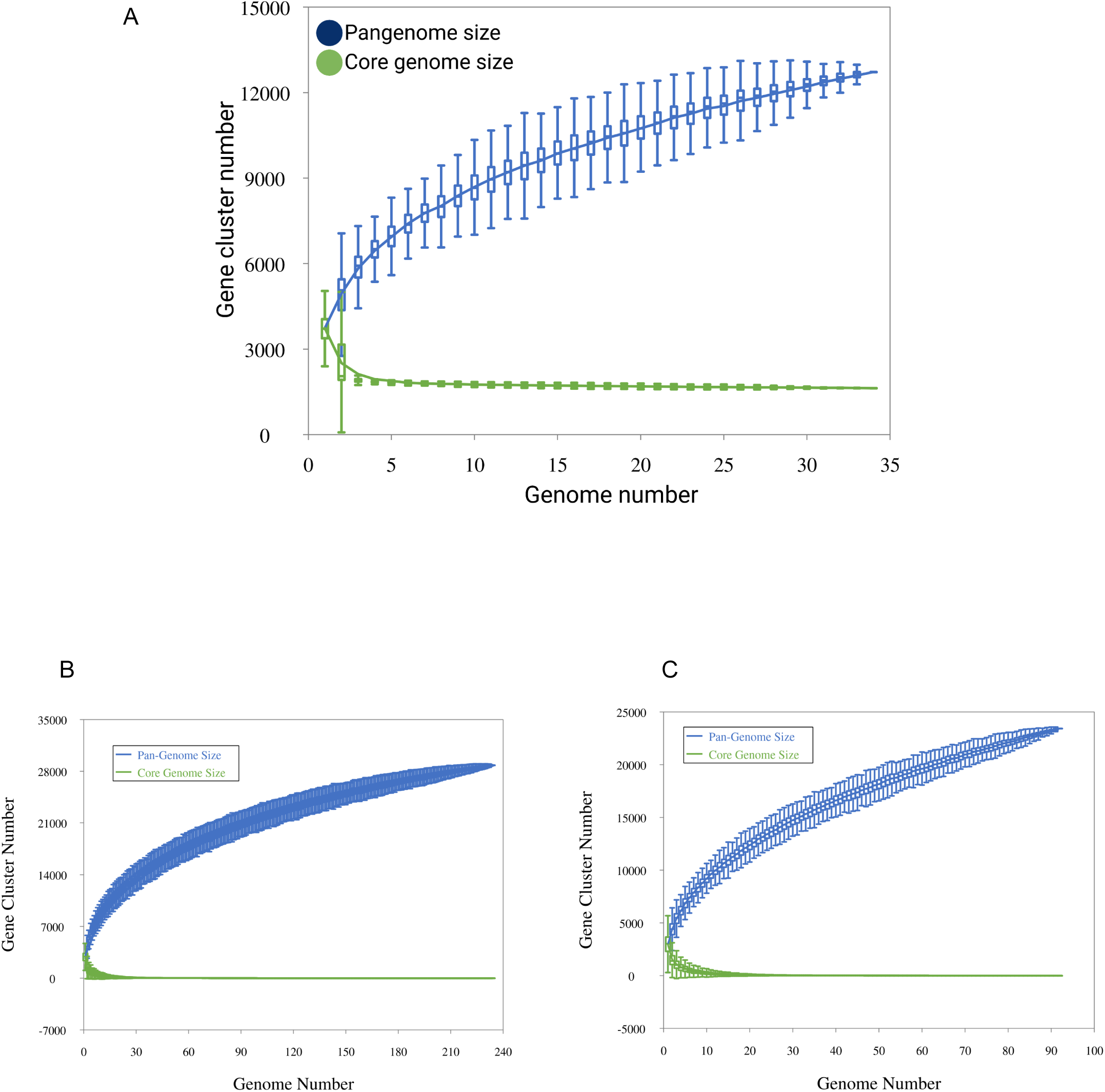
Pan-core plots for *C. scindens.* **A.** Pan-core plot of the 34 cultured strains of *C. scindens.* **B.** Pan-core plot including 200 *C. scindens* MAGs**. C.** Pan-core plot including 58 *C. scindens* dereplicated MAGs. The graphics show cumulative curves of the upward trend in the number of pangenome gene families (in blue) and the downward trend of core gene families (in green) with each consecutive addition of a *C. scindens* genome. The rising curve in blue shows an open pangenome

### Pangenome analysis after addition of C. scindens metagenome assembled genomes

Pangenome analysis of *C. scindens* MAGs was determined considering the percentage of completeness of genomes, with analysis being carried out for each level of completeness. A group of 200 MAGs and another with 58 dereplicated MAGs were analyzed. To calculate the pangenome, the 34 previously described complete genomes of *C. scindens* were also included. The analysis shows that, as MAG completeness decreases, the pangenome size tends to increase, while the number of gene groups in the core genome tends to be absent (**Table 2**), thus sampling effects due to genome incompleteness severely affect the estimation of core size. After testing multiple completeness values with the 200 MAGs, the core genome was determined using MAGs with a completeness value of at least 85%. A total of 157 genomes shows a pangenome size of 19,189 gene families and a core genome of 132 gene groups, or almost 7% of the total pangenome (**Table 2a**). Pangenomic analysis of MAGs can be affected by fragmentation, incompleteness, and contamination (Li and Yin 2022). For example, it has been found that fragmentation and incompleteness lead to a significant loss of core genes, which translates into incorrect pangenomic functional predictions and inaccurate phylogenetic trees, and on the other hand, contamination influences the accessory genome. Accordingly, in our analyses, the core genome size was reduced by at least 50% for each increase of 5% in incompleteness (**Table 2**). The use of a higher completeness threshold, such as 95 or 90%, is recommended when defining core genes and carrying out a pangenome analysis combining complete genomes and MAGs.

**Table 2:** Analysis results of the *C. scindens* MAGs pangenomes, depending on the completeness of the genomes used in the analysis. **A.** Analysis results of the 200 *C. scindens* MAGs. **B.** Analysis results of the *C. scindens* dereplicated MAGs. The size of the pangenome and *core* genome for each MAG completeness value is shown.

| Completeness of MAGs % | Pangenome | core genome |
| --- | --- | --- |
| ≥ 95 | 14,625 | 850 |
| ≥ 90 | 17,713 | 401 |
| ≥ 85 | 19,189 | 132 |
| ≥ 80 | 19,792 | 75 |
| ≥ 75 | 21,923 | 8 |
| ≥ 70 | 22,864 | 4 |

B
| Completeness of MAGs % | Pangenome | core genome |
| --- | --- | --- |
| ≥ 95 | 14,464 | 1,120 |
| ≥ 90 | 15,144 | 931 |
| ≥ 85 | 16,168 | 501 |
| ≥ 80 | 16,629 | 276 |
| ≥ 75 | 19,083 | 25 |
| ≥ 70 | 20,023 | 6 |

### Pangenome profile

The plots provided by Roary and PanGP show the relationship between the pangenome, core genome, and the number of genomes, along with the different gene family distribution within the genomes under study, allowing us to estimate the pangenome profile as open or closed. Overall, the number of gene families in the pangenome and core genome increases and decreases, respectively, with each consecutive addition of a *C. scindens* genome, as expected.

According to the analysis of the 34 cultured *C. scindens* strains, the pangenome profile is open (**Figure 3a**), since as the number of sequenced genomes increases, the total number of gene families also increases. This is confirmed by the application of the Heaps’ Law formula, which results in an alpha value of 0.845, consistent with an open pangenome. The core genome, on the other hand, seems to have stabilized. Likewise, after adding 200 total MAGs and dereplicated MAGs of *C. scindens*, the pangenome profile remains open, with a Heaps’ alpha of 0.768, and the core genome becomes nearly absent (**Figure 3b**, **Figura 3c**), due to genome incompleteness, as discussed above.

An open pangenome can be extrapolated into a sympatric lifestyle and the ability to gain new species-specific genes, which could be related to virulence, metabolism, and information storage, among others (Bhardwaj and Somvanshi 2017). An open pangenome also suggests the possibility of adding new gene sets, along with new strain-specific genes (singletons). Furthermore, the persistence of singletons might represent the ability to acquire new virulence traits, a threat to human health. Likewise, an open pangenome indicates that it is necessary to study more genomes, from a greater variety of environments and geographic locations, in order to define the entire genomic content of this species.

### Identification of C. scindens strain groups

We utilized Roary to group the 12,720 gene families of the pangenome into a presence/absence matrix and a phylogenomic tree based on the core genes, identifying a relationship between the tree and the distribution of core and accessory gene families among the 34 *C. scindens* strains (**Figure 4**), which were separated into 2 groups, referred to here as “group 1” (the *C. scindens* ATCC 35704 group) and “group 2” (the *C. scindens* VPI 12708 group).

**Figure 4.**
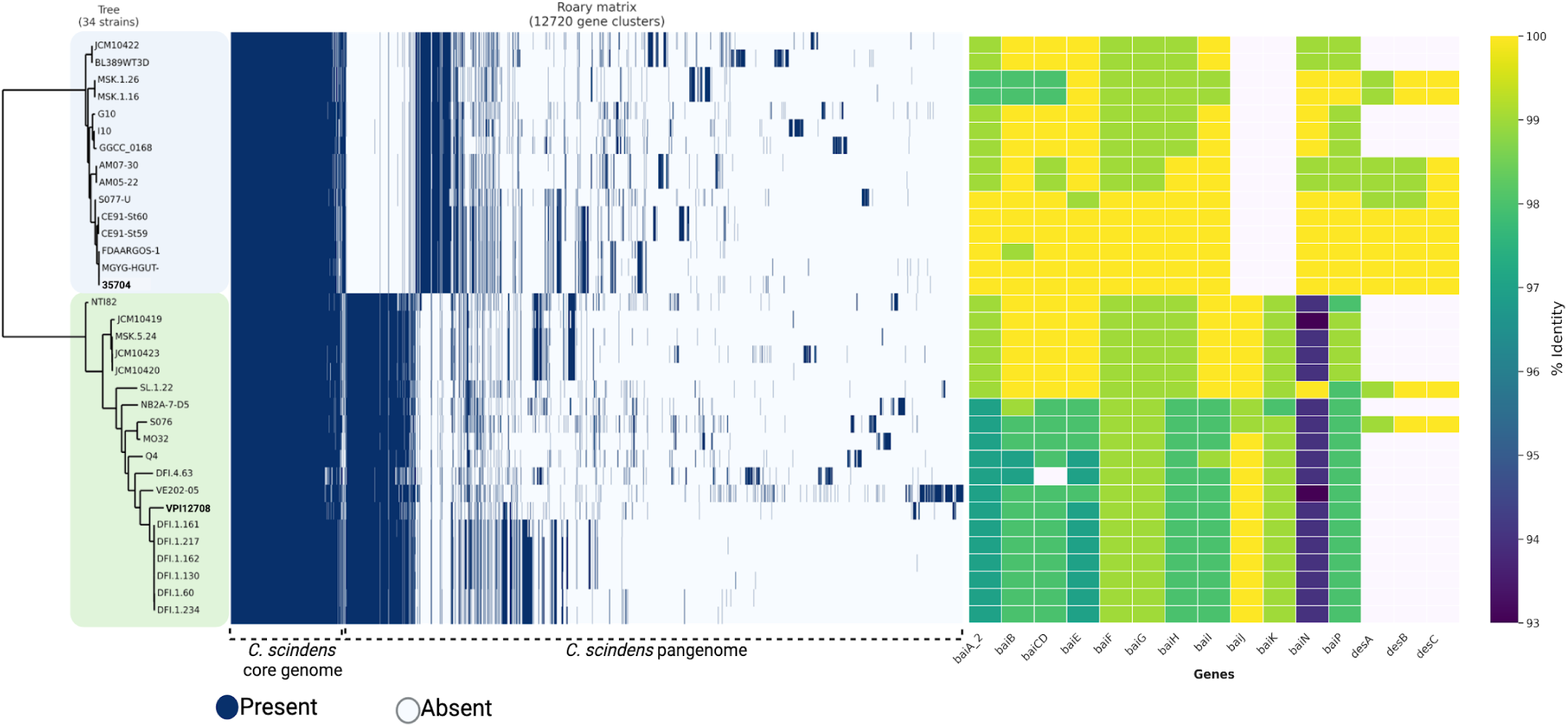
Pangenomic comparison of the 34 *C. scindens* strains. On the left: phylogenomic tree and matrix of presence (blue lines) and absence (white lines) of core and accessory genes of the pangenome. The phylogenomic tree was generated based on 1,490 single-copy orthologous genes shared by the 34 genomes of *C. scindens.* The tree was inferred using the maximum likelihood method by running 1,000 fast bootstrap pseudoreplicates. The clades formed were referred to as Group 1 and Group 2, marked on the tree by light blue and green boxes, respectively. On the right side: Heatmap of *bai* and *des* genes in the 34 *C. scindens* genomes. The percentage value of sequence identity is shown in color; the highest value in yellow and the lowest in purple. The lack of the gene is indicated by a white rectangle.

To represent a pangenome appropriately, the gene frequency spectrum function G(k) needs to be considered, which is defined as the number of orthologous groups containing genes from exactly k genomes (Moldovan and Gelfand 2018). Generally, in a set of strains belonging to the same species, the spectral function of the pangenome is U-shaped, without internal peaks that differentiate the number of gene groups found in the genomes. However, in a mixed sample, the spectrum function will have internal peaks. In other words, it is defined as “homogeneous” if the set of genomes presents a U-shaped spectrum function, and as “non-homogeneous” if the set contains internal peaks (Moldovan and Gelfand 2018). In our analysis of 34 cultured strain genomes, the number of genes specific to one strain is shown on the left (bar “1”), the core genome shared by all 34 strains is shown on the right, and the accessory genome is shown in the bars in between, present in 2 to 33 genomes (**Figure 5**). Our plot does not present the U-shape associated with a homogeneous distribution that would arise from the analysis of a single species, but rather presents internal peaks that split the graph in two main regions. This, associated with the gene content and phylogenomic analyses, suggests the presence of two different groups of *C. scindens* strains.

**Figure 5.**
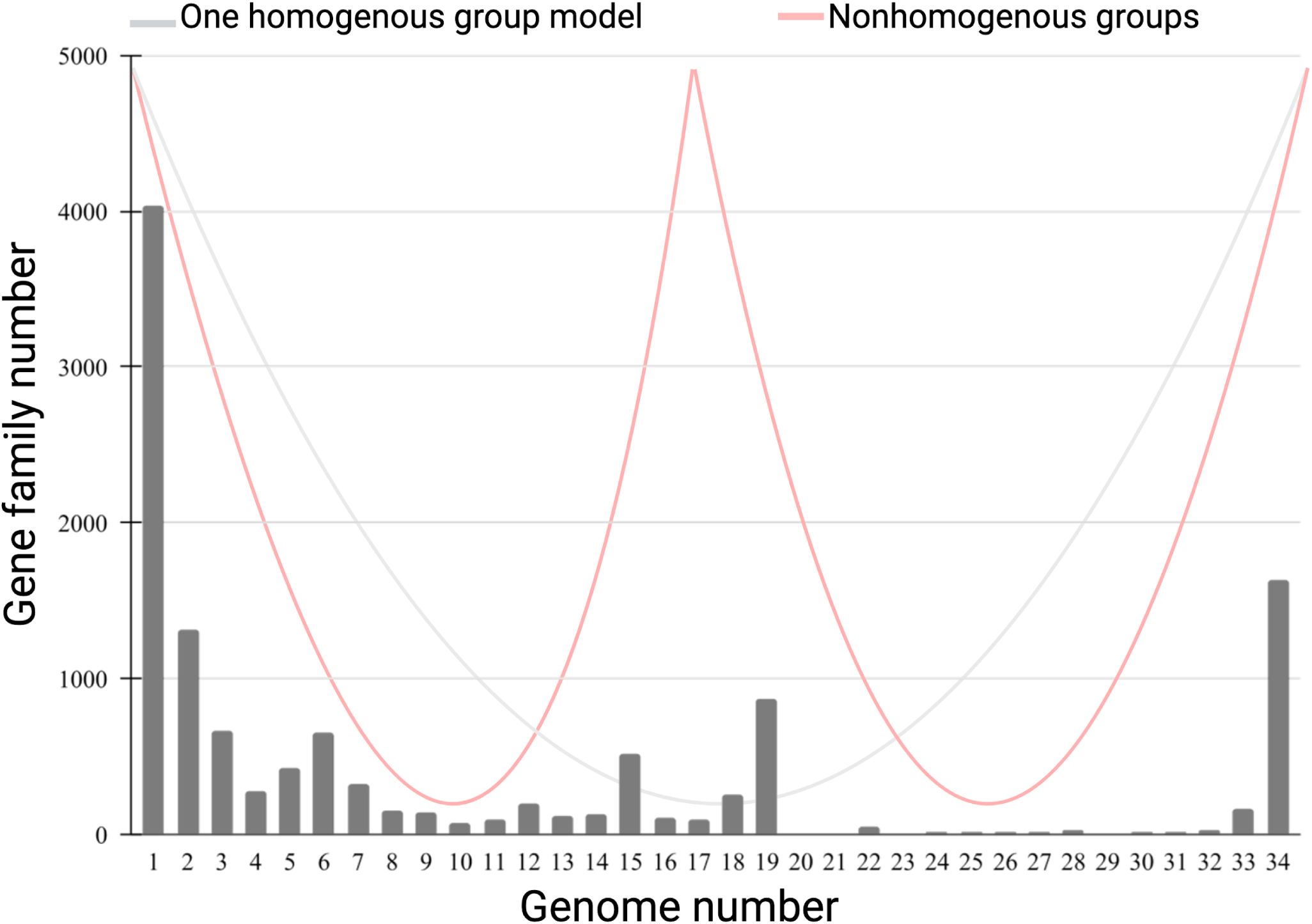
Gene frequency versus number of *C. scindens* genomes. The graph shows the frequency of gene clusters in the 34 *C. scindens* genomes, the left bar “1” represents the number of strain-specific genes, the right bar “34” indicates the core genome, the central bars refer to the accessory genome. Lines are drawn to show the theoretical distribution of a “U” shaped “homogeneous” distribution (gray) and a “W” shaped distribution (red) for “non-homogeneous” plot.

### Average nucleotide identity analysis of 34 *C. scindens* strains

The average percent nucleotide identity calculated by the PyANI tool using the BLAST ANIb method provides the genomic relationship and variation level between the recently sequenced *C. scindens* genomes and the genomes obtained from NCBI, represented by color intensities based on the calculated percentage of identity (**Supplementary** Figure 2). The suggested threshold value for species separation in an ANI analysis is equal to or greater than 95% identity (Richter and Rosselló-Móra 2009). Our results show two groups divided into 15 and 19 strains with a difference of approximately 4-5% in their genomic sequences. Identity within each group is ≥ 98%, while identity between groups is 94.5 to 96%, whereas with the *C. hylemonae* genome the identity values are between 74 and 76%. The present ANI values of *C. scindens* suggest the presence of two possible distinct microbial species or at least an ongoing speciation process.

These results provide a basis for understanding the formation of the two groups of *C. scindens* strains, group 1 and group 2 (**Figure 4**), as previously demonstrated in the pangenome analysis, thus allowing further elucidation of the relationship between the genomes.

### SSU rDNA analysis of 34 C. scindens strains

We have analyzed the SSU rRNA genes from the 34 strains of *C. scindens* and *C. hylemonae*, identifying all ribosomal cistrons, running their maximum likelihood phylogeny, and calculating pairwise sequence distances.

With the exception of two strains, all complete or nearly complete genome sequences (i.e., those containing five contigs or fewer, as shown in Table 1) presented four copies of the complete ribosomal cistron. The two exceptions, strains CE91-St59 and CE91-St60, presented three copies each of the set of ribosomal genes. Except for strain MGYG-HGUT-01303, which had two copies, all other incomplete genomes presented just one copy of the ribosomal cistron. In addition, strains SL.1.22, MSK.1.16, and VE202-05 lacked a portion of the beginning of the SSU rRNA gene sequence, strain MSK.1.26 lacked part of the end, and strain DFI.1.161 had sequence missing both from the start and the end of the gene (**Supplementary File 1**).

The distance values between all *C. scindens* SSU rRNA sequences (**Supplementary** Figure 3 and **Supplementary File 2**), expressed in terms of the uncorrected (i.e., directly observed) number of differences per 100 nucleotides, revealed an interesting pattern, with two different kinds of SSU rRNA gene sequences present in some strains, but not all of them. All SSU rRNA copies within *C. scindens* strains in group 1 (the ATCC 35704 group) genomes belonged to the same sequence type, i.e., they were all very similar, with at most 0.20% of difference (average: 0.05%) in pairwise comparisons; the SSU sequences of different strains were often identical, in fact.

Among strains from group 2 (the VPI 12708 group), the picture is more complex when we look at the SSU rRNA sequence differences, presenting two distinct ranges of dissimilarity present: 0.00 to 0.46% (type A SSU rRNA, average: 0.17%) and 0.85 to 1.18% (type B, average: 1.00%) (**Supplementary File 2**). Almost all of the differences between the two different kinds of SSU rRNA genes found in group 2 are situated between positions 75 and 115 of the alignment (**Supplementary File 1**). As can be seen in the alignment, type B has more similarities with the group 1 SSU rRNA sequences in that region of the molecule than with the type A that is also present in some genomes of group 2 strains. Strains VPI 12708 and MO32 presented two copies of each SSU type, and strain S076 presented three copies of type A and three copies of type B. Of the group 2 strains with more fragmented genome assemblies, all of which had only one ribosomal gene set identified, seven of them (VPI 12708, SL.1.22, DFI.1.130, DFI.1.161, DFI.1.127, DFI.1.234, and DFI.1.60) presented type B sequences while two (DFI.1.162 and DFI.4.63) presented type A sequences. Due to the incompleteness of these genome sequences, it cannot be discarded that some (or even all) of them possess both SSU sequence types.

The distances between group 1 and group 2 sequences agree with the presence of two types of SSU rRNA gene in group 2 genomes, with two ranges of distances present: 0.00 to 0.42% (average: 0.27%) and 1.04 to 1.39% (average: 1.19%). The traditionally suggested threshold value to separate bacterial species using the complete SSU rRNA gene is ≥3%. Therefore, this gene is not reflecting the possible ongoing speciation process as closely as the ANI results described above, since SSU rRNA gene distances never went above 1.39%.

The relatively short distances between group 2’s type B SSU sequences and those found in group 1 are reflected in the phylogenetic tree of the SSU rRNA genes when all copies are included (**Supplementary** Figure 4). By rooting the tree on the branch leading to *C. hylemonae*’s sequence, it can be seen that all group 1 sequences seem to have originated from an internal branch of group 2’s type B SSU rRNA, with its most closely related sequence being one of the copies from strain MO32. The branch lengths on the tree also show that there is much more sequence variation within the sequences from group 2 than those of group 1, indicating that group 1 might have gone through a relatively recent population bottleneck. Both of these observations suggest that group 1 probably descends from a subset of group 1 that specialized in a different niche in the host. However, given the overall very low diversity in the SSU rRNA sequences (**Supplementary File 1**), leading to low phylogenetic signal (and therefore low bootstrap support values), these hypotheses are currently tentative. Indeed, the highly supported phylogenomic analysis does not place the strains in the same manner. Further investigation, including more strains and focusing on more variable genes, will be needed in order to better understand the origin of the two different *C. scindens* groups.

### COG distributions of *C. scindens* core, accessory, and unique genes

The distribution of functional genes from the core, accessory, and unique gene families into COG categories shows that the main ones are constituted by genes for storage and information processing (classes J, K, and L), cellular processes and signaling (classes D, M, N, O, T, U, and V), metabolism (classes C, E, F, G, H, I, P and Q), and poorly characterized genes (class S) (**Figure 6a**). Determining the functional classification of the core genome through COG categories is important, as these genes are responsible for the most fundamental biological characteristics. The analysis of core genes shows 39% of genes related to metabolism, a higher percentage compared to the other categories. On the other hand, unique genes show a greater number for the storage and information processing category (34%), followed by unknown or poorly characterized gene function (26%).

**Figure 6.**
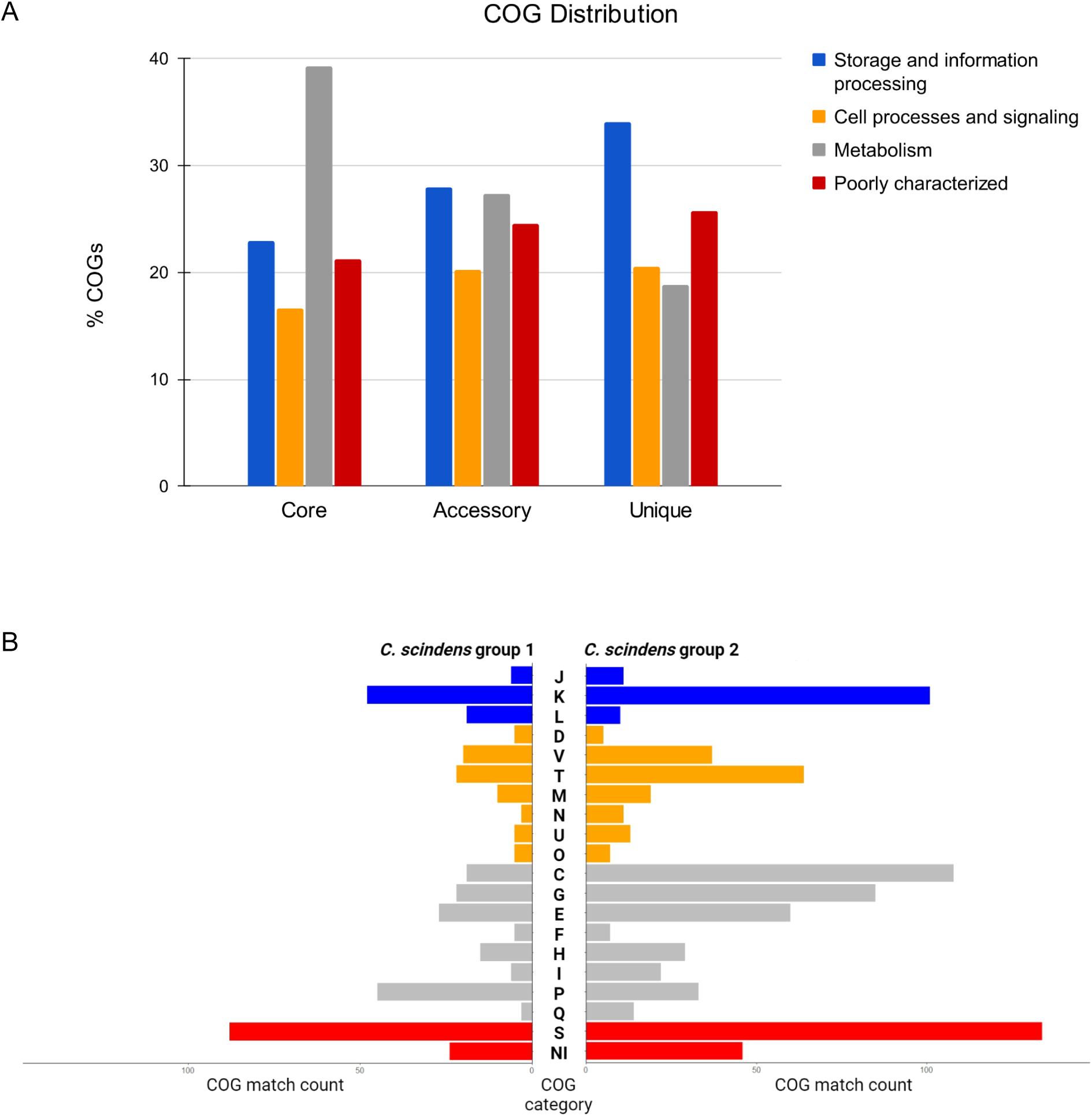
**A.** Distribution of core, accessory and unique genes in clusters of orthologous groups (COG). The distribution of the main functional categories of the pangenome is displayed in the figure. **B**. COG analysis of the accessory genome of Group 1 & Group 2 (see Fig. 4 phylogenomic analysis).

Among the genes associated with metabolism, the COG annotation of the core genes shows that category C (energy production and conversion) is the most common metabolic function (8.5%), followed by categories E (6.6%), and H (6.2%). Category K, belonging to the category of information storage and processing, presented 9.7% of genes in the core genome. The category with the highest number of gene families in the pangenome, as expected from previous studies, is category S (unknown function). The COG annotation of accessory genes and unique genes shows greater numbers in categories K and L (27% and 32%, respectively). Similar results from functional annotation of gene families have been reported in pangenome analyses in other pathogenic and non-pathogenic *Clostridium* species such as *C. perfringens* (Kiu et al. 2017), *C. butyricum* (Zou et al. 2021), and *C. botulinum* (Bhardwaj and Somvanshi 2017).

### KEGG pathway distributions of the C. scindens pangenome

KAAS functional annotation results for the comparative analysis of protein sequences representative of the core, accessory, and unique genome of *C. scindens* against the KEGG database shows a greater gene distribution percentage for metabolism-related pathways (**Supplementary** Figure 5a). This data coincides with our previous result of COG distributions related to metabolic function.

Environmental information processing genes are the second highest category (41%), followed by the genetic information processing category (39%).

In the core genome, carbohydrate metabolism and amino acid metabolism are the two subcategories with the highest number of genes (**Supplementary** Figure 4b), where the pentose-phosphate pathway, and alanine and glycine metabolism pathway collect the highest number of core genes with 18 and 20 genes, respectively. In the accessory and single genome, carbohydrate metabolism, amino acid metabolism and membrane transport are the predominant subcategories. In the second largest main category, processing of environmental information, there is a greater number of accessory and unique genes in the membrane transport subcategory, where the ABC transporter pathway owns the largest number of genes, presenting 73 accessory genes and 42 unique genes.

Furthermore, the KEGG distribution was determined in categories and subcategories of non-core shared genes by the groups of strains identified above as group 1 and group 2. Overall, no full pathways were found to be exclusive to one of the groups, but most genes complement or enrich main core genome metabolic pathways. Group 2 has a higher proportion of genes for metabolism, while group 1 has a higher proportion of genes for the environmental information processing category (**Supplementary** Figure 6a). Additionally, group 1 presented the largest number of genes for the membrane transport subcategory, and group 2, for the carbohydrate metabolism subcategory (**Supplementary** Figure 6b). Quorum sensing proteins seem to be more prevalent in group 1, suggesting more efficiency in cell-cell communication and response to microbial competition.

### Identification of bile acid-metabolizing genes in C. scindens genomes

To determine *bai* and *des* genes presence, BLASTP alignments with an identity greater than 90% and e-value of 0.0 of the 34 *C. scindens* strains and MAG proteomes were considered (**Figure 4**, **Figure 7**). The results suggest that genes located in the bai operon of *C. scindens* are highly conserved in their amino acid sequences among different strains. The *baiJ* and *baiK* genes are not present in the 15 strains in group 1, and the *baiCD* gene was absent in the DFI.4.63 strain from group 2, since this gene is fragmented. This fragmentation could be due to sequencing or assembly errors, but only resequencing of the relevant genomic region in this strain will be able to tell. On the other hand, 10 strains from group 1 feature *desA, desB,* and *desC* genes, while only two strains from group 2 contain these same genes (**Figure 4**).

**Figure 7.**
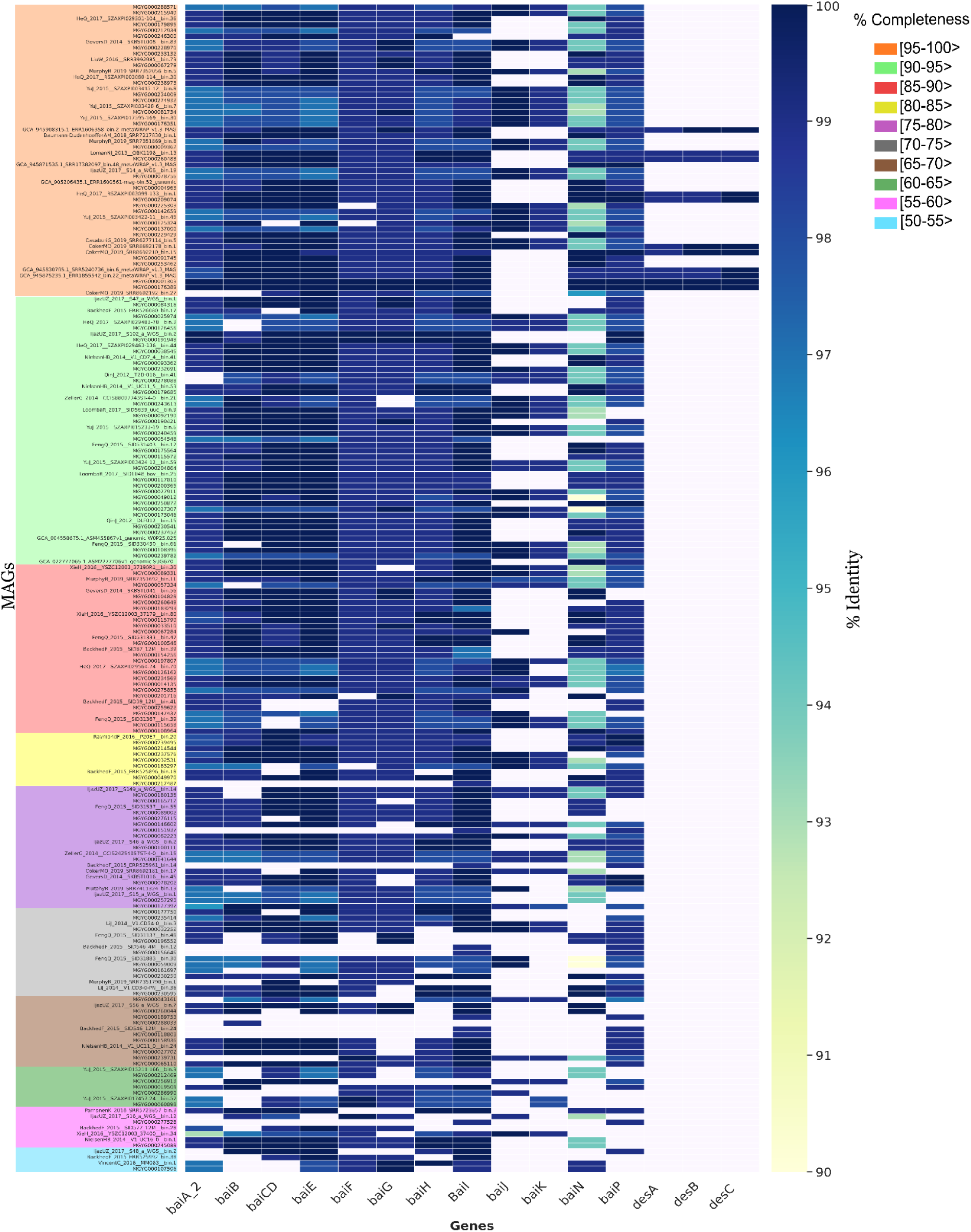
Heatmap of *bai* and *des* genes present in 200 *C. scindens* MAGs. The percentage value of sequence identity is shown in color; the highest value in dark blue and the lowest in light yellow. The absence of the gene is indicated by the color white. The completeness of the genomes is represented in descending order and indicated by the color boxes.

Previously, comparative analysis of the *bai* genes in *Eubacterium* sp. c-25 and other DCA-producing species including *C. scindens* strains ATCC 35704 and G10 (Song et al. 2021) showed that the sequence identity of the predicted Bai proteins from *Eubacterium* is low (43-55%) compared to *C. scindens*, with the exception of BaiH (83%). This suggests that *baiH* has a key role in 7α-dehydroxylation and is a required protein. The 7α-dehydroxylation of primary bile acids is a multistep pathway that involves a group of *bai* genes that encode enzymes that participate in this process (Ridlon et al. 2006). Furthermore, bile acids (BAs) are important due to their relationship with intestinal diseases, such as CRC. The distribution of *bai* genes among *Bacillota* was observed in human gut metagenomes, as determined through a phylogenetic analysis and searches in hidden Markov models, resulting in a greater abundance of *baiCD*, *baiE*, *baiJ* and *baiP* genes in CRC patients compared to healthy subjects (Lee et al. 2022). The diversity of the presence and absence of *bai* and *des* genes in MAGs shows absence of *baiJ* and *baiK* genes in most genomes, with *desA*, *desB* and *desC* genes almost completely absent (**Figure 7**). The genes involved in the biotransformation of BAs in *C. scindens* are the genes of the *baiABCDEFGHI* operon in addition to those of another operon of still unknown function, *baiJKLM*, and the *baiN* gene. *baiN* encodes a protein from the flavoprotein family involved in the BA 7α-dehydroxylation pathway; the sequence of this protein is conserved among 7α-dehydroxylating bacteria closely related to *C. scindens*. The initial characterization of *baiN* in *C. scindens* ATCC 35704 and the presence of the gene in *C. hylemonae* was reported by Harris and colleagues (Harris et al. 2018). We then determined *bai* and *des* gene frequency in 200 MAGs of *C. scindens* (**Figure 7**). The *bai* regulon genes were found in all MAGs, with more gaps as the genome completeness was reduced to ≥ 90%. Interestingly, the *desABC* genes were identified in only 20 *C. scindens* MAGs, while the *desF* gene was more prevalent (97 MAGs).

Vital and collaborators showed the diversity of intestinal bacteria with *bai* genes, using BLAST to search for possible homologues of the *baiN* gene in a set of MAGs of the *Lachnospiraceae* family and in genomes of isolates available in databases, using as a reference the *bai* sequences from *C. scindens* ATCC 35704 (Vital et al. 2019). In that study, *baiN* was identified in most genomes; however, amino acid identities were low even for *C. hiranonis*, a verified 7α-dehydroxylating bacterium that is relatively closely related to *C. scindens* in phylogenetic analyses. Furthermore, the authors propose the absence of genetic sequences homologous to the *baiN* gene outside the main clade of *Lachnospiraceae* that includes *C. scindens* and *C. hylemonae*. By contrast, we located *baiN* in 168 MAGs and all 34 isolated strain genomes (**Figures 4 and 7**).

The *baiJ* and *baiK* genes were discovered in a multigene operon conserved in *C. scindens* VPI 12708 and *C. hylemonae* (Ridlon and Hylemon 2012). The operon encodes a bile acid CoA transferase (*baiK*), and a predicted flavin-dependent oxidoreductase (*baiJ*); however, this pathway is not found in all 7α-dehydroxylating strains of *C. scindens*, as shown in our results (**Figure 4**). The *baiP* gene serves a similar function as *baiJ* in the formation of allo-secondary bile acids (Lee et al. 2022, Meibom et al. 2024). The *baiP* gene was identified in 186 *C. scindens* MAGs, and all 34 isolated strain genomes. These data indicate that *C. scindens* strains are universal in their potential to generate allo-secondary bile acids.

### Predicted Metabolic Pathways in the Core Genome

We previously developed a defined growth medium for *C. scindens* ATCC 35704 using the “leave-one-out” approach to identify essential vitamins and amino acids (Devendran et al. 2019). With this approach, we determined that the amino acid tryptophan and three vitamins (riboflavin, pantothenate, and pyridoxal) were required for the growth of *C. scindens* ATCC 35704. In this medium, *C. scindens* ATCC 35704 fermented glucose mainly to ethanol, acetate, formate, and H_2_. However, it is not clear whether other strains of *C. scindens* have similar nutritional requirements or fermentation end products. We therefore examined the core genome, particularly with respect to predicted amino acid and vitamin metabolism, determining the metabolic pathways located in the core genome of the 34 genomes of cultured *C. scindens* stains (**Figure 8**).

**Figure 8.**
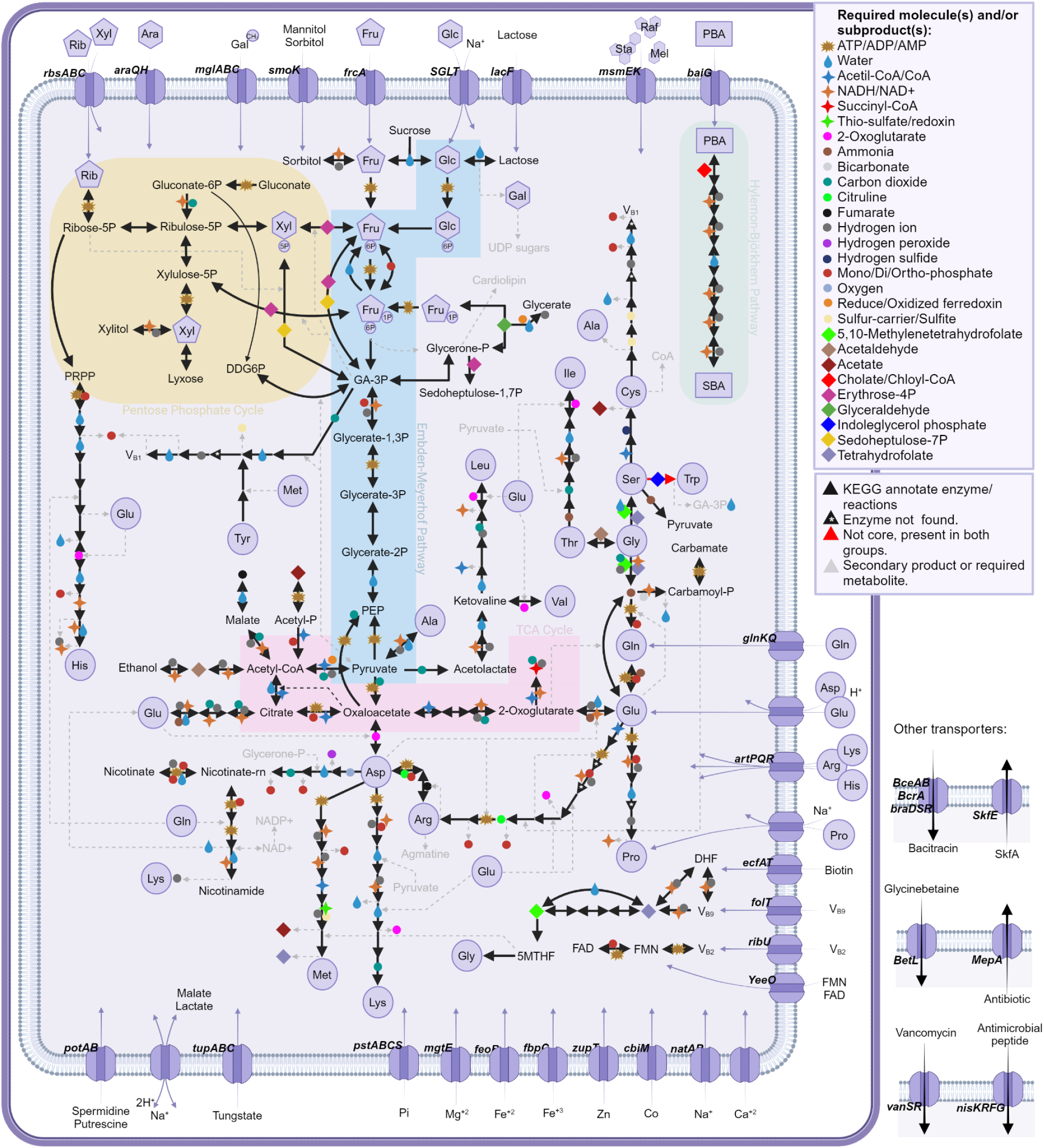
Key metabolic pathways represented in the core genome of *Clostridium scindens*.

*C. scindens* has a complete glycolytic pathway as well as Entner-Doudoroff Pathway, a complete pentose phosphate pathway, and a “horseshoe” TCA cycle from oxaloacetate to succinyl∼SCoA. Oxaloacetate is generated from phosphoenolpyruvate, and malate and fumarate from pyruvate. The complete HB pathway is a core feature of *C. scindens* strains, but steroid-17,20-desmolase, while present in genomes from both groups, is not part of the core genome. The core genome contains complete pathways for the biosynthesis of the majority of amino acids. L-histidine and L-alanine are predicted to be obtained from carnosine metabolism. As part of the accessory genomes of both groups 1 and 2, a tryptophan synthase enzyme (EC 4.2.1.20) was annotated suggesting the possibility that L-tryptophan might be synthesized from indole and L-serine, which is a major product of L-tryptophan catabolism by gut microbiota (Agus et al. 2018). The core and accessory genomes contain the near complete shikimate pathway for the biosynthesis of L-phenylalanine and L-tyrosine. A prolyl aminopeptidase is present in all strains indicating acquisition of proline from peptides; however, a *de novo* pathway for the synthesis of proline is not part of the core genome.

Pantothenate biosynthesis is not part of the core; however, a pathway from pantothenate to CoA was evident in the core genome (**Figure 8**). Synthesis of R-pantothenic acid requires B-alanine, a molecule that seems to be generated by group 1 from L-aspartate, but not by group 2. Pathways from nicotinate to NAD+ and NADP+ from L-aspartate are present. Thiamine and cobalamin biosynthesis pathways are part of the core genome. Genes encoding enzymes involved in one carbon pool by folate are complete. Genes encoding enzymes involved in converting riboflavin to FMN and FAD are present; however, biosynthesis pathways for riboflavin are absent. Folate is required, although the one carbon pool by folate cycle is present in the core genome. The complete biosynthetic pathway for thiamine is present. Lipoate salvage, but not biosynthesis, is present in the core genome. Thus, supplement of our previous defined medium for *C. scindens* ATCC 35704 (Devendran et al. 2019) with a complete set of vitamins (with exception of thiamine) and additional amino acids (tryptophan, proline, phenylalanine, tyrosine) is advisable.

## Discussion

The current study represents the first pangenome analysis of the important bile acid and steroid metabolizing gut microbial species, *Clostridium scindens.* In a phylogenetic analysis of the rRNA SSU sequences and the core genome of members of the *Lachnospiraceae* family, including species identified as *Clostridium* related to the genus *Lachnoclostridium*, it was demonstrated that *C. scindens* was not grouped with other *Clostridium* isolates, despite being classified as a member of this genus; these results showed that all members of the genus *Lachnoclostridium* analyzed formed a distinct monophyletic group, with the exception of *C. scindens*, which is more closely related to isolates of the genus *Dorea*, which includes intestinal commensal species (Sorbara et al. 2020), and has members that also harbor the *bai* operon (Song et al. 2021, Bai et al. 2022, Kim et al. 2022).

We have shown by whole genome phylogeny, gene content, and genome-wide similarity analysis that strains of *C. scindens* actually group into two distinct clades, group 1 (ATCC 35704) and group 2 (VPI 12708), thus supporting Bokkenheuser’s claims of “taxonomic value” for steroid-metabolizing activities and that *Eubacterium* sp. VPI 12708 and *C. scindens* ATCC 35704 would be considered distinct and separate species. *Eubacterium* sp. VPI 12708, isolated in the 1970s, and *C. scindens* ATCC 35704, isolated in 1985, are both known for bile acid dehydroxylation but, unlike *Eubacterium* VPI 12708, *C. scindens* ATCC 35704 is also capable of side-chain cleavage, via steroid-17,20-desmolase, of 17α-hydroxysteroids (e.g., cortisol, tetrahydrocortisol, cortisone, 11-deoxycortisol, 21-deoxycortisol, and 17α-hydroxyprogesterone). Bokkenheuser and associates who isolated *C. scindens* ATCC 35704 argued that steroid-metabolizing activities (e.g, 17, 20-desmolase activity) are species-specific and as such represent distinctive traits that are useful in bacterial identification and taxonomy. However, in 2000, *Eubacterium* VPI 12708 and 5 additional bile acid-dehydroxylating strains were all reclassified as *C. scindens* based on carbohydrate fermentation profiles, 16S rRNA sequencing, and DNA–DNA similarity tests. Steroid-metabolizing activities, other than bile acid dehydroxylation (i.e., presence or absence of *bai* genes), were not considered in their strain reassignment. What the taxonomic designation should be for *Eubacterium* sp. VPI 12708 and related strains (group 2) awaits determination.

Understanding the functional consequences of strain variation in host-associated microbiomes is a current focus in the field, particularly in building microbial consortia for therapeutic application. (Oliveira and Pamer 2023, Louie et al. 2023, Britton and Faith 2021). Bile acid metabolite profiles are important in a myriad of considerations in the germination and vegetative growth of *C. difficile* (Dsouza et al. 2022), immune cell differentiation and function in inflammatory bowel disease (Mohanty et al. 2024, Ridlon and Gaskins 2024), neurodegenerative conditions (Wang et al. 2023), and metabolic disorders (Fuchs and Trauner 2022, Callender et al. 2022). Thus, understanding the depth and breadth of strain variations among potential LBP candidate species is important in advancing microbiome engineering.

## CONCLUSION

Our results reveal that the *C. scindens* pangenome is open, containing, for the set of 34 cultured strains, 12,720 groups of gene families, distributed in 1,630 groups of core genes, 7,051 accessory groups and 4,039 unique strain groups, thus indicating the existence of genomic diversity that could be discovered with additional strains isolation and sequencing. Furthermore, inclusion of 200 *C. scindens* MAGs did not close the pangenome, revealing the high plasticity in the accessory genomes of this species. The inference, by phylogenetic, gene content, and distance methods, of two separate groups of isolates was seen among the 34 cultured strains, which differ by approximately 4-5% in their genomic sequences and in the presence of some genes from the *bai* operon. These two clades might represent species that have diverged very recently or, at the very least, an ongoing speciation process caught on the verge of completion. Functional annotation of the pangenome determined that metabolism is the predominant functional category. In summary, the analysis of the structure and function of the *C. scindens* pangenome provides new data on the genomic content and variability between strains of this species, which will hopefully help to stimulate future research on this important intestinal bile acid-metabolizing bacterium.

## MATERIALS AND METHODS

### Genomic sequences

The sequenced genomes of 34 strains of *C. scindens* were analyzed (**Table 3**). These included: (1) 8 complete genomes (number of contigs = 1) and 2 incomplete genomes (number of contigs = 2 and 5) that were recently published by our group (Olivos Caicedo et al. 2023, Fernandez-Materan et al. 2024); (2) 7 complete genomes (number of contigs = 1) obtained from the public GenBank database at the National Center for Biotechnology Information (NCBI), and (3) 17 incomplete genomes (number of contigs ranging from 21 to 797) obtained from NCBI. In **Table 3**, additional characteristics are described, such as accession number, host, and geographic origin, among others.

**Table 3.**
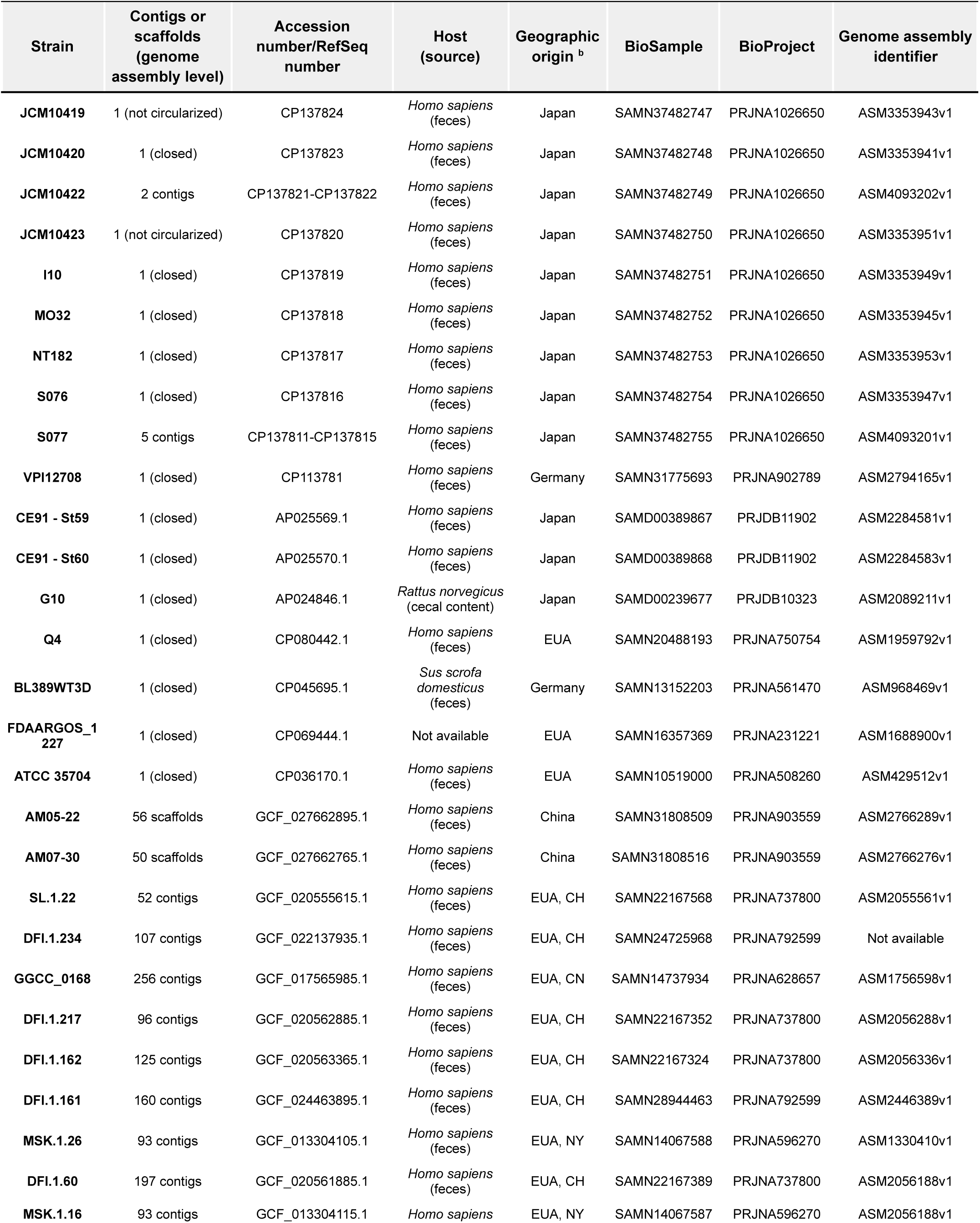
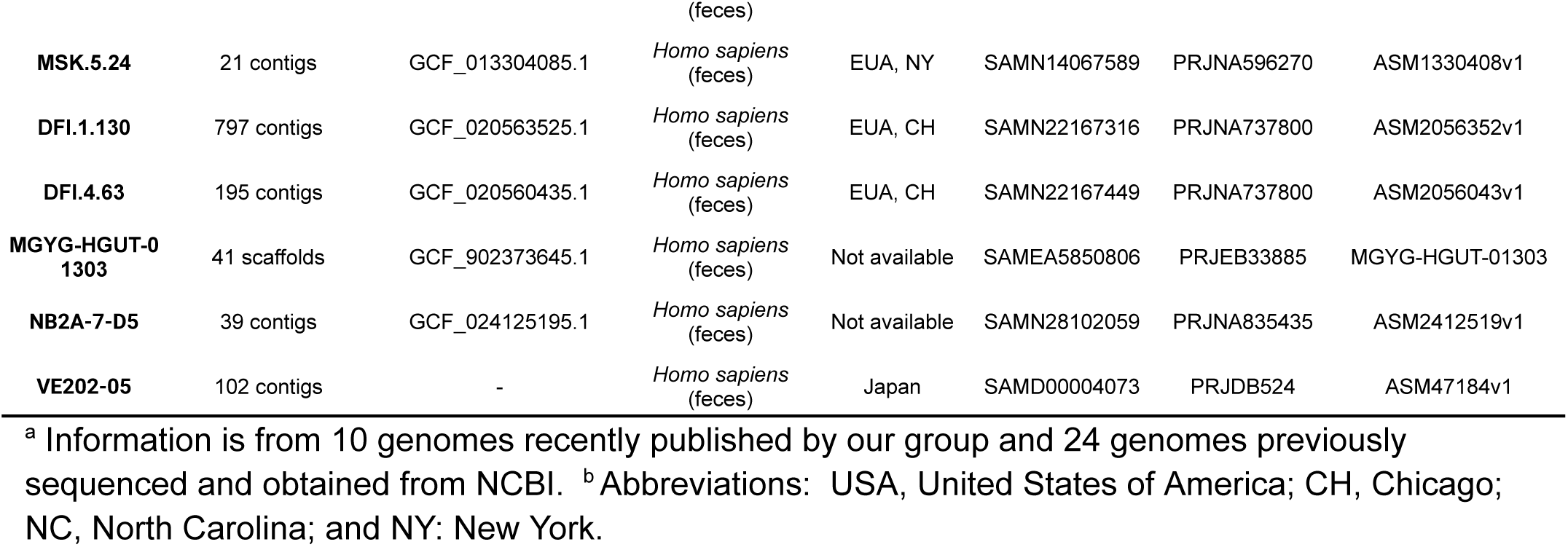
Characteristics and genomic information for 34 cultured strains of *C. scindens*.^a^.

| Strain | Contigs or scaffolds (genome assembly level) | Accession number/RefSeq number | Host (source) | Geographic origin <sup>b</sup> | BioSample | BioProject | Genome assembly identifier |
| --- | --- | --- | --- | --- | --- | --- | --- |
| JCM10419 | 1 (not circularized) | CP137824 | *Homo sapiens* (feces) | Japan | SAMN37482747 | PRJNA1026650 | ASM3353943v1 |
| JCM10420 | 1 (closed) | CP137823 | *Homo sapiens* (feces) | Japan | SAMN37482748 | PRJNA1026650 | ASM3353941v1 |
| JCM10422 | 2 contigs | CP137821-CP137822 | *Homo sapiens* (feces) | Japan | SAMN37482749 | PRJNA1026650 | ASM4093202v1 |
| JCM10423 | 1 (not circularized) | CP137820 | *Homo sapiens* (feces) | Japan | SAMN37482750 | PRJNA1026650 | ASM3353951v1 |
| I10 | 1 (closed) | CP137819 | *Homo sapiens* (feces) | Japan | SAMN37482751 | PRJNA1026650 | ASM3353949v1 |
| MO32 | 1 (closed) | CP137818 | *Homo sapiens* (feces) | Japan | SAMN37482752 | PRJNA1026650 | ASM3353945v1 |
| NT182 | 1 (closed) | CP137817 | *Homo sapiens* (feces) | Japan | SAMN37482753 | PRJNA1026650 | ASM3353953v1 |
| S076 | 1 (closed) | CP137816 | *Homo sapiens* (feces) | Japan | SAMN37482754 | PRJNA1026650 | ASM3353947v1 |
| S077 | 5 contigs | CP137811-CP137815 | *Homo sapiens* (feces) | Japan | SAMN37482755 | PRJNA1026650 | ASM4093201v1 |
| VPI12708 | 1 (closed) | CP113781 | *Homo sapiens* (feces) | Germany | SAMN31775693 | PRJNA902789 | ASM2794165v1 |
| CE91 - St59 | 1 (closed) | AP025569.1 | *Homo sapiens* (feces) | Japan | SAMD00389867 | PRJDB11902 | ASM2284581v1 |
| CE91 - St60 | 1 (closed) | AP025570.1 | *Homo sapiens* (feces) | Japan | SAMD00389868 | PRJDB11902 | ASM2284583v1 |
| G10 | 1 (closed) | AP024846.1 | *Rattus norvegicus* (cecal content) | Japan | SAMD00239677 | PRJDB10323 | ASM2089211v1 |
| Q4 | 1 (closed) | CP080442.1 | *Homo sapiens* (feces) | EUA | SAMN20488193 | PRJNA750754 | ASM1959792v1 |
| BL389WT3D | 1 (closed) | CP045695.1 | *Sus scrofa domestica* (feces) | Germany | SAMN13152203 | PRJNA561470 | ASM968469v1 |
| FDAARGOS_1227 | 1 (closed) | CP069444.1 | Not available | EUA | SAMN16357369 | PRJNA231221 | ASM1688900v1 |
| ATCC 35704 | 1 (closed) | CP036170.1 | *Homo sapiens* (feces) | EUA | SAMN10519000 | PRJNA508260 | ASM429512v1 |
| AM05-22 | 56 scaffolds | GCF_027662895.1 | *Homo sapiens* (feces) | China | SAMN31808509 | PRJNA903559 | ASM2766289v1 |
| AM07-30 | 50 scaffolds | GCF_027662765.1 | *Homo sapiens* (feces) | China | SAMN31808516 | PRJNA903559 | ASM2766276v1 |
| SL.1.22 | 52 contigs | GCF_020555615.1 | *Homo sapiens* (feces) | EUA, CH | SAMN22167568 | PRJNA737800 | ASM2055561v1 |
| DFI.1.234 | 107 contigs | GCF_022137935.1 | *Homo sapiens* (feces) | EUA, CH | SAMN24725968 | PRJNA792599 | Not available |
| GGCC_0168 | 256 contigs | GCF_017565985.1 | *Homo sapiens* (feces) | EUA, CN | SAMN14737934 | PRJNA628657 | ASM1756598v1 |
| DFI.1.217 | 96 contigs | GCF_020562885.1 | *Homo sapiens* (feces) | EUA, CH | SAMN22167352 | PRJNA737800 | ASM2056288v1 |
| DFI.1.162 | 125 contigs | GCF_020563365.1 | *Homo sapiens* (feces) | EUA, CH | SAMN22167324 | PRJNA737800 | ASM2056336v1 |
| DFI.1.161 | 160 contigs | GCF_024463895.1 | *Homo sapiens* (feces) | EUA, CH | SAMN28944463 | PRJNA792599 | ASM2446389v1 |
| MSK.1.26 | 93 contigs | GCF_013304105.1 | *Homo sapiens* (feces) | EUA, NY | SAMN14067588 | PRJNA596270 | ASM1330410v1 |
| DFI.1.60 | 197 contigs | GCF_020561885.1 | *Homo sapiens* (feces) | EUA, CH | SAMN22167389 | PRJNA737800 | ASM2056188v1 |
| MSK.1.16 | 93 contigs | GCF_013304115.1 | *Homo sapiens* (feces) | EUA, NY | SAMN14067587 | PRJNA596270 | ASM2056188v1 |
| **MSK.5.24** | 21 contigs | GCF_013304085.1 | *Homo sapiens*<br>(feces) | EUA, NY | SAMN14067589 | PRJNA596270 | ASM1330408v1 |
| **DFI.1.130** | 797 contigs | GCF_020563525.1 | *Homo sapiens*<br>(feces) | EUA, CH | SAMN22167316 | PRJNA737800 | ASM2056352v1 |
| **DFI.4.63** | 195 contigs | GCF_020560435.1 | *Homo sapiens*<br>(feces) | EUA, CH | SAMN22167449 | PRJNA737800 | ASM2056043v1 |
| **MGYG-HGUT-01303** | 41 scaffolds | GCF_902373645.1 | *Homo sapiens*<br>(feces) | Not available | SAMEA5850806 | PRJEB33885 | MGYG-HGUT-01303 |
| **NB2A-7-D5** | 39 contigs | GCF_024125195.1 | *Homo sapiens*<br>(feces) | Not available | SAMN28102059 | PRJNA835435 | ASM2412519v1 |
| **VE202-05** | 102 contigs | - | *Homo sapiens*<br>(feces) | Japan | SAMD00004073 | PRJDB524 | ASM47184v1 |

<sup>a</sup> Information is from 10 genomes recently published by our group and 24 genomes previously sequenced and obtained from NCBI. <sup>b</sup> Abbreviations: USA, United States of America; CH, Chicago; NC, North Carolina; and NY: New York.

The strains used in the present study were isolated from North America, Europe, and Asia (**Figure 9**), most from human fecal samples of healthy adults, some from unhealthy adults, and two strains isolated from fecal samples from pig and mouse (**Table 3**). The genomes of 200 genomes assembled from metagenomic data (MAGs) of *C. scindens* from the intestinal metagenome of human fecal samples were included (Pasolli et al. 2019, Almeida et al. 2021, Zeng et al. 2022).

**Figure 9.**
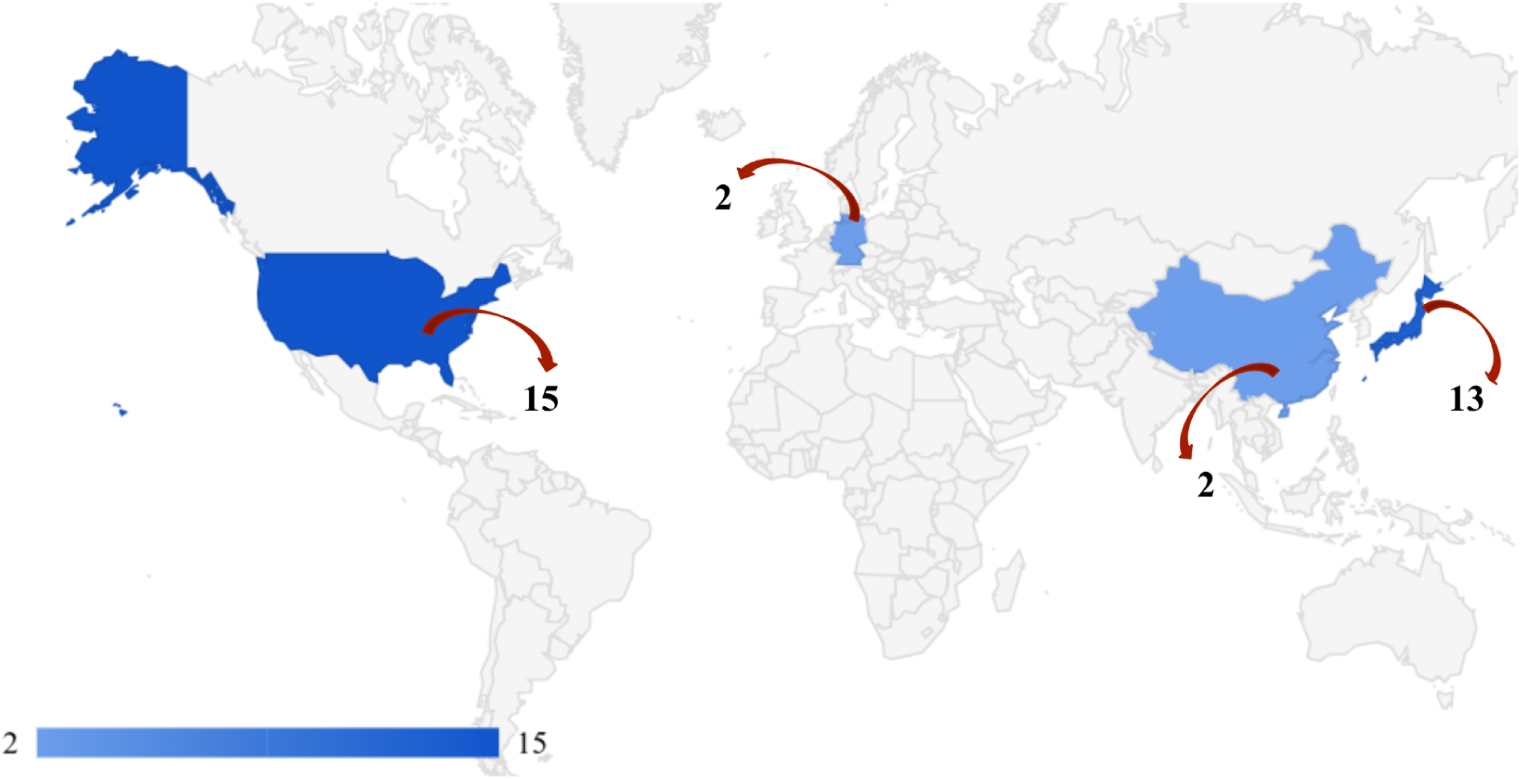
Distribution of the 34 *C. scindens* strains in 4 geographic regions. USA (15), Germany (2), China (2) and Japan (13). Two strains were of unknown geographical origin.

### Genome annotation

The methodological approach is summarized in **Figure 10**. Briefly, sequencing data quality was assessed with the FastQC tool version 0.11.8 (Andrews 2010), and the correction and *de novo* assembling of genomes were done with Unicycler (Wick et al. 2017), Flye (Kolmogorov et al. 2019), and Canu (Koren et al. 2017), resulting in either complete (i.e., circularized) or incomplete (i.e., fragmented into a few contigs) genome assemblies. Evaluation of genome assembly completeness was performed with BUSCO (Benchmarking Universal Single-Copy Ortholog) tool version 5.3.1 (Simão et al. 2015), and the *Clostridium* database of 247 orthologous genes. On the other hand, completeness and contamination of MAGs were assessed with CheckM tool version 1.2.2 (Parks et al. 2015), using a lineage-specific workflow. Functional annotation of the 34 genomes and MAGs of *C. scindens* was performed with Prokka version 1.14 (Seemann 2014), using default parameters values. Prokka performs the prediction of protein-coding genes (CDS), tRNAs, and rRNAs on bacterial, archaeal, and viral genomes, generating individual whole-genome annotation files in a GFF format for each strain, later used in diverse downstream analysis, such as the pangenome analysis detailed below.

**Figure 10.**
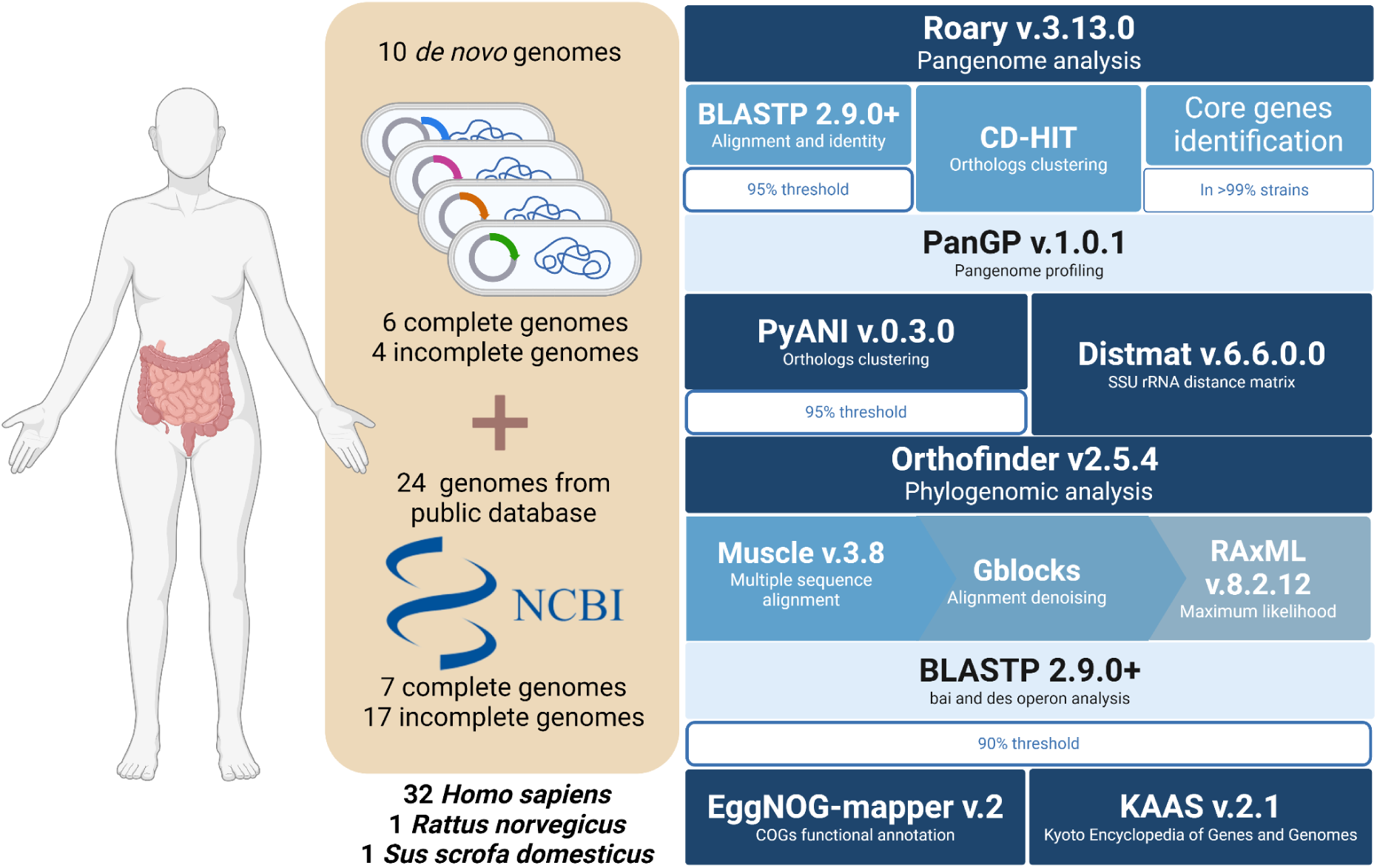
The methodological approach used in this study.

### Metagenome assembled genomes (MAGs) of *C. scindens* from public metagenomes

Genomes from human gut microbiome datasets were downloaded from the following nine different sources: 32,277 genomes from Zeng et al. 2022 (Zeng et al. 2022), 1,200 genomes from Wilkinson et al. 2020 (Wilkinson et al. 2020), 120 genomes classified as *C. scindens* from Almeida et al. 2020 (Almeida et al. 2021), 1,381 genomes from Tamburini et al. 2021 (Tamburini et al. 2022), 154,723 genomes from Pasolli et al. 2019 (Pasolli et al. 2019), 4,997 genomes from Merrill et al. 2022 (Bryan et al. 2022), 2,914 genomes from Lemos et al. 2022 (Lemos et al. 2022), 4,497 genomes from Gounot et al. 2022 (Gounot et al. 2022), and 31 genomes from NCBI. The GTDB-Tk (version 2.1.1) classify workflow (classify_wf) was run on all 202,140 genomes, which resulted in the identification of 224 *C. scindens* genomes, including 200 MAGs.

To generate a list of non-redundant *C. scindens* genomes, we used the program dRep (Olm et al. 2017) on the group of 224 MAGs. The process of dRep includes identifying genomes that are essentially the same and removing all genomes identified as the same except for the best genome that represents that cluster of identical genomes. In this case, the requirement was to have an average nucleotide identity (ANI) of 99% for two genomes to be considered the same.

Custom HMMs were generated for the proteins DesA, DesB, and DesF by using experimentally verified sequences. Briefly, in order to create the protein alignments needed to generate HMM profiles, muscle (5.1.linux64) was used with default parameters to align all amino acid sequences for each of the proteins followed by the hmmbuild function of the HMMER (version 3.3.2) package to generate the HMM profiles. Trusted HMM cutoffs were generated for each of the proteins based on the maximum F-scores based on searches with orthologous proteins. The 224 identified *C. scindens* genomes were translated into amino acid sequences using Prodigal (version 2.6.3). The generated HMM profiles were queried against the amino acid sequences for the 224 genomes using hmmsearch of the HMMER package with the flag --cut_tc.

### Determination of the pangenome of 34 C. scindens strains

The *C. scindens* pangenome and gene content variation of the 34 genomes and 200 MAGs were determined based on the annotation file generated by Prokka and the Roary tool version 3.13.0 (Page et al. 2015). Roary defines clusters of homologous proteins between genomes, thus identifying orthologous genes. To evaluate the MAGs pangenome, percentage values of genome completeness were considered for including a MAG in the analyses, testing completeness values from x to y, increasing by 5% at a time. A 95% identity and 99% definition were used as minimum standard criteria for BLASTP and core genome in this study, allowing Roary to classify genes present in ≥ 99% of the genomes as core genes, genes present in at least two strains as the shell or accessory genes, and genes specific to each strain as the cloud or unique genes.

In general, the roary pipeline iteratively filtered and pre-clustered proteins with CD-Hit (Li & Godzik 2006), then performed an all-against-all comparison using BLASTP, subsequently sequences were clustered with the Markov cluster algorithm (MCL) (Enright et al., 2002) and finally the CD-HIT pre-clustering results were merged together with the MCL results. Moreover, the FastTree tool version 2.1.11 (Price et al. 2010) and a Roary script (roary_plots.py) was used to visualize a presence-absence matrix of core and accessory genes shared between genomes, to determine the variation between the set of strains. Additionally, the Roary binary matrix file for presence-absence of orthologous genes among all strains was selected to estimate the size of the pangenome and core genome using the PanGP version 1.0.1 program (Zhao et al. 2014) with the "totally random sampling" algorithm. Heaps Law’s alpha parameter (to estimate whether the pangenome is closed or open) and core genome predicted size were also estimated using the micropan R package (Snipen and Liland 2015).

### Distance analysis of strains

An average nucleotide identity (ANI) analysis was performed with the PyANI tool version 0.3.0 (Pritchard et al. 2016) with the “-m ANIb” option to indicate genome alignment using the ANIb method with BLAST and to study the variation between the nine newly sequenced genomes and the genomes obtained from NCBI, including *C. hylemonae* species as an outgroup. The suggested threshold percentage of ANI for species identification is greater than or equal to 95% (Richter and Rosselló-Móra 2009).

First, a multiple sequence alignment was performed with Muscle version 3.8 (Edgar 2004). Subsequently, to create the distance matrix, the use method parameter for nucleotide “Uncorrected” for multiple substitutions, was selected. Finally, a table was created with the generated matrix, to be represented by a heat map. Furthermore, to create a distance matrix from multiple alignments of sequences of the SSU rRNA gene copies of all *C. scindens* strains. the Distmat tool (EMBOSS version 6.6.0.0) was used, with a *C. hylemonae* strain included as an outgroup.

### Functional annotation: prediction of COG and KEGG groups

Functional annotations were assigned to proteins using the Clusters of Orthologous Genes (COG) database from the EggNOG-mapper v.2 online tools (Cantalapiedra et al. 2021) (http://eggnog5.embl.de).

The metabolic pathways were studied with the Kyoto Encyclopedia of Genes and Genomes (KEGG) database through the online tool KAAS v.2.1 (KEGG Automatic Annotation Server) (Moriya et al. 2007). Orthologous gene families have been organized into classes and subclasses. Finally, the percentage frequencies of the COG and KEGG categories were calculated for the set of core genes, accessory genes, and unique genes.

### Identification of bai and des genes in C. scindens genomes

To identify the presence of genes from the *bai* operon (bile acid-inducible) and genes from the desABC operon (cortisol-inducible) in the 34 genomes and 200 MAGs of *C. scindens*, a similarity analysis was performed with the BLASTP 2.9.0+ tool (Altschul et al. 1997), using as queries *bai* and *des* genes from *C. scindens* ATCC 35704 and *C. scindens* VPI 12708 **(Supplementary Table 2),** and the proteome database of the analyzed strains as a database with a maximum e-value allowed set at 1e-20. Amino acid sequences with a similarity greater than 90% were considered as best matches. The sequences analyzed belong to the genes *baiA_2, baiB, baiCD, baiE, baiF, baiG, baiH, baiI, baiJ, baiK, baiN,* and *baiP,* and to the genes *desA, desB,* and *desC* obtained from *C. scindens* ATCC 35704 or *C. scindens* VPI 12708.

### Phylogenomic analysis

The Orthofinder v2.5.4 program (Emms and Kelly 2015) was used to analyze the 34 cultured *C. scindens* strains and *C. hylemonae* proteomes. All protein sequences from orthogroups of interest were aligned with the Muscle program v.3.8. Gblocks (Castresana 2000) was used to remove ambiguously aligned regions, as these regions decrease the quality of phylogenetic inference. Default parameters were used, except for keeping columns where gaps occurred in up to half of the sequences in the alignment.

After eliminating ambiguous positions, alignments smaller than 50% of the original size were discarded. Finally, FASconCaT v1.04 (Kück and Longo 2014) was used to concatenate all resulting alignments and create a supermatrix for phylogenetic inference analysis. The tree was generated applying the maximum likelihood method, with RAxML version 8.2.12 (Stamatakis 2014) and was drawn and edited manually with the online tool iTOL (itol.embl.de) and Inkscape (www.inkscape.org).

## Acknowledgements

This work was supported by NIH R03AI147127 (J.M.R; J.M.P.A), RO1 CA204808-01 (J.M.R), R01 GM134423-01A1 (J.M.R.), R01 GM145920-01 (J.M.R), and a Fulbright Scholarship (F.F.). K.Y.O.C. was supported by a Coordenação de Aperfeiçoamento de Pessoal de Nível Superior (CAPES) fellowship.

## Extended Data Figures and Tables

**Supplementary Figure 1.**
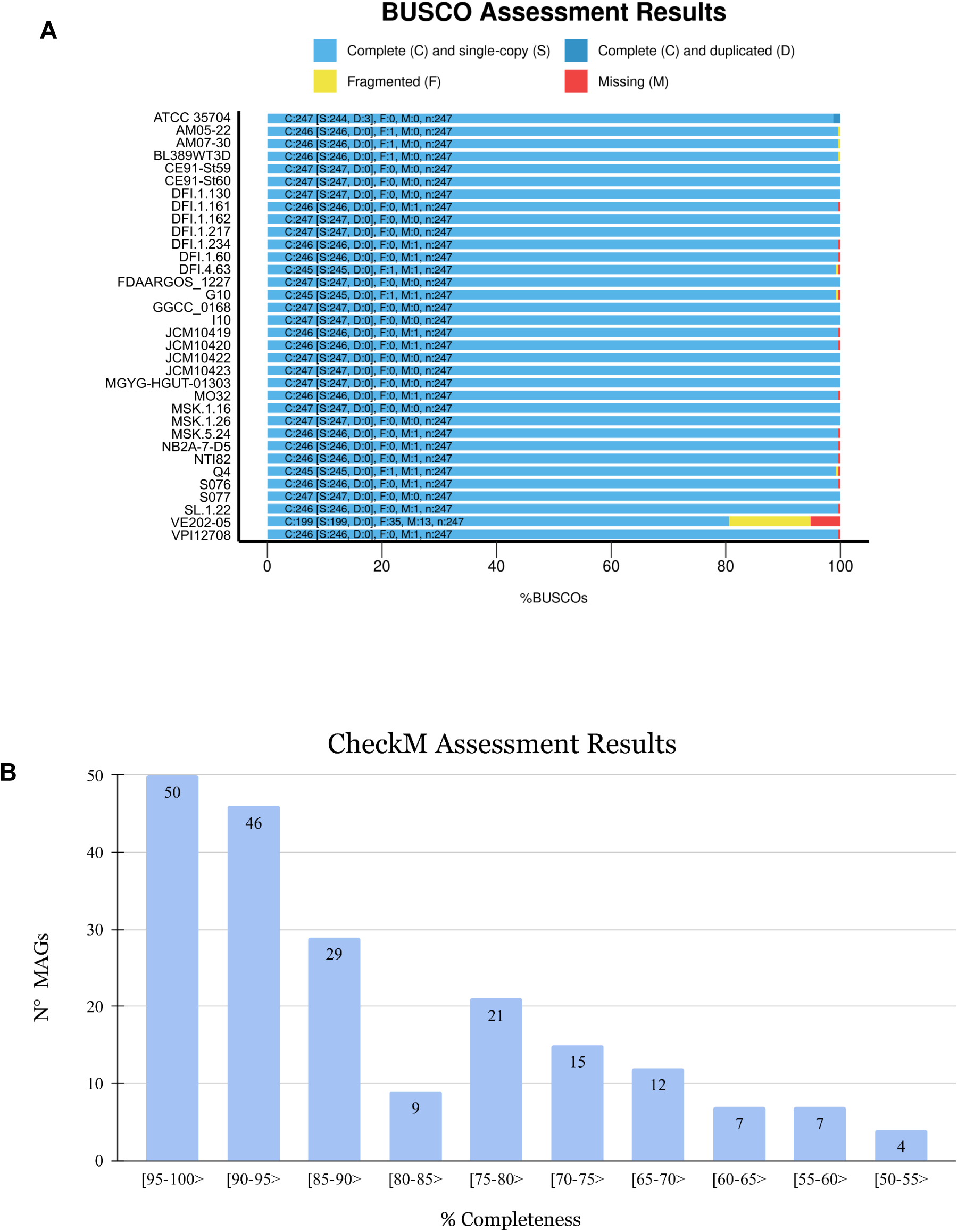
Completeness assessment of genome assembly. **A.** BUSCO results for 34 *C. scindens* strains, including the genomes recently published by our group and the complete and incomplete genomes from NCBI. **B.** CheckM results for 200 *C. scindens* MAGs.

**Supplementary Figure 2.**
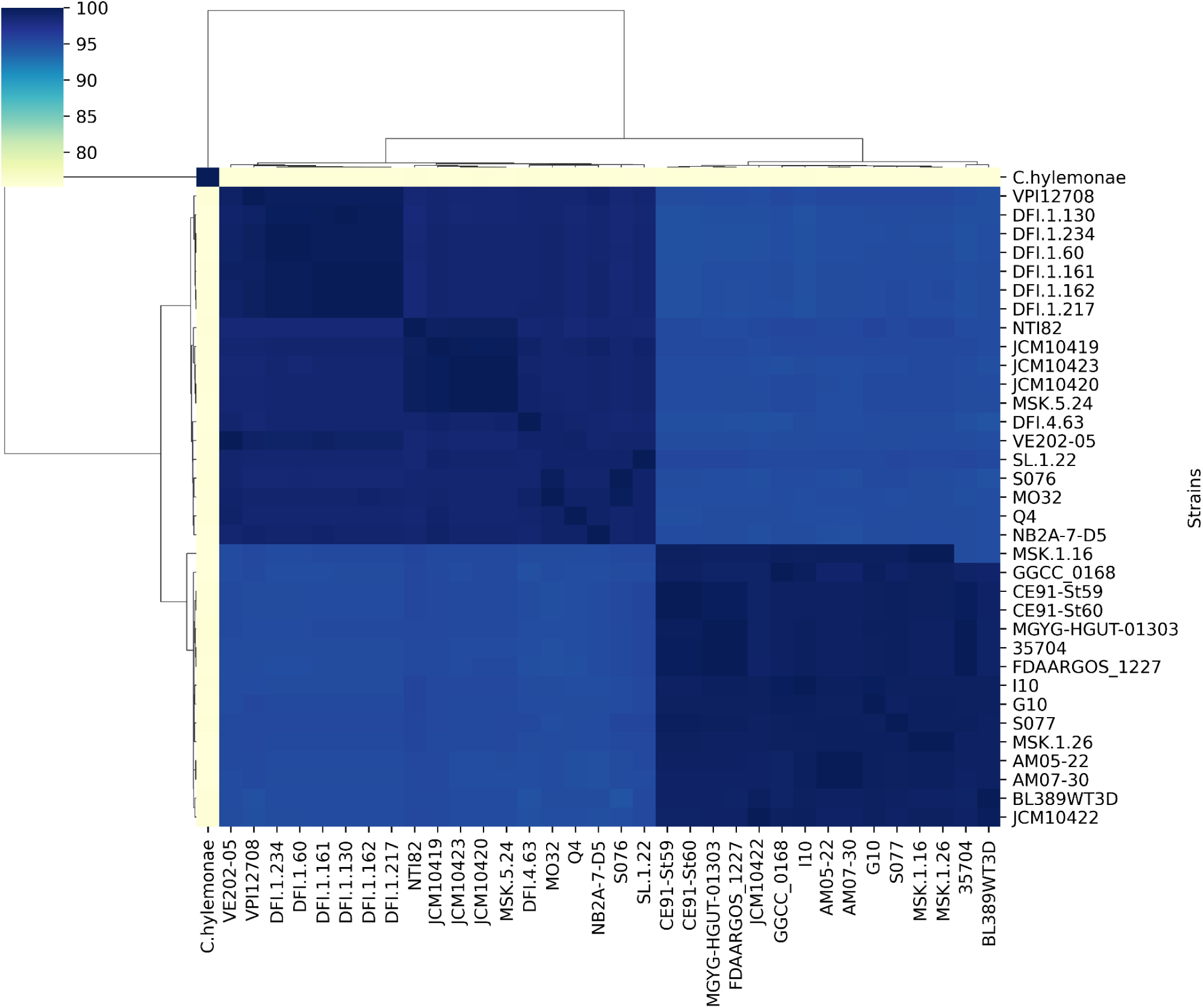
ANI analysis between the *C. scindens* genomes and the *C. hylemonae* genome. The graph shows the formation of two sets of strains with a difference of around 4-5% in their genomic sequences, represented by color intensities. One group includes 15 strains and the other 19.

**Supplementary Figure 3.**
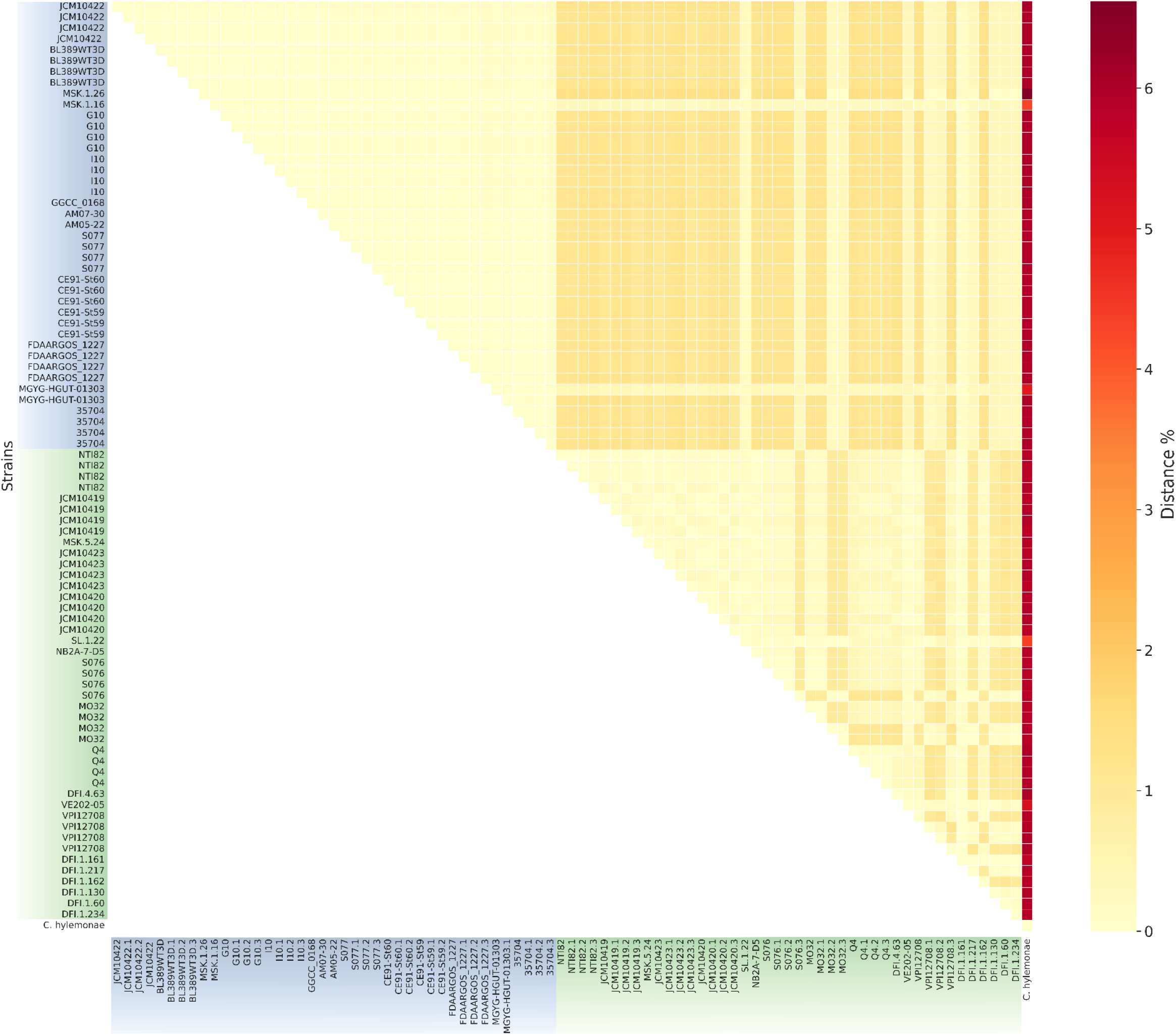
Distance matrix based on the SSU rRNA gene copies of the 34 *C. scindens* strains. The species *C. hylemonae* was included as an outgroup. The distance value is shown in color; the highest value (6) in red and the lowest (0) in yellow. Strains from groups 1 and 2 are indicated by blue and green boxes respectively. The numbers used to color the matrix are presented in **Supplementary File 2**.

**Supplementary Figure 4.**
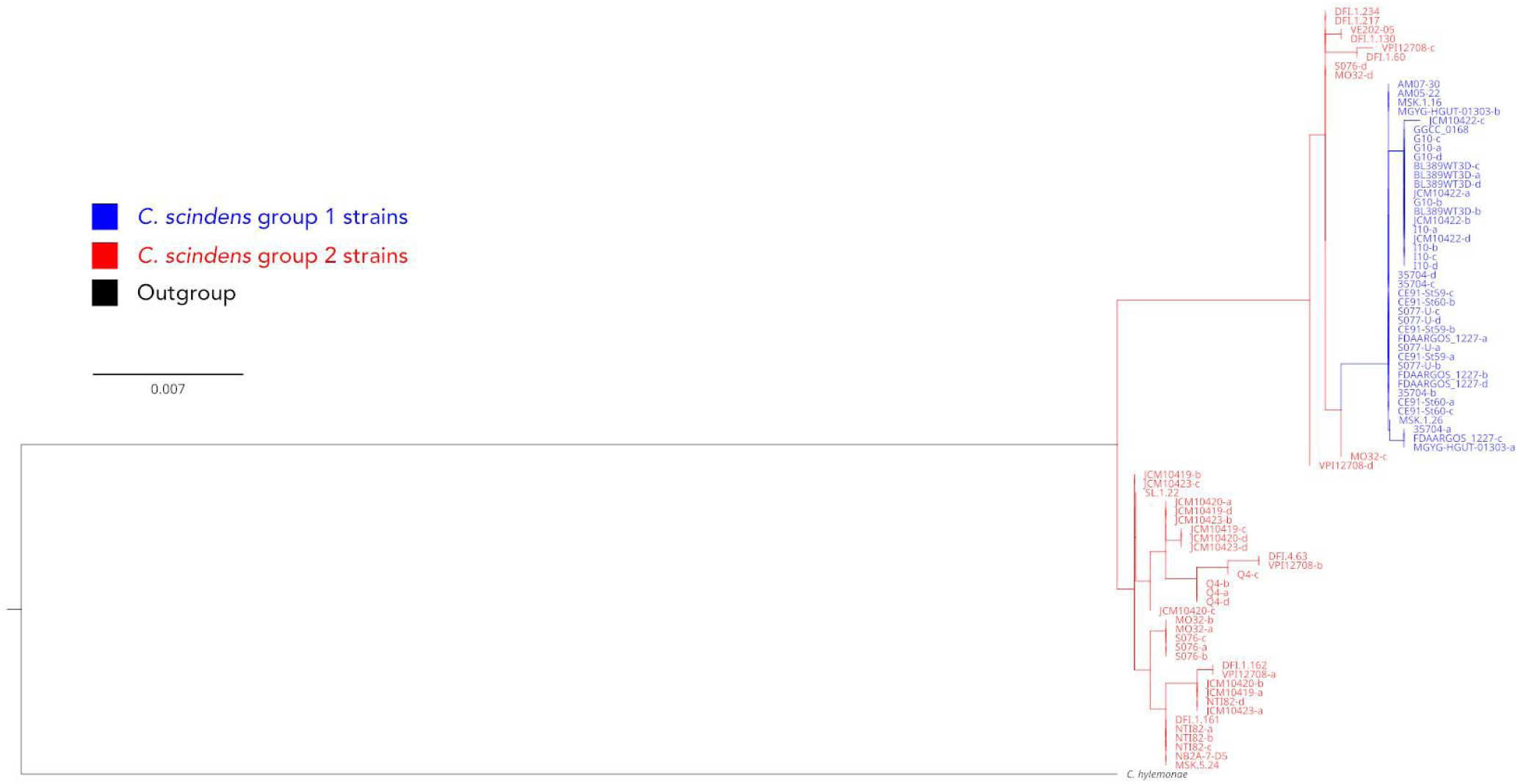
Maximum likelihood phylogenetic tree of all SSU rRNA genes identified from 34 *C. scindens* genomes, with *C. hylemonae* as the outgroup. Bootstrap values were all below 50 and are therefore not shown. Strain grouping indicated by colors, with group 1 strains in blue and group 2 strains in red.

**Supplementary Figure 5.**
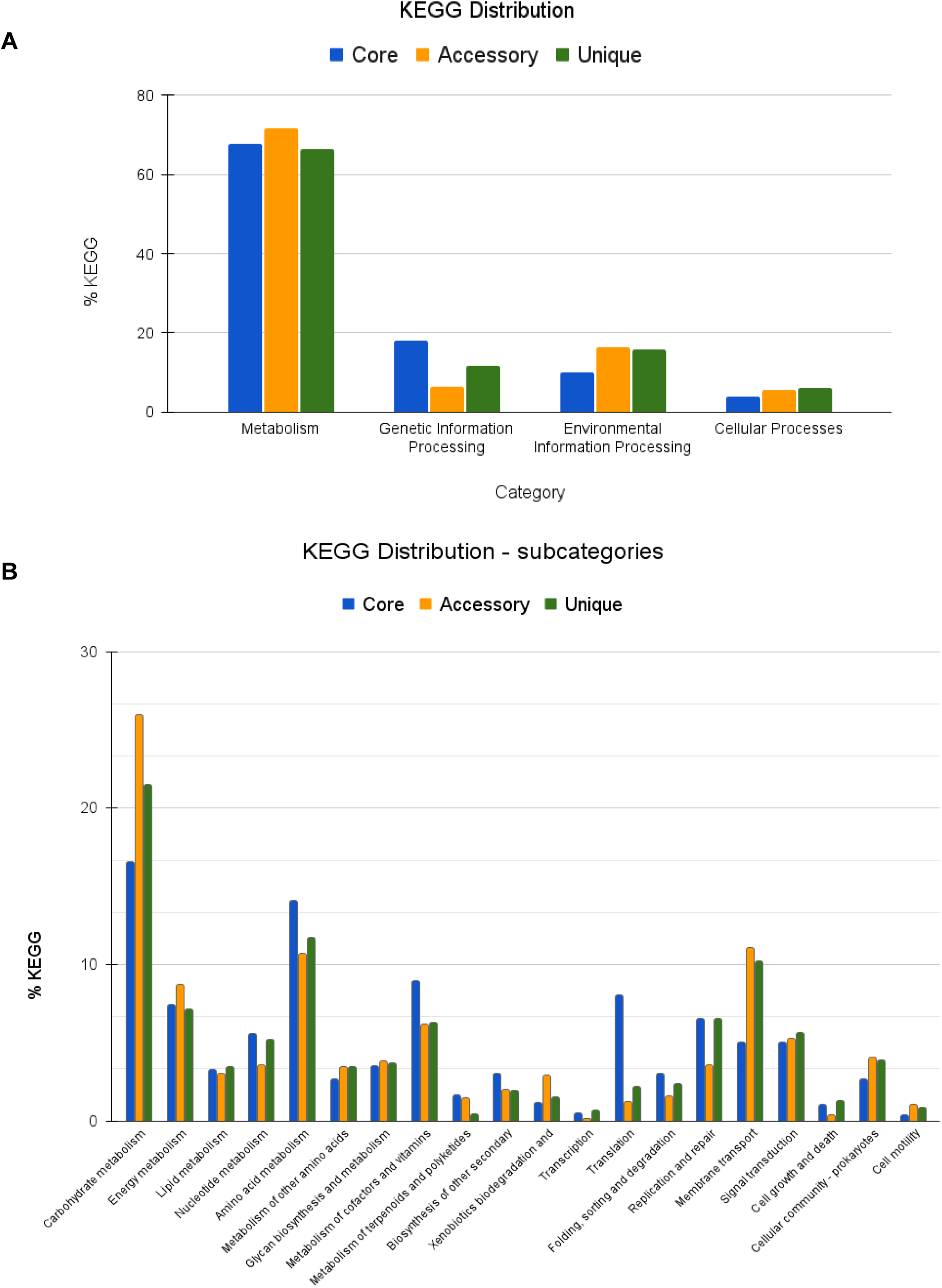
KEGG distribution of core, accessory and unique genes of the *C. scindens* pangenome. **A.** Distribution in main categories. **B.** Distribution into subcategories.

**Supplementary Figure 6.**
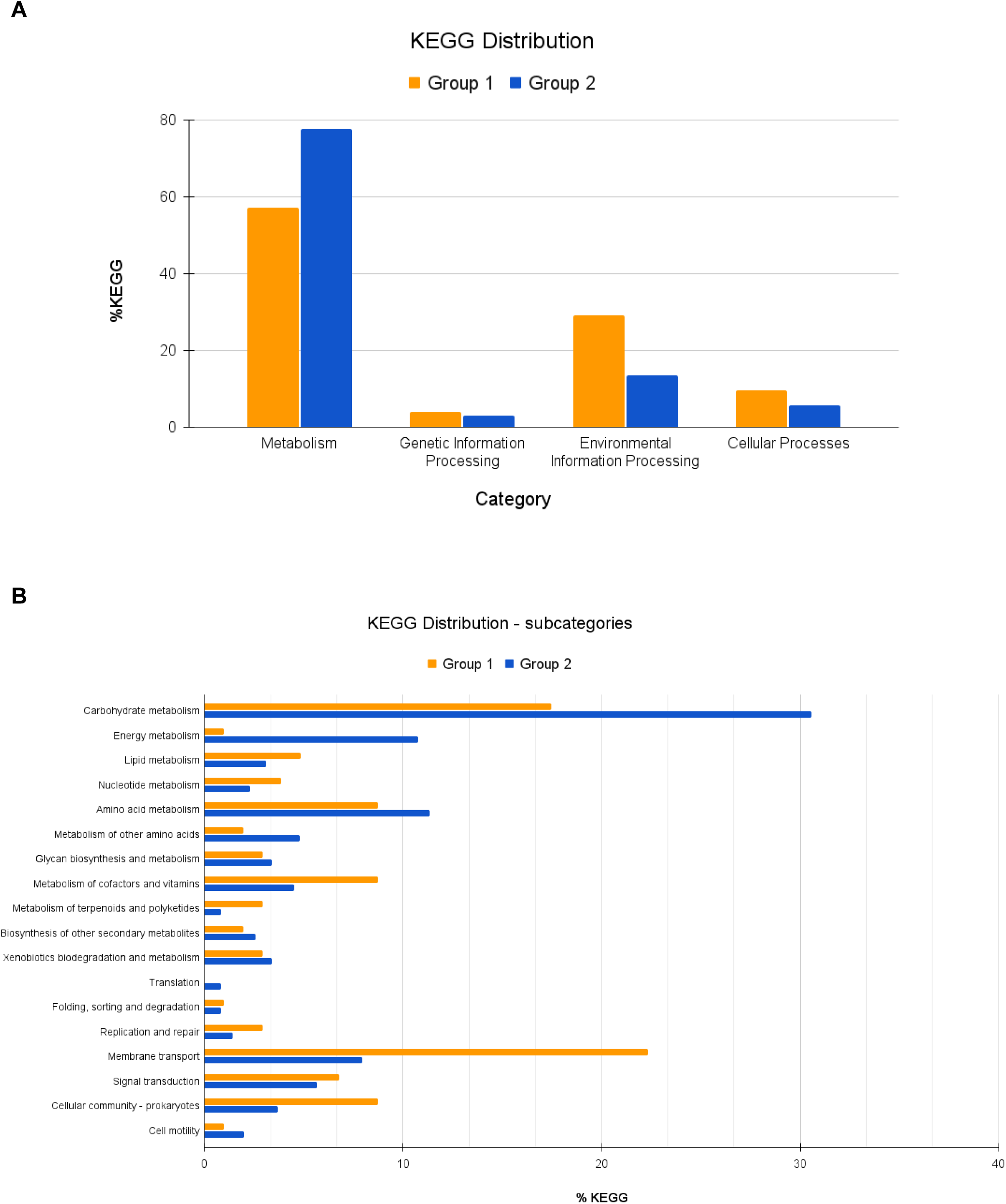
KEGG distribution of the groups of *C. scindens* strains, Group 1 and Group 2. **A.** Distribution in main categories. **B.** Distribution into subcategories.

**Supplementary table 1.** Genomic characteristics of 200 non-dereplicated *C. scindens* MAGs. Genome completeness values are shown in descending order.

| Genome ID | Completeness % | Contamination % | Genome size (bp) | N° scaffolds | N° contigs | N50 (scaffolds) | N50 (contigs) | N° predicted genes |
| --- | --- | --- | --- | --- | --- | --- | --- | --- |
| GCA_945830785.1_SRR5240736_bin.6_metaWRAP_v1.3_MAG_genomic | 99.42 | 0.00 | 3,099,469 | 82 | 82 | 87,848 | 87,848 | 3,060 |
| GCA_945875235.1_ERR1855542_bin.22_metaWRAP_v1.3_MAG_genomic | 99.42 | 0.00 | 3,127,405 | 84 | 84 | 87,848 | 87,848 | 3,086 |
| MGYG000001303 | 99.42 | 0.00 | 3,622,605 | 41 | 68 | 158,586 | 157,623 | 3,602 |
| MGYG000176389 | 99.42 | 0.00 | 3,619,905 | 68 | 68 | 157,623 | 157,623 | 3,603 |
| CokerMO_2019_SRR8692192_bin.27 | 99.42 | 0.00 | 2,981,811 | 74 | 74 | 68,939 | 68,939 | 2,913 |
| MGYG000253462 | 99.22 | 0.00 | 3,336,973 | 64 | 64 | 107,968 | 107,968 | 3,374 |
| MGYG000091745 | 99.22 | 0.58 | 3,339,557 | 51 | 64 | 165,625 | 107,968 | 3,364 |
| YuJ_2015__SZAXPI003422-11__bin.45 | 98.83 | 0.00 | 3,780,218 | 44 | 44 | 140,415 | 140,415 | 3,568 |
| MGYG000137000 | 98.83 | 0.19 | 3,780,218 | 44 | 44 | 140,415 | 140,415 | 3,568 |
| CasaburiG_2019_SRR6277114_bin.5 | 98.83 | 0.19 | 3,885,654 | 21 | 21 | 251,049 | 251,049 | 3,672 |
| MGYG000125324 | 98.83 | 0.00 | 4,056,916 | 17 | 25 | 546,338 | 291,830 | 3,836 |
| CokerMO_2019_SRR8692178_bin.1 | 98.83 | 0.19 | 3,795,272 | 22 | 22 | 290,238 | 290,238 | 3,602 |
| MGYG000229429 | 98.83 | 0.00 | 3,131,882 | 57 | 57 | 159,529 | 159,529 | 3,133 |
| CokerMO_2019_SRR8692210_bin.15 | 98.83 | 0.00 | 2,953,419 | 70 | 70 | 82,650 | 82,650 | 2,886 |
| MGYG000142659 | 98.80 | 0.00 | 3,676,932 | 23 | 23 | 249,655 | 249,655 | 3,426 |
| MGYG000225303 | 98.71 | 0.00 | 4,059,414 | 20 | 31 | 545,924 | 265,229 | 3,838 |
| HeQ_2017__RSZAXPI003099-133__bin.1 | 98.45 | 0.00 | 3,090,863 | 48 | 48 | 138,734 | 138,734 | 3,042 |
| MGYG000209074 | 98.45 | 0.00 | 3,090,863 | 48 | 48 | 138,734 | 138,734 | 3,042 |
| GCA_905206435.1_ERR1600561-mag-bin.52_genomic | 98.25 | 0.88 | 2,985,519 | 63 | 63 | 85,679 | 85,679 | 2,985 |
| MGYG000004963 | 98.25 | 0.88 | 2,985,519 | 63 | 63 | 85,679 | 85,679 | 2,985 |
| IjazUZ_2017__S14_a_WGS__bin.19 | 98.11 | 0.12 | 3,562,888 | 211 | 211 | 23,628 | 23,628 | 3,489 |
| MGYG000078756 | 98.11 | 0.12 | 3,562,888 | 211 | 211 | 23,628 | 23,628 | 3,489 |
| GCA_945871535.1_SRR17382097_bin.48__metaWRAP_v1.3_MAG_genomic | 97.66 | 0.58 | 2,878,239 | 103 | 103 | 50,587 | 50,587 | 2,896 |
| LomanNJ_2013__OBK1196__bin.13 | 97.40 | 0.58 | 2,608,426 | 185 | 185 | 21,622 | 21,622 | 2,621 |
| MGYG000260488 | 97.40 | 0.00 | 2,608,426 | 185 | 185 | 21,622 | 21,622 | 2,621 |
| MGYG000009362 | 97.37 | 0.00 | 3,915,227 | 26 | 26 | 297,304 | 297,304 | 3,882 |
| MurphyR_2019_SRR7351869_bin.8 | 97.08 | 0.00 | 3,771,567 | 53 | 53 | 112,091 | 112,091 | 3,608 |
| GCA_945908315.1_ERR1606358_bin.2__metaWRAP_v1.3_MAG_genomic | 97.08 | 0.00 | 2,946,002 | 57 | 57 | 93,160 | 93,160 | 2,902 |
| Baumann-DudenhoefferAM_2018_SRR7217830_bin.1 | 97.08 | 0.00 | 2,890,901 | 68 | 68 | 79,872 | 79,872 | 2,873 |
| YuJ_2015__SZAXPI017595-169__bin.30 | 97.05 | 0.00 | 3,673,145 | 34 | 34 | 176,479 | 176,479 | 3,423 |
| MGYG000176351 | 97.05 | 0.58 | 3,673,145 | 34 | 34 | 176,479 | 176,479 | 3,423 |
| YuJ_2015__SZAXPI003428-6__bin.7 | 96.88 | 0.00 | 3,586,629 | 151 | 151 | 35,075 | 35,075 | 3,455 |
| MGYG000081734 | 96.88 | 0.00 | 3,586,629 | 151 | 151 | 35,075 | 35,075 | 3,455 |
| YuJ_2015__SZAXPI003415-12__bin.8 | 96.72 | 0.22 | 3,591,634 | 250 | 250 | 24,910 | 24,910 | 3,589 |
| MGYG000234009 | 96.72 | 0.22 | 3,591,634 | 250 | 250 | 24,910 | 24,910 | 3,589 |
| MGYG000274932 | 96.72 | 2.24 | 3,657,773 | 35 | 35 | 160,445 | 160,445 | 3,443 |
| HeQ_2017__RSZAXPI003080-114__bin.30 | 96.67 | 2.24 | 2,763,047 | 80 | 80 | 55,843 | 55,843 | 2,747 |
| MGYG000238973 | 96.67 | 0.00 | 2,763,047 | 80 | 80 | 55,843 | 55,843 | 2,747 |
| LiuW_2016__SRR3992985__bin.73 | 96.11 | 0.39 | 2,560,880 | 200 | 200 | 17,515 | 17,515 | 2,523 |
| MGYG000067279 | 96.11 | 0.39 | 2,560,880 | 200 | 200 | 17,515 | 17,515 | 2,523 |
| MurphyR_2019_SRR7352056__bin.5 | 96.11 | 1.95 | 3,554,900 | 152 | 152 | 40,200 | 40,200 | 3,401 |
| MGYG000233132 | 96.01 | 1.95 | 2,960,856 | 79 | 79 | 66,313 | 66,313 | 2,976 |
| GeversD_2014__SKBSTL008__bin.83 | 96.00 | 0.97 | 3,536,107 | 182 | 182 | 26,649 | 26,649 | 3,421 |
| MGYG000228970 | 96.00 | 1.75 | 3,536,107 | 182 | 182 | 26,649 | 26,649 | 3,421 |
| MGYG000246300 | 95.81 | 0.00 | 2,569,623 | 357 | 357 | 10,250 | 10,250 | 2,772 |
| MGYG000212934 | 95.76 | 0.00 | 3,362,959 | 282 | 282 | 16,032 | 16,032 | 3,379 |
| HeQ_2017__SZAXPI029501-104__bin.36 | 95.75 | 1.85 | 3,082,792 | 43 | 43 | 152,185 | 152,185 | 3,056 |
| MGYG000179895 | 95.75 | 0.99 | 3,082,792 | 43 | 43 | 152,185 | 152,185 | 3,056 |
| MGYG000215940 | 95.42 | 0.00 | 3,594,021 | 92 | 92 | 62,308 | 62,308 | 3,379 |
| MGYG000288571 | 95.32 | 0.00 | 3,465,768 | 170 | 170 | 29,462 | 29,462 | 3,361 |
| GCA_022777065.1_ASM2277706v1_genomic | 94.93 | 1.56 | 2,827,450 | 33 | 33 | 151,255 | 151,255 | 2,810 |
| MGYG000239782 | 94.86 | 0.12 | 3,590,894 | 82 | 82 | 89,011 | 89,011 | 3,412 |
| GCA_004558675.1_ASM455867v1_genomic | 94.74 | 0.88 | 2,885,217 | 99 | 99 | 48,377 | 48,377 | 2,929 |
| FengQ_2015__SID530450__bin.66 | 94.74 | 0.29 | 3,296,439 | 44 | 44 | 147,477 | 147,477 | 3,090 |
| MGYG000108396 | 94.74 | 0.29 | 3,296,439 | 44 | 44 | 147,477 | 147,477 | 3,090 |
| MGYG000237452 | 94.46 | 4.18 | 2,760,783 | 82 | 82 | 53,616 | 53,616 | 2,732 |
| QinJ_2012__DLF012__bin.15 | 94.45 | 0.29 | 2,523,043 | 260 | 260 | 14,037 | 14,037 | 2,645 |
| MGYG000230541 | 94.45 | 0.29 | 2,523,043 | 260 | 260 | 14,037 | 14,037 | 2,645 |
| MGYG000173046 | 94.15 | 0.00 | 3,464,751 | 38 | 38 | 164,863 | 164,863 | 3,228 |
| MGYG000027307 | 94.13 | 0.29 | 3,912,387 | 102 | 102 | 72,302 | 72,302 | 4,593 |
| MGYG000250872 | 93.97 | 1.07 | 2,813,650 | 49 | 49 | 92,921 | 92,921 | 2,770 |
| MGYG000049012 | 93.70 | 0.44 | 3,426,874 | 286 | 286 | 18,132 | 18,132 | 3,408 |
| MGYG000027911 | 93.65 | 2.19 | 3,456,019 | 320 | 320 | 14,732 | 14,732 | 3,256 |
| MGYG000117810 | 93.54 | 0.00 | 2,698,782 | 35 | 35 | 132,487 | 132,487 | 2,622 |
| LoombaR_2017__SID1048_bav__bin.25 | 93.54 | 0.00 | 2,746,452 | 37 | 37 | 132,487 | 132,487 | 2,661 |
| MGYG000200365 | 93.54 | 0.00 | 2,746,452 | 37 | 37 | 132,487 | 132,487 | 2,661 |
| YuJ_2015__SZAXPI003424-12__bin.59 | 93.45 | 0.00 | 3,453,537 | 251 | 251 | 21,299 | 21,299 | 3,393 |
| MGYG000204864 | 93.45 | 0.00 | 3,453,537 | 251 | 251 | 21,299 | 21,299 | 3,393 |
| MGYG000115572 | 93.28 | 0.29 | 2,494,484 | 244 | 244 | 14,183 | 14,183 | 2,605 |
| FengQ_2015__SID531403__bin.12 | 93.04 | 0.58 | 2,718,000 | 270 | 270 | 14,354 | 14,354 | 2,902 |
| MGYG000175564 | 93.04 | 0.58 | 2,718,000 | 270 | 270 | 14,354 | 14,354 | 2,902 |
| MGYG000054548 | 92.90 | 1.42 | 3,104,168 | 528 | 528 | 8,293 | 8,293 | 3,332 |
| YuJ_2015__SZAXPI015233-19__bin.6 | 92.89 | 0.08 | 3,493,459 | 72 | 72 | 71,570 | 71,570 | 3,271 |
| MGYG000240459 | 92.89 | 0.08 | 3,493,459 | 72 | 72 | 71,570 | 71,570 | 3,271 |
| MGYG000190421 | 92.85 | 0.88 | 2,453,338 | 334 | 334 | 10,302 | 10,302 | 2,639 |
| LoombaR_2017__SID5639_uuc__bin.9 | 92.79 | 0.97 | 3,618,456 | 166 | 166 | 31,893 | 31,893 | 3,550 |
| MGYG000092190 | 92.79 | 0.97 | 3,618,456 | 166 | 166 | 31,893 | 31,893 | 3,550 |
| ZellerG_2014__CCIS88007743ST-4-0__bin.21 | 92.69 | 0.70 | 3,379,725 | 146 | 146 | 36,321 | 36,321 | 3,257 |
| MGYG000243613 | 92.69 | 0.70 | 3,379,725 | 146 | 146 | 36,321 | 36,321 | 3,257 |
| NielsenHB_2014__V1_UC11_5__bin.53 | 92.67 | 0.00 | 2,786,591 | 116 | 116 | 37,909 | 37,909 | 2,838 |
| MGYG000179685 | 92.67 | 0.00 | 2,786,591 | 116 | 116 | 37,909 | 37,909 | 2,838 |
| QinJ_2012__T2D-016__bin.41 | 92.23 | 0.88 | 3,334,283 | 447 | 447 | 9,994 | 9,994 | 3,473 |
| MGYG000278088 | 92.23 | 0.88 | 3,334,283 | 447 | 447 | 9,994 | 9,994 | 3,473 |
| MGYG000232691 | 91.93 | 1.46 | 3,524,914 | 100 | 100 | 5,6167 | 56,167 | 3,339 |
| NielsenHB_2014__V1_CD7_4__bin.41 | 91.81 | 0.00 | 2,729,357 | 47 | 47 | 110,798 | 110,798 | 2,685 |
| MGYG000093362 | 91.81 | 0.00 | 2,729,357 | 47 | 47 | 110,798 | 110,798 | 2,685 |
| HeQ_2017__SZAXPI029463-136__bin.44 | 91.40 | 0.58 | 3,490,822 | 47 | 47 | 111,242 | 111,242 | 3,270 |
| MGYG000038545 | 91.40 | 0.58 | 3,490,822 | 47 | 47 | 111,242 | 111,242 | 3,270 |
| IjazUZ_2017__S102_a_WGS__bin.2 | 90.74 | 2.52 | 2,389,371 | 323 | 323 | 10,060 | 10,060 | 2,601 |
| MGYG000191948 | 90.74 | 2.52 | 2,389,371 | 323 | 323 | 10,060 | 10,060 | 2,601 |
| HeQ_2017__SZAXPI029483-78__bin.3 | 90.60 | 0.16 | 3,375,410 | 285 | 285 | 17,693 | 17,693 | 3,371 |
| MGYG000126456 | 90.60 | 0.16 | 3,375,410 | 285 | 285 | 17,693 | 17,693 | 3,371 |
| MGYG000025974 | 90.35 | 1.95 | 3,549,982 | 45 | 45 | 118,974 | 118,974 | 3,291 |
| BackhedF_2015_ERR526080_bin.17 | 90.31 | 3.22 | 2,454,954 | 549 | 549 | 5,756 | 5,756 | 2,850 |
| IjazUZ_2017__S47_a_WGS__bin.1 | 90.07 | 1.66 | 2,468,661 | 376 | 376 | 8,813 | 8,813 | 2,778 |
| MGYG000084316 | 90.07 | 1.66 | 2,468,661 | 376 | 376 | 8,813 | 8,813 | 2,778 |
| XieH_2016__YSZC12003_37190R1__bin.30 | 89.93 | 0.23 | 3,463,597 | 295 | 295 | 17,602 | 17,602 | 3,585 |
| MGYG000089331 | 89.93 | 0.23 | 3,463,597 | 295 | 295 | 17,602 | 17,602 | 3,585 |
| MurphyR_2019_SRR7351692_bin.11 | 89.71 | 0.94 | 3,336,402 | 474 | 474 | 8,878 | 8,878 | 3,442 |
| MGYG000057334 | 89.60 | 4.19 | 3,406,321 | 315 | 315 | 15,510 | 15,510 | 3,410 |
| GeversD_2014__SKBSTL041__bin.56 | 88.77 | 1.42 | 2,388,451 | 333 | 333 | 9,296 | 9,296 | 2,559 |
| MGYG000104828 | 88.77 | 1.42 | 2,388,451 | 333 | 333 | 9,296 | 9,296 | 2,559 |
| MGYG000260649 | 88.71 | 1.41 | 2,329,935 | 534 | 534 | 5,404 | 5,404 | 2,682 |
| MGYG000183293 | 88.69 | 0.58 | 2,364,329 | 217 | 217 | 15,442 | 15,442 | 2,467 |
| XieH_2016__YSZC12003_37179__bin.80 | 88.54 | 1.02 | 2,424,128 | 196 | 196 | 17,990 | 17,990 | 2,522 |
| MGYG000115790 | 88.54 | 1.02 | 2,424,128 | 196 | 196 | 17,990 | 17,990 | 2,522 |
| MGYG000033510 | 88.30 | 0 | 2,350,964 | 177 | 177 | 19,126 | 19,126 | 2,279 |
| MGYG000067284 | 88.21 | 1.75 | 2,308,534 | 581 | 581 | 4,957 | 4,957 | 2,669 |
| FengQ_2015__SID531333__bin.47 | 87.76 | 1.36 | 2,352,310 | 297 | 297 | 10,508 | 10,508 | 2,518 |
| MGYG000100546 | 87.76 | 1.36 | 2,352,310 | 297 | 297 | 10,508 | 10,508 | 2,518 |
| BackhedF_2015__SID87_12M__bin.39 | 87.52 | 1.17 | 2,414,812 | 229 | 229 | 15,282 | 15,282 | 2,533 |
| MGYG000154256 | 87.52 | 1.17 | 2,414,812 | 229 | 229 | 15,282 | 15,282 | 2,533 |
| MGYG000197807 | 87.49 | 4.33 | 3,325,017 | 278 | 278 | 17,210 | 17,210 | 3,287 |
| HeQ_2017__SZAXPI029564-74__bin.70 | 87.45 | 0.49 | 3,481,952 | 348 | 348 | 14,806 | 14,806 | 3,573 |
| MGYG000126162 | 87.45 | 0.49 | 3,481,952 | 348 | 348 | 14,806 | 14,806 | 3,573 |
| MGYG000234569 | 87.23 | 1.02 | 3,405,590 | 268 | 268 | 19,246 | 19,246 | 3,385 |
| MGYG000014135 | 86.86 | 1.85 | 3,231,296 | 415 | 415 | 10,502 | 10,502 | 3,079 |
| MGYG000275853 | 86.83 | 0.15 | 3,393,119 | 147 | 147 | 38,385 | 38,385 | 3,258 |
| MGYG000201716 | 86.07 | 3.31 | 2,327,850 | 285 | 285 | 10,587 | 10,587 | 2,455 |
| BackhedF_2015__SID39_12M__bin.41 | 86.03 | 1.57 | 2,357,953 | 404 | 404 | 7,186 | 7,186 | 2,660 |
| MGYG000259622 | 86.03 | 1.57 | 2,357,953 | 404 | 404 | 7,186 | 7,186 | 2,660 |
| MGYG000147437 | 85.96 | 1.07 | 3,298,593 | 335 | 335 | 13,445 | 13,445 | 3,363 |
| FengQ_2015__SID31367__bin.39 | 85.59 | 2.24 | 3,330,554 | 434 | 434 | 10,985 | 10,985 | 3,417 |
| MGYG000115658 | 85.59 | 2.24 | 3,330,554 | 434 | 434 | 10,985 | 10,985 | 3,417 |
| MGYG000108964 | 85.01 | 1.17 | 2,446,854 | 241 | 241 | 12,275 | 12,275 | 2,604 |
| RaymondF_2016__P20E7__bin.20 | 84.73 | 0.19 | 2,229,022 | 351 | 351 | 7,954 | 7,954 | 2,237 |
| MGYG000239495 | 84.73 | 0.19 | 2,229,022 | 351 | 351 | 7,954 | 7,954 | 2,237 |
| MGYG000214544 | 84.13 | 2.25 | 2,648,863 | 44 | 44 | 88,733 | 88,733 | 2,612 |
| MGYG000237576 | 83.90 | 1.07 | 2,949,887 | 794 | 794 | 4,354 | 4,354 | 3,351 |
| MGYG000032531 | 83.33 | 0 | 2,441,556 | 69 | 69 | 53,698 | 53,698 | 2,298 |
| MGYG000183297 | 83.18 | 0.90 | 3,014,150 | 734 | 734 | 5,108 | 5,108 | 3,337 |
| BackhedF_2015_ERR525896_bin.18 | 81.85 | 2.12 | 2,115,841 | 709 | 709 | 3,558 | 3,558 | 2,630 |
| MGYG000049970 | 81.20 | 0.90 | 2,394,241 | 342 | 342 | 8,462 | 8,462 | 2,655 |
| MGYG000217487 | 81.09 | 2.11 | 2,180,542 | 669 | 669 | 3,949 | 3,949 | 2,655 |
| IjazUZ_2017__S149_a_WGS__bin.14 | 79.80 | 0.98 | 3,141,993 | 724 | 724 | 5,018 | 5,018 | 3,716 |
| MGYG000180135 | 79.80 | 0.98 | 3,141,993 | 724 | 724 | 5,018 | 5,018 | 3,716 |
| MGYG000165712 | 79.74 | 2.89 | 2,076,098 | 699 | 699 | 3,469 | 3,469 | 2,549 |
| FengQ_2015__SID31537__bin.55 | 78.98 | 0.36 | 2,147,458 | 415 | 415 | 6,408 | 6,408 | 2,428 |
| MGYG000089002 | 78.98 | 0.36 | 2,147,458 | 415 | 415 | 6,408 | 6,408 | 2,428 |
| MGYG000276115 | 78.89 | 3.49 | 2,185,273 | 355 | 355 | 7,365 | 7,365 | 2,435 |
| MGYG000146602 | 78.63 | 0.82 | 3,156,989 | 261 | 261 | 18,098 | 18,098 | 3,130 |
| MGYG000151937 | 78.37 | 1.17 | 1,981,994 | 726 | 726 | 3,156 | 3,156 | 2,561 |
| MGYG000082223 | 77.99 | 0.88 | 3,189,959 | 58 | 58 | 90,158 | 90,158 | 2,995 |
| IjazUZ_2017__S46_a_WGS__bin.2 | 77.88 | 2.90 | 2,083,701 | 554 | 554 | 4,255 | 4,255 | 2,634 |
| MGYG000100111 | 77.88 | 2.90 | 2,083,701 | 554 | 554 | 4,255 | 4,255 | 2,634 |
| BackhedF_2015_ERR525961_bin.14 | 77.75 | 1.27 | 1,913,827 | 845 | 845 | 2,498 | 2,498 | 2,598 |
| ZellerG_2014__CCIS24254057ST-4-0__bin.15 | 77.75 | 1.27 | 2,953,898 | 637 | 637 | 5,438 | 5,438 | 2,999 |
| MGYG000141644 | 77.75 | 3.57 | 2,953,898 | 637 | 637 | 5,438 | 5,438 | 2,999 |
| CokerMO_2019_SRR8692181_bin.17 | 77.46 | 0.88 | 2,790,896 | 1072 | 1072 | 3,024 | 3,024 | 3,529 |
| GeversD_2014__SKBSTL016__bin.45 | 76.81 | 0.94 | 1,960,615 | 525 | 525 | 4,158 | 4,158 | 2,312 |
| MGYG000078202 | 76.81 | 0.94 | 1,960,615 | 525 | 525 | 4,158 | 4,158 | 2,312 |
| MurphyR_2019_SRR7411324_bin.13 | 76.35 | 2.26 | 2,990,792 | 768 | 768 | 4,599 | 4,599 | 3,408 |
| IjazUZ_2017__S15_a_WGS__bin.1 | 75.57 | 3.12 | 2,843,086 | 778 | 778 | 3,986 | 3,986 | 3,403 |
| MGYG000257293 | 75.57 | 3.12 | 2,843,086 | 778 | 778 | 3,986 | 3,986 | 3,403 |
| MGYG000127397 | 75.04 | 0.68 | 2,948,052 | 429 | 429 | 7,895 | 7,895 | 3,086 |
| MGYG000177750 | 74.91 | 0.00 | 1,923,449 | 318 | 318 | 7,068 | 7,068 | 2,124 |
| MGYG000235414 | 74.61 | 3.02 | 2,032,034 | 670 | 670 | 3,349 | 3,349 | 2,500 |
| LiJ_2014__V1.CD54-0__bin.3 | 74.48 | 3.45 | 3,487,032 | 334 | 334 | 18,927 | 18,927 | 3,503 |
| MGYG000032252 | 74.48 | 3.45 | 3,487,032 | 334 | 334 | 18,927 | 18,927 | 3,503 |
| FengQ_2015__SID31137__bin.48 | 73.47 | 1.58 | 1,918,844 | 512 | 512 | 4,138 | 4,138 | 2,263 |
| MGYG000196552 | 73.47 | 1.58 | 1,918,844 | 512 | 512 | 4,138 | 4,138 | 2,263 |
| BackhedF_2015__SID546_4M__bin.12 | 73.33 | 1.46 | 2,045,906 | 510 | 510 | 4,738 | 4,738 | 2,421 |
| MGYG000156646 | 73.33 | 1.46 | 2,045,906 | 510 | 510 | 4,738 | 4,738 | 2,421 |
| FengQ_2015__SID31883__bin.30 | 72.82 | 2.40 | 2,849,860 | 688 | 688 | 4,633 | 4,633 | 3,372 |
| MGYG000059009 | 72.82 | 2.40 | 2,849,860 | 688 | 688 | 4,633 | 4,633 | 3,372 |
| MGYG000161697 | 72.24 | 1.07 | 2,572,678 | 527 | 527 | 5,408 | 5,408 | 2,880 |
| MGYG000230250 | 70.77 | 3.61 | 2,099,501 | 302 | 302 | 9,043 | 9,043 | 2,285 |
| MurphyR_2019_SRR7351790_bin.1 | 70.29 | 2.24 | 2,556,966 | 732 | 732 | 3,943 | 3,943 | 2,904 |
| LiJ_2014__V1.CD3-0-PN__bin.36 | 70.14 | 1.72 | 2,258,476 | 356 | 356 | 8,325 | 8,325 | 2,523 |
| MGYG000230595 | 70.14 | 1.72 | 2,258,476 | 356 | 356 | 8,325 | 8,325 | 2,523 |
| MGYG000043161 | 69.20 | 0.74 | 2,568,057 | 1036 | 1036 | 2,833 | 2,833 | 3,248 |
| IjazUZ_2017__S56_a_WGS__bin.7 | 68.37 | 1.38 | 1,871,279 | 535 | 535 | 3,845 | 3,845 | 2,395 |
| MGYG000260044 | 68.37 | 1.38 | 1,871,279 | 535 | 535 | 3,845 | 3,845 | 2,395 |
| MGYG000189753 | 68.02 | 0.58 | 1,888,580 | 416 | 416 | 5,171 | 5,171 | 2,204 |
| MGYG000288033 | 67.90 | 2.59 | 2,468,913 | 539 | 539 | 5,087 | 5,087 | 2,774 |
| BackhedF_2015__SID546_12M__bin.24 | 67.48 | 1.27 | 1,807,772 | 508 | 508 | 3,782 | 3,782 | 2,211 |
| MGYG000118803 | 67.48 | 1.27 | 1,807,772 | 508 | 508 | 3,782 | 3,782 | 2,211 |
| MGYG000158936 | 66.28 | 1.07 | 1,657,119 | 330 | 330 | 5,447 | 5,447 | 1,871 |
| NielsenHB_2014__V1_UC11_0__bin.24 | 65.91 | 1.88 | 2,035,703 | 466 | 466 | 5,282 | 5,282 | 2,414 |
| MGYG000027702 | 65.91 | 1.88 | 2,035,703 | 466 | 466 | 5,282 | 5,282 | 2,414 |
| MGYG000239731 | 65.50 | 2.59 | 2,745,202 | 501 | 501 | 6,282 | 6,282 | 2,970 |
| MGYG000065110 | 65.10 | 0.25 | 2,152,440 | 46 | 46 | 76,086 | 76,086 | 2,143 |
| YuJ_2015__SZAXPI015211-166__bin.3 | 64.53 | 1.17 | 2,703,826 | 720 | 720 | 4,254 | 4,254 | 3,101 |
| MGYG000212469 | 64.53 | 1.17 | 2,703,826 | 720 | 720 | 4,254 | 4,254 | 3,101 |
| MGYG000256913 | 62.28 | 1.23 | 2,683,607 | 526 | 526 | 5,439 | 5,439 | 2,930 |
| MGYG000019508 | 61.92 | 0.58 | 1,489,172 | 681 | 681 | 2,371 | 2,371 | 1,926 |
| MGYG000286990 | 61.80 | 0.12 | 2,356,565 | 442 | 442 | 5,838 | 5,838 | 2,473 |
| YuJ_2015__SZAXPI017457-24__bin.57 | 60.42 | 1.72 | 2,389,318 | 762 | 762 | 3,283 | 3,283 | 2,900 |
| MGYG000060898 | 60.42 | 1.72 | 2,389,318 | 762 | 762 | 3,283 | 3,283 | 2,900 |
| ParnanenK_2018_SRR5723857_bin.3 | 59.51 | 0.88 | 1,513,562 | 752 | 752 | 2,147 | 2,147 | 1,988 |
| IjazUZ_2017__S16_a_WGS__bin.12 | 58.33 | 2.24 | 2,275,143 | 806 | 806 | 2,886 | 2,886 | 3,013 |
| MGYG000277528 | 58.16 | 1.27 | 1,584,323 | 392 | 392 | 4,166 | 4,166 | 1,894 |
| BackhedF_2015__SID577_12M__bin.28 | 56.30 | 1.72 | 1,660,682 | 495 | 495 | 3,684 | 3,684 | 2,055 |
| XieH_2016__YSZC12003_37400__bin.34 | 55.99 | 1.75 | 2,235,680 | 647 | 647 | 3,753 | 3,753 | 2,709 |
| NielsenHB_2014__V1_UC16_0__bin.1 | 55.52 | 0.00 | 2,263,928 | 684 | 684 | 3,521 | 3,521 | 2,731 |
| MGYG000245088 | 55.52 | 0.00 | 2,263,928 | 684 | 684 | 3,521 | 3,521 | 2,731 |
| IjazUZ_2017__S48_a_WGS__bin.2 | 54.62 | 1.72 | 1,559,077 | 524 | 524 | 3,146 | 3,146 | 2,096 |
| BackhedF_2015_ERR525992_bin.38 | 54.28 | 0.00 | 1,371,348 | 732 | 732 | 1,984 | 1,984 | 1,911 |
| VincentC_2016__MM063__bin.1 | 50.78 | 0.00 | 1,583,257 | 561 | 561 | 2,874 | 2,874 | 2,021 |
| MGYG000107506 | 50.78 | 0.00 | 1,583,257 | 561 | 561 | 2,874 | 2,874 | 2,021 |

**Supplementary table 2:** Genomic characteristics of 58 dereplicated MAGs. Genome completeness values are shown in descending order.

| Genome ID | Complete<br>ness % | Contami<br>nation % | Genome<br>size (bp) | N°<br>scaffolds | N°<br>contigs | N50<br>(scaffolds) | N50<br>(contigs) | N°<br>predicted<br>genes |
| --- | --- | --- | --- | --- | --- | --- | --- | --- |
| CokerMO_2019_SRR8692192_bin.27 | 99.42 | 0.00 | 2,981,811 | 74 | 74 | 68,939 | 68,939 | 2.913 |
| GCA_009696415.1_ASM969641v1_genomic | 99.42 | 0.00 | 3,614,384 | 114 | 114 | 87,879 | 87,879 | 3.523 |
| CasaburiG_2019_SRR6277114_bin.5 | 98.83 | 0.19 | 3,885,654 | 21 | 21 | 251,049 | 251,049 | 3.672 |
| MGYG000125324 | 98.83 | 0.00 | 4,056,916 | 17 | 25 | 546,338 | 291,830 | 3.836 |
| HeQ_2017_RSZAXPI003099-133_bin.1 | 98.45 | 0.00 | 3,090,863 | 48 | 48 | 138,734 | 138,734 | 3.042 |
| IjazUZ_2017_S14_a_WGS_bin.19 | 98.11 | 0.12 | 3,562,888 | 211 | 211 | 23,628 | 23,628 | 3.489 |
| MGYG000246300 | 95.81 | 0.00 | 2,569,623 | 357 | 357 | 10,250 | 10,250 | 2.772 |
| MGYG000212934 | 95.76 | 0.00 | 3,362,959 | 282 | 282 | 16,032 | 16,032 | 3.379 |
| GCA_945830785.1_SRR5240736_bin.6_metaWRAP_v1.3_MAG_genomic | 99.42 | 0.00 | 3,099,469 | 82 | 82 | 87,848 | 87,848 | 3.060 |
| GCA_945908315.1_ERR1606358_bin.2_metaWRAP_v1.3_MAG_genomic | 97.08 | 0.00 | 2,946,002 | 57 | 57 | 93,160 | 93,160 | 2.902 |
| GCA_004558675.1_ASM455867v1_genomic | 94.74 | 0.88 | 2,885,217 | 99 | 99 | 48,377 | 48,377 | 2.929 |
| LoombaR_2017_SID5639_uuc_bin.9 | 92.79 | 0.97 | 3,618,456 | 166 | 166 | 31,893 | 31,893 | 3.550 |
| IjazUZ_2017_S102_a_WGS_bin.2 | 90.74 | 2.52 | 2,389,371 | 323 | 323 | 10,060 | 10,060 | 2.601 |
| MGYG000089331 | 89.93 | 0.23 | 3,463,597 | 295 | 295 | 17,602 | 17,602 | 3.585 |
| MurphyR_2019_SRR7351692_bin.11 | 89.71 | 0.94 | 3,336,402 | 474 | 474 | 8,878 | 8,878 | 3.442 |
| MGYG000260649 | 88.71 | 1.41 | 2,329,935 | 534 | 534 | 5,404 | 5,404 | 2.682 |
| MGYG000067284 | 88.21 | 1.75 | 2,308,534 | 581 | 581 | 4,957 | 4,957 | 2.669 |
| FengQ_2015_SID531333_bin.47 | 87.76 | 1.36 | 2,352,310 | 297 | 297 | 10,508 | 10,508 | 2.518 |
| MGYG000201716 | 86.07 | 3.31 | 2,327,850 | 285 | 285 | 10,587 | 10,587 | 2.455 |
| MGYG000239495 | 84.73 | 0.19 | 2,229,022 | 351 | 351 | 7,954 | 7,954 | 2.237 |
| MGYG000032531 | 83.33 | 0 | 2,441,556 | 69 | 69 | 53,698 | 53,698 | 2.298 |
| MGYG000049970 | 81.20 | 0.90 | 2,394,241 | 342 | 342 | 8,462 | 8,462 | 2.655 |
| MGYG000217487 | 81.09 | 2.11 | 2,180,542 | 669 | 669 | 3,949 | 3,949 | 2.655 |
| IjazUZ_2017_S149_a_WGS_bin.14 | 79.80 | 0.98 | 3,141,993 | 724 | 724 | 5,018 | 5,018 | 3.716 |
| FengQ_2015_SID31537_bin.55 | 78.98 | 0.36 | 2,147,458 | 415 | 415 | 6,408 | 6,408 | 2.428 |
| MGYG000151937 | 78.37 | 1.17 | 1,981,994 | 726 | 726 | 3,156 | 3,156 | 2.561 |
| IjazUZ_2017_S46_a_WGS_bin.2 | 77.88 | 2.90 | 2,083,701 | 554 | 554 | 4,255 | 4,255 | 2.634 |
| MGYG000141644 | 77.75 | 3.57 | 2,953,898 | 637 | 637 | 5,438 | 5,438 | 2.999 |
| CokerMO_2019_SRR8692181_bin.17 | 77.46 | 0.88 | 2,790,896 | 1072 | 1072 | 3,024 | 3,024 | 3.529 |
| GeversD_2014_SKBSTL016_bin.45 | 76.81 | 0.94 | 1,960,615 | 525 | 525 | 4,158 | 4,158 | 2.312 |
| MurphyR_2019_SRR7411324_bin.13 | 76.35 | 2.26 | 2,990,792 | 768 | 768 | 4,599 | 4,599 | 3.408 |
| IjazUZ_2017_S15_a_WGS_bin.1 | 75.57 | 3.12 | 2,843,086 | 778 | 778 | 3,986 | 3,986 | 3.403 |
| MGYG000177750 | 74.91 | 0.00 | 1,923,449 | 318 | 318 | 7,068 | 7,068 | 2.124 |
| MGYG000235414 | 74.61 | 3.02 | 2,032,034 | 670 | 670 | 3,349 | 3,349 | 2.500 |
| FengQ_2015_SID31137_bin.48 | 73.47 | 1.58 | 1,918,844 | 512 | 512 | 4,138 | 4,138 | 2.263 |
| BackhedF_2015_SID546_4M_bin.12 | 73.33 | 1.46 | 2,045,906 | 510 | 510 | 4,738 | 4,738 | 2.421 |
| MGYG000161697 | 72.24 | 1.07 | 2,572,678 | 527 | 527 | 5,408 | 5,408 | 2.880 |
| LiJ_2014_V1.CD3-0-PN_bin.36 | 70.14 | 1.72 | 2,258,476 | 356 | 356 | 8,325 | 8,325 | 2.523 |
| MurphyR_2019_SRR7351790_bin.1 | 70.29 | 2.24 | 2,556,966 | 732 | 732 | 3,943 | 3,943 | 2.904 |
| IjazUZ_2017__S56_a_WGS__bin.7 | 68.37 | 1.38 | 1,871,279 | 535 | 535 | 3,845 | 3,845 | 2.395 |
| MGYG000288033 | 67.90 | 2.59 | 2,468,913 | 539 | 539 | 5,087 | 5,087 | 2.774 |
| BackhedF_2015__SID546_12M__bin.24 | 67.48 | 1.27 | 1,807,772 | 508 | 508 | 3,782 | 3,782 | 2.211 |
| MGYG000158936 | 66.28 | 1.07 | 1,657,119 | 330 | 330 | 5,447 | 5,447 | 1.871 |
| MGYG000027702 | 65.91 | 1.88 | 2,035,703 | 466 | 466 | 5,282 | 5,282 | 2.414 |
| MGYG000239731 | 65.50 | 2.59 | 2,745,202 | 501 | 501 | 6,282 | 6,282 | 2.970 |
| MGYG000065110 | 65.10 | 0.25 | 2,152,440 | 46 | 46 | 76,086 | 76,086 | 2.143 |
| MGYG000256913 | 62.28 | 1.23 | 2,683,607 | 526 | 526 | 5,439 | 5,439 | 2.930 |
| MGYG000019508 | 61.92 | 0.58 | 1,489,172 | 681 | 681 | 2,371 | 2,371 | 1.926 |
| MGYG000286990 | 61.80 | 0.12 | 2,356,565 | 442 | 442 | 5,838 | 5,838 | 2.473 |
| MGYG000060898 | 60.42 | 1.72 | 2,389,318 | 762 | 762 | 3,283 | 3,283 | 2.900 |
| ParnanenK_2018_SRR5723857_bin.3 | 59.51 | 0.88 | 1,513,562 | 752 | 752 | 2,147 | 2,147 | 1.988 |
| IjazUZ_2017__S16_a_WGS__bin.12 | 58.33 | 2.24 | 2,275,143 | 806 | 806 | 2,886 | 2,886 | 3.013 |
| BackhedF_2015__SID577_12M__bin.28 | 56.30 | 1.72 | 1,660,682 | 495 | 495 | 3,684 | 3,684 | 2.055 |
| XieH_2016__YSZC12003_37400__bin.34 | 55.99 | 1.75 | 2,235,680 | 647 | 647 | 3,753 | 3,753 | 2.709 |
| MGYG000245088 | 55.52 | 0.00 | 2,263,928 | 684 | 684 | 3,521 | 3,521 | 2.731 |
| IjazUZ_2017__S48_a_WGS__bin.2 | 54.62 | 1.72 | 1,559,077 | 524 | 524 | 3,146 | 3,146 | 2.096 |
| BackhedF_2015_ERR525992_bin.38 | 54.28 | 0.00 | 1,371,348 | 732 | 732 | 1,984 | 1,984 | 1.911 |
| MGYG000107506 | 50.78 | 0.00 | 1,583,257 | 561 | 561 | 2,874 | 2,874 | 2.021 |

**Supplementary table 2:**
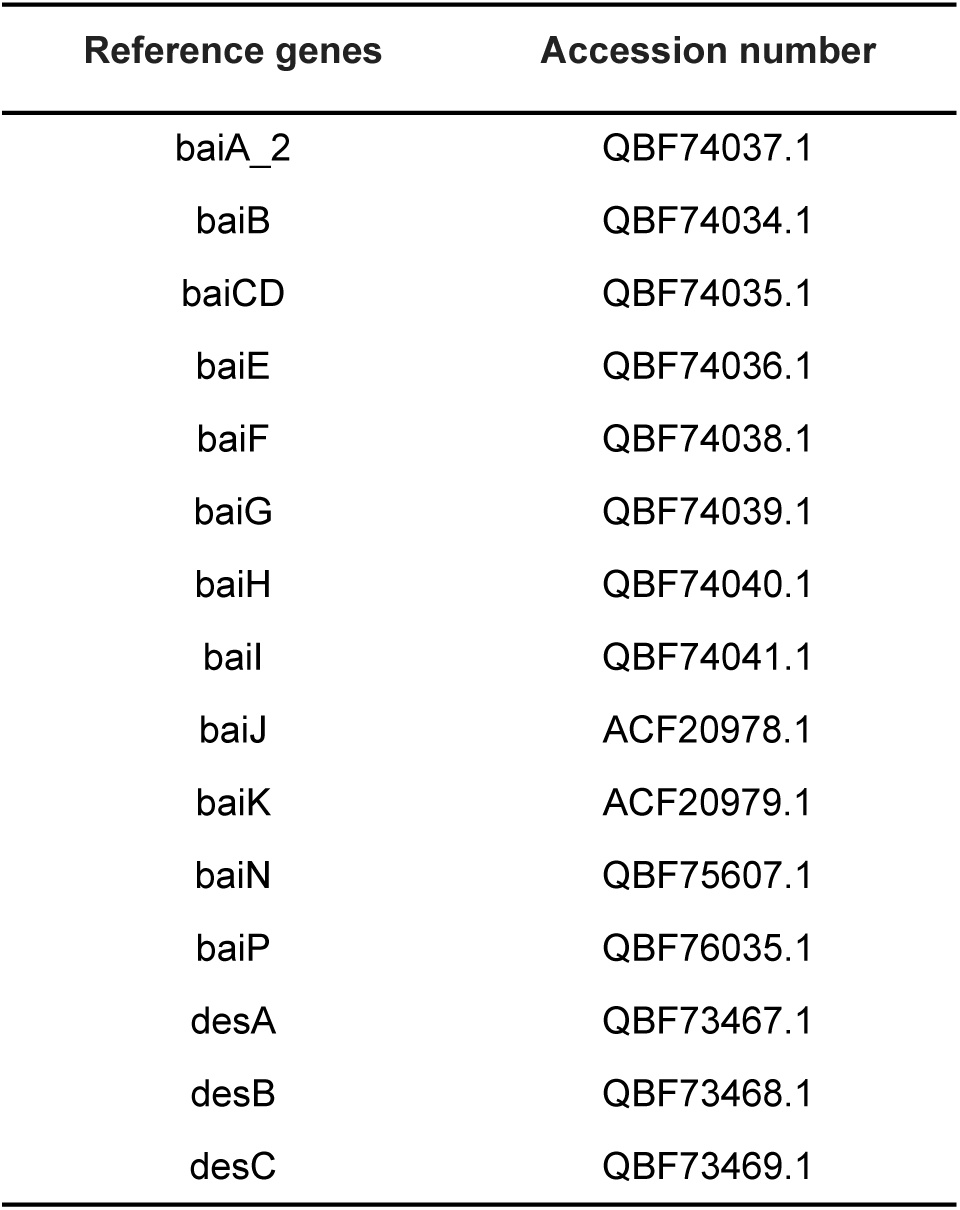
Accession numbers for all reference genes used in the Bai and Des protein searches.

**Supplementary File 1.** Sequence alignment generated by Muscle for all SSU rRNA genes identified in 34 *C. scindens* genomes and in *C. hylemonae*.

**Supplementary File 2.** Uncorrected distance matrix between all SSU rRNA gene sequences identified in 34 *C. scindens* genomes and in *C. hylemonae*, calculated by distmat of the EMBOSS suite.

