## Supplementary File 1 for "Pangenome analysis of *Clostridium scindens*: a collection of diverse bile acid and steroid metabolizing commensal gut bacterial strains"

Alignment: 16SrRNA_v2_sort.afa
Seaview [blocks=10 fontsize=8 A4] on Mon Sep  2 14:18:23 2024

```
                         1
35704-a                  ---AACGAGA GTTTGATCCT GGCTCAGGAT GAACGCTGGC GGCGTGCCTA ACACATGCAA GTCGAACGAA GCGCCTGGCC
35704-b                  ---AACGAGA GTTTGATCCT GGCTCAGGAT GAACGCTGGC GGCGTGCCTA ACACATGCAA GTCGAACGAA GCGCCTGGCC
35704-c                  ---AACGAGA GTTTGATCCT GGCTCAGGAT GAACGCTGGC GGCGTGCCTA ACACATGCAA GTCGAACGAA GCGCCTGGCC
35704-d                  ---AACGAGA GTTTGATCCT GGCTCAGGAT GAACGCTGGC GGCGTGCCTA ACACATGCAA GTCGAACGAA GCGCCTGGCC
AM05-22                  -------AGA GTTTGATCCT GGCTCAGGAT GAACGCTGGC GGCGTGCCTA ACACATGCAA GTCGAACGAA GCGCCTGGCC
AM07-30                  -------AGA GTTTGATCCT GGCTCAGGAT GAACGCTGGC GGCGTGCCTA ACACATGCAA GTCGAACGAA GCGCCTGGCC
BL389WT3D-a              ---AACGAGA GTTTGATCCT GGCTCAGGAT GAACGCTGGC GGCGTGCCTA ACACATGCAA GTCGAACGAA GCGCCTGGCC
BL389WT3D-b              ---AACGAGA GTTTGATCCT GGCTCAGGAT GAACGCTGGC GGCGTGCCTA ACACATGCAA GTCGAACGAA GCGCCTGGCC
BL389WT3D-c              ---AACGAGA GTTTGATCCT GGCTCAGGAT GAACGCTGGC GGCGTGCCTA ACACATGCAA GTCGAACGAA GCGCCTGGCC
BL389WT3D-d              ---AACGAGA GTTTGATCCT GGCTCAGGAT GAACGCTGGC GGCGTGCCTA ACACATGCAA GTCGAACGAA GCGCCTGGCC
CE91-St59-a              ---AACGAGA GTTTGATCCT GGCTCAGGAT GAACGCTGGC GGCGTGCCTA ACACATGCAA GTCGAACGAA GCGCCTGGCC
CE91-St59-b              ---AACGAGA GTTTGATCCT GGCTCAGGAT GAACGCTGGC GGCGTGCCTA ACACATGCAA GTCGAACGAA GCGCCTGGCC
CE91-St59-c              ---AACGAGA GTTTGATCCT GGCTCAGGAT GAACGCTGGC GGCGTGCCTA ACACATGCAA GTCGAACGAA GCGCCTGGCC
CE91-St60-a              T--AACGAGA GTTTGATCCT GGCTCAGGAT GAACGCTGGC GGCGTGCCTA ACACATGCAA GTCGAACGAA GCGCCTGGCC
CE91-St60-b              ---AACGAGA GTTTGATCCT GGCTCAGGAT GAACGCTGGC GGCGTGCCTA ACACATGCAA GTCGAACGAA GCGCCTGGCC
CE91-St60-c              T--AACGAGA GTTTGATCCT GGCTCAGGAT GAACGCTGGC GGCGTGCCTA ACACATGCAA GTCGAACGAA GCGCCTGGCC
FDAARGOS_1227-a          ---AACGAGA GTTTGATCCT GGCTCAGGAT GAACGCTGGC GGCGTGCCTA ACACATGCAA GTCGAACGAA GCGCCTGGCC
FDAARGOS_1227-b          ---AACGAGA GTTTGATCCT GGCTCAGGAT GAACGCTGGC GGCGTGCCTA ACACATGCAA GTCGAACGAA GCGCCTGGCC
FDAARGOS_1227-c          ---AACGAGA GTTTGATCCT GGCTCAGGAT GAACGCTGGC GGCGTGCCTA ACACATGCAA GTCGAACGAA GCGCCTGGCC
FDAARGOS_1227-d          ---AACGAGA GTTTGATCCT GGCTCAGGAT GAACGCTGGC GGCGTGCCTA ACACATGCAA GTCGAACGAA GCGCCTGGCC
G10-a                    ---AACGAGA GTTTGATCCT GGCTCAGGAT GAACGCTGGC GGCGTGCCTA ACACATGCAA GTCGAACGAA GCGCCTGGCC
G10-b                    ---AACGAGA GTTTGATCCT GGCTCAGGAT GAACGCTGGC GGCGTGCCTA ACACATGCAA GTCGAACGAA GCGCCTGGCC
G10-c                    ---AACGAGA GTTTGATCCT GGCTCAGGAT GAACGCTGGC GGCGTGCCTA ACACATGCAA GTCGAACGAA GCGCCTGGCC
G10-d                    ---AACGAGA GTTTGATCCT GGCTCAGGAT GAACGCTGGC GGCGTGCCTA ACACATGCAA GTCGAACGAA GCGCCTGGCC
GGCC_0168                -------AGA GTTTGATCCT GGCTCAGGAT GAACGCTGGC GGCGTGCCTA ACACATGCAA GTCGAACGAA GCGCCTGGCC
I10-a                    ---AACGAGA GTTTGATCCT GGCTCAGGAT GAACGCTGGC GGCGTGCCTA ACACATGCAA GTCGAACGAA GCGCCTGGCC
I10-b                    ---AACGAGA GTTTGATCCT GGCTCAGGAT GAACGCTGGC GGCGTGCCTA ACACATGCAA GTCGAACGAA GCGCCTGGCC
I10-c                    ---AACGAGA GTTTGATCCT GGCTCAGGAT GAACGCTGGC GGCGTGCCTA ACACATGCAA GTCGAACGAA GCGCCTGGCC
I10-d                    ---AACGAGA GTTTGATCCT GGCTCAGGAT GAACGCTGGC GGCGTGCCTA ACACATGCAA GTCGAACGAA GCGCCTGGCC
JCM10422-a               ---AACGAGA GTTTGATCCT GGCTCAGGAT GAACGCTGGC GGCGTGCCTA ACACATGCAA GTCGAACGAA GCGCCTGGCC
JCM10422-b               ---AACGAGA GTTTGATCCT GGCTCAGGAT GAACGCTGGC GGCGTGCCTA ACACATGCAA GTCGAACGAA GCGCCTGGCC
JCM10422-c               ---AACGAGA GTTTGATCCT GGCTCAGGAT GAACGCTGGC GGCGTGCCTA ACACATGCAA GTCGAACGAA GCGCCTGGCC
JCM10422-d               ---AACGAGA GTTTGATCCT GGCTCAGGAT GAACGCTGGC GGCGTGCCTA ACACATGCAA GTCGAACGAA GCGCCTGGCC
MGYG-HGUT-01303-a        -------AGA GTTTGATCCT GGCTCAGGAT GAACGCTGGC GGCGTGCCTA ACACATGCAA GTCGAACGAA GCGCCTGGCC
MGYG-HGUT-01303-b        ---------- ---------- ---------- ---------- ---------- ---------- ---------- ----------
MSK.1.16                 ---------- ---------- ---------- ---------- ---------- ---------- ---------- ----------
MSK.1.26                 -------AGA GTTTGATCCT GGCTCAGGAT GAACGCTGGC GGCGTGCCTA ACACATGCAA GTCGAACGAA GCGCCTGGCC
S077-U-a                 ---AACGAGA GTTTGATCCT GGCTCAGGAT GAACGCTGGC GGCGTGCCTA ACACATGCAA GTCGAACGAA GCGCCTGGCC
S077-U-b                 ---AACGAGA GTTTGATCCT GGCTCAGGAT GAACGCTGGC GGCGTGCCTA ACACATGCAA GTCGAACGAA GCGCCTGGCC
S077-U-c                 ---AACGAGA GTTTGATCCT GGCTCAGGAT GAACGCTGGC GGCGTGCCTA ACACATGCAA GTCGAACGAA GCGCCTGGCC
S077-U-d                 ---AACGAGA GTTTGATCCT GGCTCAGGAT GAACGCTGGC GGCGTGCCTA ACACATGCAA GTCGAACGAA GCGCCTGGCC
VPI12708-a               -------AGA GTTTGATCCT GGCTCAGGAT GAACGCTGGC GGCGTGCCTA ACACATGCAA GTCGAACGAA GCGCTTCCGC
VPI12708-b               -------AGA GTTTGATCCT GGCTCAGGAT GAACGCTGGC GGCGTGCCTA ACACATGCAA GTCGAACGAA GCGCTTCCGC
VPI12708-c               -------AGA GTTTGATCCT GGCTCAGGAT GAACGCTGGC GGCGTGCCTA ACACATGCAA GTCGAACGAA GCGCCTGGCC
VPI12708-d               -------AGA GTTTGATCCT GGCTCAGGAT GAACGCTGGC GGCGTGCCTA ACACATGCAA GTCGAACGAA GCGCCTGGCC
DFI.1.130                -------AGA GTTTGATCCT GGCTCAGGAT GAACGCTGGC GGCGTGCCTA ACACATGCAA GTCGAACGAA GCGCCTGGCC
DFI.1.161                ---------- ---------- ---------- ---------- ---------- ---------- ---------- ----------
DFI.1.162                -------AGA GTTTGATCCT GGCTCAGGAT GAACGCTGGC GGCGTGCCTA ACACATGCAA GTCGAACGAA GCGCTTCCGC
DFI.1.217                -------AGA GTTTGATCCT GGCTCAGGAT GAACGCTGGC GGCGTGCCTA ACACATGCAA GTCGAACGAA GCGCCTGGCC
DFI.1.234                -------AGA GTTTGATCCT GGCTCAGGAT GAACGCTGGC GGCGTGCCTA ACACATGCAA GTCGAACGAA GCGCCTGGCC
DFI.1.60                 -------AGA GTTTGATCCT GGCTCAGGAT GAACGCTGGC GGCGTGCCTA ACACATGCAA GTCGAACGAA GCGCCTGGCC
DFI.4.63                 -------AGA GTTTGATCCT GGCTCAGGAT GAACGCTGGC GGCGTGCCTA ACACATGCAA GTCGAACGAA GCGCTTCCGC
JCM10419-a               ---AACGAGA GTTTGATCCT GGCTCAGGAT GAACGCTGGC GGCGTGCCTA ACACATGCAA GTCGAACGAA GCGCTTCCGC
JCM10419-b               ---AACGAGA GTTTGATCCT GGCTCAGGAT GAACGCTGGC GGCGTGCCTA ACACATGCAA GTCGAACGAA GCGCTTCCGC
JCM10419-c               ---AACGAGA GTTTGATCCT GGCTCAGGAT GAACGCTGGC GGCGTGCCTA ACACATGCAA GTCGAACGAA GCGCTTCCGC
JCM10419-d               ---AACGAGA GTTTGATCCT GGCTCAGGAT GAACGCTGGC GGCGTGCCTA ACACATGCAA GTCGAACGAA GCGCTTCCGC
JCM10420-a               ---AACGAGA GTTTGATCCT GGCTCAGGAT GAACGCTGGC GGCGTGCCTA ACACATGCAA GTCGAACGAA GCGCTTCCGC
JCM10420-b               ---AACGAGA GTTTGATCCT GGCTCAGGAT GAACGCTGGC GGCGTGCCTA ACACATGCAA GTCGAACGAA GCGCTTCCGC
JCM10420-c               ---AACGAGA GTTTGATCCT GGCTCAGGAT GAACGCTGGC GGCGTGCCTA ACACATGCAA GTCGAACGAA GCGCTTCCGC
JCM10420-d               ---AACGAGA GTTTGATCCT GGCTCAGGAT GAACGCTGGC GGCGTGCCTA ACACATGCAA GTCGAACGAA GCGCTTCCGC
JCM10423-a               ---AACGAGA GTTTGATCCT GGCTCAGGAT GAACGCTGGC GGCGTGCCTA ACACATGCAA GTCGAACGAA GCGCTTCCGC
JCM10423-b               ---AACGAGA GTTTGATCCT GGCTCAGGAT GAACGCTGGC GGCGTGCCTA ACACATGCAA GTCGAACGAA GCGCTTCCGC
JCM10423-c               ---AACGAGA GTTTGATCCT GGCTCAGGAT GAACGCTGGC GGCGTGCCTA ACACATGCAA GTCGAACGAA GCGCTTCCGC
JCM10423-d               ---AACGAGA GTTTGATCCT GGCTCAGGAT GAACGCTGGC GGCGTGCCTA ACACATGCAA GTCGAACGAA GCGCTTCCGC
MO32-a                   ---AACGAGA GTTTGATCCT GGCTCAGGAT GAACGCTGGC GGCGTGCCTA ACACATGCAA GTCGAACGAA GCGCTTCCGC
MO32-b                   ---AACGAGA GTTTGATCCT GGCTCAGGAT GAACGCTGGC GGCGTGCCTA ACACATGCAA GTCGAACGAA GCGCTTCCGC
MO32-c                   ---AACGAGA GTTTGATCCT GGCTCAGGAT GAACGCTGGC GGCGTGCCTA ACACATGCAA GTCGAACGAA GCGCCTGGCC
MO32-d                   ---AACGAGA GTTTGATCCT GGCTCAGGAT GAACGCTGGC GGCGTGCCTA ACACATGCAA GTCGAACGAA GCGCCTGGCC
MSK.5.24                 -------AGA GTTTGATCCT GGCTCAGGAT GAACGCTGGC GGCGTGCCTA ACACATGCAA GTCGAACGAA GCGCTTCCGC
NB2A-7-D5                -------AGA GTTTGATCCT GGCTCAGGAT GAACGCTGGC GGCGTGCCTA ACACATGCAA GTCGAACGAA GCGCTTCCGC
NTI82-a                  ---AACGAGA GTTTGATCCT GGCTCAGGAT GAACGCTGGC GGCGTGCCTA ACACATGCAA GTCGAACGAA GCGCTTCCGC
NTI82-b                  ---AACGAGA GTTTGATCCT GGCTCAGGAT GAACGCTGGC GGCGTGCCTA ACACATGCAA GTCGAACGAA GCGCTTCCGC
NTI82-c                  ---AACGAGA GTTTGATCCT GGCTCAGGAT GAACGCTGGC GGCGTGCCTA ACACATGCAA GTCGAACGAA GCGCTTCCGC
NTI82-d                  ---AACGAGA GTTTGATCCT GGCTCAGGAT GAACGCTGGC GGCGTGCCTA ACACATGCAA GTCGAACGAA GCGCTTCCGC
Q4-a                     ---AACGAGA GTTTGATCCT GGCTCAGGAT GAACGCTGGC GGCGTGCCTA ACACATGCAA GTCGAACGAA GCGCTTCCGC
Q4-b                     ---AACGAGA GTTTGATCCT GGCTCAGGAT GAACGCTGGC GGCGTGCCTA ACACATGCAA GTCGAACGAA GCGCTTCCGC
Q4-c                     ---AACGAGA GTTTGATCCT GGCTCAGGAT GAACGCTGGC GGCGTGCCTA ACACATGCAA GTCGAACGAA GCGCTTCCGC
Q4-d                     ---AACGAGA GTTTGATCCT GGCTCAGGAT GAACGCTGGC GGCGTGCCTA ACACATGCAA GTCGAACGAA GCGCTTCCGC
S076-a                   ---AACGAGA GTTTGATCCT GGCTCAGGAT GAACGCTGGC GGCGTGCCTA ACACATGCAA GTCGAACGAA GCGCTTCCGC
S076-b                   ---AACGAGA GTTTGATCCT GGCTCAGGAT GAACGCTGGC GGCGTGCCTA ACACATGCAA GTCGAACGAA GCGCTTCCGC
S076-c                   ---AACGAGA GTTTGATCCT GGCTCAGGAT GAACGCTGGC GGCGTGCCTA ACACATGCAA GTCGAACGAA GCGCTTCCGC
S076-d                   ---AACGAGA GTTTGATCCT GGCTCAGGAT GAACGCTGGC GGCGTGCCTA ACACATGCAA GTCGAACGAA GCGCCTGGCC
SL.1.22                  ---------- ---------- ---------- ---------- ---------- ---------- ---------- ----------
VE202-05                 ---------- ---------- ---------- ---------- ---------- ---------- ---------- ----------
C.hylemonae              TTTAACGAGA GTTTGATCCT GGCTCAGGAT GAACGCTGGC GGCGTGCTTA ACACATGCAA GTCGAACGAA GCAGTACTGT
```

```
                      81
35704-a               CCGACT---- -----TCTTC GGA-ACGAGG AGCCTTGCGA CTGAGTGGCG GACGGGTGAG TAACGCGTGG GCAACCTGCC
35704-b               CCGACT---- -----TCTTC GGA-ACGAGG AGCCTTGCGA CTGAGTGGCG GACGGGTGAG TAACGCGTGG GCAACCTGCC
35704-c               CCGACT---- -----TCTTC GGA-ACGAGG AGCCTTGCGA CTGAGTGGCG GACGGGTGAG TAACGCGTGG GCAACCTGCC
35704-d               CCGACT---- -----TCTTC GGA-ACGAGG AGCCTTGCGA CTGAGTGGCG GACGGGTGAG TAACGCGTGG GCAACCTGCC
AM05-22               CCGACT---- -----TCTTC GGA-ACGAGG AGCCTTGCGA CTGAGTGGCG GACGGGTGAG TAACGCGTGG GCAACCTGCC
AM07-30               CCGACT---- -----TCTTC GGA-ACGAGG AGCCTTGCGA CTGAGTGGCG GACGGGTGAG TAACGCGTGG GCAACCTGCC
BL389WT3D-a           CCGACT---- -----TCTTC GGA-ACGAGG AGCCTTGCGA CTGAGTGGCG GACGGGTGAG TAACGCGTGG GCAACCTGCC
BL389WT3D-b           CCGACT---- -----TCTTC GGA-ACGAGG AGCCTTGCGA CTGAGTGGCG GACGGGTGAG TAACGCGTGG GCAACCTGCC
BL389WT3D-c           CCGACT---- -----TCTTC GGA-ACGAGG AGCCTTGCGA CTGAGTGGCG GACGGGTGAG TAACGCGTGG GCAACCTGCC
BL389WT3D-d           CCGACT---- -----TCTTC GGA-ACGAGG AGCCTTGCGA CTGAGTGGCG GACGGGTGAG TAACGCGTGG GCAACCTGCC
CE91-St59-a           CCGACT---- -----TCTTC GGA-ACGAGG AGCCTTGCGA CTGAGTGGCG GACGGGTGAG TAACGCGTGG GCAACCTGCC
CE91-St59-b           CCGACT---- -----TCTTC GGA-ACGAGG AGCCTTGCGA CTGAGTGGCG GACGGGTGAG TAACGCGTGG GCAACCTGCC
CE91-St59-c           CCGACT---- -----TCTTC GGA-ACGAGG AGCCTTGCGA CTGAGTGGCG GACGGGTGAG TAACGCGTGG GCAACCTGCC
CE91-St60-a           CCGACT---- -----TCTTC GGA-ACGAGG AGCCTTGCGA CTGAGTGGCG GACGGGTGAG TAACGCGTGG GCAACCTGCC
CE91-St60-b           CCGACT---- -----TCTTC GGA-ACGAGG AGCCTTGCGA CTGAGTGGCG GACGGGTGAG TAACGCGTGG GCAACCTGCC
CE91-St60-c           CCGACT---- -----TCTTC GGA-ACGAGG AGCCTTGCGA CTGAGTGGCG GACGGGTGAG TAACGCGTGG GCAACCTGCC
FDAARGOS_1227-a       CCGACT---- -----TCTTC GGA-ACGAGG AGCCTTGCGA CTGAGTGGCG GACGGGTGAG TAACGCGTGG GCAACCTGCC
FDAARGOS_1227-b       CCGACT---- -----TCTTC GGA-ACGAGG AGCCTTGCGA CTGAGTGGCG GACGGGTGAG TAACGCGTGG GCAACCTGCC
FDAARGOS_1227-c       CCGACT---- -----TCTTC GGA-ACGAGG AGCCTTGCGA CTGAGTGGCG GACGGGTGAG TAACGCGTGG GCAACCTGCC
FDAARGOS_1227-d       CCGACT---- -----TCTTC GGA-ACGAGG AGCCTTGCGA CTGAGTGGCG GACGGGTGAG TAACGCGTGG GCAACCTGCC
G10-a                 CCGACT---- -----TCTTC GGA-ACGAGG AGCCTTGCGA CTGAGTGGCG GACGGGTGAG TAACGCGTGG GCAACCTGCC
G10-b                 CCGACT---- -----TCTTC GGA-ACGAGG AGCCTTGCGA CTGAGTGGCG GACGGGTGAG TAACGCGTGG GCAACCTGCC
G10-c                 CCGACT---- -----TCTTC GGA-ACGAGG AGCCTTGCGA CTGAGTGGCG GACGGGTGAG TAACGCGTGG GCAACCTGCC
G10-d                 CCGACT---- -----TCTTC GGA-ACGAGG AGCCTTGCGA CTGAGTGGCG GACGGGTGAG TAACGCGTGG GCAACCTGCC
GGCC_0168             CCGACT---- -----TCTTC GGA-ACGAGG AGCCTTGCGA CTGAGTGGCG GACGGGTGAG TAACGCGTGG GCAACCTGCC
I10-a                 CCGACT---- -----TCTTC GGA-ACGAGG AGCCTTGCGA CTGAGTGGCG GACGGGTGAG TAACGCGTGG GCAACCTGCC
I10-b                 CCGACT---- -----TCTTC GGA-ACGAGG AGCCTTGCGA CTGAGTGGCG GACGGGTGAG TAACGCGTGG GCAACCTGCC
I10-c                 CCGACT---- -----TCTTC GGA-ACGAGG AGCCTTGCGA CTGAGTGGCG GACGGGTGAG TAACGCGTGG GCAACCTGCC
I10-d                 CCGACT---- -----TCTTC GGA-ACGAGG AGCCTTGCGA CTGAGTGGCG GACGGGTGAG TAACGCGTGG GCAACCTGCC
JCM10422-a            CCGACT---- -----TCTTC GGA-ACGAGG AGCCTTGCGA CTGAGTGGCG GACGGGTGAG TAACGCGTGG GCAACCTGCC
JCM10422-b            CCGACT---- -----TCTTC GGA-ACGAGG AGCCTTGCGA CTGAGTGGCG GACGGGTGAG TAACGCGTGG GCAACCTGCC
JCM10422-c            CCGACT---- -----TCTTC GGA-ACGAGG AGCCTTGCGA CTGAGTGGCG GACGGGTGAG TAACGCGTGG GCAACCTGCC
JCM10422-d            CCGACT---- -----TCTTC GGA-ACGAGG AGCCTTGCGA CTGAGTGGCG GACGGGTGAG TAACGCGTGG GCAACCTGCC
MGYG-HGUT-01303-a     CCGACT---- -----TCTTC GGA-ACGAGG AGCCTTGCGA CTGAGTGGCG GACGGGTGAG TAACGCGTGG GCAACCTGCC
MGYG-HGUT-01303-b     ---------- ---------- ---------- ---------- ---------- ---------- ---------- ----------
MSK.1.16              ---------- ---------- ---------- ---------- ---------- ---------- ---------- ----------
MSK.1.26              CCGACT---- -----TCTTC GGA-ACGAGG AGCCTTGCGA CTGAGTGGCG GACGGGTGAG TAACGCGTGG GCAACCTGCC
S077-U-a              CCGACT---- -----TCTTC GGA-ACGAGG AGCCTTGCGA CTGAGTGGCG GACGGGTGAG TAACGCGTGG GCAACCTGCC
S077-U-b              CCGACT---- -----TCTTC GGA-ACGAGG AGCCTTGCGA CTGAGTGGCG GACGGGTGAG TAACGCGTGG GCAACCTGCC
S077-U-c              CCGACT---- -----TCTTC GGA-ACGAGG AGCCTTGCGA CTGAGTGGCG GACGGGTGAG TAACGCGTGG GCAACCTGCC
S077-U-d              CCGACT---- -----TCTTC GGA-ACGAGG AGCCTTGCGA CTGAGTGGCG GACGGGTGAG TAACGCGTGG GCAACCTGCC
VPI12708-a            TAGATT---- ----TTCTTC GGAGATGAAG GCGGCTGCGA CTGAGTGGCG GACGGGTGAG TAACGCGTGG GCAACCTGCC
VPI12708-b            TAGATT---- ----TTCTTC GGAGATGAAG GCGGCTGCGA CTGAGTGGCG GACGGGTGAG TAACGCGTGG GCAACCTGCC
VPI12708-c            CCGACT---- -----TCTTC GGA-ACGAGG AGCCTTGCGA CTGAGTGGCG GACGGGTGAG TAACGCGTGG GCAACCTGCC
VPI12708-d            CCGACT---- -----TCTTC GGA-ACGAGG AGCCTTGCGA CTGAGTGGCG GACGGGTGAG TAACGCGTGG GCAACCTGCC
DFI.1.130             CCGACT---- -----TCTTC GGA-ACGAGG AGCCTTGCGA CTGAGTGGCG GACGGGTGAG TAACGCGTGG GCAACCTGCC
DFI.1.161             ---------- ---------- ---------- -----TGCGA CTGAGTGGCG GACGGGTGAG TAACGCGTGG GCAACCTGCC
DFI.1.162             TAGATT---- ----TTCTTC GGAGATGAAG GCGGCTGCGA CTGAGTGGCG GACGGGTGAG TAACGCGTGG GCAACCTGCC
DFI.1.217             CCGACT---- -----TCTTC GGA-ACGAGG AGCCTTGCGA CTGAGTGGCG GACGGGTGAG TAACGCGTGG GCAACCTGCC
DFI.1.234             CCGACT---- -----TCTTC GGA-ACGAGG AGCCTTGCGA CTGAGTGGCG GACGGGTGAG TAACGCGTGG GCAACCTGCC
DFI.1.60              CCGACT---- -----TCTTC GGA-ACGAGG AGCCTTGCGA CTGAGTGGCG GACGGGTGAG TAACGCGTGG GCAACCTGCC
DFI.4.63              TAGATT---- ----TTCTTC GGAGATGAAG GCGGCTGCGA CTGAGTGGCG GACGGGTGAG TAACGCGTGG GCAACCTGCC
JCM10419-a            TAGATT---- ----TTCTTC GGAGATGAAG GCGGCTGCGA CTGAGTGGCG GACGGGTGAG TAACGCGTGG GCAACCTGCC
JCM10419-b            CTGATT---- ----TTCTTC GGAGATGAAG GCGGCTGCGA CTGAGTGGCG GACGGGTGAG TAACGCGTGG GCAACCTGCC
JCM10419-c            CTGATT---- ----TTCTTC GGAGATGAAG GCGGCTGCGA CTGAGTGGCG GACGGGTGAG TAACGCGTGG GCAACCTGCC
JCM10419-d            CTGATT---- ----TTCTTC GGAGATGAAG GCGGCTGCGA CTGAGTGGCG GACGGGTGAG TAACGCGTGG GCAACCTGCC
JCM10420-a            CTGATT---- ----TTCTTC GGAGATGAAG GCGGCTGCGA CTGAGTGGCG GACGGGTGAG TAACGCGTGG GCAACCTGCC
JCM10420-b            TAGATT---- ----TTCTTC GGAGATGAAG GCGGCTGCGA CTGAGTGGCG GACGGGTGAG TAACGCGTGG GCAACCTGCC
JCM10420-c            CTGATT---- ----TTCTTC GGAGATGAAG GCGGCTGCGA CTGAGTGGCG GACGGGTGAG TAACGCGTGG GCAACCTGCC
JCM10420-d            CTGATT---- ----TTCTTC GGAGATGAAG GCGGCTGCGA CTGAGTGGCG GACGGGTGAG TAACGCGTGG GCAACCTGCC
JCM10423-a            TAGATT---- ----TTCTTC GGAGATGAAG GCGGCTGCGA CTGAGTGGCG GACGGGTGAG TAACGCGTGG GCAACCTGCC
JCM10423-b            CTGATT---- ----TTCTTC GGAGATGAAG GCGGCTGCGA CTGAGTGGCG GACGGGTGAG TAACGCGTGG GCAACCTGCC
JCM10423-c            CTGATT---- ----TTCTTC GGAGATGAAG GCGGCTGCGA CTGAGTGGCG GACGGGTGAG TAACGCGTGG GCAACCTGCC
JCM10423-d            CTGATT---- ----TTCTTC GGAGATGAAG GCGGCTGCGA CTGAGTGGCG GACGGGTGAG TAACGCGTGG GCAACCTGCC
MO32-a                CTGATT---- ----TTCTTC GGAGATGAAG GCGGCTGCGA CTGAGTGGCG GACGGGTGAG TAACGCGTGG GCAACCTGCC
MO32-b                CTGATT---- ----TTCTTC GGAGATGAAG GCGGCTGCGA CTGAGTGGCG GACGGGTGAG TAACGCGTGG GCAACCTGCC
MO32-c                CCGACT---- -----TCTTC GGA-ACGAGG AGCCTTGCGA CTGAGTGGCG GACGGGTGAG TAACGCGTGG GCAACCTGCC
MO32-d                CCGACT---- -----TCTTC GGA-ACGAGG AGCCTTGCGA CTGAGTGGCG GACGGGTGAG TAACGCGTGG GCAACCTGCC
MSK.5.24              CTGATT---- ----TTCTTC GGAGATGAAG GCGGCTGCGA CTGAGTGGCG GACGGGTGAG TAACGCGTGG GCAACCTGCC
NB2A-7-D5             CTGATT---- ----TTCTTC GGAGATGAAG GCGGCTGCGA CTGAGTGGCG GACGGGTGAG TAACGCGTGG GCAACCTGCC
NTI82-a               CTGATT---- ----TTCTTC GGAGATGAAG GCGGCTGCGA CTGAGTGGCG GACGGGTGAG TAACGCGTGG GCAACCTGCC
NTI82-b               CTGATT---- ----TTCTTC GGAGATGAAG GCGGCTGCGA CTGAGTGGCG GACGGGTGAG TAACGCGTGG GCAACCTGCC
NTI82-c               CTGATT---- ----TTCTTC GGAGATGAAG GCGGCTGCGA CTGAGTGGCG GACGGGTGAG TAACGCGTGG GCAACCTGCC
NTI82-d               TAGATT---- ----TTCTTC GGAGATGAAG GCGGCTGCGA CTGAGTGGCG GACGGGTGAG TAACGCGTGG GCAACCTGCC
Q4-a                  CTGATT---- ----TTCTTC GGAGATGAAG GCGGCTGCGA CTGAGTGGCG GACGGGTGAG TAACGCGTGG GCAACCTGCC
Q4-b                  CTGATT---- ----TTCTTC GGAGATGAAG GCGGCTGCGA CTGAGTGGCG GACGGGTGAG TAACGCGTGG GCAACCTGCC
Q4-c                  TAGATT---- ----TTCTTC GGAGATGAAG GCGGCTGCGA CTGAGTGGCG GACGGGTGAG TAACGCGTGG GCAACCTGCC
Q4-d                  CTGATT---- ----TTCTTC GGAGATGAAG GCGGCTGCGA CTGAGTGGCG GACGGGTGAG TAACGCGTGG GCAACCTGCC
S076-a                CTGATT---- ----TTCTTC GGAGATGAAG GCGGCTGCGA CTGAGTGGCG GACGGGTGAG TAACGCGTGG GCAACCTGCC
S076-b                CTGATT---- ----TTCTTC GGAGATGAAG GCGGCTGCGA CTGAGTGGCG GACGGGTGAG TAACGCGTGG GCAACCTGCC
S076-c                CTGATT---- ----TTCTTC GGAGATGAAG GCGGCTGCGA CTGAGTGGCG GACGGGTGAG TAACGCGTGG GCAACCTGCC
S076-d                CCGACT---- -----TCTTC GGA-ACGAGG AGCCTTGCGA CTGAGTGGCG GACGGGTGAG TAACGCGTGG GCAACCTGCC
SL.1.22               ---------- ---------- ---------- ---------- ---------- ---------- ---------- ----------
VE202-05              ---------- ---------- ---------- -------CGA CTGAGTGGCG GACGGGTGAG TAACGCGTGG GCAACCTGCC
C.hylemonae           GTGAAGAGAT TAGCTTGCTA AGATCAGAAC TTTGTATTGA CTGAGTGGCG GACGGGTGAG TAACGCGTGG GCAACCTGCC
```

```
                         161
35704-a               TTGCACTGGG GGATAACAGC CAGAAATGGC TGCTAATACC GCATAAGACC GAAGCGCCGC ATGGCGCGGC GGCCAAAGCC
35704-b               TTGCACTGGG GGATAACAGC CAGAAATGGC TGCTAATACC GCATAAGACC GAAGCGCCGC ATGGCGCGGC GGCCAAAGCC
35704-c               TTGCACTGGG GGATAACAGC CAGAAATGGC TGCTAATACC GCATAAGACC GAAGCGCCGC ATGGCGCGGC GGCCAAAGCC
35704-d               TTGCACTGGG GGATAACAGC CAGAAATGGC TGCTAATACC GCATAAGACC GAAGCGCCGC ATGGCGCGGC GGCCAAAGCC
AM05-22               TTGCACTGGG GGATAACAGC CAGAAATGGC TGCTAATACC GCATAAGACC GAAGCGCCGC ATGGCGCGGC GGCCAAAGCC
AM07-30               TTGCACTGGG GGATAACAGC CAGAAATGGC TGCTAATACC GCATAAGACC GAAGCGCCGC ATGGCGCGGC GGCCAAAGCC
BL389WT3D-a           TTGCACTGGG GGATAACAGC CAGAAATGGC TGCTAATACC GCATAAGACC GAAGCGCCGC ATGGCGCGGC GGCCAAAGCC
BL389WT3D-b           TTGCACTGGG GGATAACAGC CAGAAATGGC TGCTAATACC GCATAAGACC GAAGCGCCGC ATGGCGCGGC GGCCAAAGCC
BL389WT3D-c           TTGCACTGGG GGATAACAGC CAGAAATGGC TGCTAATACC GCATAAGACC GAAGCGCCGC ATGGCGCGGC GGCCAAAGCC
BL389WT3D-d           TTGCACTGGG GGATAACAGC CAGAAATGGC TGCTAATACC GCATAAGACC GAAGCGCCGC ATGGCGCGGC GGCCAAAGCC
CE91-St59-a           TTGCACTGGG GGATAACAGC CAGAAATGGC TGCTAATACC GCATAAGACC GAAGCGCCGC ATGGCGCGGC GGCCAAAGCC
CE91-St59-b           TTGCACTGGG GGATAACAGC CAGAAATGGC TGCTAATACC GCATAAGACC GAAGCGCCGC ATGGCGCGGC GGCCAAAGCC
CE91-St59-c           TTGCACTGGG GGATAACAGC CAGAAATGGC TGCTAATACC GCATAAGACC GAAGCGCCGC ATGGCGCGGC GGCCAAAGCC
CE91-St60-a           TTGCACTGGG GGATAACAGC CAGAAATGGC TGCTAATACC GCATAAGACC GAAGCGCCGC ATGGCGCGGC GGCCAAAGCC
CE91-St60-b           TTGCACTGGG GGATAACAGC CAGAAATGGC TGCTAATACC GCATAAGACC GAAGCGCCGC ATGGCGCGGC GGCCAAAGCC
CE91-St60-c           TTGCACTGGG GGATAACAGC CAGAAATGGC TGCTAATACC GCATAAGACC GAAGCGCCGC ATGGCGCGGC GGCCAAAGCC
FDAARGOS_1227-a       TTGCACTGGG GGATAACAGC CAGAAATGGC TGCTAATACC GCATAAGACC GAAGCGCCGC ATGGCGCGGC GGCCAAAGCC
FDAARGOS_1227-b       TTGCACTGGG GGATAACAGC CAGAAATGGC TGCTAATACC GCATAAGACC GAAGCGCCGC ATGGCGCGGC GGCCAAAGCC
FDAARGOS_1227-c       TTGCACTGGG GGATAACAGC CAGAAATGGC TGCTAATACC GCATAAGACC GAAGCGCCGC ATGGCGCGGC GGCCAAAGCC
FDAARGOS_1227-d       TTGCACTGGG GGATAACAGC CAGAAATGGC TGCTAATACC GCATAAGACC GAAGCGCCGC ATGGCGCGGC GGCCAAAGCC
G10-a                 TTGCACTGGG GGATAACAGC CAGAAATGGC TGCTAATACC GCATAAGACC GAAGCGCCGC ATGGCGCGGC GGCCAAAGCC
G10-b                 TTGCACTGGG GGATAACAGC CAGAAATGGC TGCTAATACC GCATAAGACC GAAGCGCCGC ATGGCGCGGC GGCCAAAGCC
G10-c                 TTGCACTGGG GGATAACAGC CAGAAATGGC TGCTAATACC GCATAAGACC GAAGCGCCGC ATGGCGCGGC GGCCAAAGCC
G10-d                 TTGCACTGGG GGATAACAGC CAGAAATGGC TGCTAATACC GCATAAGACC GAAGCGCCGC ATGGCGCGGC GGCCAAAGCC
GGCC_0168             TTGCACTGGG GGATAACAGC CAGAAATGGC TGCTAATACC GCATAAGACC GAAGCGCCGC ATGGCGCGGC GGCCAAAGCC
I10-a                 TTGCACTGGG GGATAACAGC CAGAAATGGC TGCTAATACC GCATAAGACC GAAGCGCCGC ATGGCGCGGC GGCCAAAGCC
I10-b                 TTGCACTGGG GGATAACAGC CAGAAATGGC TGCTAATACC GCATAAGACC GAAGCGCCGC ATGGCGCGGC GGCCAAAGCC
I10-c                 TTGCACTGGG GGATAACAGC CAGAAATGGC TGCTAATACC GCATAAGACC GAAGCGCCGC ATGGCGCGGC GGCCAAAGCC
I10-d                 TTGCACTGGG GGATAACAGC CAGAAATGGC TGCTAATACC GCATAAGACC GAAGCGCCGC ATGGCGCGGC GGCCAAAGCC
JCM10422-a            TTGCACTGGG GGATAACAGC CAGAAATGGC TGCTAATACC GCATAAGACC GAAGCGCCGC ATGGCGCGGC GGCCAAAGCC
JCM10422-b            TTGCACTGGG GGATAACAGC CAGAAATGGC TGCTAATACC GCATAAGACC GAAGCGCCGC ATGGCGCGGC GGCCAAAGCC
JCM10422-c            TTGCACTGGG GGATAACAGC CAGAAATGGC TGCTAATACC GCATAAGACC GAAGCGTCGC ATGGCGCGGC GGCCAAAGCC
JCM10422-d            TTGCACTGGG GGATAACAGC CAGAAATGGC TGCTAATACC GCATAAGACC GAAGCGCCGC ATGGCGCGGC GGCCAAAGCC
MGYG-HGUT-01303-a     TTGCACTGGG GGATAACAGC CAGAAATGGC TGCTAATACC GCATAAGACC GAAGCGCCGC ATGGCGCGGC GGCCAAAGCC
MGYG-HGUT-01303-b     ---------- ---------- ---------- ---------- ---------- ---------- ---------- ----------
MSK.1.16              ---------- ---------- ---------- ---------- ---------- ---------- ---------- ----------
MSK.1.26              TTGCACTGGG GGATAACAGC CAGAAATGGC TGCTAATACC GCATAAGACC GAAGCGCCGC ATGGCGCGGC GGCCAAAGCC
S077-U-a              TTGCACTGGG GGATAACAGC CAGAAATGGC TGCTAATACC GCATAAGACC GAAGCGCCGC ATGGCGCGGC GGCCAAAGCC
S077-U-b              TTGCACTGGG GGATAACAGC CAGAAATGGC TGCTAATACC GCATAAGACC GAAGCGCCGC ATGGCGCGGC GGCCAAAGCC
S077-U-c              TTGCACTGGG GGATAACAGC CAGAAATGGC TGCTAATACC GCATAAGACC GAAGCGCCGC ATGGCGCGGC GGCCAAAGCC
S077-U-d              TTGCACTGGG GGATAACAGC CAGAAATGGC TGCTAATACC GCATAAGACC GAAGCGCCGC ATGGCGCGGC GGCCAAAGCC
VPI12708-a            TTGCACTGGG GGATAACAGC CAGAAATGGC TGCTAATACC GCATAAGACC GAAGCGCCGC ATGGCGCAGC GGCCAAAGCC
VPI12708-b            TTGCACTGGG GGATAACAGC CAGAAATGGC TGCTAATACC GCATAAGACC GAAGCGCCGC ATGGCGCAGC GGCCAAAGCC
VPI12708-c            TTGCACTGGG GGATAACAGC CAGAAATGGC TGCTAATACC GCATAAGACC GAAGCGCCGC ATGGCGCAGC GGCCAAAGCC
VPI12708-d            TTGCACTGGG GGATAACAGC CAGAAATGGC TGCTAATACC GCATAAGACC GAAGCGCCGC ATGGCGCAGC GGCCAAAGCC
DFI.1.130             TTGCACTGGG GGATAACAGC CAGAAATGGC TGCTAATACC GCATAAGACC GAAGCGCCGC ATGGCGCAGC GGCCAAAGCC
DFI.1.161             TTGCACTGGG GGATAACAGC CAGAAATGGC TGCTAATACC GCATAAGACC GAAGCGCCGC ATGGCGCAGC GGCCAAAGCC
DFI.1.162             TTGCACTGGG GGATAACAGC CAGAAATGGC TGCTAATACC GCATAAGACC GAAGCGCCGC ATGGCGCAGC GGCCAAAGCC
DFI.1.217             TTGCACTGGG GGATAACAGC CAGAAATGGC TGCTAATACC GCATAAGACC GAAGCGCCGC ATGGCGCAGC GGCCAAAGCC
DFI.1.234             TTGCACTGGG GGATAACAGC CAGAAATGGC TGCTAATACC GCATAAGACC GAAGCGCCGC ATGGCGCAGC GGCCAAAGCC
DFI.1.60              TTGCACTGGG GGATAACAGC CAGAAATGGC TGCTAATACC GCATAAGACC GAAGCGCCGC ATGGCGCAGC GGCCAAAGCC
DFI.4.63              TTGCACTGGG GGATAACAGC CAGAAATGGC TGCTAATACC GCATAAGACC GAAGCGCCGC ATGGCGCAGC GGCCAAAGCC
JCM10419-a            TTGCACTGGG GGATAACAGC CAGAAATGGC TGCTAATACC GCATAAGACC GAAGCGCCGC ATGGCGCAGC GGCCAAAGCC
JCM10419-b            TTGCACTGGG GGATAACAGC CAGAAATGGC TGCTAATACC GCATAAGACC GAAGCGCCGC ATGGCGCAGC GGCCAAAGCC
JCM10419-c            TTGCACTGGG GGATAACAGC CAGAAATGGC TGCTAATACC GCATAAGACC GAAGCGCCGC ATGGCGCTGC GGCCAAAGCC
JCM10419-d            TTGCACTGGG GGATAACAGC CAGAAATGGC TGCTAATACC GCATAAGACC GAAGCGCCGC ATGGCGCTGC GGCCAAAGCC
JCM10420-a            TTGCACTGGG GGATAACAGC CAGAAATGGC TGCTAATACC GCATAAGACC GAAGCGCCGC ATGGCGCTGC GGCCAAAGCC
JCM10420-b            TTGCACTGGG GGATAACAGC CAGAAATGGC TGCTAATACC GCATAAGACC GAAGCGCCGC ATGGCGCAGC GGCCAAAGCC
JCM10420-c            TTGCACTGGG GGATAACAGC CAGAAATGGC TGCTAATACC GCATAAGACC GAAGCGCCGC ATGGCGCTGC GGCCAAAGCC
JCM10420-d            TTGCACTGGG GGATAACAGC CAGAAATGGC TGCTAATACC GCATAAGACC GAAGCGCCGC ATGGCGCTGC GGCCAAAGCC
JCM10423-a            TTGCACTGGG GGATAACAGC CAGAAATGGC TGCTAATACC GCATAAGACC GAAGCGCCGC ATGGCGCAGC GGCCAAAGCC
JCM10423-b            TTGCACTGGG GGATAACAGC CAGAAATGGC TGCTAATACC GCATAAGACC GAAGCGCCGC ATGGCGCTGC GGCCAAAGCC
JCM10423-c            TTGCACTGGG GGATAACAGC CAGAAATGGC TGCTAATACC GCATAAGACC GAAGCGCCGC ATGGCGCAGC GGCCAAAGCC
JCM10423-d            TTGCACTGGG GGATAACAGC CAGAAATGGC TGCTAATACC GCATAAGACC GAAGCGCCGC ATGGCGCTGC GGCCAAAGCC
MO32-a                TTGCACTGGG GGATAACAGC CAGAAATGGC TGCTAATACC GCATAAGACC GAAGCGCCGC ATGGCGCAGC GGCCAAAGCC
MO32-b                TTGCACTGGG GGATAACAGC CAGAAATGGC TGCTAATACC GCATAAGACC GAAGCGCCGC ATGGCGCAGC GGCCAAAGCC
MO32-c                TTGCACTGGG GGATAACAGC CAGAAATGGC TGCTAATACC GCATAAGACC GAAGCGCCGC ATGGCGCGGC GGCCAAAGCC
MO32-d                TTGCACTGGG GGATAACAGC CAGAAATGGC TGCTAATACC GCATAAGACC GAAGCGCCGC ATGGCGCAGC GGCCAAAGCC
MSK.5.24              TTGCACTGGG GGATAACAGC CAGAAATGGC TGCTAATACC GCATAAGACC GAAGCGCCGC ATGGCGCAGC GGCCAAAGCC
NB2A-7-D5             TTGCACTGGG GGATAACAGC CAGAAATGGC TGCTAATACC GCATAAGACC GAAGCGCCGC ATGGCGCAGC GGCCAAAGCC
NTI82-a               TTGCACTGGG GGATAACAGC CAGAAATGGC TGCTAATACC GCATAAGACC GAAGCGCCGC ATGGCGCAGC GGCCAAAGCC
NTI82-b               TTGCACTGGG GGATAACAGC CAGAAATGGC TGCTAATACC GCATAAGACC GAAGCGCCGC ATGGCGCAGC GGCCAAAGCC
NTI82-c               TTGCACTGGG GGATAACAGC CAGAAATGGC TGCTAATACC GCATAAGACC GAAGCGCCGC ATGGCGCAGC GGCCAAAGCC
NTI82-d               TTGCACTGGG GGATAACAGC CAGAAATGGC TGCTAATACC GCATAAGACC GAAGCGCCGC ATGGCGCAGC GGCCAAAGCC
Q4-a                  TTGCACTGGG GGATAACAGC CAGAAATGGC TGCTAATACC GCATAAGACC GAAGCGCCGC ATGGCGCTGC GGCCAAAGCC
Q4-b                  TTGCACTGGG GGATAACAGC CAGAAATGGC TGCTAATACC GCATAAGACC GAAGCGCCGC ATGGCGCTGC GGCCAAAGCC
Q4-c                  TTGCACTGGG GGATAACAGC CAGAAATGGC TGCTAATACC GCATAAGACC GAAGCGCCGC ATGGCGCTGC GGCCAAAGCC
Q4-d                  TTGCACTGGG GGATAACAGC CAGAAATGGC TGCTAATACC GCATAAGACC GAAGCGCCGC ATGGCGCTGC GGCCAAAGCC
S076-a                TTGCACTGGG GGATAACAGC CAGAAATGGC TGCTAATACC GCATAAGACC GAAGCGCCGC ATGGCGCAGC GGCCAAAGCC
S076-b                TTGCACTGGG GGATAACAGC CAGAAATGGC TGCTAATACC GCATAAGACC GAAGCGCCGC ATGGCGCAGC GGCCAAAGCC
S076-c                TTGCACTGGG GGATAACAGC CAGAAATGGC TGCTAATACC GCATAAGACC GAAGCGCCGC ATGGCGCAGC GGCCAAAGCC
S076-d                TTGCACTGGG GGATAACAGC CAGAAATGGC TGCTAATACC GCATAAGACC GAAGCGCCGC ATGGCGCAGC GGCCAAAGCC
SL.1.22               ---------- ---------- ---------- ---------- ---------- ---------- ---------- ----------
VE202-05              TTGCACTGGG GGATAACAGC CAGAAATGGC TGCTAATACC GCATAAGACC GAAGCGCCGC ATGGCGCAGC GGCCAAAGCC
C.hylemonae           TTACACAGGG GGATAACAGC TAGAAATGGC TGCTAATACC GCATAAGACC TCAGTACCGC ATGGTAGAGG GGTAAAAACT
```

```
                      241
35704-a               CCGGCGGTGC AAGATGGGCC CGCGTCTGAT TAGGTAGTTG GCGGGGTAAC GGCCCACCAA GCCGACGATC AGTAGCCGAC
35704-b               CCGGCGGTGC AAGATGGGCC CGCGTCTGAT TAGGTAGTTG GCGGGGTAAC GGCCCACCAA GCCGACGATC AGTAGCCGAC
35704-c               CCGGCGGTGC AAGATGGGCC CGCGTCTGAT TAGGTAGTTG GCGGGGTAAC GGCCCACCAA GCCGACGATC AGTAGCCGAC
35704-d               CCGGCGGTGC AAGATGGGCC CGCGTCTGAT TAGGTAGTTG GCGGGGTAAC GGCCCACCAA GCCGACGATC AGTAGCCGAC
AM05-22               CCGGCGGTGC AAGATGGGCC CGCGTCTGAT TAGGTAGTTG GCGGGGTAAC GGCCCACCAA GCCGACGATC AGTAGCCGAC
AM07-30               CCGGCGGTGC AAGATGGGCC CGCGTCTGAT TAGGTAGTTG GCGGGGTAAC GGCCCACCAA GCCGACGATC AGTAGCCGAC
BL389WT3D-a           CCGGCGGTGC AAGATGGGCC CGCGTCTGAT TAGGTAGTTG GCGGGGTAAC GGCCCACCAA GCCGACGATC AGTAGCCGAC
BL389WT3D-b           CCGGCGGTGC AAGATGGGCC CGCGTCTGAT TAGGTAGTTG GCGGGGTAAC GGCCCACCAA GCCGACGATC AGTAGCCGAC
BL389WT3D-c           CCGGCGGTGC AAGATGGGCC CGCGTCTGAT TAGGTAGTTG GCGGGGTAAC GGCCCACCAA GCCGACGATC AGTAGCCGAC
BL389WT3D-d           CCGGCGGTGC AAGATGGGCC CGCGTCTGAT TAGGTAGTTG GCGGGGTAAC GGCCCACCAA GCCGACGATC AGTAGCCGAC
CE91-St59-a           CCGGCGGTGC AAGATGGGCC CGCGTCTGAT TAGGTAGTTG GCGGGGTAAC GGCCCACCAA GCCGACGATC AGTAGCCGAC
CE91-St59-b           CCGGCGGTGC AAGATGGGCC CGCGTCTGAT TAGGTAGTTG GCGGGGTAAC GGCCCACCAA GCCGACGATC AGTAGCCGAC
CE91-St59-c           CCGGCGGTGC AAGATGGGCC CGCGTCTGAT TAGGTAGTTG GCGGGGTAAC GGCCCACCAA GCCGACGATC AGTAGCCGAC
CE91-St60-a           CCGGCGGTGC AAGATGGGCC CGCGTCTGAT TAGGTAGTTG GCGGGGTAAC GGCCCACCAA GCCGACGATC AGTAGCCGAC
CE91-St60-b           CCGGCGGTGC AAGATGGGCC CGCGTCTGAT TAGGTAGTTG GCGGGGTAAC GGCCCACCAA GCCGACGATC AGTAGCCGAC
CE91-St60-c           CCGGCGGTGC AAGATGGGCC CGCGTCTGAT TAGGTAGTTG GCGGGGTAAC GGCCCACCAA GCCGACGATC AGTAGCCGAC
FDAARGOS_1227-a       CCGGCGGTGC AAGATGGGCC CGCGTCTGAT TAGGTAGTTG GCGGGGTAAC GGCCCACCAA GCCGACGATC AGTAGCCGAC
FDAARGOS_1227-b       CCGGCGGTGC AAGATGGGCC CGCGTCTGAT TAGGTAGTTG GCGGGGTAAC GGCCCACCAA GCCGACGATC AGTAGCCGAC
FDAARGOS_1227-c       CCGGCGGTGC AAGATGGGCC CGCGTCTGAT TAGGTAGTTG GCGGGGTAAC GGCCCACCAA GCCGACGATC AGTAGCCGAC
FDAARGOS_1227-d       CCGGCGGTGC AAGATGGGCC CGCGTCTGAT TAGGTAGTTG GCGGGGTAAC GGCCCACCAA GCCGACGATC AGTAGCCGAC
G10-a                 CCGGCGGTGC AAGATGGGCC CGCGTCTGAT TAGGTAGTTG GCGGGGTAAC GGCCCACCAA GCCGACGATC AGTAGCCGAC
G10-b                 CCGGCGGTGC AAGATGGGCC CGCGTCTGAT TAGGTAGTTG GCGGGGTAAC GGCCCACCAA GCCGACGATC AGTAGCCGAC
G10-c                 CCGGCGGTGC AAGATGGGCC CGCGTCTGAT TAGGTAGTTG GCGGGGTAAC GGCCCACCAA GCCGACGATC AGTAGCCGAC
G10-d                 CCGGCGGTGC AAGATGGGCC CGCGTCTGAT TAGGTAGTTG GCGGGGTAAC GGCCCACCAA GCCGACGATC AGTAGCCGAC
GGCC_0168             CCGGCGGTGC AAGATGGGCC CGCGTCTGAT TAGGTAGTTG GCGGGGTAAC GGCCCACCAA GCCGACGATC AGTAGCCGAC
I10-a                 CCGGCGGTGC AAGATGGGCC CGCGTCTGAT TAGGTAGTTG GCGGGGTAAC GGCCCACCAA GCCGACGATC AGTAGCCGAC
I10-b                 CCGGCGGTGC AAGATGGGCC CGCGTCTGAT TAGGTAGTTG GCGGGGTAAC GGCCCACCAA GCCGACGATC AGTAGCCGAC
I10-c                 CCGGCGGTGC AAGATGGGCC CGCGTCTGAT TAGGTAGTTG GCGGGGTAAC GGCCCACCAA GCCGACGATC AGTAGCCGAC
I10-d                 CCGGCGGTGC AAGATGGGCC CGCGTCTGAT TAGGTAGTTG GCGGGGTAAC GGCCCACCAA GCCGACGATC AGTAGCCGAC
JCM10422-a            CCGGCGGTGC AAGATGGGCC CGCGTCTGAT TAGGTAGTTG GCGGGGTAAC GGCCCACCAA GCCGACGATC AGTAGCCGAC
JCM10422-b            CCGGCGGTGC AAGATGGGCC CGCGTCTGAT TAGGTAGTTG GCGGGGTAAC GGCCCACCAA GCCGACGATC AGTAGCCGAC
JCM10422-c            CCGGCGGTGC AAGATGGGCC CGCGTCTGAT TAGGTAGTTG GCGGGGTAAC GGCCCACCAA GCCGACGATC AGTAGCCGAC
JCM10422-d            CCGGCGGTGC AAGATGGGCC CGCGTCTGAT TAGGTAGTTG GCGGGGTAAC GGCCCACCAA GCCGACGATC AGTAGCCGAC
MGYG-HGUT-01303-a     CCGGCGGTGC AAGATGGGCC CGCGTCTGAT TAGGTAGTTG GCGGGGTAAC GGCCCACCAA GCCGACGATC AGTAGCCGAC
MGYG-HGUT-01303-b     ---------- ---------- ---------- ---------- ---------- ---------- ---------- ----------
MSK.1.16              ---------- ---------- ---------- ---------- ---------- ---------- ---------- ----------
MSK.1.26              CCGGCGGTGC AAGATGGGCC CGCGTCTGAT TAGGTAGTTG GCGGGGTAAC GGCCCACCAA GCCGACGATC AGTAGCCGAC
S077-U-a              CCGGCGGTGC AAGATGGGCC CGCGTCTGAT TAGGTAGTTG GCGGGGTAAC GGCCCACCAA GCCGACGATC AGTAGCCGAC
S077-U-b              CCGGCGGTGC AAGATGGGCC CGCGTCTGAT TAGGTAGTTG GCGGGGTAAC GGCCCACCAA GCCGACGATC AGTAGCCGAC
S077-U-c              CCGGCGGTGC AAGATGGGCC CGCGTCTGAT TAGGTAGTTG GCGGGGTAAC GGCCCACCAA GCCGACGATC AGTAGCCGAC
S077-U-d              CCGGCGGTGC AAGATGGGCC CGCGTCTGAT TAGGTAGTTG GCGGGGTAAC GGCCCACCAA GCCGACGATC AGTAGCCGAC
VPI12708-a            CCGGCGGTGC AAGATGGGCC CGCGTCTGAT TAGGTAGTTG GCGGGGTAAC GGCCCACCAA GCCGACGATC AGTAGCCGAC
VPI12708-b            CCGGCGGTGC AAGATGGGCC CGCGTCTGAT TAGGTAGTTG GCGGGGTAAC GGCCCACCAA GCCGACGATC AGTAGCCGAC
VPI12708-c            CCGGCGGTGC AAGATGGGCC CGCGTCTGAT TAGGTAGTTG GCGGGGTAAC GGCCCACCAA GCCGACGATC AGTAGCCGAC
VPI12708-d            CCGGCGGTGC AAGATGGGCC CGCGTCTGAT TAGGTAGTTG GCGGGGTAAC GGCCCACCAA GCCGACGATC AGTAGCCGAC
DFI.1.130             CCGGCGGTGC AAGATGGGCC CGCGTCTGAT TAGGTAGTTG GCGGGGTAAC GGCCCACCAA GCCGACGATC AGTAGCCGAC
DFI.1.161             CCGGCGGTGC AAGATGGGCC CGCGTCTGAT TAGGTAGTTG GCGGGGTAAC GGCCCACCAA GCCGACGATC AGTAGCCGAC
DFI.1.162             CCGGCGGTGC AAGATGGGCC CGCGTCTGAT TAGGTAGTTG GCGGGGTAAC GGCCCACCAA GCCGACGATC AGTAGCCGAC
DFI.1.217             CCGGCGGTGC AAGATGGGCC CGCGTCTGAT TAGGTAGTTG GCGGGGTAAC GGCCCACCAA GCCGACGATC AGTAGCCGAC
DFI.1.234             CCGGCGGTGC AAGATGGGCC CGCGTCTGAT TAGGTAGTTG GCGGGGTAAC GGCCCACCAA GCCGACGATC AGTAGCCGAC
DFI.1.60              CCGGCGGTGC AAGATGGGCC CGCGTCTGAT TAGGTAGTTG GCGGGGTAAC GGCCCACCAA GCCGACGATC AGTAGCCGAC
DFI.4.63              CCGGCGGTGC AAGATGGGCC CGCGTCTGAT TAGGTAGTTG GCGGGGTAAC GGCCCACCAA GCCGACGATC AGTAGCCGAC
JCM10419-a            CCGGCGGTGC AAGATGGGCC CGCGTCTGAT TAGGTAGTTG GCGGGGTAAC GGCCCACCAA GCCGACGATC AGTAGCCGAC
JCM10419-b            CCGGCGGTGC AAGATGGGCC CGCGTCTGAT TAGGTAGTTG GCGGGGTAAC GGCCCACCAA GCCGACGATC AGTAGCCGAC
JCM10419-c            CCGGCGGTGC AAGATGGGCC CGCGTCTGAT TAGGTAGTTG GCGGGGTAAC GGCCCACCAA GCCGACGATC AGTAGCCGAC
JCM10419-d            CCGGCGGTGC AAGATGGGCC CGCGTCTGAT TAGGTAGTTG GCGGGGTAAC GGCCCACCAA GCCGACGATC AGTAGCCGAC
JCM10420-a            CCGGCGGTGC AAGATGGGCC CGCGTCTGAT TAGGTAGTTG GCGGGGTAAC GGCCCACCAA GCCGACGATC AGTAGCCGAC
JCM10420-b            CCGGCGGTGC AAGATGGGCC CGCGTCTGAT TAGGTAGTTG GCGGGGTAAC GGCCCACCAA GCCGACGATC AGTAGCCGAC
JCM10420-c            CCGGCGGTGC AAGATGGGCC CGCGTCTGAT TAGGTAGTTG GCGGGGTAAC GGCCCACCAA GCCGACGATC AGTAGCCGAC
JCM10420-d            CCGGCGGTGC AAGATGGGCC CGCGTCTGAT TAGGTAGTTG GCGGGGTAAC GGCCCACCAA GCCGACGATC AGTAGCCGAC
JCM10423-a            CCGGCGGTGC AAGATGGGCC CGCGTCTGAT TAGGTAGTTG GCGGGGTAAC GGCCCACCAA GCCGACGATC AGTAGCCGAC
JCM10423-b            CCGGCGGTGC AAGATGGGCC CGCGTCTGAT TAGGTAGTTG GCGGGGTAAC GGCCCACCAA GCCGACGATC AGTAGCCGAC
JCM10423-c            CCGGCGGTGC AAGATGGGCC CGCGTCTGAT TAGGTAGTTG GCGGGGTAAC GGCCCACCAA GCCGACGATC AGTAGCCGAC
JCM10423-d            CCGGCGGTGC AAGATGGGCC CGCGTCTGAT TAGGTAGTTG GCGGGGTAAC GGCCCACCAA GCCGACGATC AGTAGCCGAC
MO32-a                CCGGCGGTGC AAGATGGGCC CGCGTCTGAT TAGGTGGTTG GCGGGGTAAC GGCCCACCAA GCCGACGATC AGTAGCCGAC
MO32-b                CCGGCGGTGC AAGATGGGCC CGCGTCTGAT TAGGTGGTTG GCGGGGTAAC GGCCCACCAA GCCGACGATC AGTAGCCGAC
MO32-c                CCGGCGGTGC AAGATGGGCC CGCGTCTGAT TAGGTAGTTG GCGGGGTAAC GGCCCACCAA GCCGACGATC AGTAGCCGAC
MO32-d                CCGGCGGTGC AAGATGGGCC CGCGTCTGAT TAGGTAGTTG GCGGGGTAAC GGCCCACCAA GCCGACGATC AGTAGCCGAC
MSK.5.24              CCGGCGGTGC AAGATGGGCC CGCGTCTGAT TAGGTAGTTG GCGGGGTAAC GGCCCACCAA GCCGACGATC AGTAGCCGAC
NB2A-7-D5             CCGGCGGTGC AAGATGGGCC CGCGTCTGAT TAGGTAGTTG GCGGGGTAAC GGCCCACCAA GCCGACGATC AGTAGCCGAC
NTI82-a               CCGGCGGTGC AAGATGGGCC CGCGTCTGAT TAGGTAGTTG GCGGGGTAAC GGCCCACCAA GCCGACGATC AGTAGCCGAC
NTI82-b               CCGGCGGTGC AAGATGGGCC CGCGTCTGAT TAGGTAGTTG GCGGGGTAAC GGCCCACCAA GCCGACGATC AGTAGCCGAC
NTI82-c               CCGGCGGTGC AAGATGGGCC CGCGTCTGAT TAGGTAGTTG GCGGGGTAAC GGCCCACCAA GCCGACGATC AGTAGCCGAC
NTI82-d               CCGGCGGTGC AAGATGGGCC CGCGTCTGAT TAGGTAGTTG GCGGGGTAAC GGCCCACCAA GCCGACGATC AGTAGCCGAC
Q4-a                  CCGGCGGTGC AAGATGGGCC CGCGTCTGAT TAGGTAGTTG GCGGGGTAAC GGCCCACCAA GCCGACGATC AGTAGCCGAC
Q4-b                  CCGGCGGTGC AAGATGGGCC CGCGTCTGAT TAGGTAGTTG GCGGGGTAAC GGCCCACCAA GCCGACGATC AGTAGCCGAC
Q4-c                  CCGGCGGTGC AAGATGGGCC CGCGTCTGAT TAGGTAGTTG GCGGGGTAAC GGCCCACCAA GCCGACGATC AGTAGCCGAC
Q4-d                  CCGGCGGTGC AAGATGGGCC CGCGTCTGAT TAGGTAGTTG GCGGGGTAAC GGCCCACCAA GCCGACGATC AGTAGCCGAC
S076-a                CCGGCGGTGC AAGATGGGCC CGCGTCTGAT TAGGTGGTTG GCGGGGTAAC GGCCCACCAA GCCGACGATC AGTAGCCGAC
S076-b                CCGGCGGTGC AAGATGGGCC CGCGTCTGAT TAGGTGGTTG GCGGGGTAAC GGCCCACCAA GCCGACGATC AGTAGCCGAC
S076-c                CCGGCGGTGC AAGATGGGCC CGCGTCTGAT TAGGTGGTTG GCGGGGTAAC GGCCCACCAA GCCGACGATC AGTAGCCGAC
S076-d                CCGGCGGTGC AAGATGGGCC CGCGTCTGAT TAGGTAGTTG GCGGGGTAAC GGCCCACCAA GCCGACGATC AGTAGCCGAC
SL.1.22               ---------- ---------- ---------- ------GTTG GCGGGGTAAC GGCCCACCAA GCCGACGATC AGTAGCCGAC
VE202-05              CCGGCGGTGC AAGATGGGCC CGCGTCTGAT TAGGTAGTTG GCGGGGTAAC GGCCCACCAA GCCGACGATC AGTAGCCGAC
C.hylemonae           CCGGTGGTGT AAGATGGGCC CGCGTCTGAT TAGGTAGTTG GTAGGGTAAC GGCCTACCAA GCCGACGATC AGTAGCCGAC
```

```
                    321
35704-a             CTGAGAGGGT GACCGGCCAC ATTGGGACTG AGACACGGCC CAGACTCCTA CGGGAGGCAG CAGTGGGGAA TATTGCACAA
35704-b             CTGAGAGGGT GACCGGCCAC ATTGGGACTG AGACACGGCC CAGACTCCTA CGGGAGGCAG CAGTGGGGAA TATTGCACAA
35704-c             CTGAGAGGGT GACCGGCCAC ATTGGGACTG AGACACGGCC CAGACTCCTA CGGGAGGCAG CAGTGGGGAA TATTGCACAA
35704-d             CTGAGAGGGT GACCGGCCAC ATTGGGACTG AGACACGGCC CAGACTCCTA CGGGAGGCAG CAGTGGGGAA TATTGCACAA
AM05-22             CTGAGAGGGT GACCGGCCAC ATTGGGACTG AGACACGGCC CAGACTCCTA CGGGAGGCAG CAGTGGGGAA TATTGCACAA
AM07-30             CTGAGAGGGT GACCGGCCAC ATTGGGACTG AGACACGGCC CAGACTCCTA CGGGAGGCAG CAGTGGGGAA TATTGCACAA
BL389WT3D-a         CTGAGAGGGT GACCGGCCAC ATTGGGACTG AGACACGGCC CAGACTCCTA CGGGAGGCAG CAGTGGGGAA TATTGCACAA
BL389WT3D-b         CTGAGAGGGT GACCGGCCAC ATTGGGACTG AGACACGGCC CAGACTCCTA CGGGAGGCAG CAGTGGGGAA TATTGCACAA
BL389WT3D-c         CTGAGAGGGT GACCGGCCAC ATTGGGACTG AGACACGGCC CAGACTCCTA CGGGAGGCAG CAGTGGGGAA TATTGCACAA
BL389WT3D-d         CTGAGAGGGT GACCGGCCAC ATTGGGACTG AGACACGGCC CAGACTCCTA CGGGAGGCAG CAGTGGGGAA TATTGCACAA
CE91-St59-a         CTGAGAGGGT GACCGGCCAC ATTGGGACTG AGACACGGCC CAGACTCCTA CGGGAGGCAG CAGTGGGGAA TATTGCACAA
CE91-St59-b         CTGAGAGGGT GACCGGCCAC ATTGGGACTG AGACACGGCC CAGACTCCTA CGGGAGGCAG CAGTGGGGAA TATTGCACAA
CE91-St59-c         CTGAGAGGGT GACCGGCCAC ATTGGGACTG AGACACGGCC CAGACTCCTA CGGGAGGCAG CAGTGGGGAA TATTGCACAA
CE91-St60-a         CTGAGAGGGT GACCGGCCAC ATTGGGACTG AGACACGGCC CAGACTCCTA CGGGAGGCAG CAGTGGGGAA TATTGCACAA
CE91-St60-b         CTGAGAGGGT GACCGGCCAC ATTGGGACTG AGACACGGCC CAGACTCCTA CGGGAGGCAG CAGTGGGGAA TATTGCACAA
CE91-St60-c         CTGAGAGGGT GACCGGCCAC ATTGGGACTG AGACACGGCC CAGACTCCTA CGGGAGGCAG CAGTGGGGAA TATTGCACAA
FDAARGOS_1227-a     CTGAGAGGGT GACCGGCCAC ATTGGGACTG AGACACGGCC CAGACTCCTA CGGGAGGCAG CAGTGGGGAA TATTGCACAA
FDAARGOS_1227-b     CTGAGAGGGT GACCGGCCAC ATTGGGACTG AGACACGGCC CAGACTCCTA CGGGAGGCAG CAGTGGGGAA TATTGCACAA
FDAARGOS_1227-c     CTGAGAGGGT GACCGGCCAC ATTGGGACTG AGACACGGCC CAGACTCCTA CGGGAGGCAG CAGTGGGGAA TATTGCACAA
FDAARGOS_1227-d     CTGAGAGGGT GACCGGCCAC ATTGGGACTG AGACACGGCC CAGACTCCTA CGGGAGGCAG CAGTGGGGAA TATTGCACAA
G10-a               CTGAGAGGGT GACCGGCCAC ATTGGGACTG AGACACGGCC CAGACTCCTA CGGGAGGCAG CAGTGGGGAA TATTGCACAA
G10-b               CTGAGAGGGT GACCGGCCAC ATTGGGACTG AGACACGGCC CAGACTCCTA CGGGAGGCAG CAGTGGGGAA TATTGCACAA
G10-c               CTGAGAGGGT GACCGGCCAC ATTGGGACTG AGACACGGCC CAGACTCCTA CGGGAGGCAG CAGTGGGGAA TATTGCACAA
G10-d               CTGAGAGGGT GACCGGCCAC ATTGGGACTG AGACACGGCC CAGACTCCTA CGGGAGGCAG CAGTGGGGAA TATTGCACAA
GGCC_0168           CTGAGAGGGT GACCGGCCAC ATTGGGACTG AGACACGGCC CAGACTCCTA CGGGAGGCAG CAGTGGGGAA TATTGCACAA
I10-a               CTGAGAGGGT GACCGGCCAC ATTGGGACTG AGACACGGCC CAGACTCCTA CGGGAGGCAG CAGTGGGGAA TATTGCACAA
I10-b               CTGAGAGGGT GACCGGCCAC ATTGGGACTG AGACACGGCC CAGACTCCTA CGGGAGGCAG CAGTGGGGAA TATTGCACAA
I10-c               CTGAGAGGGT GACCGGCCAC ATTGGGACTG AGACACGGCC CAGACTCCTA CGGGAGGCAG CAGTGGGGAA TATTGCACAA
I10-d               CTGAGAGGGT GACCGGCCAC ATTGGGACTG AGACACGGCC CAGACTCCTA CGGGAGGCAG CAGTGGGGAA TATTGCACAA
JCM10422-a          CTGAGAGGGT GACCGGCCAC ATTGGGACTG AGACACGGCC CAGACTCCTA CGGGAGGCAG CAGTGGGGAA TATTGCACAA
JCM10422-b          CTGAGAGGGT GACCGGCCAC ATTGGGACTG AGACACGGCC CAGACTCCTA CGGGAGGCAG CAGTGGGGAA TATTGCACAA
JCM10422-c          CTGAGAGGGT GACCGGCCAC ATTGGGACTG AGACACGGCC CAGACTCCTA CGGGAGGCAG CAGTGGGGAA TATTGCACAA
JCM10422-d          CTGAGAGGGT GACCGGCCAC ATTGGGACTG AGACACGGCC CAGACTCCTA CGGGAGGCAG CAGTGGGGAA TATTGCACAA
MGYG-HGUT-01303-a   CTGAGAGGGT GACCGGCCAC ATTGGGACTG AGACACGGCC CAGACTCCTA CGGGAGGCAG CAGTGGGGAA TATTGCACAA
MGYG-HGUT-01303-b   ---------- ---------- ---------- ---------- ---------- ---------- ---------- ----------
MSK.1.16            ---------T GACCGGCCAC ATTGGGACTG AGACACGGCC CAGACTCCTA CGGGAGGCAG CAGTGGGGAA TATTGCACAA
MSK.1.26            CTGAGAGGGT GACCGGCCAC ATTGGGACTG AGACACGGCC CAGACTCCTA CGGGAGGCAG CAGTGGGGAA TATTGCACAA
S077-U-a            CTGAGAGGGT GACCGGCCAC ATTGGGACTG AGACACGGCC CAGACTCCTA CGGGAGGCAG CAGTGGGGAA TATTGCACAA
S077-U-b            CTGAGAGGGT GACCGGCCAC ATTGGGACTG AGACACGGCC CAGACTCCTA CGGGAGGCAG CAGTGGGGAA TATTGCACAA
S077-U-c            CTGAGAGGGT GACCGGCCAC ATTGGGACTG AGACACGGCC CAGACTCCTA CGGGAGGCAG CAGTGGGGAA TATTGCACAA
S077-U-d            CTGAGAGGGT GACCGGCCAC ATTGGGACTG AGACACGGCC CAGACTCCTA CGGGAGGCAG CAGTGGGGAA TATTGCACAA
VPI12708-a          CTGAGAGGGT GACCGGCCAC ATTGGGACTG AGACACGGCC CAGACTCCTA CGGGAGGCAG CAGTGGGGAA TATTGCACAA
VPI12708-b          CTGAGAGGGT GACCGGCCAC ATTGGGACTG AGACACGGCC CAGACTCCTA CGGGAGGCAG CAGTGGGGAA TATTGCACAA
VPI12708-c          CTGAGAGGGT GACCGGCCAC ATTGGGACTG AGACACGGCC CAGACTCCTA CGGGAGGCAG CAGTGGGGAA TATTGCACAA
VPI12708-d          CTGAGAGGGT GACCGGCCAC ATTGGGACTG AGACACGGCC CAGACTCCTA CGGGAGGCAG CAGTGGGGAA TATTGCACAA
DFI.1.130           CTGAGAGGGT GACCGGCCAC ATTGGGACTG AGACACGGCC CAGACTCCTA CGGGAGGCAG CAGTGGGGAA TATTGCACAA
DFI.1.161           CTGAGAGGGT GACCGGCCAC ATTGGGACTG AGACACGGCC CAGACTCCTA CGGGAGGCAG CAGTGGGGAA TATTGCACAA
DFI.1.162           CTGAGAGGGT GACCGGCCAC ATTGGGACTG AGACACGGCC CAGACTCCTA CGGGAGGCAG CAGTGGGGAA TATTGCACAA
DFI.1.217           CTGAGAGGGT GACCGGCCAC ATTGGGACTG AGACACGGCC CAGACTCCTA CGGGAGGCAG CAGTGGGGAA TATTGCACAA
DFI.1.234           CTGAGAGGGT GACCGGCCAC ATTGGGACTG AGACACGGCC CAGACTCCTA CGGGAGGCAG CAGTGGGGAA TATTGCACAA
DFI.1.60            CTGAGAGGGT GACCGGCCAC ATTGGGACTG AGACACGGCC CAGACTCCTA CGGGAGGCAG CAGTGGGGAA TATTGCACAA
DFI.4.63            CTGAGAGGGT GACCGGCCAC ATTGGGACTG AGACACGGCC CAGACTCCTA CGGGAGGCAG CAGTGGGGAA TATTGCACAA
JCM10419-a          CTGAGAGGGT GACCGGCCAC ATTGGGACTG AGACACGGCC CAGACTCCTA CGGGAGGCAG CAGTGGGGAA TATTGCACAA
JCM10419-b          CTGAGAGGGT GACCGGCCAC ATTGGGACTG AGACACGGCC CAGACTCCTA CGGGAGGCAG CAGTGGGGAA TATTGCACAA
JCM10419-c          CTGAGAGGGT GACCGGCCAC ATTGGGACTG AGACACGGCC CAGACTCCTA CGGGAGGCAG CAGTGGGGAA TATTGCACAA
JCM10419-d          CTGAGAGGGT GACCGGCCAC ATTGGGACTG AGACACGGCC CAGACTCCTA CGGGAGGCAG CAGTGGGGAA TATTGCACAA
JCM10420-a          CTGAGAGGGT GACCGGCCAC ATTGGGACTG AGACACGGCC CAGACTCCTA CGGGAGGCAG CAGTGGGGAA TATTGCACAA
JCM10420-b          CTGAGAGGGT GACCGGCCAC ATTGGGACTG AGACACGGCC CAGACTCCTA CGGGAGGCAG CAGTGGGGAA TATTGCACAA
JCM10420-c          CTGAGAGGGT GACCGGCCAC ATTGGGACTG AGACACGGCC CAGACTCCTA CGGGAGGCAG CAGTGGGGAA TATTGCACAA
JCM10420-d          CTGAGAGGGT GACCGGCCAC ATTGGGACTG AGACACGGCC CAGACTCCTA CGGGAGGCAG CAGTGGGGAA TATTGCACAA
JCM10423-a          CTGAGAGGGT GACCGGCCAC ATTGGGACTG AGACACGGCC CAGACTCCTA CGGGAGGCAG CAGTGGGGAA TATTGCACAA
JCM10423-b          CTGAGAGGGT GACCGGCCAC ATTGGGACTG AGACACGGCC CAGACTCCTA CGGGAGGCAG CAGTGGGGAA TATTGCACAA
JCM10423-c          CTGAGAGGGT GACCGGCCAC ATTGGGACTG AGACACGGCC CAGACTCCTA CGGGAGGCAG CAGTGGGGAA TATTGCACAA
JCM10423-d          CTGAGAGGGT GACCGGCCAC ATTGGGACTG AGACACGGCC CAGACTCCTA CGGGAGGCAG CAGTGGGGAA TATTGCACAA
MO32-a              CTGAGAGGGT GACCGGCCAC ATTGGGACTG AGACACGGCC CAGACTCCTA CGGGAGGCAG CAGTGGGGAA TATTGCACAA
MO32-b              CTGAGAGGGT GACCGGCCAC ATTGGGACTG AGACACGGCC CAGACTCCTA CGGGAGGCAG CAGTGGGGAA TATTGCACAA
MO32-c              CTGAGAGGGT GACCGGCCAC ATTGGGACTG AGACACGGCC CAGACTCCTA CGGGAGGCAG CAGTGGGGAA TATTGCACAA
MO32-d              CTGAGAGGGT GACCGGCCAC ATTGGGACTG AGACACGGCC CAGACTCCTA CGGGAGGCAG CAGTGGGGAA TATTGCACAA
MSK.5.24            CTGAGAGGGT GACCGGCCAC ATTGGGACTG AGACACGGCC CAGACTCCTA CGGGAGGCAG CAGTGGGGAA TATTGCACAA
NB2A-7-D5           CTGAGAGGGT GACCGGCCAC ATTGGGACTG AGACACGGCC CAGACTCCTA CGGGAGGCAG CAGTGGGGAA TATTGCACAA
NTI82-a             CTGAGAGGGT GACCGGCCAC ATTGGGACTG AGACACGGCC CAGACTCCTA CGGGAGGCAG CAGTGGGGAA TATTGCACAA
NTI82-b             CTGAGAGGGT GACCGGCCAC ATTGGGACTG AGACACGGCC CAGACTCCTA CGGGAGGCAG CAGTGGGGAA TATTGCACAA
NTI82-c             CTGAGAGGGT GACCGGCCAC ATTGGGACTG AGACACGGCC CAGACTCCTA CGGGAGGCAG CAGTGGGGAA TATTGCACAA
NTI82-d             CTGAGAGGGT GACCGGCCAC ATTGGGACTG AGACACGGCC CAGACTCCTA CGGGAGGCAG CAGTGGGGAA TATTGCACAA
Q4-a                CTGAGAGGGT GACCGGCCAC ATTGGGACTG AGACACGGCC CAGACTCCTA CGGGAGGCAG CAGTGGGGAA TATTGCACAA
Q4-b                CTGAGAGGGT GACCGGCCAC ATTGGGACTG AGACACGGCC CAGACTCCTA CGGGAGGCAG CAGTGGGGAA TATTGCACAA
Q4-c                CTGAGAGGGT GACCGGCCAC ATTGGGACTG AGACACGGCC CAGACTCCTA CGGGAGGCAG CAGTGGGGAA TATTGCACAA
Q4-d                CTGAGAGGGT GACCGGCCAC ATTGGGACTG AGACACGGCC CAGACTCCTA CGGGAGGCAG CAGTGGGGAA TATTGCACAA
S076-a              CTGAGAGGGT GACCGGCCAC ATTGGGACTG AGACACGGCC CAGACTCCTA CGGGAGGCAG CAGTGGGGAA TATTGCACAA
S076-b              CTGAGAGGGT GACCGGCCAC ATTGGGACTG AGACACGGCC CAGACTCCTA CGGGAGGCAG CAGTGGGGAA TATTGCACAA
S076-c              CTGAGAGGGT GACCGGCCAC ATTGGGACTG AGACACGGCC CAGACTCCTA CGGGAGGCAG CAGTGGGGAA TATTGCACAA
S076-d              CTGAGAGGGT GACCGGCCAC ATTGGGACTG AGACACGGCC CAGACTCCTA CGGGAGGCAG CAGTGGGGAA TATTGCACAA
SL.1.22             CTGAGAGGGT GACCGGCCAC ATTGGGACTG AGACACGGCC CAGACTCCTA CGGGAGGCAG CAGTGGGGAA TATTGCACAA
VE202-05            CTGAGAGGGT GACCGGCCAC ATTGGGACTG AGACACGGCC CAGACTCCTA CGGGAGGCAG CAGTGGGGAA TATTGCACAA
C.hylemonae         CTGAGAGGGT GACCGGCCAC ATTGGGACTG AGACACGGCC CAAACTCCTA CGGGAGGCAG CAGTGGGGAA TATTGCACAA
```

```
                   401
35704-a            TGGGGGAAAC CCTGATGCAG CGACGCCGCG TGAAGGATGA AGTATTTCGG TATGTAAACT TCTATCAGCA GGGAAGAAGA
35704-b            TGGGGGAAAC CCTGATGCAG CGACGCCGCG TGAAGGATGA AGTATTTCGG TATGTAAACT TCTATCAGCA GGGAAGAAGA
35704-c            TGGGGGAAAC CCTGATGCAG CGACGCCGCG TGAAGGATGA AGTATTTCGG TATGTAAACT TCTATCAGCA GGGAAGAAGA
35704-d            TGGGGGAAAC CCTGATGCAG CGACGCCGCG TGAAGGATGA AGTATTTCGG TATGTAAACT TCTATCAGCA GGGAAGAAGA
AM05-22            TGGGGGAAAC CCTGATGCAG CGACGCCGCG TGAAGGATGA AGTATTTCGG TATGTAAACT TCTATCAGCA GGGAAGAAGA
AM07-30            TGGGGGAAAC CCTGATGCAG CGACGCCGCG TGAAGGATGA AGTATTTCGG TATGTAAACT TCTATCAGCA GGGAAGAAGA
BL389WT3D-a        TGGGGGAAAC CCTGATGCAG CGACGCCGCG TGAAGGATGA AGTATTTCGG TATGTAAACT TCTATCAGCA GGGAAGAAGA
BL389WT3D-b        TGGGGGAAAC CCTGATGCAG CGACGCCGCG TGAAGGATGA AGTATTTCGG TATGTAAACT TCTATCAGCA GGGAAGAAGA
BL389WT3D-c        TGGGGGAAAC CCTGATGCAG CGACGCCGCG TGAAGGATGA AGTATTTCGG TATGTAAACT TCTATCAGCA GGGAAGAAGA
BL389WT3D-d        TGGGGGAAAC CCTGATGCAG CGACGCCGCG TGAAGGATGA AGTATTTCGG TATGTAAACT TCTATCAGCA GGGAAGAAGA
CE91-St59-a        TGGGGGAAAC CCTGATGCAG CGACGCCGCG TGAAGGATGA AGTATTTCGG TATGTAAACT TCTATCAGCA GGGAAGAAGA
CE91-St59-b        TGGGGGAAAC CCTGATGCAG CGACGCCGCG TGAAGGATGA AGTATTTCGG TATGTAAACT TCTATCAGCA GGGAAGAAGA
CE91-St59-c        TGGGGGAAAC CCTGATGCAG CGACGCCGCG TGAAGGATGA AGTATTTCGG TATGTAAACT TCTATCAGCA GGGAAGAAGA
CE91-St60-a        TGGGGGAAAC CCTGATGCAG CGACGCCGCG TGAAGGATGA AGTATTTCGG TATGTAAACT TCTATCAGCA GGGAAGAAGA
CE91-St60-b        TGGGGGAAAC CCTGATGCAG CGACGCCGCG TGAAGGATGA AGTATTTCGG TATGTAAACT TCTATCAGCA GGGAAGAAGA
CE91-St60-c        TGGGGGAAAC CCTGATGCAG CGACGCCGCG TGAAGGATGA AGTATTTCGG TATGTAAACT TCTATCAGCA GGGAAGAAGA
FDAARGOS_1227-a    TGGGGGAAAC CCTGATGCAG CGACGCCGCG TGAAGGATGA AGTATTTCGG TATGTAAACT TCTATCAGCA GGGAAGAAGA
FDAARGOS_1227-b    TGGGGGAAAC CCTGATGCAG CGACGCCGCG TGAAGGATGA AGTATTTCGG TATGTAAACT TCTATCAGCA GGGAAGAAGA
FDAARGOS_1227-c    TGGGGGAAAC CCTGATGCAG CGACGCCGCG TGAAGGATGA AGTATTTCGG TATGTAAACT TCTATCAGCA GGGAAGAAGA
FDAARGOS_1227-d    TGGGGGAAAC CCTGATGCAG CGACGCCGCG TGAAGGATGA AGTATTTCGG TATGTAAACT TCTATCAGCA GGGAAGAAGA
G10-a              TGGGGGAAAC CCTGATGCAG CGACGCCGCG TGAAGGATGA AGTATTTCGG TATGTAAACT TCTATCAGCA GGGAAGAAGA
G10-b              TGGGGGAAAC CCTGATGCAG CGACGCCGCG TGAAGGATGA AGTATTTCGG TATGTAAACT TCTATCAGCA GGGAAGAAGA
G10-c              TGGGGGAAAC CCTGATGCAG CGACGCCGCG TGAAGGATGA AGTATTTCGG TATGTAAACT TCTATCAGCA GGGAAGAAGA
G10-d              TGGGGGAAAC CCTGATGCAG CGACGCCGCG TGAAGGATGA AGTATTTCGG TATGTAAACT TCTATCAGCA GGGAAGAAGA
GGCC_0168          TGGGGGAAAC CCTGATGCAG CGACGCCGCG TGAAGGATGA AGTATTTCGG TATGTAAACT TCTATCAGCA GGGAAGAAGA
I10-a              TGGGGGAAAC CCTGATGCAG CGACGCCGCG TGAAGGATGA AGTATTTCGG TATGTAAACT TCTATCAGCA GGGAAGAAGA
I10-b              TGGGGGAAAC CCTGATGCAG CGACGCCGCG TGAAGGATGA AGTATTTCGG TATGTAAACT TCTATCAGCA GGGAAGAAGA
I10-c              TGGGGGAAAC CCTGATGCAG CGACGCCGCG TGAAGGATGA AGTATTTCGG TATGTAAACT TCTATCAGCA GGGAAGAAGA
I10-d              TGGGGGAAAC CCTGATGCAG CGACGCCGCG TGAAGGATGA AGTATTTCGG TATGTAAACT TCTATCAGCA GGGAAGAAGA
JCM10422-a         TGGGGGAAAC CCTGATGCAG CGACGCCGCG TGAAGGATGA AGTATTTCGG TATGTAAACT TCTATCAGCA GGGAAGAAGA
JCM10422-b         TGGGGGAAAC CCTGATGCAG CGACGCCGCG TGAAGGATGA AGTATTTCGG TATGTAAACT TCTATCAGCA GGGAAGAAGA
JCM10422-c         TGGGGGAAAC CCTGATGCAG CGACGCCGCG TGAAGGATGA AGTATTTCGG TATGTAAACT TCTATCAGCA GGGAAGAAGA
JCM10422-d         TGGGGGAAAC CCTGATGCAG CGACGCCGCG TGAAGGATGA AGTATTTCGG TATGTAAACT TCTATCAGCA GGGAAGAAGA
MGYG-HGUT-01303-a  TGGGGGAAAC CCTGATGCAG CGACGCCGCG TGAAGGATGA AGTATTTCGG TATGTAAACT TCTATCAGCA GGGAAGAAGA
MGYG-HGUT-01303-b  ---------- ---------- ---------- ---------- ---------- ---------- ---------- ----------
MSK.1.16           TGGGGGAAAC CCTGATGCAG CGACGCCGCG TGAAGGATGA AGTATTTCGG TATGTAAACT TCTATCAGCA GGGAAGAAGA
MSK.1.26           TGGGGGAAAC CCTGATGCAG CGACGCCGCG TGAAGGATGA AGTATTTCGG TATGTAAACT TCTATCAGCA GGGAAGAAGA
S077-U-a           TGGGGGAAAC CCTGATGCAG CGACGCCGCG TGAAGGATGA AGTATTTCGG TATGTAAACT TCTATCAGCA GGGAAGAAGA
S077-U-b           TGGGGGAAAC CCTGATGCAG CGACGCCGCG TGAAGGATGA AGTATTTCGG TATGTAAACT TCTATCAGCA GGGAAGAAGA
S077-U-c           TGGGGGAAAC CCTGATGCAG CGACGCCGCG TGAAGGATGA AGTATTTCGG TATGTAAACT TCTATCAGCA GGGAAGAAGA
S077-U-d           TGGGGGAAAC CCTGATGCAG CGACGCCGCG TGAAGGATGA AGTATTTCGG TATGTAAACT TCTATCAGCA GGGAAGAAGA
VPI12708-a         TGGGGGAAAC CCTGATGCAG CGACGCCGCG TGAAGGATGA AGTATTTCGG TATGTAAACT TCTATCAGCA GGGAAGAAGA
VPI12708-b         TGGGGGAAAC CCTGATGCAG CGACGCCGCG TGAAGGATGA AGTATTTCGG TATGTAAACT TCTATCAGCA GGGAAGAAGA
VPI12708-c         TGGGGGAAAC CCTGATGCAG CGACGCCGCG TGAAGGATGA AGTATTTCGG TATGTAAACT TCTATCAGCA GGGAAGAAGA
VPI12708-d         TGGGGGAAAC CCTGATGCAG CGACGCCGCG TGAAGGATGA AGTATTTCGG TATGTAAACT TCTATCAGCA GGGAAGAAGA
DFI.1.130          TGGGGGAAAC CCTGATGCAG CGACGCCGCG TGAAGGATGA AGTATTTCGG TATGTAAACT TCTATCAGCA GGGAAGAAGA
DFI.1.161          TGGGGGAAAC CCTGATGCAG CGACGCCGCG TGAAGGATGA AGTATTTCGG TATGTAAACT TCTATCAGCA GGGAAGAAGA
DFI.1.162          TGGGGGAAAC CCTGATGCAG CGACGCCGCG TGAAGGATGA AGTATTTCGG TATGTAAACT TCTATCAGCA GGGAAGAAGA
DFI.1.217          TGGGGGAAAC CCTGATGCAG CGACGCCGCG TGAAGGATGA AGTATTTCGG TATGTAAACT TCTATCAGCA GGGAAGAAGA
DFI.1.234          TGGGGGAAAC CCTGATGCAG CGACGCCGCG TGAAGGATGA AGTATTTCGG TATGTAAACT TCTATCAGCA GGGAAGAAGA
DFI.1.60           TGGGGGAAAC CCTGATGCAG CGACGCCGCG TGAAGGATGA AGTATTTCGG TATGTAAACT TCTATCAGCA GGGAAGAAGA
DFI.4.63           TGGGGGAAAC CCTGATGCAG CGACGCCGCG TGAAGGATGA AGTATTTCGG TATGTAAACT TCTATCAGCA GGGAAGAAGA
JCM10419-a         TGGGGGAAAC CCTGATGCAG CGACGCCGCG TGAAGGATGA AGTATTTCGG TATGTAAACT TCTATCAGCA GGGAAGAAGA
JCM10419-b         TGGGGGAAAC CCTGATGCAG CGACGCCGCG TGAAGGATGA AGTATTTCGG TATGTAAACT TCTATCAGCA GGGAAGAAGA
JCM10419-c         TGGGGGAAAC CCTGATGCAG CGACGCCGCG TGAAGGATGA AGTATTTCGG TATGTAAACT TCTATCAGCA GGGAAGAAGA
JCM10419-d         TGGGGGAAAC CCTGATGCAG CGACGCCGCG TGAAGGATGA AGTATTTCGG TATGTAAACT TCTATCAGCA GGGAAGAAGA
JCM10420-a         TGGGGGAAAC CCTGATGCAG CGACGCCGCG TGAAGGATGA AGTATTTCGG TATGTAAACT TCTATCAGCA GGGAAGAAGA
JCM10420-b         TGGGGGAAAC CCTGATGCAG CGACGCCGCG TGAAGGATGA AGTATTTCGG TATGTAAACT TCTATCAGCA GGGAAGAAGA
JCM10420-c         TGGGGGAAAC CCTGATGCAG CGACGCCGCG TGAAGGATGA AGTATTTCGG TATGTAAACT TCTATCAGCA GGGAAGAAGA
JCM10420-d         TGGGGGAAAC CCTGATGCAG CGACGCCGCG TGAAGGATGA AGTATTTCGG TATGTAAACT TCTATCAGCA GGGAAGAAGA
JCM10423-a         TGGGGGAAAC CCTGATGCAG CGACGCCGCG TGAAGGATGA AGTATTTCGG TATGTAAACT TCTATCAGCA GGGAAGAAGA
JCM10423-b         TGGGGGAAAC CCTGATGCAG CGACGCCGCG TGAAGGATGA AGTATTTCGG TATGTAAACT TCTATCAGCA GGGAAGAAGA
JCM10423-c         TGGGGGAAAC CCTGATGCAG CGACGCCGCG TGAAGGATGA AGTATTTCGG TATGTAAACT TCTATCAGCA GGGAAGAAGA
JCM10423-d         TGGGGGAAAC CCTGATGCAG CGACGCCGCG TGAAGGATGA AGTATTTCGG TATGTAAACT TCTATCAGCA GGGAAGAAGA
MO32-a             TGGGGGAAAC CCTGATGCAG CGACGCCGCG TGAAGGATGA AGTATTTCGG TATGTAAACT TCTATCAGCA GGGAAGAAGA
MO32-b             TGGGGGAAAC CCTGATGCAG CGACGCCGCG TGAAGGATGA AGTATTTCGG TATGTAAACT TCTATCAGCA GGGAAGAAGA
MO32-c             TGGGGGAAAC CCTGATGCAG CGACGCCGCG TGAAGGATGA AGTATTTCGG TATGTAAACT TCTATCAGCA GGGAAGAAGA
MO32-d             TGGGGGAAAC CCTGATGCAG CGACGCCGCG TGAAGGATGA AGTATTTCGG TATGTAAACT TCTATCAGCA GGGAAGAAGA
MSK.5.24           TGGGGGAAAC CCTGATGCAG CGACGCCGCG TGAAGGATGA AGTATTTCGG TATGTAAACT TCTATCAGCA GGGAAGAAGA
NB2A-7-D5          TGGGGGAAAC CCTGATGCAG CGACGCCGCG TGAAGGATGA AGTATTTCGG TATGTAAACT TCTATCAGCA GGGAAGAAGA
NTI82-a            TGGGGGAAAC CCTGATGCAG CGACGCCGCG TGAAGGATGA AGTATTTCGG TATGTAAACT TCTATCAGCA GGGAAGAAGA
NTI82-b            TGGGGGAAAC CCTGATGCAG CGACGCCGCG TGAAGGATGA AGTATTTCGG TATGTAAACT TCTATCAGCA GGGAAGAAGA
NTI82-c            TGGGGGAAAC CCTGATGCAG CGACGCCGCG TGAAGGATGA AGTATTTCGG TATGTAAACT TCTATCAGCA GGGAAGAAGA
NTI82-d            TGGGGGAAAC CCTGATGCAG CGACGCCGCG TGAAGGATGA AGTATTTCGG TATGTAAACT TCTATCAGCA GGGAAGAAGA
Q4-a               TGGGGGAAAC CCTGATGCAG CGACGCCGCG TGAAGGATGA AGTATTTCGG TATGTAAACT TCTATCAGCA GGGAAGAAGA
Q4-b               TGGGGGAAAC CCTGATGCAG CGACGCCGCG TGAAGGATGA AGTATTTCGG TATGTAAACT TCTATCAGCA GGGAAGAAGA
Q4-c               TGGGGGAAAC CCTGATGCAG CGACGCCGCG TGAAGGATGA AGTATTTCGG TATGTAAACT TCTATCAGCA GGGAAGAAGA
Q4-d               TGGGGGAAAC CCTGATGCAG CGACGCCGCG TGAAGGATGA AGTATTTCGG TATGTAAACT TCTATCAGCA GGGAAGAAGA
S076-a             TGGGGGAAAC CCTGATGCAG CGACGCCGCG TGAAGGATGA AGTATTTCGG TATGTAAACT TCTATCAGCA GGGAAGAAGA
S076-b             TGGGGGAAAC CCTGATGCAG CGACGCCGCG TGAAGGATGA AGTATTTCGG TATGTAAACT TCTATCAGCA GGGAAGAAGA
S076-c             TGGGGGAAAC CCTGATGCAG CGACGCCGCG TGAAGGATGA AGTATTTCGG TATGTAAACT TCTATCAGCA GGGAAGAAGA
S076-d             TGGGGGAAAC CCTGATGCAG CGACGCCGCG TGAAGGATGA AGTATTTCGG TATGTAAACT TCTATCAGCA GGGAAGAAGA
SL.1.22            TGGGGGAAAC CCTGATGCAG CGACGCCGCG TGAAGGATGA AGTATTTCGG TATGTAAACT TCTATCAGCA GGGAAGAAGA
VE202-05           TGGGGGAAAC CCTGATGCAG CGACGCCGCG TGAAGGATGA AGTATTTCGG TATGTAAACT TCTATCAGCA GGGAAGAAGA
C.hylemonae        TGGGGGAAAC CCTGATGCAG CGACGCCGCG TGAAGGATGA AGTATTTCGG TATGTAAACT TCTATCAGCA GGGAAGAAGA
```

```
                    481
35704-a             TGACGGTACC TGACTAAGAA GCCCCGGCTA ACTACGTGCC AGCAGCCGCG GTAATACGTA GGGGGCAAGC GTTATCCGGA
35704-b             TGACGGTACC TGACTAAGAA GCCCCGGCTA ACTACGTGCC AGCAGCCGCG GTAATACGTA GGGGGCAAGC GTTATCCGGA
35704-c             TGACGGTACC TGACTAAGAA GCCCCGGCTA ACTACGTGCC AGCAGCCGCG GTAATACGTA GGGGGCAAGC GTTATCCGGA
35704-d             TGACGGTACC TGACTAAGAA GCCCCGGCTA ACTACGTGCC AGCAGCCGCG GTAATACGTA GGGGGCAAGC GTTATCCGGA
AM05-22             TGACGGTACC TGACTAAGAA GCCCCGGCTA ACTACGTGCC AGCAGCCGCG GTAATACGTA GGGGGCAAGC GTTATCCGGA
AM07-30             TGACGGTACC TGACTAAGAA GCCCCGGCTA ACTACGTGCC AGCAGCCGCG GTAATACGTA GGGGGCAAGC GTTATCCGGA
BL389WT3D-a         TGACGGTACC TGACTAAGAA GCCCCGGCTA ACTACGTGCC AGCAGCCGCG GTAATACGTA GGGGGCAAGC GTTATCCGGA
BL389WT3D-b         TGACGGTACC TGACTAAGAA GCCCCGGCTA ACTACGTGCC AGCAGCCGCG GTAATACGTA GGGGGCAAGC GTTATCCGGA
BL389WT3D-c         TGACGGTACC TGACTAAGAA GCCCCGGCTA ACTACGTGCC AGCAGCCGCG GTAATACGTA GGGGGCAAGC GTTATCCGGA
BL389WT3D-d         TGACGGTACC TGACTAAGAA GCCCCGGCTA ACTACGTGCC AGCAGCCGCG GTAATACGTA GGGGGCAAGC GTTATCCGGA
CE91-St59-a         TGACGGTACC TGACTAAGAA GCCCCGGCTA ACTACGTGCC AGCAGCCGCG GTAATACGTA GGGGGCAAGC GTTATCCGGA
CE91-St59-b         TGACGGTACC TGACTAAGAA GCCCCGGCTA ACTACGTGCC AGCAGCCGCG GTAATACGTA GGGGGCAAGC GTTATCCGGA
CE91-St59-c         TGACGGTACC TGACTAAGAA GCCCCGGCTA ACTACGTGCC AGCAGCCGCG GTAATACGTA GGGGGCAAGC GTTATCCGGA
CE91-St60-a         TGACGGTACC TGACTAAGAA GCCCCGGCTA ACTACGTGCC AGCAGCCGCG GTAATACGTA GGGGGCAAGC GTTATCCGGA
CE91-St60-b         TGACGGTACC TGACTAAGAA GCCCCGGCTA ACTACGTGCC AGCAGCCGCG GTAATACGTA GGGGGCAAGC GTTATCCGGA
CE91-St60-c         TGACGGTACC TGACTAAGAA GCCCCGGCTA ACTACGTGCC AGCAGCCGCG GTAATACGTA GGGGGCAAGC GTTATCCGGA
FDAARGOS_1227-a     TGACGGTACC TGACTAAGAA GCCCCGGCTA ACTACGTGCC AGCAGCCGCG GTAATACGTA GGGGGCAAGC GTTATCCGGA
FDAARGOS_1227-b     TGACGGTACC TGACTAAGAA GCCCCGGCTA ACTACGTGCC AGCAGCCGCG GTAATACGTA GGGGGCAAGC GTTATCCGGA
FDAARGOS_1227-c     TGACGGTACC TGACTAAGAA GCCCCGGCTA ACTACGTGCC AGCAGCCGCG GTAATACGTA GGGGGCAAGC GTTATCCGGA
FDAARGOS_1227-d     TGACGGTACC TGACTAAGAA GCCCCGGCTA ACTACGTGCC AGCAGCCGCG GTAATACGTA GGGGGCAAGC GTTATCCGGA
G10-a               TGACGGTACC TGACTAAGAA GCCCCGGCTA ACTACGTGCC AGCAGCCGCG GTAATACGTA GGGGGCAAGC GTTATCCGGA
G10-b               TGACGGTACC TGACTAAGAA GCCCCGGCTA ACTACGTGCC AGCAGCCGCG GTAATACGTA GGGGGCAAGC GTTATCCGGA
G10-c               TGACGGTACC TGACTAAGAA GCCCCGGCTA ACTACGTGCC AGCAGCCGCG GTAATACGTA GGGGGCAAGC GTTATCCGGA
G10-d               TGACGGTACC TGACTAAGAA GCCCCGGCTA ACTACGTGCC AGCAGCCGCG GTAATACGTA GGGGGCAAGC GTTATCCGGA
GGCC_0168           TGACGGTACC TGACTAAGAA GCCCCGGCTA ACTACGTGCC AGCAGCCGCG GTAATACGTA GGGGGCAAGC GTTATCCGGA
I10-a               TGACGGTACC TGACTAAGAA GCCCCGGCTA ACTACGTGCC AGCAGCCGCG GTAATACGTA GGGGGCAAGC GTTATCCGGA
I10-b               TGACGGTACC TGACTAAGAA GCCCCGGCTA ACTACGTGCC AGCAGCCGCG GTAATACGTA GGGGGCAAGC GTTATCCGGA
I10-c               TGACGGTACC TGACTAAGAA GCCCCGGCTA ACTACGTGCC AGCAGCCGCG GTAATACGTA GGGGGCAAGC GTTATCCGGA
I10-d               TGACGGTACC TGACTAAGAA GCCCCGGCTA ACTACGTGCC AGCAGCCGCG GTAATACGTA GGGGGCAAGC GTTATCCGGA
JCM10422-a          TGACGGTACC TGACTAAGAA GCCCCGGCTA ACTACGTGCC AGCAGCCGCG GTAATACGTA GGGGGCAAGC GTTATCCGGA
JCM10422-b          TGACGGTACC TGACTAAGAA GCCCCGGCTA ACTACGTGCC AGCAGCCGCG GTAATACGTA GGGGGCAAGC GTTATCCGGA
JCM10422-c          TGACGGTACC TGACTAAGAA GCCCCGGCTA ACTACGTGCC AGCAGCCGCG GTAATACGTA GGGGGCAAGC GTTATCCGGA
JCM10422-d          TGACGGTACC TGACTAAGAA GCCCCGGCTA ACTACGTGCC AGCAGCCGCG GTAATACGTA GGGGGCAAGC GTTATCCGGA
MGYG-HGUT-01303-a   TGACGGTACC TGACTAAGAA GCCCCGGCTA ACTACGTGCC AGCAGCCGCG GTAATACGTA GGGGGCAAGC GTTATCCGGA
MGYG-HGUT-01303-b   ---------- ---------- ---------- ---------- AGCAGCCGCG GTAATACGTA GGGGGCAAGC GTTATCCGGA
MSK.1.16            TGACGGTACC TGACTAAGAA GCCCCGGCTA ACTACGTGCC AGCAGCCGCG GTAATACGTA GGGGGCAAGC GTTATCCGGA
MSK.1.26            TGACGGTACC TGACTAAGAA GCCCCGGCTA ACTACGTGCC AGCAGCCGCG GTAATACGTA GGGGGCAAGC GTTATCCGGA
S077-U-a            TGACGGTACC TGACTAAGAA GCCCCGGCTA ACTACGTGCC AGCAGCCGCG GTAATACGTA GGGGGCAAGC GTTATCCGGA
S077-U-b            TGACGGTACC TGACTAAGAA GCCCCGGCTA ACTACGTGCC AGCAGCCGCG GTAATACGTA GGGGGCAAGC GTTATCCGGA
S077-U-c            TGACGGTACC TGACTAAGAA GCCCCGGCTA ACTACGTGCC AGCAGCCGCG GTAATACGTA GGGGGCAAGC GTTATCCGGA
S077-U-d            TGACGGTACC TGACTAAGAA GCCCCGGCTA ACTACGTGCC AGCAGCCGCG GTAATACGTA GGGGGCAAGC GTTATCCGGA
VPI12708-a          TGACGGTACC TGACTAAGAA GCCCCGGCTA ACTACGTGCC AGCAGCCGCG GTAATACGTA GGGGGCAAGC GTTATCCGGA
VPI12708-b          TGACGGTACC TGACTAAGAA GCCCCGGCTA ACTACGTGCC AGCAGCCGCG GTAATACGTA GGGGGCAAGC GTTATCCGGA
VPI12708-c          TGACGGTACC TGACTAAGAA GCCCCGGCTA ACTACGTGCC AGCAGCCGCG GTAATACGTA GGGGGCAAGC GTTATCCGGA
VPI12708-d          TGACGGTACC TGACTAAGAA GCCCCGGCTA ACTACGTGCC AGCAGCCGCG GTAATACGTA GGGGGCAAGC GTTATCCGGA
DFI.1.130           TGACGGTACC TGACTAAGAA GCCCCGGCTA ACTACGTGCC AGCAGCCGCG GTAATACGTA GGGGGCAAGC GTTATCCGGA
DFI.1.161           TGACGGTACC TGACTAAGAA GCCCCGGCTA ACTACGTGCC AGCAGCCGCG GTAATACGTA GGGGGCAAGC GTTATCCGGA
DFI.1.162           TGACGGTACC TGACTAAGAA GCCCCGGCTA ACTACGTGCC AGCAGCCGCG GTAATACGTA GGGGGCAAGC GTTATCCGGA
DFI.1.217           TGACGGTACC TGACTAAGAA GCCCCGGCTA ACTACGTGCC AGCAGCCGCG GTAATACGTA GGGGGCAAGC GTTATCCGGA
DFI.1.234           TGACGGTACC TGACTAAGAA GCCCCGGCTA ACTACGTGCC AGCAGCCGCG GTAATACGTA GGGGGCAAGC GTTATCCGGA
DFI.1.60            TGACGGTACC TGACTAAGAA GCCCCGGCTA ACTACGTGCC AGCAGCCGCG GTAATACGTA GGGGGCAAGC GTTATCCGGA
DFI.4.63            TGACGGTACC TGACTAAGAA GCCCCGGCTA ACTACGTGCC AGCAGCCGCG GTAATACGTA GGGGGCAAGC GTTATCCGGA
JCM10419-a          TGACGGTACC TGACTAAGAA GCCCCGGCTA ACTACGTGCC AGCAGCCGCG GTAATACGTA GGGGGCAAGC GTTATCCGGA
JCM10419-b          TGACGGTACC TGACTAAGAA GCCCCGGCTA ACTACGTGCC AGCAGCCGCG GTAATACGTA GGGGGCAAGC GTTATCCGGA
JCM10419-c          TGACGGTACC TGACTAAGAA GCCCCGGCTA ACTACGTGCC AGCAGCCGCG GTAATACGTA GGGGGCAAGC GTTATCCGGA
JCM10419-d          TGACGGTACC TGACTAAGAA GCCCCGGCTA ACTACGTGCC AGCAGCCGCG GTAATACGTA GGGGGCAAGC GTTATCCGGA
JCM10420-a          TGACGGTACC TGACTAAGAA GCCCCGGCTA ACTACGTGCC AGCAGCCGCG GTAATACGTA GGGGGCAAGC GTTATCCGGA
JCM10420-b          TGACGGTACC TGACTAAGAA GCCCCGGCTA ACTACGTGCC AGCAGCCGCG GTAATACGTA GGGGGCAAGC GTTATCCGGA
JCM10420-c          TGACGGTACC TGACTAAGAA GCCCCGGCTA ACTACGTGCC AGCAGCCGCG GTAATACGTA GGGGGCAAGC GTTATCCGGA
JCM10420-d          TGACGGTACC TGACTAAGAA GCCCCGGCTA ACTACGTGCC AGCAGCCGCG GTAATACGTA GGGGGCAAGC GTTATCCGGA
JCM10423-a          TGACGGTACC TGACTAAGAA GCCCCGGCTA ACTACGTGCC AGCAGCCGCG GTAATACGTA GGGGGCAAGC GTTATCCGGA
JCM10423-b          TGACGGTACC TGACTAAGAA GCCCCGGCTA ACTACGTGCC AGCAGCCGCG GTAATACGTA GGGGGCAAGC GTTATCCGGA
JCM10423-c          TGACGGTACC TGACTAAGAA GCCCCGGCTA ACTACGTGCC AGCAGCCGCG GTAATACGTA GGGGGCAAGC GTTATCCGGA
JCM10423-d          TGACGGTACC TGACTAAGAA GCCCCGGCTA ACTACGTGCC AGCAGCCGCG GTAATACGTA GGGGGCAAGC GTTATCCGGA
MO32-a              TGACGGTACC TGACTAAGAA GCCCCGGCTA ACTACGTGCC AGCAGCCGCG GTAATACGTA GGGGGCAAGC GTTATCCGGA
MO32-b              TGACGGTACC TGACTAAGAA GCCCCGGCTA ACTACGTGCC AGCAGCCGCG GTAATACGTA GGGGGCAAGC GTTATCCGGA
MO32-c              TGACGGTACC TGACTAAGAA GCCCCGGCTA ACTACGTGCC AGCAGCCGCG GTAATACGTA GGGGGCAAGC GTTATCCGGA
MO32-d              TGACGGTACC TGACTAAGAA GCCCCGGCTA ACTACGTGCC AGCAGCCGCG GTAATACGTA GGGGGCAAGC GTTATCCGGA
MSK.5.24            TGACGGTACC TGACTAAGAA GCCCCGGCTA ACTACGTGCC AGCAGCCGCG GTAATACGTA GGGGGCAAGC GTTATCCGGA
NB2A-7-D5           TGACGGTACC TGACTAAGAA GCCCCGGCTA ACTACGTGCC AGCAGCCGCG GTAATACGTA GGGGGCAAGC GTTATCCGGA
NTI82-a             TGACGGTACC TGACTAAGAA GCCCCGGCTA ACTACGTGCC AGCAGCCGCG GTAATACGTA GGGGGCAAGC GTTATCCGGA
NTI82-b             TGACGGTACC TGACTAAGAA GCCCCGGCTA ACTACGTGCC AGCAGCCGCG GTAATACGTA GGGGGCAAGC GTTATCCGGA
NTI82-c             TGACGGTACC TGACTAAGAA GCCCCGGCTA ACTACGTGCC AGCAGCCGCG GTAATACGTA GGGGGCAAGC GTTATCCGGA
NTI82-d             TGACGGTACC TGACTAAGAA GCCCCGGCTA ACTACGTGCC AGCAGCCGCG GTAATACGTA GGGGGCAAGC GTTATCCGGA
Q4-a                TGACGGTACC TGACTAAGAA GCCCCGGCTA ACTACGTGCC AGCAGCCGCG GTAATACGTA GGGGGCAAGC GTTATCCGGA
Q4-b                TGACGGTACC TGACTAAGAA GCCCCGGCTA ACTACGTGCC AGCAGCCGCG GTAATACGTA GGGGGCAAGC GTTATCCGGA
Q4-c                TGACGGTACC TGACTAAGAA GCCCCGGCTA ACTACGTGCC AGCAGCCGCG GTAATACGTA GGGGGCAAGC GTTATCCGGA
Q4-d                TGACGGTACC TGACTAAGAA GCCCCGGCTA ACTACGTGCC AGCAGCCGCG GTAATACGTA GGGGGCAAGC GTTATCCGGA
S076-a              TGACGGTACC TGACTAAGAA GCCCCGGCTA ACTACGTGCC AGCAGCCGCG GTAATACGTA GGGGGCAAGC GTTATCCGGA
S076-b              TGACGGTACC TGACTAAGAA GCCCCGGCTA ACTACGTGCC AGCAGCCGCG GTAATACGTA GGGGGCAAGC GTTATCCGGA
S076-c              TGACGGTACC TGACTAAGAA GCCCCGGCTA ACTACGTGCC AGCAGCCGCG GTAATACGTA GGGGGCAAGC GTTATCCGGA
S076-d              TGACGGTACC TGACTAAGAA GCCCCGGCTA ACTACGTGCC AGCAGCCGCG GTAATACGTA GGGGGCAAGC GTTATCCGGA
SL.1.22             TGACGGTACC TGACTAAGAA GCCCCGGCTA ACTACGTGCC AGCAGCCGCG GTAATACGTA GGGGGCAAGC GTTATCCGGA
VE202-05            TGACGGTACC TGACTAAGAA GCCCCGGCTA ACTACGTGCC AGCAGCCGCG GTAATACGTA GGGGGCAAGC GTTATCCGGA
C.hylemonae         TGACGGTACC TGACTAAGAA GCCCCGGCTA ACTACGTGCC AGCAGCCGCG GTAATACGTA GGGGGCAAGC GTTATCCGGA
```

```
                        561
35704-a                 TTTACTGGGT GTAAAGGGAG CGTAGACGGC GATGCAAGCC AGATGTGAAA GCCCGGGGCT CAACCCCGGG ACTGCATTTG
35704-b                 TTTACTGGGT GTAAAGGGAG CGTAGACGGC GATGCAAGCC AGATGTGAAA GCCCGGGGCT CAACCCCGGG ACTGCATTTG
35704-c                 TTTACTGGGT GTAAAGGGAG CGTAGACGGC GATGCAAGCC AGATGTGAAA GCCCGGGGCT CAACCCCGGG ACTGCATTTG
35704-d                 TTTACTGGGT GTAAAGGGAG CGTAGACGGC GATGCAAGCC AGATGTGAAA GCCCGGGGCT CAACCCCGGG ACTGCATTTG
AM05-22                 TTTACTGGGT GTAAAGGGAG CGTAGACGGC GATGCAAGCC AGATGTGAAA GCCCGGGGCT CAACCCCGGG ACTGCATTTG
AM07-30                 TTTACTGGGT GTAAAGGGAG CGTAGACGGC GATGCAAGCC AGATGTGAAA GCCCGGGGCT CAACCCCGGG ACTGCATTTG
BL389WT3D-a             TTTACTGGGT GTAAAGGGAG CGTAGACGGC GATGCAAGCC AGATGTGAAA GCCCGGGGCT CAACCCCGGG ACTGCATTTG
BL389WT3D-b             TTTACTGGGT GTAAAGGGAG CGTAGACGGC GATGCAAGCC AGATGTGAAA GCCCGGGGCT CAACCCCGGG ACTGCATTTG
BL389WT3D-c             TTTACTGGGT GTAAAGGGAG CGTAGACGGC GATGCAAGCC AGATGTGAAA GCCCGGGGCT CAACCCCGGG ACTGCATTTG
BL389WT3D-d             TTTACTGGGT GTAAAGGGAG CGTAGACGGC GATGCAAGCC AGATGTGAAA GCCCGGGGCT CAACCCCGGG ACTGCATTTG
CE91-St59-a             TTTACTGGGT GTAAAGGGAG CGTAGACGGC GATGCAAGCC AGATGTGAAA GCCCGGGGCT CAACCCCGGG ACTGCATTTG
CE91-St59-b             TTTACTGGGT GTAAAGGGAG CGTAGACGGC GATGCAAGCC AGATGTGAAA GCCCGGGGCT CAACCCCGGG ACTGCATTTG
CE91-St59-c             TTTACTGGGT GTAAAGGGAG CGTAGACGGC GATGCAAGCC AGATGTGAAA GCCCGGGGCT CAACCCCGGG ACTGCATTTG
CE91-St60-a             TTTACTGGGT GTAAAGGGAG CGTAGACGGC GATGCAAGCC AGATGTGAAA GCCCGGGGCT CAACCCCGGG ACTGCATTTG
CE91-St60-b             TTTACTGGGT GTAAAGGGAG CGTAGACGGC GATGCAAGCC AGATGTGAAA GCCCGGGGCT CAACCCCGGG ACTGCATTTG
CE91-St60-c             TTTACTGGGT GTAAAGGGAG CGTAGACGGC GATGCAAGCC AGATGTGAAA GCCCGGGGCT CAACCCCGGG ACTGCATTTG
FDAARGOS_1227-a         TTTACTGGGT GTAAAGGGAG CGTAGACGGC GATGCAAGCC AGATGTGAAA GCCCGGGGCT CAACCCCGGG ACTGCATTTG
FDAARGOS_1227-b         TTTACTGGGT GTAAAGGGAG CGTAGACGGC GATGCAAGCC AGATGTGAAA GCCCGGGGCT CAACCCCGGG ACTGCATTTG
FDAARGOS_1227-c         TTTACTGGGT GTAAAGGGAG CGTAGACGGC GATGCAAGCC AGATGTGAAA GCCCGGGGCT CAACCCCGGG ACTGCATTTG
FDAARGOS_1227-d         TTTACTGGGT GTAAAGGGAG CGTAGACGGC GATGCAAGCC AGATGTGAAA GCCCGGGGCT CAACCCCGGG ACTGCATTTG
G10-a                   TTTACTGGGT GTAAAGGGAG CGTAGACGGC GATGCAAGCC AGATGTGAAA GCCCGGGGCT CAACCCCGGG ACTGCATTTG
G10-b                   TTTACTGGGT GTAAAGGGAG CGTAGACGGC GATGCAAGCC AGATGTGAAA GCCCGGGGCT CAACCCCGGG ACTGCATTTG
G10-c                   TTTACTGGGT GTAAAGGGAG CGTAGACGGC GATGCAAGCC AGATGTGAAA GCCCGGGGCT CAACCCCGGG ACTGCATTTG
G10-d                   TTTACTGGGT GTAAAGGGAG CGTAGACGGC GATGCAAGCC AGATGTGAAA GCCCGGGGCT CAACCCCGGG ACTGCATTTG
GGCC_0168               TTTACTGGGT GTAAAGGGAG CGTAGACGGC GATGCAAGCC AGATGTGAAA GCCCGGGGCT CAACCCCGGG ACTGCATTTG
I10-a                   TTTACTGGGT GTAAAGGGAG CGTAGACGGC GATGCAAGCC AGATGTGAAA GCCCGGGGCT CAACCCCGGG ACTGCATTTG
I10-b                   TTTACTGGGT GTAAAGGGAG CGTAGACGGC GATGCAAGCC AGATGTGAAA GCCCGGGGCT CAACCCCGGG ACTGCATTTG
I10-c                   TTTACTGGGT GTAAAGGGAG CGTAGACGGC GATGCAAGCC AGATGTGAAA GCCCGGGGCT CAACCCCGGG ACTGCATTTG
I10-d                   TTTACTGGGT GTAAAGGGAG CGTAGACGGC GATGCAAGCC AGATGTGAAA GCCCGGGGCT CAACCCCGGG ACTGCATTTG
JCM10422-a              TTTACTGGGT GTAAAGGGAG CGTAGACGGC GATGCAAGCC AGATGTGAAA GCCCGGGGCT CAACCCCGGG ACTGCATTTG
JCM10422-b              TTTACTGGGT GTAAAGGGAG CGTAGACGGC GATGCAAGCC AGATGTGAAA GCCCGGGGCT CAACCCCGGG ACTGCATTTG
JCM10422-c              TTTACTGGGT GTAAAGGGAG CGTAGACGGC GATGCAAGCC AGATGTGAAA GCCCGGGGCT CAACCCCGGG ACTGCATTTG
JCM10422-d              TTTACTGGGT GTAAAGGGAG CGTAGACGGC GATGCAAGCC AGATGTGAAA GCCCGGGGCT CAACCCCGGG ACTGCATTTG
MGYG-HGUT-01303-a       TTTACTGGGT GTAAAGGGAG CGTAGACGGC GATGCAAGCC AGATGTGAAA GCCCGGGGCT CAACCCCGGG ACTGCATTTG
MGYG-HGUT-01303-b       TTTACTGGGT GTAAAGGGAG CGTAGACGGC GATGCAAGCC AGATGTGAAA GCCCGGGGCT CAACCCCGGG ACTGCATTTG
MSK.1.16                TTTACTGGGT GTAAAGGGAG CGTAGACGGC GATGCAAGCC AGATGTGAAA GCCCGGGGCT CAACCCCGGG ACTGCATTTG
MSK.1.26                TTTACTGGGT GTAAAGGGAG CGTAGACGGC GATGCAAGCC AGATGTGAAA GCCCGGGGCT CAACCCCGGG ACTGCATTTG
S077-U-a                TTTACTGGGT GTAAAGGGAG CGTAGACGGC GATGCAAGCC AGATGTGAAA GCCCGGGGCT CAACCCCGGG ACTGCATTTG
S077-U-b                TTTACTGGGT GTAAAGGGAG CGTAGACGGC GATGCAAGCC AGATGTGAAA GCCCGGGGCT CAACCCCGGG ACTGCATTTG
S077-U-c                TTTACTGGGT GTAAAGGGAG CGTAGACGGC GATGCAAGCC AGATGTGAAA GCCCGGGGCT CAACCCCGGG ACTGCATTTG
S077-U-d                TTTACTGGGT GTAAAGGGAG CGTAGACGGC GATGCAAGCC AGATGTGAAA GCCCGGGGCT CAACCCCGGG ACTGCATTTG
VPI12708-a              TTTACTGGGT GTAAAGGGAG CGTAGACGGC GATGCAAGCC AGATGTGAAA GCCCGGGGCT CAACCCCGGG ACTGCATTTG
VPI12708-b              TTTACTGGGT GTAAAGGGAG CGTAGACGGC GATGCAAGCC AGATGTGAAA GCCCGGGGCT CAACCCCGGG ACTGCATTTG
VPI12708-c              TTTACTGGGT GTAAAGGGAG CGTAGACGGC GATGCAAGCC AGATGTGAAA GCCCGGGGCT CAACCCCGGG ACTGCATTTG
VPI12708-d              TTTACTGGGT GTAAAGGGAG CGTAGACGGC GATGCAAGCC AGATGTGAAA GCCCGGGGCT CAACCCCGGG ACTGCATTTG
DFI.1.130               TTTACTGGGT GTAAAGGGAG CGTAGACGGC GATGCAAGCC AGATGTGAAA GCCCGGGGCT CAACCCCGGG ACTGCATTTG
DFI.1.161               TTTACTGGGT GTAAAGGGAG CGTAGACGGC GATGCAAGCC AGATGTGAAA GCCCGGGGCT CAACCCCGGG ACTGCATTTG
DFI.1.162               TTTACTGGGT GTAAAGGGAG CGTAGACGGC GATGCAAGCC AGATGTGAAA GCCCGGGGCT CAACCCCGGG ACTGCATTTG
DFI.1.217               TTTACTGGGT GTAAAGGGAG CGTAGACGGC GATGCAAGCC AGATGTGAAA GCCCGGGGCT CAACCCCGGG ACTGCATTTG
DFI.1.234               TTTACTGGGT GTAAAGGGAG CGTAGACGGC GATGCAAGCC AGATGTGAAA GCCCGGGGCT CAACCCCGGG ACTGCATTTG
DFI.1.60                TTTACTGGGT GTAAAGGGAG CGTAGACGGC GATGCAAGCC AGATGTGAAA GCCCGGGGCT CAACCCCGGG ACTGCATTTG
DFI.4.63                TTTACTGGGT GTAAAGGGAG CGTAGACGGC GATGCAAGCC AGATGTGAAA GCCCGGGGCT CAACCCCGGG ACTGCATTTG
JCM10419-a              TTTACTGGGT GTAAAGGGAG CGTAGACGGC GATGCAAGCC AGATGTGAAA GCCCGGGGCT CAACCCCGGG ACTGCATTTG
JCM10419-b              TTTACTGGGT GTAAAGGGAG CGTAGACGGC GATGCAAGCC AGATGTGAAA GCCCGGGGCT CAACCCCGGG ACTGCATTTG
JCM10419-c              TTTACTGGGT GTAAAGGGAG CGTAGACGGC GATGCAAGCC AGATGTGAAA GCCCGGGGCT CAACCCCGGG ACTGCATTTG
JCM10419-d              TTTACTGGGT GTAAAGGGAG CGTAGACGGC GATGCAAGCC AGATGTGAAA GCCCGGGGCT CAACCCCGGG ACTGCATTTG
JCM10420-a              TTTACTGGGT GTAAAGGGAG CGTAGACGGC GATGCAAGCC AGATGTGAAA GCCCGGGGCT CAACCCCGGG ACTGCATTTG
JCM10420-b              TTTACTGGGT GTAAAGGGAG CGTAGACGGC GATGCAAGCC AGATGTGAAA GCCCGGGGCT CAACCCCGGG ACTGCATTTG
JCM10420-c              TTTACTGGGT GTAAAGGGAG CGTAGACGGC GATGCAAGCC AGATGTGAAA GCCCGGGGCT CAACCCCGGG ACTGCATTTG
JCM10420-d              TTTACTGGGT GTAAAGGGAG CGTAGACGGC GATGCAAGCC AGATGTGAAA GCCCGGGGCT CAACCCCGGG ACTGCATTTG
JCM10423-a              TTTACTGGGT GTAAAGGGAG CGTAGACGGC GATGCAAGCC AGATGTGAAA GCCCGGGGCT CAACCCCGGG ACTGCATTTG
JCM10423-b              TTTACTGGGT GTAAAGGGAG CGTAGACGGC GATGCAAGCC AGATGTGAAA GCCCGGGGCT CAACCCCGGG ACTGCATTTG
JCM10423-c              TTTACTGGGT GTAAAGGGAG CGTAGACGGC GATGCAAGCC AGATGTGAAA GCCCGGGGCT CAACCCCGGG ACTGCATTTG
JCM10423-d              TTTACTGGGT GTAAAGGGAG CGTAGACGGC GATGCAAGCC AGATGTGAAA GCCCGGGGCT CAACCCCGGG ACTGCATTTG
MO32-a                  TTTACTGGGT GTAAAGGGAG CGTAGACGGC GATGCAAGCC AGATGTGAAA GCCCGGGGCT CAACCCCGGG ACTGCATTTG
MO32-b                  TTTACTGGGT GTAAAGGGAG CGTAGACGGC GATGCAAGCC AGATGTGAAA GCCCGGGGCT CAACCCCGGG ACTGCATTTG
MO32-c                  TTTACTGGGT GTAAAGGGAG CGTAGACGGC GATGCAAGCC AGATGTGAAA GCCCGGGGCT CAACCCCGGG ACTGCATTTG
MO32-d                  TTTACTGGGT GTAAAGGGAG CGTAGACGGC GATGCAAGCC AGATGTGAAA GCCCGGGGCT CAACCCCGGG ACTGCATTTG
MSK.5.24                TTTACTGGGT GTAAAGGGAG CGTAGACGGC GATGCAAGCC AGATGTGAAA GCCCGGGGCT CAACCCCGGG ACTGCATTTG
NB2A-7-D5               TTTACTGGGT GTAAAGGGAG CGTAGACGGC GATGCAAGCC AGATGTGAAA GCCCGGGGCT CAACCCCGGG ACTGCATTTG
NTI82-a                 TTTACTGGGT GTAAAGGGAG CGTAGACGGC GATGCAAGCC AGATGTGAAA GCCCGGGGCT CAACCCCGGG ACTGCATTTG
NTI82-b                 TTTACTGGGT GTAAAGGGAG CGTAGACGGC GATGCAAGCC AGATGTGAAA GCCCGGGGCT CAACCCCGGG ACTGCATTTG
NTI82-c                 TTTACTGGGT GTAAAGGGAG CGTAGACGGC GATGCAAGCC AGATGTGAAA GCCCGGGGCT CAACCCCGGG ACTGCATTTG
NTI82-d                 TTTACTGGGT GTAAAGGGAG CGTAGACGGC GATGCAAGCC AGATGTGAAA GCCCGGGGCT CAACCCCGGG ACTGCATTTG
Q4-a                    TTTACTGGGT GTAAAGGGAG CGTAGACGGC GATGCAAGCC AGATGTGAAA GCCCGGGGCT CAACCCCGGG ACTGCATTTG
Q4-b                    TTTACTGGGT GTAAAGGGAG CGTAGACGGC GATGCAAGCC AGATGTGAAA GCCCGGGGCT CAACCCCGGG ACTGCATTTG
Q4-c                    TTTACTGGGT GTAAAGGGAG CGTAGACGGC GATGCAAGCC AGATGTGAAA GCCCGGGGCT CAACCCCGGG ACTGCATTTG
Q4-d                    TTTACTGGGT GTAAAGGGAG CGTAGACGGC GATGCAAGCC AGATGTGAAA GCCCGGGGCT CAACCCCGGG ACTGCATTTG
S076-a                  TTTACTGGGT GTAAAGGGAG CGTAGACGGC GATGCAAGCC AGATGTGAAA GCCCGGGGCT CAACCCCGGG ACTGCATTTG
S076-b                  TTTACTGGGT GTAAAGGGAG CGTAGACGGC GATGCAAGCC AGATGTGAAA GCCCGGGGCT CAACCCCGGG ACTGCATTTG
S076-c                  TTTACTGGGT GTAAAGGGAG CGTAGACGGC GATGCAAGCC AGATGTGAAA GCCCGGGGCT CAACCCCGGG ACTGCATTTG
S076-d                  TTTACTGGGT GTAAAGGGAG CGTAGACGGC GATGCAAGCC AGATGTGAAA GCCCGGGGCT CAACCCCGGG ACTGCATTTG
SL.1.22                 TTTACTGGGT GTAAAGGGAG CGTAGACGGC GATGCAAGCC AGATGTGAAA GCCCGGGGCT CAACCCCGGG ACTGCATTTG
VE202-05                TTTACTGGGT GTAAAGGGAG CGTAGACGGC GATGCAAGCC AGATGTGAAA GCCCGGGGCT CAACCCCGGG ACTGCATTTG
C.hylemonae             TTTACTGGGT GTAAAGGGAG CGTAGACGGC ATGGCAAGTC TGAAGTGAAA GCCCGGGGCT CAACCCCGGG ACTGCTTTGG
```

```
                              641
35704-a                       GAACTGCGTG GCTGGAGTGT CGGAGAGGCA GGCGGAATTC CTAGTGTAGC GGTGAAATGC GTAGATATTA GGAGGAACAC
35704-b                       GAACTGCGTG GCTGGAGTGT CGGAGAGGCA GGCGGAATTC CTAGTGTAGC GGTGAAATGC GTAGATATTA GGAGGAACAC
35704-c                       GAACTGCGTG GCTGGAGTGT CGGAGAGGCA GGCGGAATTC CTAGTGTAGC GGTGAAATGC GTAGATATTA GGAGGAACAC
35704-d                       GAACTGCGTG GCTGGAGTGT CGGAGAGGCA GGCGGAATTC CTAGTGTAGC GGTGAAATGC GTAGATATTA GGAGGAACAC
AM05-22                       GAACTGCGTG GCTGGAGTGT CGGAGAGGCA GGCGGAATTC CTAGTGTAGC GGTGAAATGC GTAGATATTA GGAGGAACAC
AM07-30                       GAACTGCGTG GCTGGAGTGT CGGAGAGGCA GGCGGAATTC CTAGTGTAGC GGTGAAATGC GTAGATATTA GGAGGAACAC
BL389WT3D-a                   GAACTGCGTG GCTGGAGTGT CGGAGAGGCA GGCGGAATTC CTAGTGTAGC GGTGAAATGC GTAGATATTA GGAGGAACAC
BL389WT3D-b                   GAACTGCGTG GCTGGAGTGT CGGAGAGGCA GGCGGAATTC CTAGTGTAGC GGTGAAATGC GTAGATATTA GGAGGAACAC
BL389WT3D-c                   GAACTGCGTG GCTGGAGTGT CGGAGAGGCA GGCGGAATTC CTAGTGTAGC GGTGAAATGC GTAGATATTA GGAGGAACAC
BL389WT3D-d                   GAACTGCGTG GCTGGAGTGT CGGAGAGGCA GGCGGAATTC CTAGTGTAGC GGTGAAATGC GTAGATATTA GGAGGAACAC
CE91-St59-a                   GAACTGCGTG GCTGGAGTGT CGGAGAGGCA GGCGGAATTC CTAGTGTAGC GGTGAAATGC GTAGATATTA GGAGGAACAC
CE91-St59-b                   GAACTGCGTG GCTGGAGTGT CGGAGAGGCA GGCGGAATTC CTAGTGTAGC GGTGAAATGC GTAGATATTA GGAGGAACAC
CE91-St59-c                   GAACTGCGTG GCTGGAGTGT CGGAGAGGCA GGCGGAATTC CTAGTGTAGC GGTGAAATGC GTAGATATTA GGAGGAACAC
CE91-St60-a                   GAACTGCGTG GCTGGAGTGT CGGAGAGGCA GGCGGAATTC CTAGTGTAGC GGTGAAATGC GTAGATATTA GGAGGAACAC
CE91-St60-b                   GAACTGCGTG GCTGGAGTGT CGGAGAGGCA GGCGGAATTC CTAGTGTAGC GGTGAAATGC GTAGATATTA GGAGGAACAC
CE91-St60-c                   GAACTGCGTG GCTGGAGTGT CGGAGAGGCA GGCGGAATTC CTAGTGTAGC GGTGAAATGC GTAGATATTA GGAGGAACAC
FDAARGOS_1227-a               GAACTGCGTG GCTGGAGTGT CGGAGAGGCA GGCGGAATTC CTAGTGTAGC GGTGAAATGC GTAGATATTA GGAGGAACAC
FDAARGOS_1227-b               GAACTGCGTG GCTGGAGTGT CGGAGAGGCA GGCGGAATTC CTAGTGTAGC GGTGAAATGC GTAGATATTA GGAGGAACAC
FDAARGOS_1227-c               GAACTGCGTG GCTGGAGTGT CGGAGAGGCA GGCGGAATTC CTAGTGTAGC GGTGAAATGC GTAGATATTA GGAGGAACAC
FDAARGOS_1227-d               GAACTGCGTG GCTGGAGTGT CGGAGAGGCA GGCGGAATTC CTAGTGTAGC GGTGAAATGC GTAGATATTA GGAGGAACAC
G10-a                         GAACTGCGTG GCTGGAGTGT CGGAGAGGCA GGCGGAATTC CTAGTGTAGC GGTGAAATGC GTAGATATTA GGAGGAACAC
G10-b                         GAACTGCGTG GCTGGAGTGT CGGAGAGGCA GGCGGAATTC CTAGTGTAGC GGTGAAATGC GTAGATATTA GGAGGAACAC
G10-c                         GAACTGCGTG GCTGGAGTGT CGGAGAGGCA GGCGGAATTC CTAGTGTAGC GGTGAAATGC GTAGATATTA GGAGGAACAC
G10-d                         GAACTGCGTG GCTGGAGTGT CGGAGAGGCA GGCGGAATTC CTAGTGTAGC GGTGAAATGC GTAGATATTA GGAGGAACAC
GGCC_0168                     GAACTGCGTG GCTGGAGTGT CGGAGAGGCA GGCGGAATTC CTAGTGTAGC GGTGAAATGC GTAGATATTA GGAGGAACAC
I10-a                         GAACTGCGTG GCTGGAGTGT CGGAGAGGCA GGCGGAATTC CTAGTGTAGC GGTGAAATGC GTAGATATTA GGAGGAACAC
I10-b                         GAACTGCGTG GCTGGAGTGT CGGAGAGGCA GGCGGAATTC CTAGTGTAGC GGTGAAATGC GTAGATATTA GGAGGAACAC
I10-c                         GAACTGCGTG GCTGGAGTGT CGGAGAGGCA GGCGGAATTC CTAGTGTAGC GGTGAAATGC GTAGATATTA GGAGGAACAC
I10-d                         GAACTGCGTG GCTGGAGTGT CGGAGAGGCA GGCGGAATTC CTAGTGTAGC GGTGAAATGC GTAGATATTA GGAGGAACAC
JCM10422-a                    GAACTGCGTG GCTGGAGTGT CGGAGAGGCA GGCGGAATTC CTAGTGTAGC GGTGAAATGC GTAGATATTA GGAGGAACAC
JCM10422-b                    GAACTGCGTG GCTGGAGTGT CGGAGAGGCA GGCGGAATTC CTAGTGTAGC GGTGAAATGC GTAGATATTA GGAGGAACAC
JCM10422-c                    GAACTGCGTG GCTGGAGTGT CGGAGAGGCA GGCGGAATTC CTAGTGTAGC GGTGAAATGC GTAGATATTA GGAGGAACAC
JCM10422-d                    GAACTGCGTG GCTGGAGTGT CGGAGAGGCA GGCGGAATTC CTAGTGTAGC GGTGAAATGC GTAGATATTA GGAGGAACAC
MGYG-HGUT-01303-a             GAACTGCGTG GCTGGAGTGT CGGAGAGGCA GGCGGAATTC CTAGTGTAGC GGTGAAATGC GTAGATATTA GGAGGAACAC
MGYG-HGUT-01303-b             GAACTGCGTG GCTGGAGTGT CGGAGAGGCA GGCGGAATTC CTAGTGTAGC GGTGAAATGC GTAGATATTA GGAGGAACAC
MSK.1.16                      GAACTGCGTG GCTGGAGTGT CGGAGAGGCA GGCGGAATTC CTAGTGTAGC GGTGAAATGC GTAGATATTA GGAGGAACAC
MSK.1.26                      GAACTGCGTG GCTGGAGTGT CGGAGAGGCA GGCGGAATTC CTAGTGTAGC GGTGAAATGC GTAGATATTA GGAGGAACAC
S077-U-a                      GAACTGCGTG GCTGGAGTGT CGGAGAGGCA GGCGGAATTC CTAGTGTAGC GGTGAAATGC GTAGATATTA GGAGGAACAC
S077-U-b                      GAACTGCGTG GCTGGAGTGT CGGAGAGGCA GGCGGAATTC CTAGTGTAGC GGTGAAATGC GTAGATATTA GGAGGAACAC
S077-U-c                      GAACTGCGTG GCTGGAGTGT CGGAGAGGCA GGCGGAATTC CTAGTGTAGC GGTGAAATGC GTAGATATTA GGAGGAACAC
S077-U-d                      GAACTGCGTG GCTGGAGTGT CGGAGAGGCA GGCGGAATTC CTAGTGTAGC GGTGAAATGC GTAGATATTA GGAGGAACAC
VPI12708-a                    GAACTGCGTG GCTGGAGTGT CGGAGAGGCA GGCGGAATTC CTAGTGTAGC GGTGAAATGC GTAGATATTA GGAGGAACAC
VPI12708-b                    GAACTGCGTG GCTGGAGTGT CGGAGAGGCA GGCGGAATTC CTAGTGTAGC GGTGAAATGC GTAGATATTA GGAGGAACAC
VPI12708-c                    GAACTGCGTG GCTGGAGTGT CGGAGAGGCA GGCGGAATTC CTAGTGTAGC GGTGAAATGC GTAGATATTA GGAGGAACAC
VPI12708-d                    GAACTGCGTG GCTGGAGTGT CGGAGAGGCA GGCGGAATTC CTAGTGTAGC GGTGAAATGC GTAGATATTA GGAGGAACAC
DFI.1.130                     GAACTGCGTG GCTGGAGTGT CGGAGAGGCA GGCGGAATTC CTAGTGTAGC GGTGAAATGC GTAGATATTA GGAGGAACAC
DFI.1.161                     GAACTGCGTG GCTGGAGTGT CGGAGAGGCA GGCGGAATTC CTAGTGTAGC GGTGAAATGC GTAGATATTA GGAGGAACAC
DFI.1.162                     GAACTGCGTG GCTGGAGTGT CGGAGAGGCA GGCGGAATTC CTAGTGTAGC GGTGAAATGC GTAGATATTA GGAGGAACAC
DFI.1.217                     GAACTGCGTG GCTGGAGTGT CGGAGAGGCA GGCGGAATTC CTAGTGTAGC GGTGAAATGC GTAGATATTA GGAGGAACAC
DFI.1.234                     GAACTGCGTG GCTGGAGTGT CGGAGAGGCA GGCGGAATTC CTAGTGTAGC GGTGAAATGC GTAGATATTA GGAGGAACAC
DFI.1.60                      GAACTGCGTG GCTGGAGTGT CGGAGAGGCA GGCGGAATTC CTAGTGTAGC GGTGAAATGC GTAGATATTA GGAGGAACAC
DFI.4.63                      GAACTGCGTG GCTGGAGTGT CGGAGAGGCA GGCGGAATTC CTAGTGTAGC GGTGAAATGC GTAGATATTA GGAGGAACAC
JCM10419-a                    GAACTGCGTG GCTGGAGTGT CGGAGAGGCA GGCGGAATTC CTAGTGTAGC GGTGAAATGC GTAGATATTA GGAGGAACAC
JCM10419-b                    GAACTGCGTG GCTGGAGTGT CGGAGAGGCA GGCGGAATTC CTAGTGTAGC GGTGAAATGC GTAGATATTA GGAGGAACAC
JCM10419-c                    GAACTGCGTG GCTGGAGTGT CGGAGAGGCA GGCGGAATTC CTAGTGTAGC GGTGAAATGC GTAGATATTA GGAGGAACAC
JCM10419-d                    GAACTGCGTG GCTGGAGTGT CGGAGAGGCA GGCGGAATTC CTAGTGTAGC GGTGAAATGC GTAGATATTA GGAGGAACAC
JCM10420-a                    GAACTGCGTG GCTGGAGTGT CGGAGAGGCA GGCGGAATTC CTAGTGTAGC GGTGAAATGC GTAGATATTA GGAGGAACAC
JCM10420-b                    GAACTGCGTG GCTGGAGTGT CGGAGAGGCA GGCGGAATTC CTAGTGTAGC GGTGAAATGC GTAGATATTA GGAGGAACAC
JCM10420-c                    GAACTGCGTG GCTGGAGTGT CGGAGAGGCA GGCGGAATTC CTAGTGTAGC GGTGAAATGC GTAGATATTA GGAGGAACAC
JCM10420-d                    GAACTGCGTG GCTGGAGTGT CGGAGAGGCA GGCGGAATTC CTAGTGTAGC GGTGAAATGC GTAGATATTA GGAGGAACAC
JCM10423-a                    GAACTGCGTG GCTGGAGTGT CGGAGAGGCA GGCGGAATTC CTAGTGTAGC GGTGAAATGC GTAGATATTA GGAGGAACAC
JCM10423-b                    GAACTGCGTG GCTGGAGTGT CGGAGAGGCA GGCGGAATTC CTAGTGTAGC GGTGAAATGC GTAGATATTA GGAGGAACAC
JCM10423-c                    GAACTGCGTG GCTGGAGTGT CGGAGAGGCA GGCGGAATTC CTAGTGTAGC GGTGAAATGC GTAGATATTA GGAGGAACAC
JCM10423-d                    GAACTGCGTG GCTGGAGTGT CGGAGAGGCA GGCGGAATTC CTAGTGTAGC GGTGAAATGC GTAGATATTA GGAGGAACAC
MO32-a                        GAACTGCGTG GCTGGAGTGT CGGAGAGGCA GGCGGAATTC CTAGTGTAGC GGTGAAATGC GTAGATATTA GGAGGAACAC
MO32-b                        GAACTGCGTG GCTGGAGTGT CGGAGAGGCA GGCGGAATTC CTAGTGTAGC GGTGAAATGC GTAGATATTA GGAGGAACAC
MO32-c                        GAACTGCGTG GCTGGAGTGT CGGAGAGGCA GGCGGAATTC CTAGTGTAGC GGTGAAATGC GTAGATATTA GGAGGAACAC
MO32-d                        GAACTGCGTG GCTGGAGTGT CGGAGAGGCA GGCGGAATTC CTAGTGTAGC GGTGAAATGC GTAGATATTA GGAGGAACAC
MSK.5.24                      GAACTGCGTG GCTGGAGTGT CGGAGAGGCA GGCGGAATTC CTAGTGTAGC GGTGAAATGC GTAGATATTA GGAGGAACAC
NB2A-7-D5                     GAACTGCGTG GCTGGAGTGT CGGAGAGGCA GGCGGAATTC CTAGTGTAGC GGTGAAATGC GTAGATATTA GGAGGAACAC
NTI82-a                       GAACTGCGTG GCTGGAGTGT CGGAGAGGCA GGCGGAATTC CTAGTGTAGC GGTGAAATGC GTAGATATTA GGAGGAACAC
NTI82-b                       GAACTGCGTG GCTGGAGTGT CGGAGAGGCA GGCGGAATTC CTAGTGTAGC GGTGAAATGC GTAGATATTA GGAGGAACAC
NTI82-c                       GAACTGCGTG GCTGGAGTGT CGGAGAGGCA GGCGGAATTC CTAGTGTAGC GGTGAAATGC GTAGATATTA GGAGGAACAC
NTI82-d                       GAACTGCGTG GCTGGAGTGT CGGAGAGGCA GGCGGAATTC CTAGTGTAGC GGTGAAATGC GTAGATATTA GGAGGAACAC
Q4-a                          GAACTGCGTG GCTGGAGTGT CGGAGAGGCA GGCGGAATTC CTAGTGTAGC GGTGAAATGC GTAGATATTA GGAGGAACAC
Q4-b                          GAACTGCGTG GCTGGAGTGT CGGAGAGGCA GGCGGAATTC CTAGTGTAGC GGTGAAATGC GTAGATATTA GGAGGAACAC
Q4-c                          GAACTGCGTG GCTGGAGTGT CGGAGAGGCA GGCGGAATTC CTAGTGTAGC GGTGAAATGC GTAGATATTA GGAGGAACAC
Q4-d                          GAACTGCGTG GCTGGAGTGT CGGAGAGGCA GGCGGAATTC CTAGTGTAGC GGTGAAATGC GTAGATATTA GGAGGAACAC
S076-a                        GAACTGCGTG GCTGGAGTGT CGGAGAGGCA GGCGGAATTC CTAGTGTAGC GGTGAAATGC GTAGATATTA GGAGGAACAC
S076-b                        GAACTGCGTG GCTGGAGTGT CGGAGAGGCA GGCGGAATTC CTAGTGTAGC GGTGAAATGC GTAGATATTA GGAGGAACAC
S076-c                        GAACTGCGTG GCTGGAGTGT CGGAGAGGCA GGCGGAATTC CTAGTGTAGC GGTGAAATGC GTAGATATTA GGAGGAACAC
S076-d                        GAACTGCGTG GCTGGAGTGT CGGAGAGGCA GGCGGAATTC CTAGTGTAGC GGTGAAATGC GTAGATATTA GGAGGAACAC
SL.1.22                       GAACTGCGTG GCTGGAGTGT CGGAGAGGCA GGCGGAATTC CTAGTGTAGC GGTGAAATGC GTAGATATTA GGAGGAACAC
VE202-05                      GAACTGCGTG GCTGGAGTGT CGGAGAGGCA GGCGGAATTC CTAGTGTAGC GGTGAAATGC GTAGATATTA GGAGGAACAC
C.hylemonae                   AAACTGTCAG GCTAGAGTGT CGGAGAGGCA AGTGGAATTC CTAGTGTAGC GGTGAAATGC GTAGATATTA GGAGGAACAC
```

```
                      721
35704-a               CAGTGGCGAA GGCGGCCTGC TGGACGATGA CTGACGTTGA GGCTCGAAAG CGTGGGGAGC AAACAGGATT AGATACCCTG
35704-b               CAGTGGCGAA GGCGGCCTGC TGGACGATGA CTGACGTTGA GGCTCGAAAG CGTGGGGAGC AAACAGGATT AGATACCCTG
35704-c               CAGTGGCGAA GGCGGCCTGC TGGACGATGA CTGACGTTGA GGCTCGAAAG CGTGGGGAGC AAACAGGATT AGATACCCTG
35704-d               CAGTGGCGAA GGCGGCCTGC TGGACGATGA CTGACGTTGA GGCTCGAAAG CGTGGGGAGC AAACAGGATT AGATACCCTG
AM05-22               CAGTGGCGAA GGCGGCCTGC TGGACGATGA CTGACGTTGA GGCTCGAAAG CGTGGGGAGC AAACAGGATT AGATACCCTG
AM07-30               CAGTGGCGAA GGCGGCCTGC TGGACGATGA CTGACGTTGA GGCTCGAAAG CGTGGGGAGC AAACAGGATT AGATACCCTG
BL389WT3D-a           CAGTGGCGAA GGCGGCCTGC TGGACGATGA CTGACGTTGA GGCTCGAAAG CGTGGGGAGC AAACAGGATT AGATACCCTG
BL389WT3D-b           CAGTGGCGAA GGCGGCCTGC TGGACGATGA CTGACGTTGA GGCTCGAAAG CGTGGGGAGC AAACAGGATT AGATACCCTG
BL389WT3D-c           CAGTGGCGAA GGCGGCCTGC TGGACGATGA CTGACGTTGA GGCTCGAAAG CGTGGGGAGC AAACAGGATT AGATACCCTG
BL389WT3D-d           CAGTGGCGAA GGCGGCCTGC TGGACGATGA CTGACGTTGA GGCTCGAAAG CGTGGGGAGC AAACAGGATT AGATACCCTG
CE91-St59-a           CAGTGGCGAA GGCGGCCTGC TGGACGATGA CTGACGTTGA GGCTCGAAAG CGTGGGGAGC AAACAGGATT AGATACCCTG
CE91-St59-b           CAGTGGCGAA GGCGGCCTGC TGGACGATGA CTGACGTTGA GGCTCGAAAG CGTGGGGAGC AAACAGGATT AGATACCCTG
CE91-St59-c           CAGTGGCGAA GGCGGCCTGC TGGACGATGA CTGACGTTGA GGCTCGAAAG CGTGGGGAGC AAACAGGATT AGATACCCTG
CE91-St60-a           CAGTGGCGAA GGCGGCCTGC TGGACGATGA CTGACGTTGA GGCTCGAAAG CGTGGGGAGC AAACAGGATT AGATACCCTG
CE91-St60-b           CAGTGGCGAA GGCGGCCTGC TGGACGATGA CTGACGTTGA GGCTCGAAAG CGTGGGGAGC AAACAGGATT AGATACCCTG
CE91-St60-c           CAGTGGCGAA GGCGGCCTGC TGGACGATGA CTGACGTTGA GGCTCGAAAG CGTGGGGAGC AAACAGGATT AGATACCCTG
FDAARGOS_1227-a       CAGTGGCGAA GGCGGCCTGC TGGACGATGA CTGACGTTGA GGCTCGAAAG CGTGGGGAGC AAACAGGATT AGATACCCTG
FDAARGOS_1227-b       CAGTGGCGAA GGCGGCCTGC TGGACGATGA CTGACGTTGA GGCTCGAAAG CGTGGGGAGC AAACAGGATT AGATACCCTG
FDAARGOS_1227-c       CAGTGGCGAA GGCGGCCTGC TGGACGATGA CTGACGTTGA GGCTCGAAAG CGTGGGGAGC AAACAGGATT AGATACCCTG
FDAARGOS_1227-d       CAGTGGCGAA GGCGGCCTGC TGGACGATGA CTGACGTTGA GGCTCGAAAG CGTGGGGAGC AAACAGGATT AGATACCCTG
G10-a                 CAGTGGCGAA GGCGGCCTGC TGGACGATGA CTGACGTTGA GGCTCGAAAG CGTGGGGAGC AAACAGGATT AGATACCCTG
G10-b                 CAGTGGCGAA GGCGGCCTGC TGGACGATGA CTGACGTTGA GGCTCGAAAG CGTGGGGAGC AAACAGGATT AGATACCCTG
G10-c                 CAGTGGCGAA GGCGGCCTGC TGGACGATGA CTGACGTTGA GGCTCGAAAG CGTGGGGAGC AAACAGGATT AGATACCCTG
G10-d                 CAGTGGCGAA GGCGGCCTGC TGGACGATGA CTGACGTTGA GGCTCGAAAG CGTGGGGAGC AAACAGGATT AGATACCCTG
GGCC_0168             CAGTGGCGAA GGCGGCCTGC TGGACGATGA CTGACGTTGA GGCTCGAAAG CGTGGGGAGC AAACAGGATT AGATACCCTG
I10-a                 CAGTGGCGAA GGCGGCCTGC TGGACGATGA CTGACGTTGA GGCTCGAAAG CGTGGGGAGC AAACAGGATT AGATACCCTG
I10-b                 CAGTGGCGAA GGCGGCCTGC TGGACGATGA CTGACGTTGA GGCTCGAAAG CGTGGGGAGC AAACAGGATT AGATACCCTG
I10-c                 CAGTGGCGAA GGCGGCCTGC TGGACGATGA CTGACGTTGA GGCTCGAAAG CGTGGGGAGC AAACAGGATT AGATACCCTG
I10-d                 CAGTGGCGAA GGCGGCCTGC TGGACGATGA CTGACGTTGA GGCTCGAAAG CGTGGGGAGC AAACAGGATT AGATACCCTG
JCM10422-a            CAGTGGCGAA GGCGGCCTGC TGGACGATGA CTGACGTTGA GGCTCGAAAG CGTGGGGAGC AAACAGGATT AGATACCCTG
JCM10422-b            CAGTGGCGAA GGCGGCCTGC TGGACGATGA CTGACGTTGA GGCTCGAAAG CGTGGGGAGC AAACAGGATT AGATACCCTG
JCM10422-c            CAGTGGCGAA GGCGGCCTGC TGGACGATGA CTGACGTTGA GGCTCGAAAG CGTGGGGAGC AAACAGGATT AGATACCCTG
JCM10422-d            CAGTGGCGAA GGCGGCCTGC TGGACGATGA CTGACGTTGA GGCTCGAAAG CGTGGGGAGC AAACAGGATT AGATACCCTG
MGYG-HGUT-01303-a     CAGTGGCGAA GGCGGCCTGC TGGACGATGA CTGACGTTGA GGCTCGAAAG CGTGGGGAGC AAACAGGATT AGATACCCTG
MGYG-HGUT-01303-b     CAGTGGCGAA GGCGGCCTGC TGGACGATGA CTGACGTTGA GGCTCGAAAG CGTGGGGAGC AAACAGGATT AGATACCCTG
MSK.1.16              CAGTGGCGAA GGCGGCCTGC TGGACGATGA CTGACGTTGA GGCTCGAAAG CGTGGGGAGC AAACAGGATT AGATACCCTG
MSK.1.26              CAGTGGCGAA GGCGGCCTGC TGGACGATGA CTGACGTTGA GGCTCGAAAG CGTGGGGAGC AAACAGGATT AGATACCCTG
S077-U-a              CAGTGGCGAA GGCGGCCTGC TGGACGATGA CTGACGTTGA GGCTCGAAAG CGTGGGGAGC AAACAGGATT AGATACCCTG
S077-U-b              CAGTGGCGAA GGCGGCCTGC TGGACGATGA CTGACGTTGA GGCTCGAAAG CGTGGGGAGC AAACAGGATT AGATACCCTG
S077-U-c              CAGTGGCGAA GGCGGCCTGC TGGACGATGA CTGACGTTGA GGCTCGAAAG CGTGGGGAGC AAACAGGATT AGATACCCTG
S077-U-d              CAGTGGCGAA GGCGGCCTGC TGGACGATGA CTGACGTTGA GGCTCGAAAG CGTGGGGAGC AAACAGGATT AGATACCCTG
VPI12708-a            CAGTGGCGAA GGCGGCCTGC TGGACGATGA CTGACGTTGA GGCTCGAAAG CGTGGGGAGC AAACAGGATT AGATACCCTG
VPI12708-b            CAGTGGCGAA GGCGGCCTGC TGGACGATGA CTGACGTTGA GGCTCGAAAG CGTGGGGAGC AAACAGGATT AGATACCCTG
VPI12708-c            CAGTGGCGAA GGCGGCCTGC TGGACGATGA CTGACGTTGA GGCTCGAAAG CGTGGGGAGC AAACAGGATT AGATACCCTG
VPI12708-d            CAGTGGCGAA GGCGGCCTGC TGGACGATGA CTGACGTTGA GGCTCGAAAG CGTGGGGAGC AAACAGGATT AGATACCCTG
DFI.1.130             CAGTGGCGAA GGCGGCCTGC TGGACGATGA CTGACGTTGA GGCTCGAAAG CGTGGGGAGC AAACAGGATT AGATACCCTG
DFI.1.161             CAGTGGCGAA GGCGGCCTGC TGGACGATGA CTGACGTTGA GGCTCGAAAG CGTGGGGAGC AAACAGGATT AGATACCCTG
DFI.1.162             CAGTGGCGAA GGCGGCCTGC TGGACGATGA CTGACGTTGA GGCTCGAAAG CGTGGGGAGC AAACAGGATT AGATACCCTG
DFI.1.217             CAGTGGCGAA GGCGGCCTGC TGGACGATGA CTGACGTTGA GGCTCGAAAG CGTGGGGAGC AAACAGGATT AGATACCCTG
DFI.1.234             CAGTGGCGAA GGCGGCCTGC TGGACGATGA CTGACGTTGA GGCTCGAAAG CGTGGGGAGC AAACAGGATT AGATACCCTG
DFI.1.60              CAGTGGCGAA GGCGGCCTGC TGGACGATGA CTGACGTTGA GGCTCGAAAG CGTGGGGAGC AAACAGGATT AGATACCCTG
DFI.4.63              CAGTGGCGAA GGCGGCCTGC TGGACGATGA CTGACGTTGA GGCTCGAAAG CGTGGGGAGC AAACAGGATT AGATACCCTG
JCM10419-a            CAGTGGCGAA GGCGGCCTGC TGGACGATGA CTGACGTTGA GGCTCGAAAG CGTGGGGAGC AAACAGGATT AGATACCCTG
JCM10419-b            CAGTGGCGAA GGCGGCCTGC TGGACGATGA CTGACGTTGA GGCTCGAAAG CGTGGGGAGC AAACAGGATT AGATACCCTG
JCM10419-c            CAGTGGCGAA GGCGGCCTGC TGGACGATGA CTGACGTTGA GGCTCGAAAG CGTGGGGAGC AAACAGGATT AGATACCCTG
JCM10419-d            CAGTGGCGAA GGCGGCCTGC TGGACGATGA CTGACGTTGA GGCTCGAAAG CGTGGGGAGC AAACAGGATT AGATACCCTG
JCM10420-a            CAGTGGCGAA GGCGGCCTGC TGGACGATGA CTGACGTTGA GGCTCGAAAG CGTGGGGAGC AAACAGGATT AGATACCCTG
JCM10420-b            CAGTGGCGAA GGCGGCCTGC TGGACGATGA CTGACGTTGA GGCTCGAAAG CGTGGGGAGC AAACAGGATT AGATACCCTG
JCM10420-c            CAGTGGCGAA GGCGGCCTGC TGGACGATGA CTGACGTTGA GGCTCGAAAG CGTGGGGAGC AAACAGGATT AGATACCCTG
JCM10420-d            CAGTGGCGAA GGCGGCCTGC TGGACGATGA CTGACGTTGA GGCTCGAAAG CGTGGGGAGC AAACAGGATT AGATACCCTG
JCM10423-a            CAGTGGCGAA GGCGGCCTGC TGGACGATGA CTGACGTTGA GGCTCGAAAG CGTGGGGAGC AAACAGGATT AGATACCCTG
JCM10423-b            CAGTGGCGAA GGCGGCCTGC TGGACGATGA CTGACGTTGA GGCTCGAAAG CGTGGGGAGC AAACAGGATT AGATACCCTG
JCM10423-c            CAGTGGCGAA GGCGGCCTGC TGGACGATGA CTGACGTTGA GGCTCGAAAG CGTGGGGAGC AAACAGGATT AGATACCCTG
JCM10423-d            CAGTGGCGAA GGCGGCCTGC TGGACGATGA CTGACGTTGA GGCTCGAAAG CGTGGGGAGC AAACAGGATT AGATACCCTG
MO32-a                CAGTGGCGAA GGCGGCCTGC TGGACGATGA CTGACGTTGA GGCTCGAAAG CGTGGGGAGC AAACAGGATT AGATACCCTG
MO32-b                CAGTGGCGAA GGCGGCCTGC TGGACGATGA CTGACGTTGA GGCTCGAAAG CGTGGGGAGC AAACAGGATT AGATACCCTG
MO32-c                CAGTGGCGAA GGCGGCCTGC TGGACGATGA CTGACGTTGA GGCTCGAAAG CGTGGGGAGC AAACAGGATT AGATACCCTG
MO32-d                CAGTGGCGAA GGCGGCCTGC TGGACGATGA CTGACGTTGA GGCTCGAAAG CGTGGGGAGC AAACAGGATT AGATACCCTG
MSK.5.24              CAGTGGCGAA GGCGGCCTGC TGGACGATGA CTGACGTTGA GGCTCGAAAG CGTGGGGAGC AAACAGGATT AGATACCCTG
NB2A-7-D5             CAGTGGCGAA GGCGGCCTGC TGGACGATGA CTGACGTTGA GGCTCGAAAG CGTGGGGAGC AAACAGGATT AGATACCCTG
NTI82-a               CAGTGGCGAA GGCGGCCTGC TGGACGATGA CTGACGTTGA GGCTCGAAAG CGTGGGGAGC AAACAGGATT AGATACCCTG
NTI82-b               CAGTGGCGAA GGCGGCCTGC TGGACGATGA CTGACGTTGA GGCTCGAAAG CGTGGGGAGC AAACAGGATT AGATACCCTG
NTI82-c               CAGTGGCGAA GGCGGCCTGC TGGACGATGA CTGACGTTGA GGCTCGAAAG CGTGGGGAGC AAACAGGATT AGATACCCTG
NTI82-d               CAGTGGCGAA GGCGGCCTGC TGGACGATGA CTGACGTTGA GGCTCGAAAG CGTGGGGAGC AAACAGGATT AGATACCCTG
Q4-a                  CAGTGGCGAA GGCGGCCTGC TGGACGATGA CTGACGTTGA GGCTCGAAAG CGTGGGGAGC AAACAGGATT AGATACCCTG
Q4-b                  CAGTGGCGAA GGCGGCCTGC TGGACGATGA CTGACGTTGA GGCTCGAAAG CGTGGGGAGC AAACAGGATT AGATACCCTG
Q4-c                  CAGTGGCGAA GGCGGCCTGC TGGACGATGA CTGACGTTGA GGCTCGAAAG CGTGGGGAGC AAACAGGATT AGATACCCTG
Q4-d                  CAGTGGCGAA GGCGGCCTGC TGGACGATGA CTGACGTTGA GGCTCGAAAG CGTGGGGAGC AAACAGGATT AGATACCCTG
S076-a                CAGTGGCGAA GGCGGCCTGC TGGACGATGA CTGACGTTGA GGCTCGAAAG CGTGGGGAGC AAACAGGATT AGATACCCTG
S076-b                CAGTGGCGAA GGCGGCCTGC TGGACGATGA CTGACGTTGA GGCTCGAAAG CGTGGGGAGC AAACAGGATT AGATACCCTG
S076-c                CAGTGGCGAA GGCGGCCTGC TGGACGATGA CTGACGTTGA GGCTCGAAAG CGTGGGGAGC AAACAGGATT AGATACCCTG
S076-d                CAGTGGCGAA GGCGGCCTGC TGGACGATGA CTGACGTTGA GGCTCGAAAG CGTGGGGAGC AAACAGGATT AGATACCCTG
SL.1.22               CAGTGGCGAA GGCGGCCTGC TGGACGATGA CTGACGTTGA GGCTCGAAAG CGTGGGGAGC AAACAGGATT AGATACCCTG
VE202-05              CAGTGGCGAA GGCGGCCTGC TGGACGATGA CTGACGTTGA GGCTCGAAAG CGTGGGGAGC AAACAGGATT AGATACCCTG
C.hylemonae           CAGTGGCGAA GGCGGCTTGC TGGACGATGA CTGACGTTGA GGCTCGAAAG CGTGGGGAGC AAACAGGATT AGATACCCTG
```

```
                    801
35704-a             GTAGTCCACG CCGTAAACGA TGACTACTAG GTGTCGGGTG GCAAGGCCAT TCGGTGCCGC AGCAAACGCA ATAAGTAGTC
35704-b             GTAGTCCACG CCGTAAACGA TGACTACTAG GTGTCGGGTG GCAAGGCCAT TCGGTGCCGC AGCAAACGCA ATAAGTAGTC
35704-c             GTAGTCCACG CCGTAAACGA TGACTACTAG GTGTCGGGTG GCAAGGCCAT TCGGTGCCGC AGCAAACGCA ATAAGTAGTC
35704-d             GTAGTCCACG CCGTAAACGA TGACTACTAG GTGTCGGGTG GCAAGGCCAT TCGGTGCCGC AGCAAACGCA ATAAGTAGTC
AM05-22             GTAGTCCACG CCGTAAACGA TGACTACTAG GTGTCGGGTG GCAAGGCCAT TCGGTGCCGC AGCAAACGCA ATAAGTAGTC
AM07-30             GTAGTCCACG CCGTAAACGA TGACTACTAG GTGTCGGGTG GCAAGGCCAT TCGGTGCCGC AGCAAACGCA ATAAGTAGTC
BL389WT3D-a         GTAGTCCACG CCGTAAACGA TGACTACTAG GTGTCGGGTG GCAAGGCCAT TCGGTGCCGC AGCAAACGCA ATAAGTAGTC
BL389WT3D-b         GTAGTCCACG CCGTAAACGA TGACTACTAG GTGTCGGGTG GCAAGGCCAT TCGGTGCCGC AGCAAACGCA ATAAGTAGTC
BL389WT3D-c         GTAGTCCACG CCGTAAACGA TGACTACTAG GTGTCGGGTG GCAAGGCCAT TCGGTGCCGC AGCAAACGCA ATAAGTAGTC
BL389WT3D-d         GTAGTCCACG CCGTAAACGA TGACTACTAG GTGTCGGGTG GCAAGGCCAT TCGGTGCCGC AGCAAACGCA ATAAGTAGTC
CE91-St59-a         GTAGTCCACG CCGTAAACGA TGACTACTAG GTGTCGGGTG GCAAGGCCAT TCGGTGCCGC AGCAAACGCA ATAAGTAGTC
CE91-St59-b         GTAGTCCACG CCGTAAACGA TGACTACTAG GTGTCGGGTG GCAAGGCCAT TCGGTGCCGC AGCAAACGCA ATAAGTAGTC
CE91-St59-c         GTAGTCCACG CCGTAAACGA TGACTACTAG GTGTCGGGTG GCAAGGCCAT TCGGTGCCGC AGCAAACGCA ATAAGTAGTC
CE91-St60-a         GTAGTCCACG CCGTAAACGA TGACTACTAG GTGTCGGGTG GCAAGGCCAT TCGGTGCCGC AGCAAACGCA ATAAGTAGTC
CE91-St60-b         GTAGTCCACG CCGTAAACGA TGACTACTAG GTGTCGGGTG GCAAGGCCAT TCGGTGCCGC AGCAAACGCA ATAAGTAGTC
CE91-St60-c         GTAGTCCACG CCGTAAACGA TGACTACTAG GTGTCGGGTG GCAAGGCCAT TCGGTGCCGC AGCAAACGCA ATAAGTAGTC
FDAARGOS_1227-a     GTAGTCCACG CCGTAAACGA TGACTACTAG GTGTCGGGTG GCAAGGCCAT TCGGTGCCGC AGCAAACGCA ATAAGTAGTC
FDAARGOS_1227-b     GTAGTCCACG CCGTAAACGA TGACTACTAG GTGTCGGGTG GCAAGGCCAT TCGGTGCCGC AGCAAACGCA ATAAGTAGTC
FDAARGOS_1227-c     GTAGTCCACG CCGTAAACGA TGACTACTAG GTGTCGGGTG GCAAGGCCAT TCGGTGCCGC AGCAAACGCA ATAAGTAGTC
FDAARGOS_1227-d     GTAGTCCACG CCGTAAACGA TGACTACTAG GTGTCGGGTG GCAAGGCCAT TCGGTGCCGC AGCAAACGCA ATAAGTAGTC
G10-a               GTAGTCCACG CCGTAAACGA TGACTACTAG GTGTCGGGTG GCAAGGCCAT TCGGTGCCGC AGCAAACGCA ATAAGTAGTC
G10-b               GTAGTCCACG CCGTAAACGA TGACTACTAG GTGTCGGGTG GCAAGGCCAT TCGGTGCCGC AGCAAACGCA ATAAGTAGTC
G10-c               GTAGTCCACG CCGTAAACGA TGACTACTAG GTGTCGGGTG GCAAGGCCAT TCGGTGCCGC AGCAAACGCA ATAAGTAGTC
G10-d               GTAGTCCACG CCGTAAACGA TGACTACTAG GTGTCGGGTG GCAAGGCCAT TCGGTGCCGC AGCAAACGCA ATAAGTAGTC
GGCC_0168           GTAGTCCACG CCGTAAACGA TGACTACTAG GTGTCGGGTG GCAAGGCCAT TCGGTGCCGC AGCAAACGCA ATAAGTAGTC
I10-a               GTAGTCCACG CCGTAAACGA TGACTACTAG GTGTCGGGTG GCAAGGCCAT TCGGTGCCGC AGCAAACGCA ATAAGTAGTC
I10-b               GTAGTCCACG CCGTAAACGA TGACTACTAG GTGTCGGGTG GCAAGGCCAT TCGGTGCCGC AGCAAACGCA ATAAGTAGTC
I10-c               GTAGTCCACG CCGTAAACGA TGACTACTAG GTGTCGGGTG GCAAGGCCAT TCGGTGCCGC AGCAAACGCA ATAAGTAGTC
I10-d               GTAGTCCACG CCGTAAACGA TGACTACTAG GTGTCGGGTG GCAAGGCCAT TCGGTGCCGC AGCAAACGCA ATAAGTAGTC
JCM10422-a          GTAGTCCACG CCGTAAACGA TGACTACTAG GTGTCGGGTG GCAAGGCCAT TCGGTGCCGC AGCAAACGCA ATAAGTAGTC
JCM10422-b          GTAGTCCACG CCGTAAACGA TGACTACTAG GTGTCGGGTG GCAAGGCCAT TCGGTGCCGC AGCAAACGCA ATAAGTAGTC
JCM10422-c          GTAGTCCACG CCGTAAACGA TGACTACTAG GTGTCGGGTG GCAAGGCCAT TCGGTGCCGC AGCAAACGCA ATAAGTAGTC
JCM10422-d          GTAGTCCACG CCGTAAACGA TGACTACTAG GTGTCGGGTG GCAAGGCCAT TCGGTGCCGC AGCAAACGCA ATAAGTAGTC
MGYG-HGUT-01303-a   GTAGTCCACG CCGTAAACGA TGACTACTAG GTGTCGGGTG GCAAGGCCAT TCGGTGCCGC AGCAAACGCA ATAAGTAGTC
MGYG-HGUT-01303-b   GTAGTCCACG CCGTAAACGA TGACTACTAG GTGTCGGGTG GCAAGGCCAT TCGGTGCCGC AGCAAACGCA ATAAGTAGTC
MSK.1.16            GTAGTCCACG CCGTAAACGA TGACTACTAG GTGTCGGGTG GCAAGGCCAT TCGGTGCCGC AGCAAACGCA ATAAGTAGTC
MSK.1.26            GTAGTCCACG CCGTAAACGA TGACTACTAG GTGTCGGGTG GCAAGGCCAT TCGGTGCCGC AGCAAACGCA ATAAGTAGTC
S077-U-a            GTAGTCCACG CCGTAAACGA TGACTACTAG GTGTCGGGTG GCAAGGCCAT TCGGTGCCGC AGCAAACGCA ATAAGTAGTC
S077-U-b            GTAGTCCACG CCGTAAACGA TGACTACTAG GTGTCGGGTG GCAAGGCCAT TCGGTGCCGC AGCAAACGCA ATAAGTAGTC
S077-U-c            GTAGTCCACG CCGTAAACGA TGACTACTAG GTGTCGGGTG GCAAGGCCAT TCGGTGCCGC AGCAAACGCA ATAAGTAGTC
S077-U-d            GTAGTCCACG CCGTAAACGA TGACTACTAG GTGTCGGGTG GCAAGGCCAT TCGGTGCCGC AGCAAACGCA ATAAGTAGTC
VPI12708-a          GTAGTCCACG CCGTAAACGA TGACTACTAG GTGTCGGGTG GCAAGGCCAT TCGGTGCCGC AGCAAACGCA ATAAGTAGTC
VPI12708-b          GTAGTCCACG CCGTAAACGA TGACTACTAG GTGTCGGGTG GCAAGGCCAT TCGGTGCCGC AGCAAACGCA ATAAGTAGTC
VPI12708-c          GTAGTCCACG CCGTAAACGA TGACTACTAG GTGTCGGGTG GCAAGGCCAT TCGGTGCCGC AGCAAACGCA ATAAGTAGTC
VPI12708-d          GTAGTCCACG CCGTAAACGA TGACTACTAG GTGTCGGGTG GCAAGGCCAT TCGGTGCCGC AGCAAACGCA ATAAGTAGTC
DFI.1.130           GTAGTCCACG CCGTAAACGA TGACTACTAG GTGTCGGGTG GCAAGGCCAT TCGGTGCCGC AGCAAACGCA ATAAGTAGTC
DFI.1.161           GTAGTCCACG CCGTAAACGA TGACTACTAG GTGTCGGGTG GCAAGGCCAT TCGGTGCCGC AGCAAACGCA ATAAGTAGTC
DFI.1.162           GTAGTCCACG CCGTAAACGA TGACTACTAG GTGTCGGGTG GCAAGGCCAT TCGGTGCCGC AGCAAACGCA ATAAGTAGTC
DFI.1.217           GTAGTCCACG CCGTAAACGA TGACTACTAG GTGTCGGGTG GCAAGGCCAT TCGGTGCCGC AGCAAACGCA ATAAGTAGTC
DFI.1.234           GTAGTCCACG CCGTAAACGA TGACTACTAG GTGTCGGGTG GCAAGGCCAT TCGGTGCCGC AGCAAACGCA ATAAGTAGTC
DFI.1.60            GTAGTCCACG CCGTAAACGA TGACTACTAG GTGTCGGGTG GCAAGGCCAT TCGGTGCCGC AGCAAACGCA ATAAGTAGTC
DFI.4.63            GTAGTCCACG CCGTAAACGA TGACTACTAG GTGTCGGGTG GCAAGGCCAT TCGGTGCCGC AGCAAACGCA ATAAGTAGTC
JCM10419-a          GTAGTCCACG CCGTAAACGA TGACTACTAG GTGTCGGGTG GCAAGGCCAT TCGGTGCCGC AGCAAACGCA ATAAGTAGTC
JCM10419-b          GTAGTCCACG CCGTAAACGA TGACTACTAG GTGTCGGGTG GCAAGGCCAT TCGGTGCCGC AGCAAACGCA ATAAGTAGTC
JCM10419-c          GTAGTCCACG CCGTAAACGA TGACTACTAG GTGTCGGGTG GCAAGGCCAT TCGGTGCCGC AGCAAACGCA ATAAGTAGTC
JCM10419-d          GTAGTCCACG CCGTAAACGA TGACTACTAG GTGTCGGGTG GCAAGGCCAT TCGGTGCCGC AGCAAACGCA ATAAGTAGTC
JCM10420-a          GTAGTCCACG CCGTAAACGA TGACTACTAG GTGTCGGGTG GCAAGGCCAT TCGGTGCCGC AGCAAACGCA ATAAGTAGTC
JCM10420-b          GTAGTCCACG CCGTAAACGA TGACTACTAG GTGTCGGGTG GCAAGGCCAT TCGGTGCCGC AGCAAACGCA ATAAGTAGTC
JCM10420-c          GTAGTCCACG CCGTAAACGA TGACTACTAG GTGTCGGGTG GCAAGGCCAT TCGGTGCCGC AGCAAACGCA ATAAGTAGTC
JCM10420-d          GTAGTCCACG CCGTAAACGA TGACTACTAG GTGTCGGGTG GCAAGGCCAT TCGGTGCCGC AGCAAACGCA ATAAGTAGTC
JCM10423-a          GTAGTCCACG CCGTAAACGA TGACTACTAG GTGTCGGGTG GCAAGGCCAT TCGGTGCCGC AGCAAACGCA ATAAGTAGTC
JCM10423-b          GTAGTCCACG CCGTAAACGA TGACTACTAG GTGTCGGGTG GCAAGGCCAT TCGGTGCCGC AGCAAACGCA ATAAGTAGTC
JCM10423-c          GTAGTCCACG CCGTAAACGA TGACTACTAG GTGTCGGGTG GCAAGGCCAT TCGGTGCCGC AGCAAACGCA ATAAGTAGTC
JCM10423-d          GTAGTCCACG CCGTAAACGA TGACTACTAG GTGTCGGGTG GCAAGGCCAT TCGGTGCCGC AGCAAACGCA ATAAGTAGTC
MO32-a              GTAGTCCACG CCGTAAACGA TGACTACTAG GTGTCGGGTG GCAAGGCCAT TCGGTGCCGC AGCAAACGCA ATAAGTAGTC
MO32-b              GTAGTCCACG CCGTAAACGA TGACTACTAG GTGTCGGGTG GCAAGGCCAT TCGGTGCCGC AGCAAACGCA ATAAGTAGTC
MO32-c              GTAGTCCACG CCGTAAACGA TGACTACTAG GTGTCGGGTG GCAAGGCCAT TCGGTGCCGC AGCAAACGCA ATAAGTAGTC
MO32-d              GTAGTCCACG CCGTAAACGA TGACTACTAG GTGTCGGGTG GCAAGGCCAT TCGGTGCCGC AGCAAACGCA ATAAGTAGTC
MSK.5.24            GTAGTCCACG CCGTAAACGA TGACTACTAG GTGTCGGGTG GCAAGGCCAT TCGGTGCCGC AGCAAACGCA ATAAGTAGTC
NB2A-7-D5           GTAGTCCACG CCGTAAACGA TGACTACTAG GTGTCGGGTG GCAAGGCCAT TCGGTGCCGC AGCAAACGCA ATAAGTAGTC
NTI82-a             GTAGTCCACG CCGTAAACGA TGACTACTAG GTGTCGGGTG GCAAGGCCAT TCGGTGCCGC AGCAAACGCA ATAAGTAGTC
NTI82-b             GTAGTCCACG CCGTAAACGA TGACTACTAG GTGTCGGGTG GCAAGGCCAT TCGGTGCCGC AGCAAACGCA ATAAGTAGTC
NTI82-c             GTAGTCCACG CCGTAAACGA TGACTACTAG GTGTCGGGTG GCAAGGCCAT TCGGTGCCGC AGCAAACGCA ATAAGTAGTC
NTI82-d             GTAGTCCACG CCGTAAACGA TGACTACTAG GTGTCGGGTG GCAAGGCCAT TCGGTGCCGC AGCAAACGCA ATAAGTAGTC
Q4-a                GTAGTCCACG CCGTAAACGA TGACTACTAG GTGTCGGGTG GCAAGGCCAT TCGGTGCCGC AGCAAACGCA ATAAGTAGTC
Q4-b                GTAGTCCACG CCGTAAACGA TGACTACTAG GTGTCGGGTG GCAAGGCCAT TCGGTGCCGC AGCAAACGCA ATAAGTAGTC
Q4-c                GTAGTCCACG CCGTAAACGA TGACTACTAG GTGTCGGGTG GCAAGGCCAT TCGGTGCCGC AGCAAACGCA ATAAGTAGTC
Q4-d                GTAGTCCACG CCGTAAACGA TGACTACTAG GTGTCGGGTG GCAAGGCCAT TCGGTGCCGC AGCAAACGCA ATAAGTAGTC
S076-a              GTAGTCCACG CCGTAAACGA TGACTACTAG GTGTCGGGTG GCAAGGCCAT TCGGTGCCGC AGCAAACGCA ATAAGTAGTC
S076-b              GTAGTCCACG CCGTAAACGA TGACTACTAG GTGTCGGGTG GCAAGGCCAT TCGGTGCCGC AGCAAACGCA ATAAGTAGTC
S076-c              GTAGTCCACG CCGTAAACGA TGACTACTAG GTGTCGGGTG GCAAGGCCAT TCGGTGCCGC AGCAAACGCA ATAAGTAGTC
S076-d              GTAGTCCACG CCGTAAACGA TGACTACTAG GTGTCGGGTG GCAAGGCCAT TCGGTGCCGC AGCAAACGCA ATAAGTAGTC
SL.1.22             GTAGTCCACG CCGTAAACGA TGACTACTAG GTGTCGGGTG GCAAGGCCAT TCGGTGCCGC AGCAAACGCA ATAAGTAGTC
VE202-05            GTAGTCCACG CCGTAAACGA TGACTACTAG GTGTCGGGTG GCAAGGCCAT TCGGTGCCGC AGCAAACGCA ATAAGTAGTC
C.hylemonae         GTAGTCCACG CCGTAAACGA TGATTACTAG GTGTCGGGAA GCAAAGCTTT TCGGTGCCGC AGCCAACGCA ATAAGTAATC
```

```
                     881
35704-a              CACCTGGGGA GTACGTTCGC AAGAATGAAA CTCAAAGGAA TTGACGGGGA CCCGCACAAG CGGTGGAGCA TGTGGTTTAA
35704-b              CACCTGGGGA GTACGTTCGC AAGAATGAAA CTCAAAGGAA TTGACGGGGA CCCGCACAAG CGGTGGAGCA TGTGGTTTAA
35704-c              CACCTGGGGA GTACGTTCGC AAGAATGAAA CTCAAAGGAA TTGACGGGGA CCCGCACAAG CGGTGGAGCA TGTGGTTTAA
35704-d              CACCTGGGGA GTACGTTCGC AAGAATGAAA CTCAAAGGAA TTGACGGGGA CCCGCACAAG CGGTGGAGCA TGTGGTTTAA
AM05-22              CACCTGGGGA GTACGTTCGC AAGAATGAAA CTCAAAGGAA TTGACGGGGA CCCGCACAAG CGGTGGAGCA TGTGGTTTAA
AM07-30              CACCTGGGGA GTACGTTCGC AAGAATGAAA CTCAAAGGAA TTGACGGGGA CCCGCACAAG CGGTGGAGCA TGTGGTTTAA
BL389WT3D-a          CACCTGGGGA GTACGTTCGC AAGAATGAAA CTCAAAGGAA TTGACGGGGA CCCGCACAAG CGGTGGAGCA TGTGGTTTAA
BL389WT3D-b          CACCTGGGGA GTACGTTCGC AAGAATGAAA CTCAAAGGAA TTGACGGGGA CCCGCACAAG CGGTGGAGCA TGTGGTTTAA
BL389WT3D-c          CACCTGGGGA GTACGTTCGC AAGAATGAAA CTCAAAGGAA TTGACGGGGA CCCGCACAAG CGGTGGAGCA TGTGGTTTAA
BL389WT3D-d          CACCTGGGGA GTACGTTCGC AAGAATGAAA CTCAAAGGAA TTGACGGGGA CCCGCACAAG CGGTGGAGCA TGTGGTTTAA
CE91-St59-a          CACCTGGGGA GTACGTTCGC AAGAATGAAA CTCAAAGGAA TTGACGGGGA CCCGCACAAG CGGTGGAGCA TGTGGTTTAA
CE91-St59-b          CACCTGGGGA GTACGTTCGC AAGAATGAAA CTCAAAGGAA TTGACGGGGA CCCGCACAAG CGGTGGAGCA TGTGGTTTAA
CE91-St59-c          CACCTGGGGA GTACGTTCGC AAGAATGAAA CTCAAAGGAA TTGACGGGGA CCCGCACAAG CGGTGGAGCA TGTGGTTTAA
CE91-St60-a          CACCTGGGGA GTACGTTCGC AAGAATGAAA CTCAAAGGAA TTGACGGGGA CCCGCACAAG CGGTGGAGCA TGTGGTTTAA
CE91-St60-b          CACCTGGGGA GTACGTTCGC AAGAATGAAA CTCAAAGGAA TTGACGGGGA CCCGCACAAG CGGTGGAGCA TGTGGTTTAA
CE91-St60-c          CACCTGGGGA GTACGTTCGC AAGAATGAAA CTCAAAGGAA TTGACGGGGA CCCGCACAAG CGGTGGAGCA TGTGGTTTAA
FDAARGOS_1227-a      CACCTGGGGA GTACGTTCGC AAGAATGAAA CTCAAAGGAA TTGACGGGGA CCCGCACAAG CGGTGGAGCA TGTGGTTTAA
FDAARGOS_1227-b      CACCTGGGGA GTACGTTCGC AAGAATGAAA CTCAAAGGAA TTGACGGGGA CCCGCACAAG CGGTGGAGCA TGTGGTTTAA
FDAARGOS_1227-c      CACCTGGGGA GTACGTTCGC AAGAATGAAA CTCAAAGGAA TTGACGGGGA CCCGCACAAG CGGTGGAGCA TGTGGTTTAA
FDAARGOS_1227-d      CACCTGGGGA GTACGTTCGC AAGAATGAAA CTCAAAGGAA TTGACGGGGA CCCGCACAAG CGGTGGAGCA TGTGGTTTAA
G10-a                CACCTGGGGA GTACGTTCGC AAGAATGAAA CTCAAAGGAA TTGACGGGGA CCCGCACAAG CGGTGGAGCA TGTGGTTTAA
G10-b                CACCTGGGGA GTACGTTCGC AAGAATGAAA CTCAAAGGAA TTGACGGGGA CCCGCACAAG CGGTGGAGCA TGTGGTTTAA
G10-c                CACCTGGGGA GTACGTTCGC AAGAATGAAA CTCAAAGGAA TTGACGGGGA CCCGCACAAG CGGTGGAGCA TGTGGTTTAA
G10-d                CACCTGGGGA GTACGTTCGC AAGAATGAAA CTCAAAGGAA TTGACGGGGA CCCGCACAAG CGGTGGAGCA TGTGGTTTAA
GGCC_0168            CACCTGGGGA GTACGTTCGC AAGAATGAAA CTCAAAGGAA TTGACGGGGA CCCGCACAAG CGGTGGAGCA TGTGGTTTAA
I10-a                CACCTGGGGA GTACGTTCGC AAGAATGAAA CTCAAAGGAA TTGACGGGGA CCCGCACAAG CGGTGGAGCA TGTGGTTTAA
I10-b                CACCTGGGGA GTACGTTCGC AAGAATGAAA CTCAAAGGAA TTGACGGGGA CCCGCACAAG CGGTGGAGCA TGTGGTTTAA
I10-c                CACCTGGGGA GTACGTTCGC AAGAATGAAA CTCAAAGGAA TTGACGGGGA CCCGCACAAG CGGTGGAGCA TGTGGTTTAA
I10-d                CACCTGGGGA GTACGTTCGC AAGAATGAAA CTCAAAGGAA TTGACGGGGA CCCGCACAAG CGGTGGAGCA TGTGGTTTAA
JCM10422-a           CACCTGGGGA GTACGTTCGC AAGAATGAAA CTCAAAGGAA TTGACGGGGA CCCGCACAAG CGGTGGAGCA TGTGGTTTAA
JCM10422-b           CACCTGGGGA GTACGTTCGC AAGAATGAAA CTCAAAGGAA TTGACGGGGA CCCGCACAAG CGGTGGAGCA TGTGGTTTAA
JCM10422-c           CACCTGGGGA GTACGTTCGC AAGAATGAAA CTCAAAGGAA TTGACGGGGA CCCGCACAAG CGGTGGAGCA TGTGGTTTAA
JCM10422-d           CACCTGGGGA GTACGTTCGC AAGAATGAAA CTCAAAGGAA TTGACGGGGA CCCGCACAAG CGGTGGAGCA TGTGGTTTAA
MGYG-HGUT-01303-a    CACCTGGGGA GTACGTTCGC AAGAATGAAA CTCAAAGGAA TTGACGGGGA CCCGCACAAG CGGTGGAGCA TGTGGTTTAA
MGYG-HGUT-01303-b    CACCTGGGGA GTACGTTCGC AAGAATGAAA CTCAAAGGAA TTGACGGGGA CCCGCACAAG CGGTGGAGCA TGTGGTTTAA
MSK.1.16             CACCTGGGGA GTACGTTCGC AAGAATGAAA CTCAAAGGAA TTGACGGGGA CCCGCACAAG CGGTGGAGCA TGTGGTTTAA
MSK.1.26             CACCTGGGGA GTACGTTCGC AAGAATGAAA CTCAAAGGAA TTGACGGGGA CCCGCACAAG CGGTGGAGCA TGTGGTTTAA
S077-U-a             CACCTGGGGA GTACGTTCGC AAGAATGAAA CTCAAAGGAA TTGACGGGGA CCCGCACAAG CGGTGGAGCA TGTGGTTTAA
S077-U-b             CACCTGGGGA GTACGTTCGC AAGAATGAAA CTCAAAGGAA TTGACGGGGA CCCGCACAAG CGGTGGAGCA TGTGGTTTAA
S077-U-c             CACCTGGGGA GTACGTTCGC AAGAATGAAA CTCAAAGGAA TTGACGGGGA CCCGCACAAG CGGTGGAGCA TGTGGTTTAA
S077-U-d             CACCTGGGGA GTACGTTCGC AAGAATGAAA CTCAAAGGAA TTGACGGGGA CCCGCACAAG CGGTGGAGCA TGTGGTTTAA
VPI12708-a           CACCTGGGGA GTACGTTCGC AAGAATGAAA CTCAAAGGAA TTGACGGGGA CCCGCACAAG CGGTGGAGCA TGTGGTTTAA
VPI12708-b           CACCTGGGGA GTACGTTCGC AAGAATGAAA CTCAAAGGAA TTGACGGGGA CCCGCACAAG CGGTGGAGCA TGTGGTTTAA
VPI12708-c           CACCTGGGGA GTACGTTCGC AAGAATGAAA CTCAAAGGAA TTGACGGGGA CCCGCACAAG CGGTGGAGCA TGTGGTTTAA
VPI12708-d           CACCTGGGGA GTACGTTCGC AAGAATGAAA CTCAAAGGAA TTGACGGGGA CCCGCACAAG CGGTGGAGCA TGTGGTTTAA
DFI.1.130            CACCTGGGGA GTACGTTCGC AAGAATGAAA CTCAAAGGAA TTGACGGGGA CCCGCACAAG CGGTGGAGCA TGTGGTTTAA
DFI.1.161            CACCTGGGGA GTACGTTCGC AAGAATGAAA CTCAAAGGAA TTGA------ ---------- ---------- ----------
DFI.1.162            CACCTGGGGA GTACGTTCGC AAGAATGAAA CTCAAAGGAA TTGACGGGGA CCCGCACAAG CGGTGGAGCA TGTGGTTTAA
DFI.1.217            CACCTGGGGA GTACGTTCGC AAGAATGAAA CTCAAAGGAA TTGACGGGGA CCCGCACAAG CGGTGGAGCA TGTGGTTTAA
DFI.1.234            CACCTGGGGA GTACGTTCGC AAGAATGAAA CTCAAAGGAA TTGACGGGGA CCCGCACAAG CGGTGGAGCA TGTGGTTTAA
DFI.1.60             CACCTGGGGA GTACGTTCGC AAGAATGAAA CTCAAAGGAA TTGACGGGGA CCCGCACAAG CGGTGGAGCA TGTGGTTTAA
DFI.4.63             CACCTGGGGA GTACGTTCGC AAGAATGAAA CTCAAAGGAA TTGACGGGGA CCCGCACAAG CGGTGGAGCA TGTGGTTTAA
JCM10419-a           CACCTGGGGA GTACGTTCGC AAGAATGAAA CTCAAAGGAA TTGACGGGGA CCCGCACAAG CGGTGGAGCA TGTGGTTTAA
JCM10419-b           CACCTGGGGA GTACGTTCGC AAGAATGAAA CTCAAAGGAA TTGACGGGGA CCCGCACAAG CGGTGGAGCA TGTGGTTTAA
JCM10419-c           CACCTGGGGA GTACGTTCGC AAGAATGAAA CTCAAAGGAA TTGACGGGGA CCCGCACAAG CGGTGGAGCA TGTGGTTTAA
JCM10419-d           CACCTGGGGA GTACGTTCGC AAGAATGAAA CTCAAAGGAA TTGACGGGGA CCCGCACAAG CGGTGGAGCA TGTGGTTTAA
JCM10420-a           CACCTGGGGA GTACGTTCGC AAGAATGAAA CTCAAAGGAA TTGACGGGGA CCCGCACAAG CGGTGGAGCA TGTGGTTTAA
JCM10420-b           CACCTGGGGA GTACGTTCGC AAGAATGAAA CTCAAAGGAA TTGACGGGGA CCCGCACAAG CGGTGGAGCA TGTGGTTTAA
JCM10420-c           CACCTGGGGA GTACGTTCGC AAGAATGAAA CTCAAAGGAA TTGACGGGGA CCCGCACAAG CGGTGGAGCA TGTGGTTTAA
JCM10420-d           CACCTGGGGA GTACGTTCGC AAGAATGAAA CTCAAAGGAA TTGACGGGGA CCCGCACAAG CGGTGGAGCA TGTGGTTTAA
JCM10423-a           CACCTGGGGA GTACGTTCGC AAGAATGAAA CTCAAAGGAA TTGACGGGGA CCCGCACAAG CGGTGGAGCA TGTGGTTTAA
JCM10423-b           CACCTGGGGA GTACGTTCGC AAGAATGAAA CTCAAAGGAA TTGACGGGGA CCCGCACAAG CGGTGGAGCA TGTGGTTTAA
JCM10423-c           CACCTGGGGA GTACGTTCGC AAGAATGAAA CTCAAAGGAA TTGACGGGGA CCCGCACAAG CGGTGGAGCA TGTGGTTTAA
JCM10423-d           CACCTGGGGA GTACGTTCGC AAGAATGAAA CTCAAAGGAA TTGACGGGGA CCCGCACAAG CGGTGGAGCA TGTGGTTTAA
MO32-a               CACCTGGGGA GTACGTTCGC AAGAATGAAA CTCAAAGGAA TTGACGGGGA CCCGCACAAG CGGTGGAGCA TGTGGTTTAA
MO32-b               CACCTGGGGA GTACGTTCGC AAGAATGAAA CTCAAAGGAA TTGACGGGGA CCCGCACAAG CGGTGGAGCA TGTGGTTTAA
MO32-c               CACCTGGGGA GTACGTTCGC AAGAATGAAA CTCAAAGGAA TTGACGGGGA CCCGCACAAG CGGTGGAGCA TGTGGTTTAA
MO32-d               CACCTGGGGA GTACGTTCGC AAGAATGAAA CTCAAAGGAA TTGACGGGGA CCCGCACAAG CGGTGGAGCA TGTGGTTTAA
MSK.5.24             CACCTGGGGA GTACGTTCGC AAGAATGAAA CTCAAAGGAA TTGACGGGGA CCCGCACAAG CGGTGGAGCA TGTGGTTTAA
NB2A-7-D5            CACCTGGGGA GTACGTTCGC AAGAATGAAA CTCAAAGGAA TTGACGGGGA CCCGCACAAG CGGTGGAGCA TGTGGTTTAA
NTI82-a              CACCTGGGGA GTACGTTCGC AAGAATGAAA CTCAAAGGAA TTGACGGGGA CCCGCACAAG CGGTGGAGCA TGTGGTTTAA
NTI82-b              CACCTGGGGA GTACGTTCGC AAGAATGAAA CTCAAAGGAA TTGACGGGGA CCCGCACAAG CGGTGGAGCA TGTGGTTTAA
NTI82-c              CACCTGGGGA GTACGTTCGC AAGAATGAAA CTCAAAGGAA TTGACGGGGA CCCGCACAAG CGGTGGAGCA TGTGGTTTAA
NTI82-d              CACCTGGGGA GTACGTTCGC AAGAATGAAA CTCAAAGGAA TTGACGGGGA CCCGCACAAG CGGTGGAGCA TGTGGTTTAA
Q4-a                 CACCTGGGGA GTACGTTCGC AAGAATGAAA CTCAAAGGAA TTGACGGGGA CCCGCACAAG CGGTGGAGCA TGTGGTTTAA
Q4-b                 CACCTGGGGA GTACGTTCGC AAGAATGAAA CTCAAAGGAA TTGACGGGGA CCCGCACAAG CGGTGGAGCA TGTGGTTTAA
Q4-c                 CACCTGGGGA GTACGTTCGC AAGAATGAAA CTCAAAGGAA TTGACGGGGA CCCGCACAAG CGGTGGAGCA TGTGGTTTAA
Q4-d                 CACCTGGGGA GTACGTTCGC AAGAATGAAA CTCAAAGGAA TTGACGGGGA CCCGCACAAG CGGTGGAGCA TGTGGTTTAA
S076-a               CACCTGGGGA GTACGTTCGC AAGAATGAAA CTCAAAGGAA TTGACGGGGA CCCGCACAAG CGGTGGAGCA TGTGGTTTAA
S076-b               CACCTGGGGA GTACGTTCGC AAGAATGAAA CTCAAAGGAA TTGACGGGGA CCCGCACAAG CGGTGGAGCA TGTGGTTTAA
S076-c               CACCTGGGGA GTACGTTCGC AAGAATGAAA CTCAAAGGAA TTGACGGGGA CCCGCACAAG CGGTGGAGCA TGTGGTTTAA
S076-d               CACCTGGGGA GTACGTTCGC AAGAATGAAA CTCAAAGGAA TTGACGGGGA CCCGCACAAG CGGTGGAGCA TGTGGTTTAA
SL.1.22              CACCTGGGGA GTACGTTCGC AAGAATGAAA CTCAAAGGAA TTGACGGGGA CCCGCACAAG CGGTGGAGCA TGTGGTTTAA
VE202-05             CACCTGGGGA GTACGTTCGC AAGAATGAAA CTCAAAGGAA TTGACGGGGA CCCGCACAAG CGGTGGAGCA TGTGGTTTAA
C.hylemonae          CACCTGGGGA GTACGTTCGC AAGAATGAAA CTCAAAGGAA TTGACGGGGA CCCGCACAAG CGGTGGAGCA TGTGGTTTAA
```

```
                   961
35704-a            TTCGAAGCAA CGCGAAGAAC CTTACCTGAT CTTGACATCC CGATGCCAAA GCGCGTAACG CGCTCTTTCT TCGGAACATC
35704-b            TTCGAAGCAA CGCGAAGAAC CTTACCTGAT CTTGACATCC CGATGCCAAA GCGCGTAACG CGCTCTTTCT TCGGAACATC
35704-c            TTCGAAGCAA CGCGAAGAAC CTTACCTGAT CTTGACATCC CGATGCCAAA GCGCGTAACG CGCTCTTTCT TCGGAACATC
35704-d            TTCGAAGCAA CGCGAAGAAC CTTACCTGAT CTTGACATCC CGATGCCAAA GCGCGTAACG CGCTCTTTCT TCGGAACATC
AM05-22            TTCGAAGCAA CGCGAAGAAC CTTACCTGAT CTTGACATCC CGATGCCAAA GCGCGTAACG CGCTCTTTCT TCGGAACATC
AM07-30            TTCGAAGCAA CGCGAAGAAC CTTACCTGAT CTTGACATCC CGATGCCAAA GCGCGTAACG CGCTCTTTCT TCGGAACATC
BL389WT3D-a        TTCGAAGCAA CGCGAAGAAC CTTACCTGAT CTTGACATCC CGATGCCAAA GCGCGTAACG CGCTCTTTCT TCGGAACATC
BL389WT3D-b        TTCGAAGCAA CGCGAAGAAC CTTACCTGAT CTTGACATCC CGATGCCAAA GCGCGTAACG CGCTCTTTCT TCGGAACATC
BL389WT3D-c        TTCGAAGCAA CGCGAAGAAC CTTACCTGAT CTTGACATCC CGATGCCAAA GCGCGTAACG CGCTCTTTCT TCGGAACATC
BL389WT3D-d        TTCGAAGCAA CGCGAAGAAC CTTACCTGAT CTTGACATCC CGATGCCAAA GCGCGTAACG CGCTCTTTCT TCGGAACATC
CE91-St59-a        TTCGAAGCAA CGCGAAGAAC CTTACCTGAT CTTGACATCC CGATGCCAAA GCGCGTAACG CGCTCTTTCT TCGGAACATC
CE91-St59-b        TTCGAAGCAA CGCGAAGAAC CTTACCTGAT CTTGACATCC CGATGCCAAA GCGCGTAACG CGCTCTTTCT TCGGAACATC
CE91-St59-c        TTCGAAGCAA CGCGAAGAAC CTTACCTGAT CTTGACATCC CGATGCCAAA GCGCGTAACG CGCTCTTTCT TCGGAACATC
CE91-St60-a        TTCGAAGCAA CGCGAAGAAC CTTACCTGAT CTTGACATCC CGATGCCAAA GCGCGTAACG CGCTCTTTCT TCGGAACATC
CE91-St60-b        TTCGAAGCAA CGCGAAGAAC CTTACCTGAT CTTGACATCC CGATGCCAAA GCGCGTAACG CGCTCTTTCT TCGGAACATC
CE91-St60-c        TTCGAAGCAA CGCGAAGAAC CTTACCTGAT CTTGACATCC CGATGCCAAA GCGCGTAACG CGCTCTTTCT TCGGAACATC
FDAARGOS_1227-a    TTCGAAGCAA CGCGAAGAAC CTTACCTGAT CTTGACATCC CGATGCCAAA GCGCGTAACG CGCTCTTTCT TCGGAACATC
FDAARGOS_1227-b    TTCGAAGCAA CGCGAAGAAC CTTACCTGAT CTTGACATCC CGATGCCAAA GCGCGTAACG CGCTCTTTCT TCGGAACATC
FDAARGOS_1227-c    TTCGAAGCAA CGCGAAGAAC CTTACCTGAT CTTGACATCC CGATGCCAAA GCGCGTAACG CGCTCTTTCT TCGGAACATC
FDAARGOS_1227-d    TTCGAAGCAA CGCGAAGAAC CTTACCTGAT CTTGACATCC CGATGCCAAA GCGCGTAACG CGCTCTTTCT TCGGAACATC
G10-a              TTCGAAGCAA CGCGAAGAAC CTTACCTGAT CTTGACATCC CGATGCCAAA GCGCGTAACG CGCTCTTTCT TCGGAACATC
G10-b              TTCGAAGCAA CGCGAAGAAC CTTACCTGAT CTTGACATCC CGATGCCAAA GCGCGTAACG CGCTCTTTCT TCGGAACATC
G10-c              TTCGAAGCAA CGCGAAGAAC CTTACCTGAT CTTGACATCC CGATGCCAAA GCGCGTAACG CGCTCTTTCT TCGGAACATC
G10-d              TTCGAAGCAA CGCGAAGAAC CTTACCTGAT CTTGACATCC CGATGCCAAA GCGCGTAACG CGCTCTTTCT TCGGAACATC
GGCC_0168          TTCGAAGCAA CGCGAAGAAC CTTACCTGAT CTTGACATCC CGATGCCAAA GCGCGTAACG CGCTCTTTCT TCGGAACATC
I10-a              TTCGAAGCAA CGCGAAGAAC CTTACCTGAT CTTGACATCC CGATGCCAAA GCGCGTAACG CGCTCTTTCT TCGGAACATC
I10-b              TTCGAAGCAA CGCGAAGAAC CTTACCTGAT CTTGACATCC CGATGCCAAA GCGCGTAACG CGCTCTTTCT TCGGAACATC
I10-c              TTCGAAGCAA CGCGAAGAAC CTTACCTGAT CTTGACATCC CGATGCCAAA GCGCGTAACG CGCTCTTTCT TCGGAACATC
I10-d              TTCGAAGCAA CGCGAAGAAC CTTACCTGAT CTTGACATCC CGATGCCAAA GCGCGTAACG CGCTCTTTCT TCGGAACATC
JCM10422-a         TTCGAAGCAA CGCGAAGAAC CTTACCTGAT CTTGACATCC CGATGCCAAA GCGCGTAACG CGCTCTTTCT TCGGAACATC
JCM10422-b         TTCGAAGCAA CGCGAAGAAC CTTACCTGAT CTTGACATCC CGATGCCAAA GCGCGTAACG CGCTCTTTCT TCGGAACATC
JCM10422-c         TTCGAAGCAA CGCGAAGAAC CTTACCTGAT CTTGACATCC CGATGCCAAA GCGCGTAACG CGCTCTTTCT TCGGAACATC
JCM10422-d         TTCGAAGCAA CGCGAAGAAC CTTACCTGAT CTTGACATCC CGATGCCAAA GCGCGTAACG CGCTCTTTCT TCGGAACATC
MGYG-HGUT-01303-a  TTCGAAGCAA CGCGAAGAAC CTTACCTGAT CTTGACATCC CGATGCCAAA GCGCGTAACG CGCTCTTTCT TCGGAACATC
MGYG-HGUT-01303-b  TTCGAAGCAA CGCGAAGAAC CTTACCTGAT CTTGACATCC CGATGCCAAA GCGCGTAACG CGCTCTTTCT TCGGAACATC
MSK.1.16           TTCGAAGCAA CGCGAAGAAC CTTACCTGAT CTTGACATCC CGATGCCAAA GCGCGTAACG CGCTCTTTCT TCGGAACATC
MSK.1.26           TTCGAAGCAA CGCGAAGAAC CTTACCTGAT CTTGACATCC CGATGCCAAA GCGCGTAACG CGCTCTTTCT TCGGAACATC
S077-U-a           TTCGAAGCAA CGCGAAGAAC CTTACCTGAT CTTGACATCC CGATGCCAAA GCGCGTAACG CGCTCTTTCT TCGGAACATC
S077-U-b           TTCGAAGCAA CGCGAAGAAC CTTACCTGAT CTTGACATCC CGATGCCAAA GCGCGTAACG CGCTCTTTCT TCGGAACATC
S077-U-c           TTCGAAGCAA CGCGAAGAAC CTTACCTGAT CTTGACATCC CGATGCCAAA GCGCGTAACG CGCTCTTTCT TCGGAACATC
S077-U-d           TTCGAAGCAA CGCGAAGAAC CTTACCTGAT CTTGACATCC CGATGCCAAA GCGCGTAACG CGCTCTTTCT TCGGAACATC
VPI12708-a         TTCGAAGCAA CGCGAAGAAC CTTACCTGAT CTTGACATCC CGATGCCAAA GCGCGTAACG CGCTCTTTCT TCGGAACATC
VPI12708-b         TTCGAAGCAA CGCGAAGAAC CTTACCTGAT CTTGACATCC CGATGCCAAA GCGCGTAACG CGCTCTTTCT TCGGAACATC
VPI12708-c         TTCGAAGCAA CGCGAAGAAC CTTACCTGAT CTTGACATCC CGATGCCAAA GCGCGTAACG CGCTCTTTCT TCGGAACATC
VPI12708-d         TTCGAAGCAA CGCGAAGAAC CTTACCTGAT CTTGACATCC CGATGCCAAA GCGCGTAACG CGCTCTTTCT TCGGAACATC
DFI.1.130          TTCGAAGCAA CGCGAAGAAC CTTACCTGAT CTTGACATCC CGATGCCAAA GCGCGTAACG CGCTCTTTCT TCGGAACATC
DFI.1.161          ---------- ---------- ---------- ---------- ---------- ---------- ---------- ----------
DFI.1.162          TTCGAAGCAA CGCGAAGAAC CTTACCTGAT CTTGACATCC CGATGCCAAA GCGCGTAACG CGCTCTTTCT TCGGAACATC
DFI.1.217          TTCGAAGCAA CGCGAAGAAC CTTACCTGAT CTTGACATCC CGATGCCAAA GCGCGTAACG CGCTCTTTCT TCGGAACATC
DFI.1.234          TTCGAAGCAA CGCGAAGAAC CTTACCTGAT CTTGACATCC CGATGCCAAA GCGCGTAACG CGCTCTTTCT TCGGAACATC
DFI.1.60           TTCGAAGCAA CGCGAAGAAC CTTACCTGAT CTTGACATCC CGATGCCAAA GCGCGTAACG CGCTCTTTCT TCGGAACATC
DFI.4.63           TTCGAAGCAA CGCGAAGAAC CTTACCTGAT CTTGACATCC CGATGCCAAA GCGCGTAACG CGCTCTTTCT TCGGAACATC
JCM10419-a         TTCGAAGCAA CGCGAAGAAC CTTACCTGAT CTTGACATCC CGATGCCAAA GCGCGTAACG CGCTCTTTCT TCGGAACATC
JCM10419-b         TTCGAAGCAA CGCGAAGAAC CTTACCTGAT CTTGACATCC CGATGCCAAA GCGCGTAACG CGCTCTTTCT TCGGAACATC
JCM10419-c         TTCGAAGCAA CGCGAAGAAC CTTACCTGAT CTTGACATCC CGATGCCAAA GCGCGTAACG CGCTCTTTCT TCGGAACATC
JCM10419-d         TTCGAAGCAA CGCGAAGAAC CTTACCTGAT CTTGACATCC CGATGCCAAA GCGCGTAACG CGCTCTTTCT TCGGAACATC
JCM10420-a         TTCGAAGCAA CGCGAAGAAC CTTACCTGAT CTTGACATCC CGATGCCAAA GCGCGTAACG CGCTCTTTCT TCGGAACATC
JCM10420-b         TTCGAAGCAA CGCGAAGAAC CTTACCTGAT CTTGACATCC CGATGCCAAA GCGCGTAACG CGCTCTTTCT TCGGAACATC
JCM10420-c         TTCGAAGCAA CGCGAAGAAC CTTACCTGAT CTTGACATCC CGATGCCAAA GCGCGTAACG CGCTCTTTCT TCGGAACATC
JCM10420-d         TTCGAAGCAA CGCGAAGAAC CTTACCTGAT CTTGACATCC CGATGCCAAA GCGCGTAACG CGCTCTTTCT TCGGAACATC
JCM10423-a         TTCGAAGCAA CGCGAAGAAC CTTACCTGAT CTTGACATCC CGATGCCAAA GCGCGTAACG CGCTCTTTCT TCGGAACATC
JCM10423-b         TTCGAAGCAA CGCGAAGAAC CTTACCTGAT CTTGACATCC CGATGCCAAA GCGCGTAACG CGCTCTTTCT TCGGAACATC
JCM10423-c         TTCGAAGCAA CGCGAAGAAC CTTACCTGAT CTTGACATCC CGATGCCAAA GCGCGTAACG CGCTCTTTCT TCGGAACATC
JCM10423-d         TTCGAAGCAA CGCGAAGAAC CTTACCTGAT CTTGACATCC CGATGCCAAA GCGCGTAACG CGCTCTTTCT TCGGAACATC
MO32-a             TTCGAAGCAA CGCGAAGAAC CTTACCTGAT CTTGACATCC CGATGCCAAA GCGCGTAACG CGCTCTTTCT TCGGAACATC
MO32-b             TTCGAAGCAA CGCGAAGAAC CTTACCTGAT CTTGACATCC CGATGCCAAA GCGCGTAACG CGCTCTTTCT TCGGAACATC
MO32-c             TTCGAAGCAA CGCGAAGAAC CTTACCTGAT CTTGACATCC CGATGCCAAA GCGCGTAACG CGCTCTTTCT TCGGAACATC
MO32-d             TTCGAAGCAA CGCGAAGAAC CTTACCTGAT CTTGACATCC CGATGCCAAA GCGCGTAACG CGCTCTTTCT TCGGAACATC
MSK.5.24           TTCGAAGCAA CGCGAAGAAC CTTACCTGAT CTTGACATCC CGATGCCAAA GCGCGTAACG CGCTCTTTCT TCGGAACATC
NB2A-7-D5          TTCGAAGCAA CGCGAAGAAC CTTACCTGAT CTTGACATCC CGATGCCAAA GCGCGTAACG CGCTCTTTCT TCGGAACATC
NTI82-a            TTCGAAGCAA CGCGAAGAAC CTTACCTGAT CTTGACATCC CGATGCCAAA GCGCGTAACG CGCTCTTTCT TCGGAACATC
NTI82-b            TTCGAAGCAA CGCGAAGAAC CTTACCTGAT CTTGACATCC CGATGCCAAA GCGCGTAACG CGCTCTTTCT TCGGAACATC
NTI82-c            TTCGAAGCAA CGCGAAGAAC CTTACCTGAT CTTGACATCC CGATGCCAAA GCGCGTAACG CGCTCTTTCT TCGGAACATC
NTI82-d            TTCGAAGCAA CGCGAAGAAC CTTACCTGAT CTTGACATCC CGATGCCAAA GCGCGTAACG CGCTCTTTCT TCGGAACATC
Q4-a               TTCGAAGCAA CGCGAAGAAC CTTACCTGAT CTTGACATCC CGATGCCAAA GCGCGTAACG CGCTCTTTCT TCGGAACATC
Q4-b               TTCGAAGCAA CGCGAAGAAC CTTACCTGAT CTTGACATCC CGATGCCAAA GCGCGTAACG CGCTCTTTCT TCGGAACATC
Q4-c               TTCGAAGCAA CGCGAAGAAC CTTACCTGAT CTTGACATCC CGATGCCAAA GCGCGTAACG CGCTCTTTCT TCGGAACATC
Q4-d               TTCGAAGCAA CGCGAAGAAC CTTACCTGAT CTTGACATCC CGATGCCAAA GCGCGTAACG CGCTCTTTCT TCGGAACATC
S076-a             TTCGAAGCAA CGCGAAGAAC CTTACCTGAT CTTGACATCC CGATGCCAAA GCGCGTAACG CGCTCTTTCT TCGGAACATC
S076-b             TTCGAAGCAA CGCGAAGAAC CTTACCTGAT CTTGACATCC CGATGCCAAA GCGCGTAACG CGCTCTTTCT TCGGAACATC
S076-c             TTCGAAGCAA CGCGAAGAAC CTTACCTGAT CTTGACATCC CGATGCCAAA GCGCGTAACG CGCTCTTTCT TCGGAACATC
S076-d             TTCGAAGCAA CGCGAAGAAC CTTACCTGAT CTTGACATCC CGATGCCAAA GCGCGTAACG CGCTCTTTCT TCGGAACATC
SL.1.22            TTCGAAGCAA CGCGAAGAAC CTTACCTGAT CTTGACATCC CGATGCCAAA GCGCGTAACG CGCTCTTTCT TCGGAACATC
VE202-05           TTCGAAGCAA CGCGAAGAAC CTTACCTGAT CTTGACATCC CGATGCCAAA GCGCGTAACG CGCTCTTTCT TCGGAACATC
C.hylemonae        TTCGAAGCAA CGCGAAGAAC CTTACCTGAT CTTGACATCC CGGTGACAAA GTATGTAACG TACTCTTTCT TCGGAACACC
```

```
                        1041
35704-a              GGTGACAGGT GGTGCATGGT TGTCGTCAGC TCGTGTCGTG AGATGTTGGG TTAAGTCCCG CAACGAGCGC AACCCCTATC
35704-b              GGTGACAGGT GGTGCATGGT TGTCGTCAGC TCGTGTCGTG AGATGTTGGG TTAAGTCCCG CAACGAGCGC AACCCCTATC
35704-c              GGTGACAGGT GGTGCATGGT TGTCGTCAGC TCGTGTCGTG AGATGTTGGG TTAAGTCCCG CAACGAGCGC AACCCCTATC
35704-d              GGTGACAGGT GGTGCATGGT TGTCGTCAGC TCGTGTCGTG AGATGTTGGG TTAAGTCCCG CAACGAGCGC AACCCCTATC
AM05-22              GGTGACAGGT GGTGCATGGT TGTCGTCAGC TCGTGTCGTG AGATGTTGGG TTAAGTCCCG CAACGAGCGC AACCCCTATC
AM07-30              GGTGACAGGT GGTGCATGGT TGTCGTCAGC TCGTGTCGTG AGATGTTGGG TTAAGTCCCG CAACGAGCGC AACCCCTATC
BL389WT3D-a          GGTGACAGGT GGTGCATGGT TGTCGTCAGC TCGTGTCGTG AGATGTTGGG TTAAGTCCCG CAACGAGCGC AACCCCTATC
BL389WT3D-b          GGTGACAGGT GGTGCATGGT TGTCGTCAGC TCGTGTCGTG AGATGTTGGG TTAAGTCCCG CAACGAGCGC AACCCCTATC
BL389WT3D-c          GGTGACAGGT GGTGCATGGT TGTCGTCAGC TCGTGTCGTG AGATGTTGGG TTAAGTCCCG CAACGAGCGC AACCCCTATC
BL389WT3D-d          GGTGACAGGT GGTGCATGGT TGTCGTCAGC TCGTGTCGTG AGATGTTGGG TTAAGTCCCG CAACGAGCGC AACCCCTATC
CE91-St59-a          GGTGACAGGT GGTGCATGGT TGTCGTCAGC TCGTGTCGTG AGATGTTGGG TTAAGTCCCG CAACGAGCGC AACCCCTATC
CE91-St59-b          GGTGACAGGT GGTGCATGGT TGTCGTCAGC TCGTGTCGTG AGATGTTGGG TTAAGTCCCG CAACGAGCGC AACCCCTATC
CE91-St59-c          GGTGACAGGT GGTGCATGGT TGTCGTCAGC TCGTGTCGTG AGATGTTGGG TTAAGTCCCG CAACGAGCGC AACCCCTATC
CE91-St60-a          GGTGACAGGT GGTGCATGGT TGTCGTCAGC TCGTGTCGTG AGATGTTGGG TTAAGTCCCG CAACGAGCGC AACCCCTATC
CE91-St60-b          GGTGACAGGT GGTGCATGGT TGTCGTCAGC TCGTGTCGTG AGATGTTGGG TTAAGTCCCG CAACGAGCGC AACCCCTATC
CE91-St60-c          GGTGACAGGT GGTGCATGGT TGTCGTCAGC TCGTGTCGTG AGATGTTGGG TTAAGTCCCG CAACGAGCGC AACCCCTATC
FDAARGOS_1227-a      GGTGACAGGT GGTGCATGGT TGTCGTCAGC TCGTGTCGTG AGATGTTGGG TTAAGTCCCG CAACGAGCGC AACCCCTATC
FDAARGOS_1227-b      GGTGACAGGT GGTGCATGGT TGTCGTCAGC TCGTGTCGTG AGATGTTGGG TTAAGTCCCG CAACGAGCGC AACCCCTATC
FDAARGOS_1227-c      GGTGACAGGT GGTGCATGGT TGTCGTCAGC TCGTGTCGTG AGATGTTGGG TTAAGTCCCG CAACGAGCGC AACCCCTATC
FDAARGOS_1227-d      GGTGACAGGT GGTGCATGGT TGTCGTCAGC TCGTGTCGTG AGATGTTGGG TTAAGTCCCG CAACGAGCGC AACCCCTATC
G10-a                GGTGACAGGT GGTGCATGGT TGTCGTCAGC TCGTGTCGTG AGATGTTGGG TTAAGTCCCG CAACGAGCGC AACCCCTATC
G10-b                GGTGACAGGT GGTGCATGGT TGTCGTCAGC TCGTGTCGTG AGATGTTGGG TTAAGTCCCG CAACGAGCGC AACCCCTATC
G10-c                GGTGACAGGT GGTGCATGGT TGTCGTCAGC TCGTGTCGTG AGATGTTGGG TTAAGTCCCG CAACGAGCGC AACCCCTATC
G10-d                GGTGACAGGT GGTGCATGGT TGTCGTCAGC TCGTGTCGTG AGATGTTGGG TTAAGTCCCG CAACGAGCGC AACCCCTATC
GGCC_0168            GGTGACAGGT GGTGCATGGT TGTCGTCAGC TCGTGTCGTG AGATGTTGGG TTAAGTCCCG CAACGAGCGC AACCCCTATC
I10-a                GGTGACAGGT GGTGCATGGT TGTCGTCAGC TCGTGTCGTG AGATGTTGGG TTAAGTCCCG CAACGAGCGC AACCCCTATC
I10-b                GGTGACAGGT GGTGCATGGT TGTCGTCAGC TCGTGTCGTG AGATGTTGGG TTAAGTCCCG CAACGAGCGC AACCCCTATC
I10-c                GGTGACAGGT GGTGCATGGT TGTCGTCAGC TCGTGTCGTG AGATGTTGGG TTAAGTCCCG CAACGAGCGC AACCCCTATC
I10-d                GGTGACAGGT GGTGCATGGT TGTCGTCAGC TCGTGTCGTG AGATGTTGGG TTAAGTCCCG CAACGAGCGC AACCCCTATC
JCM10422-a           GGTGACAGGT GGTGCATGGT TGTCGTCAGC TCGTGTCGTG AGATGTTGGG TTAAGTCCCG CAACGAGCGC AACCCCTATC
JCM10422-b           GGTGACAGGT GGTGCATGGT TGTCGTCAGC TCGTGTCGTG AGATGTTGGG TTAAGTCCCG CAACGAGCGC AACCCCTATC
JCM10422-c           GGTGACAGGT GGTGCATGGT TGTCGTCAGC TCGTGTCGTG AGATGTTGGG TTAAGTCCCG CAACGAGCGC AACCCCTATC
JCM10422-d           GGTGACAGGT GGTGCATGGT TGTCGTCAGC TCGTGTCGTG AGATGTTGGG TTAAGTCCCG CAACGAGCGC AACCCCTATC
MGYG-HGUT-01303-a    GGTGACAGGT GGTGCATGGT TGTCGTCAGC TCGTGTCGTG AGATGTTGGG TTAAGTCCCG CAACGAGCGC AACCCCTATC
MGYG-HGUT-01303-b    GGTGACAGGT GGTGCATGGT TGTCGTCAGC TCGTGTCGTG AGATGTTGGG TTAAGTCCCG CAACGAGCGC AACCCCTATC
MSK.1.16             GGTGACAGGT GGTGCATGGT TGTCGTCAGC TCGTGTCGTG AGATGTTGGG TTAAGTCCCG CAACGAGCGC AACCCCTATC
MSK.1.26             GGTGACAGGT GGTGCATGGT TGTCGTCAGC TCGTGTCGTG AGATGTTGGG TTAAGTC--- ---------- ----------
S077-U-a             GGTGACAGGT GGTGCATGGT TGTCGTCAGC TCGTGTCGTG AGATGTTGGG TTAAGTCCCG CAACGAGCGC AACCCCTATC
S077-U-b             GGTGACAGGT GGTGCATGGT TGTCGTCAGC TCGTGTCGTG AGATGTTGGG TTAAGTCCCG CAACGAGCGC AACCCCTATC
S077-U-c             GGTGACAGGT GGTGCATGGT TGTCGTCAGC TCGTGTCGTG AGATGTTGGG TTAAGTCCCG CAACGAGCGC AACCCCTATC
S077-U-d             GGTGACAGGT GGTGCATGGT TGTCGTCAGC TCGTGTCGTG AGATGTTGGG TTAAGTCCCG CAACGAGCGC AACCCCTATC
VPI12708-a           GGTGACAGGT GGTGCATGGT TGTCGTCAGC TCGTGTCGTG AGATGTTGGG TTAAGTCCCG CAACGAGCGC AACCCCTATC
VPI12708-b           GGTGACAGGT GGTGCATGGT TGTCGTCAGC TCGTGTCGTG AGATGTTGGG TTAAGTCCCG CAACGAGCGC AACCCCTATC
VPI12708-c           GGTGACAGGT GGTGCATGGT TGTCGTCAGC TCGTGTCGTG AGATGTTGGG TTAAGTCCCG CAACGAGCGC AACCCCTATC
VPI12708-d           GGTGACAGGT GGTGCATGGT TGTCGTCAGC TCGTGTCGTG AGATGTTGGG TTAAGTCCCG CAACGAGCGC AACCCCTATC
DFI.1.130            GGTGACAGGT GGTGCATGGT TGTCGTCAGC TCGTGTCGTG AGATGTTGGG TTAAGTCCCG CAACGAGCGC AACCCCTATC
DFI.1.161            ---------- ---------- ---------- ---------- ---------- ---------- ---------- ----------
DFI.1.162            GGTGACAGGT GGTGCATGGT TGTCGTCAGC TCGTGTCGTG AGATGTTGGG TTAAGTCCCG CAACGAGCGC AACCCCTATC
DFI.1.217            GGTGACAGGT GGTGCATGGT TGTCGTCAGC TCGTGTCGTG AGATGTTGGG TTAAGTCCCG CAACGAGCGC AACCCCTATC
DFI.1.234            GGTGACAGGT GGTGCATGGT TGTCGTCAGC TCGTGTCGTG AGATGTTGGG TTAAGTCCCG CAACGAGCGC AACCCCTATC
DFI.1.60             GGTGACAGGT GGTGCATGGT TGTCGTCAGC TCGTGTCGTG AGATGTTGGG TTAAGTCCCG CAACGAGCGC AACCCCTATC
DFI.4.63             GGTGACAGGT GGTGCATGGT TGTCGTCAGC TCGTGTCGTG AGATGTTGGG TTAAGTCCCG CAACGAGCGC AACCCCTATC
JCM10419-a           GGTGACAGGT GGTGCATGGT TGTCGTCAGC TCGTGTCGTG AGATGTTGGG TTAAGTCCCG CAACGAGCGC AACCCCTATC
JCM10419-b           GGTGACAGGT GGTGCATGGT TGTCGTCAGC TCGTGTCGTG AGATGTTGGG TTAAGTCCCG CAACGAGCGC AACCCCTATC
JCM10419-c           GGTGACAGGT GGTGCATGGT TGTCGTCAGC TCGTGTCGTG AGATGTTGGG TTAAGTCCCG CAACGAGCGC AACCCCTATC
JCM10419-d           GGTGACAGGT GGTGCATGGT TGTCGTCAGC TCGTGTCGTG AGATGTTGGG TTAAGTCCCG CAACGAGCGC AACCCCTATC
JCM10420-a           GGTGACAGGT GGTGCATGGT TGTCGTCAGC TCGTGTCGTG AGATGTTGGG TTAAGTCCCG CAACGAGCGC AACCCCTATC
JCM10420-b           GGTGACAGGT GGTGCATGGT TGTCGTCAGC TCGTGTCGTG AGATGTTGGG TTAAGTCCCG CAACGAGCGC AACCCCTATC
JCM10420-c           GGTGACAGGT GGTGCATGGT TGTCGTCAGC TCGTGTCGTG AGATGTTGGG TTAAGTCCCG CAACGAGCGC AACCCCTATC
JCM10420-d           GGTGACAGGT GGTGCATGGT TGTCGTCAGC TCGTGTCGTG AGATGTTGGG TTAAGTCCCG CAACGAGCGC AACCCCTATC
JCM10423-a           GGTGACAGGT GGTGCATGGT TGTCGTCAGC TCGTGTCGTG AGATGTTGGG TTAAGTCCCG CAACGAGCGC AACCCCTATC
JCM10423-b           GGTGACAGGT GGTGCATGGT TGTCGTCAGC TCGTGTCGTG AGATGTTGGG TTAAGTCCCG CAACGAGCGC AACCCCTATC
JCM10423-c           GGTGACAGGT GGTGCATGGT TGTCGTCAGC TCGTGTCGTG AGATGTTGGG TTAAGTCCCG CAACGAGCGC AACCCCTATC
JCM10423-d           GGTGACAGGT GGTGCATGGT TGTCGTCAGC TCGTGTCGTG AGATGTTGGG TTAAGTCCCG CAACGAGCGC AACCCCTATC
MO32-a               GGTGACAGGT GGTGCATGGT TGTCGTCAGC TCGTGTCGTG AGATGTTGGG TTAAGTCCCG CAACGAGCGC AACCCCTATC
MO32-b               GGTGACAGGT GGTGCATGGT TGTCGTCAGC TCGTGTCGTG AGATGTTGGG TTAAGTCCCG CAACGAGCGC AACCCCTATC
MO32-c               GGTGACAGGT GGTGCATGGT TGTCGTCAGC TCGTGTCGTG AGATGTTGGG TTAAGTCCCG CAACGAGCGC AACCCCTATC
MO32-d               GGTGACAGGT GGTGCATGGT TGTCGTCAGC TCGTGTCGTG AGATGTTGGG TTAAGTCCCG CAACGAGCGC AACCCCTATC
MSK.5.24             GGTGACAGGT GGTGCATGGT TGTCGTCAGC TCGTGTCGTG AGATGTTGGG TTAAGTCCCG CAACGAGCGC AACCCCTATC
NB2A-7-D5            GGTGACAGGT GGTGCATGGT TGTCGTCAGC TCGTGTCGTG AGATGTTGGG TTAAGTCCCG CAACGAGCGC AACCCCTATC
NTI82-a              GGTGACAGGT GGTGCATGGT TGTCGTCAGC TCGTGTCGTG AGATGTTGGG TTAAGTCCCG CAACGAGCGC AACCCCTATC
NTI82-b              GGTGACAGGT GGTGCATGGT TGTCGTCAGC TCGTGTCGTG AGATGTTGGG TTAAGTCCCG CAACGAGCGC AACCCCTATC
NTI82-c              GGTGACAGGT GGTGCATGGT TGTCGTCAGC TCGTGTCGTG AGATGTTGGG TTAAGTCCCG CAACGAGCGC AACCCCTATC
NTI82-d              GGTGACAGGT GGTGCATGGT TGTCGTCAGC TCGTGTCGTG AGATGTTGGG TTAAGTCCCG CAACGAGCGC AACCCCTATC
Q4-a                 GGTGACAGGT GGTGCATGGT TGTCGTCAGC TCGTGTCGTG AGATGTTGGG TTAAGTCCCG CAACGAGCGC AACCCCTATC
Q4-b                 GGTGACAGGT GGTGCATGGT TGTCGTCAGC TCGTGTCGTG AGATGTTGGG TTAAGTCCCG CAACGAGCGC AACCCCTATC
Q4-c                 GGTGACAGGT GGTGCATGGT TGTCGTCAGC TCGTGTCGTG AGATGTTGGG TTAAGTCCCG CAACGAGCGC AACCCCTATC
Q4-d                 GGTGACAGGT GGTGCATGGT TGTCGTCAGC TCGTGTCGTG AGATGTTGGG TTAAGTCCCG CAACGAGCGC AACCCCTATC
S076-a               GGTGACAGGT GGTGCATGGT TGTCGTCAGC TCGTGTCGTG AGATGTTGGG TTAAGTCCCG CAACGAGCGC AACCCCTATC
S076-b               GGTGACAGGT GGTGCATGGT TGTCGTCAGC TCGTGTCGTG AGATGTTGGG TTAAGTCCCG CAACGAGCGC AACCCCTATC
S076-c               GGTGACAGGT GGTGCATGGT TGTCGTCAGC TCGTGTCGTG AGATGTTGGG TTAAGTCCCG CAACGAGCGC AACCCCTATC
S076-d               GGTGACAGGT GGTGCATGGT TGTCGTCAGC TCGTGTCGTG AGATGTTGGG TTAAGTCCCG CAACGAGCGC AACCCCTATC
SL.1.22              GGTGACAGGT GGTGCATGGT TGTCGTCAGC TCGTGTCGTG AGATGTTGGG TTAAGTCCCG CAACGAGCGC AACCCCTATC
VE202-05             GGTGACAGGT GGTGCATGGT TGTCGTCAGC TCGTGTCGTG AGATGTTGGG TTAAGTCCCG CAACGAGCGC AACCCCTATC
C.hylemonae          GGTGACAGGT GGTGCATGGT TGTCGTCAGC TCGTGTCGTG AGATGTTGGG TTAAGTCCCG CAACGAGCGC AACCCTTATC
```

```
                     1121
35704-a              TTCAGTAGCC AGCATTTTGG ATGGGCACTC TGGAGAGACT GCCAGGGAGA ACCTGGAGGA AGGTGGGGAT GACGTCAAAT
35704-b              TTCAGTAGCC AGCATTTTGG ATGGGCACTC TGGAGAGACT GCCAGGGAGA ACCTGGAGGA AGGTGGGGAT GACGTCAAAT
35704-c              TTCAGTAGCC AGCATTTTGG ATGGGCACTC TGGAGAGACT GCCAGGGAGA ACCTGGAGGA AGGTGGGGAT GACGTCAAAT
35704-d              TTCAGTAGCC AGCATTTTGG ATGGGCACTC TGGAGAGACT GCCAGGGAGA ACCTGGAGGA AGGTGGGGAT GACGTCAAAT
AM05-22              TTCAGTAGCC AGCATTTTGG ATGGGCACTC TGGAGAGACT GCCAGGGAGA ACCTGGAGGA AGGTGGGGAT GACGTCAAAT
AM07-30              TTCAGTAGCC AGCATTTTGG ATGGGCACTC TGGAGAGACT GCCAGGGAGA ACCTGGAGGA AGGTGGGGAT GACGTCAAAT
BL389WT3D-a          TTCAGTAGCC AGCATTATGG ATGGGCACTC TGGAGAGACT GCCAGGGAGA ACCTGGAGGA AGGTGGGGAT GACGTCAAAT
BL389WT3D-b          TTCAGTAGCC AGCATTATGG ATGGGCACTC TGGAGAGACT GCCAGGGAGA ACCTGGAGGA AGGTGGGGAT GACGTCAAAT
BL389WT3D-c          TTCAGTAGCC AGCATTATGG ATGGGCACTC TGGAGAGACT GCCAGGGAGA ACCTGGAGGA AGGTGGGGAT GACGTCAAAT
BL389WT3D-d          TTCAGTAGCC AGCATTATGG ATGGGCACTC TGGAGAGACT GCCAGGGAGA ACCTGGAGGA AGGTGGGGAT GACGTCAAAT
CE91-St59-a          TTCAGTAGCC AGCATTTTGG ATGGGCACTC TGGAGAGACT GCCAGGGAGA ACCTGGAGGA AGGTGGGGAT GACGTCAAAT
CE91-St59-b          TTCAGTAGCC AGCATTTTGG ATGGGCACTC TGGAGAGACT GCCAGGGAGA ACCTGGAGGA AGGTGGGGAT GACGTCAAAT
CE91-St59-c          TTCAGTAGCC AGCATTTTGG ATGGGCACTC TGGAGAGACT GCCAGGGAGA ACCTGGAGGA AGGTGGGGAT GACGTCAAAT
CE91-St60-a          TTCAGTAGCC AGCATTTTGG ATGGGCACTC TGGAGAGACT GCCAGGGAGA ACCTGGAGGA AGGTGGGGAT GACGTCAAAT
CE91-St60-b          TTCAGTAGCC AGCATTTTGG ATGGGCACTC TGGAGAGACT GCCAGGGAGA ACCTGGAGGA AGGTGGGGAT GACGTCAAAT
CE91-St60-c          TTCAGTAGCC AGCATTTTGG ATGGGCACTC TGGAGAGACT GCCAGGGAGA ACCTGGAGGA AGGTGGGGAT GACGTCAAAT
FDAARGOS_1227-a      TTCAGTAGCC AGCATTTTGG ATGGGCACTC TGGAGAGACT GCCAGGGAGA ACCTGGAGGA AGGTGGGGAT GACGTCAAAT
FDAARGOS_1227-b      TTCAGTAGCC AGCATTTTGG ATGGGCACTC TGGAGAGACT GCCAGGGAGA ACCTGGAGGA AGGTGGGGAT GACGTCAAAT
FDAARGOS_1227-c      TTCAGTAGCC AGCATTTTGG ATGGGCACTC TGGAGAGACT GCCAGGGAGA ACCTGGAGGA AGGTGGGGAT GACGTCAAAT
FDAARGOS_1227-d      TTCAGTAGCC AGCATTTTGG ATGGGCACTC TGGAGAGACT GCCAGGGAGA ACCTGGAGGA AGGTGGGGAT GACGTCAAAT
G10-a                TTCAGTAGCC AGCATTATGG ATGGGCACTC TGGAGAGACT GCCAGGGAGA ACCTGGAGGA AGGTGGGGAT GACGTCAAAT
G10-b                TTCAGTAGCC AGCATTATGG ATGGGCACTC TGGAGAGACT GCCAGGGAGA ACCTGGAGGA AGGTGGGGAT GACGTCAAAT
G10-c                TTCAGTAGCC AGCATTATGG ATGGGCACTC TGGAGAGACT GCCAGGGAGA ACCTGGAGGA AGGTGGGGAT GACGTCAAAT
G10-d                TTCAGTAGCC AGCATTATGG ATGGGCACTC TGGAGAGACT GCCAGGGAGA ACCTGGAGGA AGGTGGGGAT GACGTCAAAT
GGCC_0168            TTCAGTAGCC AGCATTATGG ATGGGCACTC TGGAGAGACT GCCAGGGAGA ACCTGGAGGA AGGTGGGGAT GACGTCAAAT
I10-a                TTCAGTAGCC AGCATTATGG ATGGGCACTC TGGAGAGACT GCCAGGGAGA ACCTGGAGGA AGGTGGGGAT GACGTCAAAT
I10-b                TTCAGTAGCC AGCATTATGG ATGGGCACTC TGGAGAGACT GCCAGGGAGA ACCTGGAGGA AGGTGGGGAT GACGTCAAAT
I10-c                TTCAGTAGCC AGCATTATGG ATGGGCACTC TGGAGAGACT GCCAGGGAGA ACCTGGAGGA AGGTGGGGAT GACGTCAAAT
I10-d                TTCAGTAGCC AGCATTATGG ATGGGCACTC TGGAGAGACT GCCAGGGAGA ACCTGGAGGA AGGTGGGGAT GACGTCAAAT
JCM10422-a           TTCAGTAGCC AGCATTATGG ATGGGCACTC TGGAGAGACT GCCAGGGAGA ACCTGGAGGA AGGTGGGGAT GACGTCAAAT
JCM10422-b           TTCAGTAGCC AGCATTATGG ATGGGCACTC TGGAGAGACT GCCAGGGAGA ACCTGGAGGA AGGTGGGGAT GACGTCAAAT
JCM10422-c           TTCAGTAGCC AGCATTATGG ATGGGCACTC TGGAGAGACT GCCAGGGAGA ACCTGGAGGA AGGTGGGGAT GACGTCAAAT
JCM10422-d           TTCAGTAGCC AGCATTATGG ATGGGCACTC TGGAGAGACT GCCAGGGAGA ACCTGGAGGA AGGTGGGGAT GACGTCAAAT
MGYG-HGUT-01303-a    TTCAGTAGCC AGCATTTTGG ATGGGCACTC TGGAGAGACT GCCAGGGAGA ACCTGGAGGA AGGTGGGGAT GACGTCAAAT
MGYG-HGUT-01303-b    TTCAGTAGCC AGCATTTTGG ATGGGCACTC TGGAGAGACT GCCAGGGAGA ACCTGGAGGA AGGTGGGGAT GACGTCAAAT
MSK.1.16             TTCAGTAGCC AGCATTTTGG ATGGGCACTC TGGAGAGACT GCCAGGGAGA ACCTGGAGGA AGGTGGGGAT GACGTCAAAT
MSK.1.26             ---------- ---------- ---------- ---------- ---------- ---------- ---------- ----------
S077-U-a             TTCAGTAGCC AGCATTTTGG ATGGGCACTC TGGAGAGACT GCCAGGGAGA ACCTGGAGGA AGGTGGGGAT GACGTCAAAT
S077-U-b             TTCAGTAGCC AGCATTTTGG ATGGGCACTC TGGAGAGACT GCCAGGGAGA ACCTGGAGGA AGGTGGGGAT GACGTCAAAT
S077-U-c             TTCAGTAGCC AGCATTTTGG ATGGGCACTC TGGAGAGACT GCCAGGGAGA ACCTGGAGGA AGGTGGGGAT GACGTCAAAT
S077-U-d             TTCAGTAGCC AGCATTTTGG ATGGGCACTC TGGAGAGACT GCCAGGGAGA ACCTGGAGGA AGGTGGGGAT GACGTCAAAT
VPI12708-a           TTCAGTAGCC AGCATTTCGG ATGGGCACTC TGGAGAGACT GCCAGGGACA ACCTGGAGGA AGGTGGGGAT GACGTCAAAT
VPI12708-b           TTCAGTAGCC AGCATTCCGG ATGGGCACTC TGGAGAGACT GCCAGGGACA ACCTGGAGGA AGGTGGGGAT GACGTCAAAT
VPI12708-c           TTCAGTAGCC AGCATTCCGG ATGGGCACTC TGGAGAGACT GCCAGGGACA ACCTGGAGGA AGGTGGGGAT GACGTCAAAT
VPI12708-d           TTCAGTAGCC AGCATTTTGG ATGGGCACTC TGGAGAGACT GCCAGGGACA ACCTGGAGGA AGGTGGGGAT GACGTCAAAT
DFI.1.130            TTCAGTAGCC AGCATTCCGG ATGGGCACTC TGGAGAGACT GCCAGGGACA ACCTGGAGGA AGGTGGGGAT GACGTCAAAT
DFI.1.161            ---------- ---------- ---------- ---------- ---------- ---------- ---------- ----------
DFI.1.162            TTCAGTAGCC AGCATTTCGG ATGGGCACTC TGGAGAGACT GCCAGGGACA ACCTGGAGGA AGGTGGGGAT GACGTCAAAT
DFI.1.217            TTCAGTAGCC AGCATTTCGG ATGGGCACTC TGGAGAGACT GCCAGGGACA ACCTGGAGGA AGGTGGGGAT GACGTCAAAT
DFI.1.234            TTCAGTAGCC AGCATTTCGG ATGGGCACTC TGGAGAGACT GCCAGGGACA ACCTGGAGGA AGGTGGGGAT GACGTCAAAT
DFI.1.60             TTCAGTAGCC AGCATTTCGG ATGGGCACTC TGGAGAGACT GCCAGGGACA ACCTGGAGGA AGGTGGGGAT GACGTCAAAT
DFI.4.63             TTCAGTAGCC AGCATTCCGG ATGGGCACTC TGGAGAGACT GCCAGGGACA ACCTGGAGGA AGGTGGGGAT GACGTCAAAT
JCM10419-a           TTCAGTAGCC AGCATTCCGG ATGGGCACTC TGGAGAGACT GCCAGGGACA ACCTGGAGGA AGGTGGGGAT GACGTCAAAT
JCM10419-b           TTCAGTAGCC AGCATTTTGG ATGGGCACTC TGGAGAGACT GCCAGGGACA ACCTGGAGGA AGGTGGGGAT GACGTCAAAT
JCM10419-c           TTCAGTAGCC AGCATTCCGG ATGGGCACTC TGGAGAGACT GCCAGGGACA ACCTGGAGGA AGGTGGGGAT GACGTCAAAT
JCM10419-d           TTCAGTAGCC AGCATTTCGG ATGGGCACTC TGGAGAGACT GCCAGGGACA ACCTGGAGGA AGGTGGGGAT GACGTCAAAT
JCM10420-a           TTCAGTAGCC AGCATTTCGG ATGGGCACTC TGGAGAGACT GCCAGGGACA ACCTGGAGGA AGGTGGGGAT GACGTCAAAT
JCM10420-b           TTCAGTAGCC AGCATTCCGG ATGGGCACTC TGGAGAGACT GCCAGGGACA ACCTGGAGGA AGGTGGGGAT GACGTCAAAT
JCM10420-c           TTCAGTAGCC AGCATTTTGG ATGGGCACTC TGGAGAGACT GCCAGGGACA ACCTGGAGGA AGGTGGGGAT GACGTCAAAT
JCM10420-d           TTCAGTAGCC AGCATTCCGG ATGGGCACTC TGGAGAGACT GCCAGGGACA ACCTGGAGGA AGGTGGGGAT GACGTCAAAT
JCM10423-a           TTCAGTAGCC AGCATTCCGG ATGGGCACTC TGGAGAGACT GCCAGGGACA ACCTGGAGGA AGGTGGGGAT GACGTCAAAT
JCM10423-b           TTCAGTAGCC AGCATTTCGG ATGGGCACTC TGGAGAGACT GCCAGGGACA ACCTGGAGGA AGGTGGGGAT GACGTCAAAT
JCM10423-c           TTCAGTAGCC AGCATTTTGG ATGGGCACTC TGGAGAGACT GCCAGGGACA ACCTGGAGGA AGGTGGGGAT GACGTCAAAT
JCM10423-d           TTCAGTAGCC AGCATTCCGG ATGGGCACTC TGGAGAGACT GCCAGGGACA ACCTGGAGGA AGGTGGGGAT GACGTCAAAT
MO32-a               TTCAGTAGCC AGCATTTCGG ATGGGCACTC TGGAGAGACT GCCAGGGACA ACCTGGAGGA AGGTGGGGAT GACGTCAAAT
MO32-b               TTCAGTAGCC AGCATTTCGG ATGGGCACTC TGGAGAGACT GCCAGGGACA ACCTGGAGGA AGGTGGGGAT GACGTCAAAT
MO32-c               TTCAGTAGCC AGCATTTCGG ATGGGCACTC TGGAGAGACT GCCAGGGACA ACCTGGAGGA AGGTGGGGAT GACGTCAAAT
MO32-d               TTCAGTAGCC AGCATTTCGG ATGGGCACTC TGGAGAGACT GCCAGGGACA ACCTGGAGGA AGGTGGGGAT GACGTCAAAT
MSK.5.24             TTCAGTAGCC AGCATTCCGG ATGGGCACTC TGGAGAGACT GCCAGGGACA ACCTGGAGGA AGGTGGGGAT GACGTCAAAT
NB2A-7-D5            TTCAGTAGCC AGCATTCCGG ATGGGCACTC TGGAGAGACT GCCAGGGACA ACCTGGAGGA AGGTGGGGAT GACGTCAAAT
NTI82-a              TTCAGTAGCC AGCATTCCGG ATGGGCACTC TGGAGAGACT GCCAGGGACA ACCTGGAGGA AGGTGGGGAT GACGTCAAAT
NTI82-b              TTCAGTAGCC AGCATTCCGG ATGGGCACTC TGGAGAGACT GCCAGGGACA ACCTGGAGGA AGGTGGGGAT GACGTCAAAT
NTI82-c              TTCAGTAGCC AGCATTCCGG ATGGGCACTC TGGAGAGACT GCCAGGGACA ACCTGGAGGA AGGTGGGGAT GACGTCAAAT
NTI82-d              TTCAGTAGCC AGCATTCCGG ATGGGCACTC TGGAGAGACT GCCAGGGACA ACCTGGAGGA AGGTGGGGAT GACGTCAAAT
Q4-a                 TTCAGTAGCC AGCATTTCGG ATGGGCACTC TGGAGAGACT GCCAGGGACA ACCTGGAGGA AGGTGGGGAT GACGTCAAAT
Q4-b                 TTCAGTAGCC AGCATTTCGG ATGGGCACTC TGGAGAGACT GCCAGGGACA ACCTGGAGGA AGGTGGGGAT GACGTCAAAT
Q4-c                 TTCAGTAGCC AGCATTTCGG ATGGGCACTC TGGAGAGACT GCCAGGGACA ACCTGGAGGA AGGTGGGGAT GACGTCAAAT
Q4-d                 TTCAGTAGCC AGCATTTCGG ATGGGCACTC TGGAGAGACT GCCAGGGACA ACCTGGAGGA AGGTGGGGAT GACGTCAAAT
S076-a               TTCAGTAGCC AGCATTTCGG ATGGGCACTC TGGAGAGACT GCCAGGGACA ACCTGGAGGA AGGTGGGGAT GACGTCAAAT
S076-b               TTCAGTAGCC AGCATTTCGG ATGGGCACTC TGGAGAGACT GCCAGGGACA ACCTGGAGGA AGGTGGGGAT GACGTCAAAT
S076-c               TTCAGTAGCC AGCATTTCGG ATGGGCACTC TGGAGAGACT GCCAGGGACA ACCTGGAGGA AGGTGGGGAT GACGTCAAAT
S076-d               TTCAGTAGCC AGCATTTCGG ATGGGCACTC TGGAGAGACT GCCAGGGACA ACCTGGAGGA AGGTGGGGAT GACGTCAAAT
SL.1.22              TTCAGTAGCC AGCATTTTGG ATGGGCACTC TGGAGAGACT GCCAGGGACA ACCTGGAGGA AGGTGGGGAT GACGTCAAAT
VE202-05             TTCAGTAGCC AGCATTCCGG ATGGGCACTC TGGAGAGACT GCCAGGGACA ACCTGGAGGA AGGTGGGGAT GACGTCAAAT
C.hylemonae          TTTAGTAGCC AGCATTTGAG GTGGGCACTC TAGAGAGACT GCCAGGGATA ACCTGGAGGA AGGTGGGGAT GACGTCAAAT
```

```
                         1201
35704-a              CATCATGCCC CTTATGACCA GGGCTACACA CGTGCTACAA TGGCGTAAAC AAAGGGAGGC GAACCCGCGA GGGTGGGCAA
35704-b              CATCATGCCC CTTATGACCA GGGCTACACA CGTGCTACAA TGGCGTAAAC AAAGGGAGGC GAACCCGCGA GGGTGGGCAA
35704-c              CATCATGCCC CTTATGACCA GGGCTACACA CGTGCTACAA TGGCGTAAAC AAAGGGAGGC GAACCCGCGA GGGTGGGCAA
35704-d              CATCATGCCC CTTATGACCA GGGCTACACA CGTGCTACAA TGGCGTAAAC AAAGGGAGGC GAACCCGCGA GGGTGGGCAA
AM05-22              CATCATGCCC CTTATGACCA GGGCTACACA CGTGCTACAA TGGCGTAAAC AAAGGGAGGC GAACCCGCGA GGGTGGGCAA
AM07-30              CATCATGCCC CTTATGACCA GGGCTACACA CGTGCTACAA TGGCGTAAAC AAAGGGAGGC GAACCCGCGA GGGTGGGCAA
BL389WT3D-a          CATCATGCCC CTTATGACCA GGGCTACACA CGTGCTACAA TGGCGTAAAC AAAGGGAGGC GAACCCGCGA GGGTGGGCAA
BL389WT3D-b          CATCATGCCC CTTATGACCA GGGCTACACA CGTGCTACAA TGGCGTAAAC AAAGGGAGGC GAACCCGCGA GGGTGGGCAA
BL389WT3D-c          CATCATGCCC CTTATGACCA GGGCTACACA CGTGCTACAA TGGCGTAAAC AAAGGGAGGC GAACCCGCGA GGGTGGGCAA
BL389WT3D-d          CATCATGCCC CTTATGACCA GGGCTACACA CGTGCTACAA TGGCGTAAAC AAAGGGAGGC GAACCCGCGA GGGTGGGCAA
CE91-St59-a          CATCATGCCC CTTATGACCA GGGCTACACA CGTGCTACAA TGGCGTAAAC AAAGGGAGGC GAACCCGCGA GGGTGGGCAA
CE91-St59-b          CATCATGCCC CTTATGACCA GGGCTACACA CGTGCTACAA TGGCGTAAAC AAAGGGAGGC GAACCCGCGA GGGTGGGCAA
CE91-St59-c          CATCATGCCC CTTATGACCA GGGCTACACA CGTGCTACAA TGGCGTAAAC AAAGGGAGGC GAACCCGCGA GGGTGGGCAA
CE91-St60-a          CATCATGCCC CTTATGACCA GGGCTACACA CGTGCTACAA TGGCGTAAAC AAAGGGAGGC GAACCCGCGA GGGTGGGCAA
CE91-St60-b          CATCATGCCC CTTATGACCA GGGCTACACA CGTGCTACAA TGGCGTAAAC AAAGGGAGGC GAACCCGCGA GGGTGGGCAA
CE91-St60-c          CATCATGCCC CTTATGACCA GGGCTACACA CGTGCTACAA TGGCGTAAAC AAAGGGAGGC GAACCCGCGA GGGTGGGCAA
FDAARGOS_1227-a      CATCATGCCC CTTATGACCA GGGCTACACA CGTGCTACAA TGGCGTAAAC AAAGGGAGGC GAACCCGCGA GGGTGGGCAA
FDAARGOS_1227-b      CATCATGCCC CTTATGACCA GGGCTACACA CGTGCTACAA TGGCGTAAAC AAAGGGAGGC GAACCCGCGA GGGTGGGCAA
FDAARGOS_1227-c      CATCATGCCC CTTATGACCA GGGCTACACA CGTGCTACAA TGGCGTAAAC AAAGGGAGGC GAACCCGCGA GGGTGGGCAA
FDAARGOS_1227-d      CATCATGCCC CTTATGACCA GGGCTACACA CGTGCTACAA TGGCGTAAAC AAAGGGAGGC GAACCCGCGA GGGTGGGCAA
G10-a                CATCATGCCC CTTATGACCA GGGCTACACA CGTGCTACAA TGGCGTAAAC AAAGGGAGGC GAACCCGCGA GGGTGGGCAA
G10-b                CATCATGCCC CTTATGACCA GGGCTACACA CGTGCTACAA TGGCGTAAAC AAAGGGAGGC GAACCCGCGA GGGTGGGCAA
G10-c                CATCATGCCC CTTATGACCA GGGCTACACA CGTGCTACAA TGGCGTAAAC AAAGGGAGGC GAACCCGCGA GGGTGGGCAA
G10-d                CATCATGCCC CTTATGACCA GGGCTACACA CGTGCTACAA TGGCGTAAAC AAAGGGAGGC GAACCCGCGA GGGTGGGCAA
GGCC_0168            CATCATGCCC CTTATGACCA GGGCTACACA CGTGCTACAA TGGCGTAAAC AAAGGGAGGC GAACCCGCGA GGGTGGGCAA
I10-a                CATCATGCCC CTTATGACCA GGGCTACACA CGTGCTACAA TGGCGTAAAC AAAGGGAGGC GAACCCGCGA GGGTGGGCAA
I10-b                CATCATGCCC CTTATGACCA GGGCTACACA CGTGCTACAA TGGCGTAAAC AAAGGGAGGC GAACCCGCGA GGGTGGGCAA
I10-c                CATCATGCCC CTTATGACCA GGGCTACACA CGTGCTACAA TGGCGTAAAC AAAGGGAGGC GAACCCGCGA GGGTGGGCAA
I10-d                CATCATGCCC CTTATGACCA GGGCTACACA CGTGCTACAA TGGCGTAAAC AAAGGGAGGC GAACCCGCGA GGGTGGGCAA
JCM10422-a           CATCATGCCC CTTATGACCA GGGCTACACA CGTGCTACAA TGGCGTAAAC AAAGGGAGGC GAACCCGCGA GGGTGGGCAA
JCM10422-b           CATCATGCCC CTTATGACCA GGGCTACACA CGTGCTACAA TGGCGTAAAC AAAGGGAGGC GAACCCGCGA GGGTGGGCAA
JCM10422-c           CATCATGCCC CTTATGACCA GGGCTACACA CGTGCTACAA TGGCGTAAAC AAAGGGAGGC GAACCCGCGA GGGTGGGCAA
JCM10422-d           CATCATGCCC CTTATGACCA GGGCTACACA CGTGCTACAA TGGCGTAAAC AAAGGGAGGC GAACCCGCGA GGGTGGGCAA
MGYG-HGUT-01303-a    CATCATGCCC CTTATGACCA GGGCTACACA CGTGCTACAA TGGCGTAAAC AAAGGGAGGC GAACCCGCGA GGGTGGGCAA
MGYG-HGUT-01303-b    CATCATGCCC CTTATGACCA GGGCTACACA CGTGCTACAA TGGCGTAAAC AAAGGGAGGC GAACCCGCGA GGGTGGGCAA
MSK.1.16             CATCATGCCC CTTATGACCA GGGCTACACA CGTGCTACAA TGGCGTAAAC AAAGGGAGGC GAACCCGCGA GGGTGGGCAA
MSK.1.26             ---------- ---------- ---------- ---------- ---------- ---------- ---------- ----------
S077-U-a             CATCATGCCC CTTATGACCA GGGCTACACA CGTGCTACAA TGGCGTAAAC AAAGGGAGGC GAACCCGCGA GGGTGGGCAA
S077-U-b             CATCATGCCC CTTATGACCA GGGCTACACA CGTGCTACAA TGGCGTAAAC AAAGGGAGGC GAACCCGCGA GGGTGGGCAA
S077-U-c             CATCATGCCC CTTATGACCA GGGCTACACA CGTGCTACAA TGGCGTAAAC AAAGGGAGGC GAACCCGCGA GGGTGGGCAA
S077-U-d             CATCATGCCC CTTATGACCA GGGCTACACA CGTGCTACAA TGGCGTAAAC AAAGGGAGGC GAACCCGCGA GGGTGGGCAA
VPI12708-a           CATCATGCCC CTTATGACCA GGGCTACACA CGTGCTACAA TGGCGTAAAC AAAGGGAGGC GAACCCGCGA GGGTGGGCAA
VPI12708-b           CATCATGCCC CTTATGACCA GGGCTACACA CGTGCTACAA TGGCGTAAAC AAAGGGAGGC GAACCCGCGA GGGTGGGCAA
VPI12708-c           CATCATGCCC CTTATGACCA GGGCTACACA CGTGCTACAA TGGCGTAAAC AAAGGGAGGC GAACCCGCGA GGGTGGGCAA
VPI12708-d           CATCATGCCC CTTATGACCA GGGCTACACA CGTGCTACAA TGGCGTAAAC AAAGGGAGGC GAACCCGCGA GGGTGGGCAA
DFI.1.130            CATCATGCCC CTTATGACCA GGGCTACACA CGTGCTACAA TGGCGTAAAC AAAGGGAGGC GAACCCGCGA GGGTGGGCAA
DFI.1.161            ---------- ---------- ---------- ---------- ---------- ---------- ---------- ----------
DFI.1.162            CATCATGCCC CTTATGACCA GGGCTACACA CGTGCTACAA TGGCGTAAAC AAAGGGAGGC GAACCCGCGA GGGTGGGCAA
DFI.1.217            CATCATGCCC CTTATGACCA GGGCTACACA CGTGCTACAA TGGCGTAAAC AAAGGGAGGC GAACCCGCGA GGGTGGGCAA
DFI.1.234            CATCATGCCC CTTATGACCA GGGCTACACA CGTGCTACAA TGGCGTAAAC AAAGGGAGGC GAACCCGCGA GGGTGGGCAA
DFI.1.60             CATCATGCCC CTTATGACCA GGGCTACACA CGTGCTACAA TGGCGTAAAC AAAGGGAGGC GAACCCGCGA GGGTGGGCAA
DFI.4.63             CATCATGCCC CTTATGACCA GGGCTACACA CGTGCTACAA TGGCGTAAAC AAAGGGAGGC GAACCCGCGA GGGTGGGCAA
JCM10419-a           CATCATGCCC CTTATGACCA GGGCTACACA CGTGCTACAA TGGCGTAAAC AAAGGGAGGC GAACCCGCGA GGGTGGGCAA
JCM10419-b           CATCATGCCC CTTATGACCA GGGCTACACA CGTGCTACAA TGGCGTAAAC AAAGGGAGGC GAACCCGCGA GGGTGGGCAA
JCM10419-c           CATCATGCCC CTTATGACCA GGGCTACACA CGTGCTACAA TGGCGTAAAC AAAGGGAGGC GAACCCGCGA GGGTGGGCAA
JCM10419-d           CATCATGCCC CTTATGACCA GGGCTACACA CGTGCTACAA TGGCGTAAAC AAAGGGAGGC GAACCCGCGA GGGTGGGCAA
JCM10420-a           CATCATGCCC CTTATGACCA GGGCTACACA CGTGCTACAA TGGCGTAAAC AAAGGGAGGC GAACCCGCGA GGGTGGGCAA
JCM10420-b           CATCATGCCC CTTATGACCA GGGCTACACA CGTGCTACAA TGGCGTAAAC AAAGGGAGGC GAACCCGCGA GGGTGGGCAA
JCM10420-c           CATCATGCCC CTTATGACCA GGGCTACACA CGTGCTACAA TGGCGTAAAC AAAGGGAGGC GAACCCGCGA GGGTGGGCAA
JCM10420-d           CATCATGCCC CTTATGACCA GGGCTACACA CGTGCTACAA TGGCGTAAAC AAAGGGAGGC GAACCCGCGA GGGTGGGCAA
JCM10423-a           CATCATGCCC CTTATGACCA GGGCTACACA CGTGCTACAA TGGCGTAAAC AAAGGGAGGC GAACCCGCGA GGGTGGGCAA
JCM10423-b           CATCATGCCC CTTATGACCA GGGCTACACA CGTGCTACAA TGGCGTAAAC AAAGGGAGGC GAACCCGCGA GGGTGGGCAA
JCM10423-c           CATCATGCCC CTTATGACCA GGGCTACACA CGTGCTACAA TGGCGTAAAC AAAGGGAGGC GAACCCGCGA GGGTGGGCAA
JCM10423-d           CATCATGCCC CTTATGACCA GGGCTACACA CGTGCTACAA TGGCGTAAAC AAAGGGAGGC GAACCCGCGA GGGTGGGCAA
MO32-a               CATCATGCCC CTTATGACCA GGGCTACACA CGTGCTACAA TGGCGTAAAC AAAGGGAGGC GAACCCGCGA GGGTGGGCAA
MO32-b               CATCATGCCC CTTATGACCA GGGCTACACA CGTGCTACAA TGGCGTAAAC AAAGGGAGGC GAACCCGCGA GGGTGGGCAA
MO32-c               CATCATGCCC CTTATGACCA GGGCTACACA CGTGCTACAA TGGCGTAAAC AAAGGGAGGC GAACCCGCGA GGGTGGGCAA
MO32-d               CATCATGCCC CTTATGACCA GGGCTACACA CGTGCTACAA TGGCGTAAAC AAAGGGAGGC GAACCCGCGA GGGTGGGCAA
MSK.5.24             CATCATGCCC CTTATGACCA GGGCTACACA CGTGCTACAA TGGCGTAAAC AAAGGGAGGC GAACCCGCGA GGGTGGGCAA
NB2A-7-D5            CATCATGCCC CTTATGACCA GGGCTACACA CGTGCTACAA TGGCGTAAAC AAAGGGAGGC GAACCCGCGA GGGTGGGCAA
NTI82-a              CATCATGCCC CTTATGACCA GGGCTACACA CGTGCTACAA TGGCGTAAAC AAAGGGAGGC GAACCCGCGA GGGTGGGCAA
NTI82-b              CATCATGCCC CTTATGACCA GGGCTACACA CGTGCTACAA TGGCGTAAAC AAAGGGAGGC GAACCCGCGA GGGTGGGCAA
NTI82-c              CATCATGCCC CTTATGACCA GGGCTACACA CGTGCTACAA TGGCGTAAAC AAAGGGAGGC GAACCCGCGA GGGTGGGCAA
NTI82-d              CATCATGCCC CTTATGACCA GGGCTACACA CGTGCTACAA TGGCGTAAAC AAAGGGAGGC GAACCCGCGA GGGTGGGCAA
Q4-a                 CATCATGCCC CTTATGACCA GGGCTACACA CGTGCTACAA TGGCGTAAAC AAAGGGAGGC GAACCCGCGA GGGTGGGCAA
Q4-b                 CATCATGCCC CTTATGACCA GGGCTACACA CGTGCTACAA TGGCGTAAAC AAAGGGAGGC GAACCCGCGA GGGTGGGCAA
Q4-c                 CATCATGCCC CTTATGACCA GGGCTACACA CGTGCTACAA TGGCGTAAAC AAAGGGAGGC GAACCCGCGA GGGTGGGCAA
Q4-d                 CATCATGCCC CTTATGACCA GGGCTACACA CGTGCTACAA TGGCGTAAAC AAAGGGAGGC GAACCCGCGA GGGTGGGCAA
S076-a               CATCATGCCC CTTATGACCA GGGCTACACA CGTGCTACAA TGGCGTAAAC AAAGGGAGGC GAACCCGCGA GGGTGGGCAA
S076-b               CATCATGCCC CTTATGACCA GGGCTACACA CGTGCTACAA TGGCGTAAAC AAAGGGAGGC GAACCCGCGA GGGTGGGCAA
S076-c               CATCATGCCC CTTATGACCA GGGCTACACA CGTGCTACAA TGGCGTAAAC AAAGGGAGGC GAACCCGCGA GGGTGGGCAA
S076-d               CATCATGCCC CTTATGACCA GGGCTACACA CGTGCTACAA TGGCGTAAAC AAAGGGAGGC GAACCCGCGA GGGTGGGCAA
SL.1.22              CATCATGCCC CTTATGACCA GGGCTACACA CGTGCTACAA TGGCGTAAAC AAAGGGAGGC GAACCCGCGA GGGTGGGCAA
VE202-05             CATCATGCCC CTTATGACCA GGGCTACACA CGTGCTACAA TGGCGTAAAC AAAGGGAGGC GAACCCGCGA GGGTGGGCAA
C.hylemonae          CATCATGCCC CTTATGACCA GGGCTACACA CGTGCTACAA TGGCGTAAAC AAAGGGAAGC GACCCTGTGA AGGCAAGCAA
```

```
                      1281
35704-a               ATCCCAAAAA TAACGTCTCA GTTCGGATTG TAGTCTGCAA CTCGACTACA TGAAGTTGGA ATCGCTAGTA ATCGCGAATC
35704-b               ATCCCAAAAA TAACGTCTCA GTTCGGATTG TAGTCTGCAA CTCGACTACA TGAAGCTGGA ATCGCTAGTA ATCGCGAATC
35704-c               ATCCCAAAAA TAACGTCTCA GTTCGGATTG TAGTCTGCAA CTCGACTACA TGAAGCTGGA ATCGCTAGTA ATCGCGAATC
35704-d               ATCCCAAAAA TAACGTCTCA GTTCGGATTG TAGTCTGCAA CTCGACTACA TGAAGCTGGA ATCGCTAGTA ATCGCGAATC
AM05-22               ATCCCAAAAA TAACGTCTCA GTTCGGATTG TAGTCTGCAA CTCGACTACA TGAAGCTGGA ATCGCTAGTA ATCGCGAATC
AM07-30               ATCCCAAAAA TAACGTCTCA GTTCGGATTG TAGTCTGCAA CTCGACTACA TGAAGCTGGA ATCGCTAGTA ATCGCGAATC
BL389WT3D-a           ATCCCAAAAA TAACGTCTCA GTTCGGATTG TAGTCTGCAA CTCGACTACA TGAAGCTGGA ATCGCTAGTA ATCGCGAATC
BL389WT3D-b           ATCCCAAAAA TAACGTCTCA GTTCGGATTG TAGTCTGCAA CTCGACTACA TGAAGCTGGA ATCGCTAGTA ATCGCGAATC
BL389WT3D-c           ATCCCAAAAA TAACGTCTCA GTTCGGATTG TAGTCTGCAA CTCGACTACA TGAAGCTGGA ATCGCTAGTA ATCGCGAATC
BL389WT3D-d           ATCCCAAAAA TAACGTCTCA GTTCGGATTG TAGTCTGCAA CTCGACTACA TGAAGCTGGA ATCGCTAGTA ATCGCGAATC
CE91-St59-a           ATCCCAAAAA TAACGTCTCA GTTCGGATTG TAGTCTGCAA CTCGACTACA TGAAGCTGGA ATCGCTAGTA ATCGCGAATC
CE91-St59-b           ATCCCAAAAA TAACGTCTCA GTTCGGATTG TAGTCTGCAA CTCGACTACA TGAAGCTGGA ATCGCTAGTA ATCGCGAATC
CE91-St59-c           ATCCCAAAAA TAACGTCTCA GTTCGGATTG TAGTCTGCAA CTCGACTACA TGAAGCTGGA ATCGCTAGTA ATCGCGAATC
CE91-St60-a           ATCCCAAAAA TAACGTCTCA GTTCGGATTG TAGTCTGCAA CTCGACTACA TGAAGCTGGA ATCGCTAGTA ATCGCGAATC
CE91-St60-b           ATCCCAAAAA TAACGTCTCA GTTCGGATTG TAGTCTGCAA CTCGACTACA TGAAGCTGGA ATCGCTAGTA ATCGCGAATC
CE91-St60-c           ATCCCAAAAA TAACGTCTCA GTTCGGATTG TAGTCTGCAA CTCGACTACA TGAAGCTGGA ATCGCTAGTA ATCGCGAATC
FDAARGOS_1227-a       ATCCCAAAAA TAACGTCTCA GTTCGGATTG TAGTCTGCAA CTCGACTACA TGAAGCTGGA ATCGCTAGTA ATCGCGAATC
FDAARGOS_1227-b       ATCCCAAAAA TAACGTCTCA GTTCGGATTG TAGTCTGCAA CTCGACTACA TGAAGCTGGA ATCGCTAGTA ATCGCGAATC
FDAARGOS_1227-c       ATCCCAAAAA TAACGTCTCA GTTCGGATTG TAGTCTGCAA CTCGACTACA TGAAGTTGGA ATCGCTAGTA ATCGCGAATC
FDAARGOS_1227-d       ATCCCAAAAA TAACGTCTCA GTTCGGATTG TAGTCTGCAA CTCGACTACA TGAAGCTGGA ATCGCTAGTA ATCGCGAATC
G10-a                 ATCCCAAAAA TAACGTCTCA GTTCGGATTG TAGTCTGCAA CTCGACTACA TGAAGCTGGA ATCGCTAGTA ATCGCGAATC
G10-b                 ATCCCAAAAA TAACGTCTCA GTTCGGATTG TAGTCTGCAA CTCGACTACA TGAAGCTGGA ATCGCTAGTA ATCGCGAATC
G10-c                 ATCCCAAAAA TAACGTCTCA GTTCGGATTG TAGTCTGCAA CTCGACTACA TGAAGCTGGA ATCGCTAGTA ATCGCGAATC
G10-d                 ATCCCAAAAA TAACGTCTCA GTTCGGATTG TAGTCTGCAA CTCGACTACA TGAAGCTGGA ATCGCTAGTA ATCGCGAATC
GGCC_0168             ATCCCAAAAA TAACGTCTCA GTTCGGATTG TAGTCTGCAA CTCGACTACA TGAAGCTGGA ATCGCTAGTA ATCGCGAATC
I10-a                 ATCCCAAAAA TAACGTCTCA GTTCGGATTG TAGTCTGCAA CTCGACTACA TGAAGCTGGA ATCGCTAGTA ATCGCGAATC
I10-b                 ATCCCAAAAA TAACGTCTCA GTTCGGATTG TAGTCTGCAA CTCGACTACA TGAAGCTGGA ATCGCTAGTA ATCGCGAATC
I10-c                 ATCCCAAAAA TAACGTCTCA GTTCGGATTG TAGTCTGCAA CTCGACTACA TGAAGCTGGA ATCGCTAGTA ATCGCGAATC
I10-d                 ATCCCAAAAA TAACGTCTCA GTTCGGATTG TAGTCTGCAA CTCGACTACA TGAAGCTGGA ATCGCTAGTA ATCGCGAATC
JCM10422-a            ATCCCAAAAA TAACGTCTCA GTTCGGATTG TAGTCTGCAA CTCGACTACA TGAAGCTGGA ATCGCTAGTA ATCGCGAATC
JCM10422-b            ATCCCAAAAA TAACGTCTCA GTTCGGATTG TAGTCTGCAA CTCGACTACA TGAAGCTGGA ATCGCTAGTA ATCGCGAATC
JCM10422-c            ATCCCAAAAA TAACGTCTCA GTTCGGATTG TAGTCTGCAA CTCGACTACA TGAAGCTGGA ATCGCTAGTA ATCGCGAATC
JCM10422-d            ATCCCAAAAA TAACGTCTCA GTTCGGATTG TAGTCTGCAA CTCGACTACA TGAAGCTGGA ATCGCTAGTA ATCGCGAATC
MGYG-HGUT-01303-a     ATCCCAAAAA TAACGTCTCA GTTCGGATTG TAGTCTGCAA CTCGACTACA TGAAGTTGGA ATCGCTAGTA ATCGCGAATC
MGYG-HGUT-01303-b     ATCCCAAAAA TAACGTCTCA GTTCGGATTG TAGTCTGCAA CTCGACTACA TGAAGCTGGA ATCGCTAGTA ATCGCGAATC
MSK.1.16              ATCCCAAAAA TAACGTCTCA GTTCGGATTG TAGTCTGCAA CTCGACTACA TGAAGCTGGA ATCGCTAGTA ATCGCGAATC
MSK.1.26              ---------- ---------- ---------- ---------- ---------- ---------- ---------- ----------
S077-U-a              ATCCCAAAAA TAACGTCTCA GTTCGGATTG TAGTCTGCAA CTCGACTACA TGAAGCTGGA ATCGCTAGTA ATCGCGAATC
S077-U-b              ATCCCAAAAA TAACGTCTCA GTTCGGATTG TAGTCTGCAA CTCGACTACA TGAAGCTGGA ATCGCTAGTA ATCGCGAATC
S077-U-c              ATCCCAAAAA TAACGTCTCA GTTCGGATTG TAGTCTGCAA CTCGACTACA TGAAGCTGGA ATCGCTAGTA ATCGCGAATC
S077-U-d              ATCCCAAAAA TAACGTCTCA GTTCGGATTG TAGTCTGCAA CTCGACTACA TGAAGCTGGA ATCGCTAGTA ATCGCGAATC
VPI12708-a            ATCCCAAAAA TAACGTCTCA GTTCGGATTG TAGTCTGCAA CTCGACTACA TGAAGCTGGA ATCGCTAGTA ATCGCGAATC
VPI12708-b            ATCCCAAAAA TAACGTCTCA GTTCGGATTG TAGTCTGCAA CTCGACTACA TGAAGCTGGA ATCGCTAGTA ATCGCGAATC
VPI12708-c            ATCCCAAAAA TAACGTCTCA GTTCGGATTG TAGTCTGCAA CTCGACTACA TGAAGCTGGA ATCGCTAGTA ATCGCGAATC
VPI12708-d            ATCCCAAAAA TAACGTCTCA GTTCGGATTG TAGTCTGCAA CTCGACTACA TGAAGCTGGA ATCGCTAGTA ATCGCGAATC
DFI.1.130             ATCCCAAAAA TAACGTCTCA GTTCGGATTG TAGTCTGCAA CTCGACTACA TGAAGCTGGA ATCGCTAGTA ATCGCGAATC
DFI.1.161             ---------- ---------- ---------- ---------- ---------- ---------- ---------- ----------
DFI.1.162             ATCCCAAAAA TAACGTCTCA GTTCGGATTG TAGTCTGCAA CTCGACTACA TGAAGCTGGA ATCGCTAGTA ATCGCGAATC
DFI.1.217             ATCCCAAAAA TAACGTCTCA GTTCGGATTG TAGTCTGCAA CTCGACTACA TGAAGCTGGA ATCGCTAGTA ATCGCGAATC
DFI.1.234             ATCCCAAAAA TAACGTCTCA GTTCGGATTG TAGTCTGCAA CTCGACTACA TGAAGCTGGA ATCGCTAGTA ATCGCGAATC
DFI.1.60              ATCCCAAAAA TAACGTCTCA GTTCGGATTG TAGTCTGCAA CTCGACTACA TGAAGCTGGA ATCGCTAGTA ATCGCGAATC
DFI.4.63              ATCCCAAAAA TAACGTCTCA GTTCGGATTG TAGTCTGCAA CTCGACTACA TGAAGCTGGA ATCGCTAGTA ATCGCGAATC
JCM10419-a            ATCCCAAAAA TAACGTCTCA GTTCGGATTG TAGTCTGCAA CTCGACTACA TGAAGCTGGA ATCGCTAGTA ATCGCGAATC
JCM10419-b            ATCCCAAAAA TAACGTCTCA GTTCGGATTG TAGTCTGCAA CTCGACTACA TGAAGCTGGA ATCGCTAGTA ATCGCGAATC
JCM10419-c            ATCCCAAAAA TAACGTCTCA GTTCGGATTG TAGTCTGCAA CTCGACTACA TGAAGCTGGA ATCGCTAGTA ATCGCGAATC
JCM10419-d            ATCCCAAAAA TAACGTCTCA GTTCGGATTG TAGTCTGCAA CTCGACTACA TGAAGCTGGA ATCGCTAGTA ATCGCGAATC
JCM10420-a            ATCCCAAAAA TAACGTCTCA GTTCGGATTG TAGTCTGCAA CTCGACTACA TGAAGCTGGA ATCGCTAGTA ATCGCGAATC
JCM10420-b            ATCCCAAAAA TAACGTCTCA GTTCGGATTG TAGTCTGCAA CTCGACTACA TGAAGCTGGA ATCGCTAGTA ATCGCGAATC
JCM10420-c            ATCCCAAAAA TAACGTCTCA GTTCGGATTG TAGTCTGCAA CTCGACTACA TGAAGCTGGA ATCGCTAGTA ATCGCGAATC
JCM10420-d            ATCCCAAAAA TAACGTCTCA GTTCGGATTG TAGTCTGCAA CTCGACTACA TGAAGCTGGA ATCGCTAGTA ATCGCGAATC
JCM10423-a            ATCCCAAAAA TAACGTCTCA GTTCGGATTG TAGTCTGCAA CTCGACTACA TGAAGCTGGA ATCGCTAGTA ATCGCGAATC
JCM10423-b            ATCCCAAAAA TAACGTCTCA GTTCGGATTG TAGTCTGCAA CTCGACTACA TGAAGCTGGA ATCGCTAGTA ATCGCGAATC
JCM10423-c            ATCCCAAAAA TAACGTCTCA GTTCGGATTG TAGTCTGCAA CTCGACTACA TGAAGCTGGA ATCGCTAGTA ATCGCGAATC
JCM10423-d            ATCCCAAAAA TAACGTCTCA GTTCGGATTG TAGTCTGCAA CTCGACTACA TGAAGCTGGA ATCGCTAGTA ATCGCGAATC
MO32-a                ATCCCAAAAA TAACGTCTCA GTTCGGATTG TAGTCTGCAA CTCGACTACA TGAAGCTGGA ATCGCTAGTA ATCGCGAATC
MO32-b                ATCCCAAAAA TAACGTCTCA GTTCGGATTG TAGTCTGCAA CTCGACTACA TGAAGCTGGA ATCGCTAGTA ATCGCGAATC
MO32-c                ATCCCAAAAA TAACGTCTCA GTTCGGATTG TAGTCTGCAA CTCGACTACA TGAAGCTGGA ATCGCTAGTA ATCGCGAATC
MO32-d                ATCCCAAAAA TAACGTCTCA GTTCGGATTG TAGTCTGCAA CTCGACTACA TGAAGCTGGA ATCGCTAGTA ATCGCGAATC
MSK.5.24              ATCCCAAAAA TAACGTCTCA GTTCGGATTG TAGTCTGCAA CTCGACTACA TGAAGCTGGA ATCGCTAGTA ATCGCGAATC
NB2A-7-D5             ATCCCAAAAA TAACGTCTCA GTTCGGATTG TAGTCTGCAA CTCGACTACA TGAAGCTGGA ATCGCTAGTA ATCGCGAATC
NTI82-a               ATCCCAAAAA TAACGTCTCA GTTCGGATTG TAGTCTGCAA CTCGACTACA TGAAGCTGGA ATCGCTAGTA ATCGCGAATC
NTI82-b               ATCCCAAAAA TAACGTCTCA GTTCGGATTG TAGTCTGCAA CTCGACTACA TGAAGCTGGA ATCGCTAGTA ATCGCGAATC
NTI82-c               ATCCCAAAAA TAACGTCTCA GTTCGGATTG TAGTCTGCAA CTCGACTACA TGAAGCTGGA ATCGCTAGTA ATCGCGAATC
NTI82-d               ATCCCAAAAA TAACGTCTCA GTTCGGATTG TAGTCTGCAA CTCGACTACA TGAAGCTGGA ATCGCTAGTA ATCGCGAATC
Q4-a                  ATCCCAAAAA TAACGTCTCA GTTCGGATTG TAGTCTGCAA CTCGACTACA TGAAGCTGGA ATCGCTAGTA ATCGCGAATC
Q4-b                  ATCCCAAAAA TAACGTCTCA GTTCGGATTG TAGTCTGCAA CTCGACTACA TGAAGCTGGA ATCGCTAGTA ATCGCGAATC
Q4-c                  ATCCCAAAAA TAACGTCTCA GTTCGGATTG TAGTCTGCAA CTCGACTACA TGAAGCTGGA ATCGCTAGTA ATCGCGAATC
Q4-d                  ATCCCAAAAA TAACGTCTCA GTTCGGATTG TAGTCTGCAA CTCGACTACA TGAAGCTGGA ATCGCTAGTA ATCGCGAATC
S076-a                ATCCCAAAAA TAACGTCTCA GTTCGGATTG TAGTCTGCAA CTCGACTACA TGAAGCTGGA ATCGCTAGTA ATCGCGAATC
S076-b                ATCCCAAAAA TAACGTCTCA GTTCGGATTG TAGTCTGCAA CTCGACTACA TGAAGCTGGA ATCGCTAGTA ATCGCGAATC
S076-c                ATCCCAAAAA TAACGTCTCA GTTCGGATTG TAGTCTGCAA CTCGACTACA TGAAGCTGGA ATCGCTAGTA ATCGCGAATC
S076-d                ATCCCAAAAA TAACGTCTCA GTTCGGATTG TAGTCTGCAA CTCGACTACA TGAAGCTGGA ATCGCTAGTA ATCGCGAATC
SL.1.22               ATCCCAAAAA TAACGTCTCA GTTCGGATTG TAGTCTGCAA CTCGACTACA TGAAGCTGGA ATCGCTAGTA ATCGCGAATC
VE202-05              ATCCCAAAAA TAACGTCTCA GTTCGGATTG TAGTCTGCAA CTCGACTACA TGAAGCTGGA ATCGCTAGTA ATCGCGAATC
C.hylemonae           ATCCCAAAAA TAACGTCTCA GTTCGGATTG TAGTCTGCAA CTCGACTACA TGAAGCTGGA ATCGCTAGTA ATCGCGAATC
```

```
                      1361
35704-a               AGAATGTCGC GGTGAATACG TTCCCGGGTC TTGTACACAC CGCCCGTCAC ACCATGGGAG TCAGTAACGC CCGAAGCCGG
35704-b               AGAATGTCGC GGTGAATACG TTCCCGGGTC TTGTACACAC CGCCCGTCAC ACCATGGGAG TCAGTAACGC CCGAAGCCGG
35704-c               AGAATGTCGC GGTGAATACG TTCCCGGGTC TTGTACACAC CGCCCGTCAC ACCATGGGAG TCAGTAACGC CCGAAGCCGG
35704-d               AGAATGTCGC GGTGAATACG TTCCCGGGTC TTGTACACAC CGCCCGTCAC ACCATGGGAG TCAGTAACGC CCGAAGCCGG
AM05-22               AGAATGTCGC GGTGAATACG TTCCCGGGTC TTGTACACAC CGCCCGTCAC ACCATGGGAG TCAGTAACGC CCGAAGCCGG
AM07-30               AGAATGTCGC GGTGAATACG TTCCCGGGTC TTGTACACAC CGCCCGTCAC ACCATGGGAG TCAGTAACGC CCGAAGCCGG
BL389WT3D-a           AGAATGTCGC GGTGAATACG TTCCCGGGTC TTGTACACAC CGCCCGTCAC ACCATGGGAG TCAGTAACGC CCGAAGCCGG
BL389WT3D-b           AGAATGTCGC GGTGAATACG TTCCCGGGTC TTGTACACAC CGCCCGTCAC ACCATGGGAG TCAGTAACGC CCGAAGCCGG
BL389WT3D-c           AGAATGTCGC GGTGAATACG TTCCCGGGTC TTGTACACAC CGCCCGTCAC ACCATGGGAG TCAGTAACGC CCGAAGCCGG
BL389WT3D-d           AGAATGTCGC GGTGAATACG TTCCCGGGTC TTGTACACAC CGCCCGTCAC ACCATGGGAG TCAGTAACGC CCGAAGCCGG
CE91-St59-a           AGAATGTCGC GGTGAATACG TTCCCGGGTC TTGTACACAC CGCCCGTCAC ACCATGGGAG TCAGTAACGC CCGAAGCCGG
CE91-St59-b           AGAATGTCGC GGTGAATACG TTCCCGGGTC TTGTACACAC CGCCCGTCAC ACCATGGGAG TCAGTAACGC CCGAAGCCGG
CE91-St59-c           AGAATGTCGC GGTGAATACG TTCCCGGGTC TTGTACACAC CGCCCGTCAC ACCATGGGAG TCAGTAACGC CCGAAGCCGG
CE91-St60-a           AGAATGTCGC GGTGAATACG TTCCCGGGTC TTGTACACAC CGCCCGTCAC ACCATGGGAG TCAGTAACGC CCGAAGCCGG
CE91-St60-b           AGAATGTCGC GGTGAATACG TTCCCGGGTC TTGTACACAC CGCCCGTCAC ACCATGGGAG TCAGTAACGC CCGAAGCCGG
CE91-St60-c           AGAATGTCGC GGTGAATACG TTCCCGGGTC TTGTACACAC CGCCCGTCAC ACCATGGGAG TCAGTAACGC CCGAAGCCGG
FDAARGOS_1227-a       AGAATGTCGC GGTGAATACG TTCCCGGGTC TTGTACACAC CGCCCGTCAC ACCATGGGAG TCAGTAACGC CCGAAGCCGG
FDAARGOS_1227-b       AGAATGTCGC GGTGAATACG TTCCCGGGTC TTGTACACAC CGCCCGTCAC ACCATGGGAG TCAGTAACGC CCGAAGCCGG
FDAARGOS_1227-c       AGAATGTCGC GGTGAATACG TTCCCGGGTC TTGTACACAC CGCCCGTCAC ACCATGGGAG TCAGTAACGC CCGAAGCCGG
FDAARGOS_1227-d       AGAATGTCGC GGTGAATACG TTCCCGGGTC TTGTACACAC CGCCCGTCAC ACCATGGGAG TCAGTAACGC CCGAAGCCGG
G10-a                 AGAATGTCGC GGTGAATACG TTCCCGGGTC TTGTACACAC CGCCCGTCAC ACCATGGGAG TCAGTAACGC CCGAAGCCGG
G10-b                 AGAATGTCGC GGTGAATACG TTCCCGGGTC TTGTACACAC CGCCCGTCAC ACCATGGGAG TCAGTAACGC CCGAAGCCGG
G10-c                 AGAATGTCGC GGTGAATACG TTCCCGGGTC TTGTACACAC CGCCCGTCAC ACCATGGGAG TCAGTAACGC CCGAAGCCGG
G10-d                 AGAATGTCGC GGTGAATACG TTCCCGGGTC TTGTACACAC CGCCCGTCAC ACCATGGGAG TCAGTAACGC CCGAAGCCGG
GGCC_0168             AGAATGTCGC GGTGAATACG TTCCCGGGTC TTGTACACAC CGCCCGTCAC ACCATGGGAG TCAGTAACGC CCGAAGCCGG
I10-a                 AGAATGTCGC GGTGAATACG TTCCCGGGTC TTGTACACAC CGCCCGTCAC ACCATGGGAG TCAGTAACGC CCGAAGCCGG
I10-b                 AGAATGTCGC GGTGAATACG TTCCCGGGTC TTGTACACAC CGCCCGTCAC ACCATGGGAG TCAGTAACGC CCGAAGCCGG
I10-c                 AGAATGTCGC GGTGAATACG TTCCCGGGTC TTGTACACAC CGCCCGTCAC ACCATGGGAG TCAGTAACGC CCGAAGCCGG
I10-d                 AGAATGTCGC GGTGAATACG TTCCCGGGTC TTGTACACAC CGCCCGTCAC ACCATGGGAG TCAGTAACGC CCGAAGCCGG
JCM10422-a            AGAATGTCGC GGTGAATACG TTCCCGGGTC TTGTACACAC CGCCCGTCAC ACCATGGGAG TCAGTAACGC CCGAAGCCGG
JCM10422-b            AGAATGTCGC GGTGAATACG TTCCCGGGTC TTGTACACAC CGCCCGTCAC ACCATGGGAG TCAGTAACGC CCGAAGCCGG
JCM10422-c            AGAATGTCGC GGTGAATACG TTCCCGGGTC TTGTACACAC CGCCCGTCAC ACCATGGGAG TCAGTAACGC CCGAAGCCGG
JCM10422-d            AGAATGTCGC GGTGAATACG TTCCCGGGTC TTGTACACAC CGCCCGTCAC ACCATGGGAG TCAGTAACGC CCGAAGCCGG
MGYG-HGUT-01303-a     AGAATGTCGC GGTGAATACG TTCCCGGGTC TTGTACACAC CGCCCGTCAC ACCATGGGAG TCAGTAACGC CCGAAGCCGG
MGYG-HGUT-01303-b     AGAATGTCGC GGTGAATACG TTCCCGGGTC TTGTACACAC CGCCCGTCAC ACCATGGGAG TCAGTAACGC CCGAAGCCGG
MSK.1.16              AGAATGTCGC GGTGAATACG TTCCCGGGTC TTGTACACAC CGCCCGTCAC ACCATGGGAG TCAGTAACGC CCGAAGCCGG
MSK.1.26              ---------- ---------- ---------- ---------- ---------- ---------- ---------- ----------
S077-U-a              AGAATGTCGC GGTGAATACG TTCCCGGGTC TTGTACACAC CGCCCGTCAC ACCATGGGAG TCAGTAACGC CCGAAGCCGG
S077-U-b              AGAATGTCGC GGTGAATACG TTCCCGGGTC TTGTACACAC CGCCCGTCAC ACCATGGGAG TCAGTAACGC CCGAAGCCGG
S077-U-c              AGAATGTCGC GGTGAATACG TTCCCGGGTC TTGTACACAC CGCCCGTCAC ACCATGGGAG TCAGTAACGC CCGAAGCCGG
S077-U-d              AGAATGTCGC GGTGAATACG TTCCCGGGTC TTGTACACAC CGCCCGTCAC ACCATGGGAG TCAGTAACGC CCGAAGCCGG
VPI12708-a            AGAATGTCGC GGTGAATACG TTCCCGGGTC TTGTACACAC CGCCCGTCAC ACCATGGGAG TCAGTAACGC CCGAAGCCGG
VPI12708-b            AGAATGTCGC GGTGAATACG TTCCCGGGTC TTGTACACAC CGCCCGTCAC ACCATGGGAG TCAGTAACGC CCGAAGCCGG
VPI12708-c            AGAATGTCGC GGTGAATACG TTCCCGGGTC TTGTACACAC CGCCCGTCAC ACCATGGGAG TCAGTAACGC CCGAAGCCGG
VPI12708-d            AGAATGTCGC GGTGAATACG TTCCCGGGTC TTGTACACAC CGCCCGTCAC ACCATGGGAG TCAGTAACGC CCGAAGCCGG
DFI.1.130             AGAATGTCGC GGTGAATACG TTCCCGGGTC TTGTACACAC CGCCCGTCAC ACCATGGGAG TCAGTAACGC CCGAAGCCGG
DFI.1.161             ---------- ---------- ---------- ---------- ---------- ---------- ---------- ----------
DFI.1.162             AGAATGTCGC GGTGAATACG TTCCCGGGTC TTGTACACAC CGCCCGTCAC ACCATGGGAG TCAGTAACGC CCGAAGCCGG
DFI.1.217             AGAATGTCGC GGTGAATACG TTCCCGGGTC TTGTACACAC CGCCCGTCAC ACCATGGGAG TCAGTAACGC CCGAAGCCGG
DFI.1.234             AGAATGTCGC GGTGAATACG TTCCCGGGTC TTGTACACAC CGCCCGTCAC ACCATGGGAG TCAGTAACGC CCGAAGCCGG
DFI.1.60              AGAATGTCGC GGTGAATACG TTCCCGGGTC TTGTACACAC CGCCCGTCAC ACCATGGGAG TCAGTAACGC CCGAAGCCGG
DFI.4.63              AGAATGTCGC GGTGAATACG TTCCCGGGTC TTGTACACAC CGCCCGTCAC ACCATGGGAG TCAGTAACGC CCGAAGCCGG
JCM10419-a            AGAATGTCGC GGTGAATACG TTCCCGGGTC TTGTACACAC CGCCCGTCAC ACCATGGGAG TCAGTAACGC CCGAAGCCGG
JCM10419-b            AGAATGTCGC GGTGAATACG TTCCCGGGTC TTGTACACAC CGCCCGTCAC ACCATGGGAG TCAGTAACGC CCGAAGCCGG
JCM10419-c            AGAATGTCGC GGTGAATACG TTCCCGGGTC TTGTACACAC CGCCCGTCAC ACCATGGGAG TCAGTAACGC CCGAAGCCGG
JCM10419-d            AGAATGTCGC GGTGAATACG TTCCCGGGTC TTGTACACAC CGCCCGTCAC ACCATGGGAG TCAGTAACGC CCGAAGCCGG
JCM10420-a            AGAATGTCGC GGTGAATACG TTCCCGGGTC TTGTACACAC CGCCCGTCAC ACCATGGGAG TCAGTAACGC CCGAAGCCGG
JCM10420-b            AGAATGTCGC GGTGAATACG TTCCCGGGTC TTGTACACAC CGCCCGTCAC ACCATGGGAG TCAGTAACGC CCGAAGCCGG
JCM10420-c            AGAATGTCGC GGTGAATACG TTCCCGGGTC TTGTACACAC CGCCCGTCAC ACCATGGGAG TCAGTAACGC CCGAAGCCGG
JCM10420-d            AGAATGTCGC GGTGAATACG TTCCCGGGTC TTGTACACAC CGCCCGTCAC ACCATGGGAG TCAGTAACGC CCGAAGCCGG
JCM10423-a            AGAATGTCGC GGTGAATACG TTCCCGGGTC TTGTACACAC CGCCCGTCAC ACCATGGGAG TCAGTAACGC CCGAAGCCGG
JCM10423-b            AGAATGTCGC GGTGAATACG TTCCCGGGTC TTGTACACAC CGCCCGTCAC ACCATGGGAG TCAGTAACGC CCGAAGCCGG
JCM10423-c            AGAATGTCGC GGTGAATACG TTCCCGGGTC TTGTACACAC CGCCCGTCAC ACCATGGGAG TCAGTAACGC CCGAAGCCGG
JCM10423-d            AGAATGTCGC GGTGAATACG TTCCCGGGTC TTGTACACAC CGCCCGTCAC ACCATGGGAG TCAGTAACGC CCGAAGCCGG
MO32-a                AGAATGTCGC GGTGAATACG TTCCCGGGTC TTGTACACAC CGCCCGTCAC ACCATGGGAG TCAGTAACGC CCGAAGCCGG
MO32-b                AGAATGTCGC GGTGAATACG TTCCCGGGTC TTGTACACAC CGCCCGTCAC ACCATGGGAG TCAGTAACGC CCGAAGCCGG
MO32-c                AGAATGTCGC GGTGAATACG TTCCCGGGTC TTGTACACAC CGCCCGTCAC ACCATGGGAG TCAGTAACGC CCGAAGCCGG
MO32-d                AGAATGTCGC GGTGAATACG TTCCCGGGTC TTGTACACAC CGCCCGTCAC ACCATGGGAG TCAGTAACGC CCGAAGCCGG
MSK.5.24              AGAATGTCGC GGTGAATACG TTCCCGGGTC TTGTACACAC CGCCCGTCAC ACCATGGGAG TCAGTAACGC CCGAAGCCGG
NB2A-7-D5             AGAATGTCGC GGTGAATACG TTCCCGGGTC TTGTACACAC CGCCCGTCAC ACCATGGGAG TCAGTAACGC CCGAAGCCGG
NTI82-a               AGAATGTCGC GGTGAATACG TTCCCGGGTC TTGTACACAC CGCCCGTCAC ACCATGGGAG TCAGTAACGC CCGAAGCCGG
NTI82-b               AGAATGTCGC GGTGAATACG TTCCCGGGTC TTGTACACAC CGCCCGTCAC ACCATGGGAG TCAGTAACGC CCGAAGCCGG
NTI82-c               AGAATGTCGC GGTGAATACG TTCCCGGGTC TTGTACACAC CGCCCGTCAC ACCATGGGAG TCAGTAACGC CCGAAGCCGG
NTI82-d               AGAATGTCGC GGTGAATACG TTCCCGGGTC TTGTACACAC CGCCCGTCAC ACCATGGGAG TCAGTAACGC CCGAAGCCGG
Q4-a                  AGAATGTCGC GGTGAATACG TTCCCGGGTC TTGTACACAC CGCCCGTCAC ACCATGGGAG TCAGTAACGC CCGAAGCCGG
Q4-b                  AGAATGTCGC GGTGAATACG TTCCCGGGTC TTGTACACAC CGCCCGTCAC ACCATGGGAG TCAGTAACGC CCGAAGCCGG
Q4-c                  AGAATGTCGC GGTGAATACG TTCCCGGGTC TTGTACACAC CGCCCGTCAC ACCATGGGAG TCAGTAACGC CCGAAGCCGG
Q4-d                  AGAATGTCGC GGTGAATACG TTCCCGGGTC TTGTACACAC CGCCCGTCAC ACCATGGGAG TCAGTAACGC CCGAAGCCGG
S076-a                AGAATGTCGC GGTGAATACG TTCCCGGGTC TTGTACACAC CGCCCGTCAC ACCATGGGAG TCAGTAACGC CCGAAGCCGG
S076-b                AGAATGTCGC GGTGAATACG TTCCCGGGTC TTGTACACAC CGCCCGTCAC ACCATGGGAG TCAGTAACGC CCGAAGCCGG
S076-c                AGAATGTCGC GGTGAATACG TTCCCGGGTC TTGTACACAC CGCCCGTCAC ACCATGGGAG TCAGTAACGC CCGAAGCCGG
S076-d                AGAATGTCGC GGTGAATACG TTCCCGGGTC TTGTACACAC CGCCCGTCAC ACCATGGGAG TCAGTAACGC CCGAAGCCGG
SL.1.22               AGAATGTCGC GGTGAATACG TTCCCGGGTC TTGTACACAC CGCCCGTCAC ACCATGGGAG TCAGTAACGC CCGAAGCCGG
VE202-05              AGAATGTCGC GGTGAATACG TTCCCGGGTC TTGTACACAC CGCCCGTCAC ACCATGGGAG TCAGTAACGC CCGAAGCCGG
C.hylemonae           AGAATGTCGC GGTGAATACG TTCCCGGGTC TTGTACACAC CGCCCGTCAC ACCATGGGAG TCAGTAACGC CCGAAGCCGG
```

```
                     1441
35704-a              TGACCCAACC CGTAAGGGAG GGAGCCGTCG AAGGTGGGAC CGATAACTGG GGTGAAGTCG TAACAAGGTA GCCGTATCGG
35704-b              TGACCCAACC CGTAAGGGAG GGAGCCGTCG AAGGTGGGAC CGATAACTGG GGTGAAGTCG TAACAAGGTA GCCGTATCGG
35704-c              TGACCCAACC CGTAAGGGAG GGAGCCGTCG AAGGTGGGAC CGATAACTGG GGTGAAGTCG TAACAAGGTA GCCGTATCGG
35704-d              TGACCCAACC CGTAAGGGAG GGAGCCGTCG AAGGTGGGAC CGATAACTGG GGTGAAGTCG TAACAAGGTA GCCGTATCGG
AM05-22              TGACCCAACC CGTAAGGGAG GGAGCCGTCG AAGGTGGGAC CGATAACTGG GGTGAAGTCG TAACAAGGTA GCCGTATCGG
AM07-30              TGACCCAACC CGTAAGGGAG GGAGCCGTCG AAGGTGGGAC CGATAACTGG GGTGAAGTCG TAACAAGGTA GCCGTATCGG
BL389WT3D-a          TGACCCAACC CGTAAGGGAG GGAGCCGTCG AAGGTGGGAC CGATAACTGG GGTGAAGTCG TAACAAGGTA GCCGTATCGG
BL389WT3D-b          TGACCCAACC CGTAAGGGAG GGAGCCGTCG AAGGTGGGAC CGATAACTGG GGTGAAGTCG TAACAAGGTA GCCGTATCGG
BL389WT3D-c          TGACCCAACC CGTAAGGGAG GGAGCCGTCG AAGGTGGGAC CGATAACTGG GGTGAAGTCG TAACAAGGTA GCCGTATCGG
BL389WT3D-d          TGACCCAACC CGTAAGGGAG GGAGCCGTCG AAGGTGGGAC CGATAACTGG GGTGAAGTCG TAACAAGGTA GCCGTATCGG
CE91-St59-a          TGACCCAACC CGTAAGGGAG GGAGCCGTCG AAGGTGGGAC CGATAACTGG GGTGAAGTCG TAACAAGGTA GCCGTATCGG
CE91-St59-b          TGACCCAACC CGTAAGGGAG GGAGCCGTCG AAGGTGGGAC CGATAACTGG GGTGAAGTCG TAACAAGGTA GCCGTATCGG
CE91-St59-c          TGACCCAACC CGTAAGGGAG GGAGCCGTCG AAGGTGGGAC CGATAACTGG GGTGAAGTCG TAACAAGGTA GCCGTATCGG
CE91-St60-a          TGACCCAACC CGTAAGGGAG GGAGCCGTCG AAGGTGGGAC CGATAACTGG GGTGAAGTCG TAACAAGGTA GCCGTATCGG
CE91-St60-b          TGACCCAACC CGTAAGGGAG GGAGCCGTCG AAGGTGGGAC CGATAACTGG GGTGAAGTCG TAACAAGGTA GCCGTATCGG
CE91-St60-c          TGACCCAACC CGTAAGGGAG GGAGCCGTCG AAGGTGGGAC CGATAACTGG GGTGAAGTCG TAACAAGGTA GCCGTATCGG
FDAARGOS_1227-a      TGACCCAACC CGTAAGGGAG GGAGCCGTCG AAGGTGGGAC CGATAACTGG GGTGAAGTCG TAACAAGGTA GCCGTATCGG
FDAARGOS_1227-b      TGACCCAACC CGTAAGGGAG GGAGCCGTCG AAGGTGGGAC CGATAACTGG GGTGAAGTCG TAACAAGGTA GCCGTATCGG
FDAARGOS_1227-c      TGACCCAACC CGTAAGGGAG GGAGCCGTCG AAGGTGGGAC CGATAACTGG GGTGAAGTCG TAACAAGGTA GCCGTATCGG
FDAARGOS_1227-d      TGACCCAACC CGTAAGGGAG GGAGCCGTCG AAGGTGGGAC CGATAACTGG GGTGAAGTCG TAACAAGGTA GCCGTATCGG
G10-a                TGACCCAACC CGTAAGGGAG GGAGCCGTCG AAGGTGGGAC CGATAACTGG GGTGAAGTCG TAACAAGGTA GCCGTATCGG
G10-b                TGACCCAACC CGTAAGGGAG GGAGCCGTCG AAGGTGGGAC CGATAACTGG GGTGAAGTCG TAACAAGGTA GCCGTATCGG
G10-c                TGACCCAACC CGTAAGGGAG GGAGCCGTCG AAGGTGGGAC CGATAACTGG GGTGAAGTCG TAACAAGGTA GCCGTATCGG
G10-d                TGACCCAACC CGTAAGGGAG GGAGCCGTCG AAGGTGGGAC CGATAACTGG GGTGAAGTCG TAACAAGGTA GCCGTATCGG
GGCC_0168            TGACCCAACC CGTAAGGGAG GGAGCCGTCG AAGGTGGGAC CGATAACTGG GGTGAAGTCG TAACAAGGTA GCCGTATCGG
I10-a                TGACCCAACC CGTAAGGGAG GGAGCCGTCG AAGGTGGGAC CGATAACTGG GGTGAAGTCG TAACAAGGTA GCCGTATCGG
I10-b                TGACCCAACC CGTAAGGGAG GGAGCCGTCG AAGGTGGGAC CGATAACTGG GGTGAAGTCG TAACAAGGTA GCCGTATCGG
I10-c                TGACCCAACC CGTAAGGGAG GGAGCCGTCG AAGGTGGGAC CGATAACTGG GGTGAAGTCG TAACAAGGTA GCCGTATCGG
I10-d                TGACCCAACC CGTAAGGGAG GGAGCCGTCG AAGGTGGGAC CGATAACTGG GGTGAAGTCG TAACAAGGTA GCCGTATCGG
JCM10422-a           TGACCCAACC CGTAAGGGAG GGAGCCGTCG AAGGTGGGAC CGATAACTGG GGTGAAGTCG TAACAAGGTA GCCGTATCGG
JCM10422-b           TGACCCAACC CGTAAGGGAG GGAGCCGTCG AAGGTGGGAC CGATAACTGG GGTGAAGTCG TAACAAGGTA GCCGTATCGG
JCM10422-c           TGACCCAACC CGTAAGGGAG GGAGCCGTCG AAGGTGGGAC CGATAACTGG GGTGAAGTCG TAACAAGGTA GCCGTATCGG
JCM10422-d           TGACCCAACC CGTAAGGGAG GGAGCCGTCG AAGGTGGGAC CGATAACTGG GGTGAAGTCG TAACAAGGTA GCCGTATCGG
MGYG-HGUT-01303-a    TGACCCAACC CGTAAGGGAG GGAGCCGTCG AAGGTGGGAC CGATAACTGG GGTGAAGTCG TAACAAGGTA GCCGTATCGG
MGYG-HGUT-01303-b    TGACCCAACC CGTAAGGGAG GGAGCCGTCG AAGGTGGGAC CGATAACTGG GGTGAAGTCG TAACAAGGTA GCCGTATCGG
MSK.1.16             TGACCCAACC CGTAAGGGAG GGAGCCGTCG AAGGTGGGAC CGATAACTGG GGTGAAGTCG TAACAAGGTA GCCGTATCGG
MSK.1.26             ---------- ---------- ---------- ---------- ---------- ---------- ---------- ----------
S077-U-a             TGACCCAACC CGTAAGGGAG GGAGCCGTCG AAGGTGGGAC CGATAACTGG GGTGAAGTCG TAACAAGGTA GCCGTATCGG
S077-U-b             TGACCCAACC CGTAAGGGAG GGAGCCGTCG AAGGTGGGAC CGATAACTGG GGTGAAGTCG TAACAAGGTA GCCGTATCGG
S077-U-c             TGACCCAACC CGTAAGGGAG GGAGCCGTCG AAGGTGGGAC CGATAACTGG GGTGAAGTCG TAACAAGGTA GCCGTATCGG
S077-U-d             TGACCCAACC CGTAAGGGAG GGAGCCGTCG AAGGTGGGAC CGATAACTGG GGTGAAGTCG TAACAAGGTA GCCGTATCGG
VPI12708-a           TGACCCAACC CGCAAGGGAG GGAGCCGTCG AAGGTGGGAC CGATAACTGG GGTGAAGTCG TAACAAGGTA GCCGTATCGG
VPI12708-b           TGACCCAACC CGTATGGGAG GGAGCCGTCG AAGGTGGGAC CGATAACTGG GGTGAAGTCG TAACAAGGTA GCCGTATCGG
VPI12708-c           TGACCCAACC CGTATGGGAG GGAGCCGTCG AAGGTGGGAC CGATAACTGG GGTGAAGTCG TAACAAGGTA GCCGTATCGG
VPI12708-d           TGACCCAACC CGCAAGGGAG GGAGCCGTCG AAGGTGGGAC CGATAACTGG GGTGAAGTCG TAACAAGGTA GCCGTATCGG
DFI.1.130            TGACCCAACC CGCAAGGGAG GGAGCCGTCG AAGGTGGGAC CGATAACTGG GGTGAAGTCG TAACAAGGTA GCCGTATCGG
DFI.1.161            ---------- ---------- ---------- ---------- ---------- ---------- ---------- ----------
DFI.1.162            TGACCCAACC CGCAAGGGAG GGAGCCGTCG AAGGTGGGAC CGATAACTGG GGTGAAGTCG TAACAAGGTA GCCGTATCGG
DFI.1.217            TGACCCAACC CGCAAGGGAG GGAGCCGTCG AAGGTGGGAC CGATAACTGG GGTGAAGTCG TAACAAGGTA GCCGTATCGG
DFI.1.234            TGACCCAACC CGCAAGGGAG GGAGCCGTCG AAGGTGGGAC CGATAACTGG GGTGAAGTCG TAACAAGGTA GCCGTATCGG
DFI.1.60             TGACCCAACC CGTATGGGAG GGAGCCGTCG AAGGTGGGAC CGATAACTGG GGTGAAGTCG TAACAAGGTA GCCGTATCGG
DFI.4.63             TGACCCAACC CGTATGGGAG GGAGCCGTCG AAGGTGGGAC CGATAACTGG GGTGAAGTCG TAACAAGGTA GCCGTATCGG
JCM10419-a           TGACCCAACC CGCAAGGGAG GGAGCCGTCG AAGGTGGGAC CGATAACTGG GGTGAAGTCG TAACAAGGTA GCCGTATCGG
JCM10419-b           TGACCCAACC CGCAAGGGAG GGAGCCGTCG AAGGTGGGAC CGATAACTGG GGTGAAGTCG TAACAAGGTA GCCGTATCGG
JCM10419-c           TGACCCAACC CGCAAGGGAG GGAGCCGTCG AAGGTGGGAC CGATAACTGG GGTGAAGTCG TAACAAGGTA GCCGTATCGG
JCM10419-d           TGACCCAACC CGCAAGGGAG GGAGCCGTCG AAGGTGGGAC CGATAACTGG GGTGAAGTCG TAACAAGGTA GCCGTATCGG
JCM10420-a           TGACCCAACC CGCAAGGGAG GGAGCCGTCG AAGGTGGGAC CGATAACTGG GGTGAAGTCG TAACAAGGTA GCCGTATCGG
JCM10420-b           TGACCCAACC CGCAAGGGAG GGAGCCGTCG AAGGTGGGAC CGATAACTGG GGTGAAGTCG TAACAAGGTA GCCGTATCGG
JCM10420-c           TGACCCAACC CGCAAGGGAG GGAGCCGTCG AAGGTGGGAC CGATAACTGG GGTGAAGTCG TAACAAGGTA GCCGTATCGG
JCM10420-d           TGACCCAACC CGCAAGGGAG GGAGCCGTCG AAGGTGGGAC CGATAACTGG GGTGAAGTCG TAACAAGGTA GCCGTATCGG
JCM10423-a           TGACCCAACC CGCAAGGGAG GGAGCCGTCG AAGGTGGGAC CGATAACTGG GGTGAAGTCG TAACAAGGTA GCCGTATCGG
JCM10423-b           TGACCCAACC CGCAAGGGAG GGAGCCGTCG AAGGTGGGAC CGATAACTGG GGTGAAGTCG TAACAAGGTA GCCGTATCGG
JCM10423-c           TGACCCAACC CGCAAGGGAG GGAGCCGTCG AAGGTGGGAC CGATAACTGG GGTGAAGTCG TAACAAGGTA GCCGTATCGG
JCM10423-d           TGACCCAACC CGCAAGGGAG GGAGCCGTCG AAGGTGGGAC CGATAACTGG GGTGAAGTCG TAACAAGGTA GCCGTATCGG
MO32-a               TGACCCAACC CGCAAGGGAG GGAGCCGTCG AAGGTGGGAC CGATAACTGG GGTGAAGTCG TAACAAGGTA GCCGTATCGG
MO32-b               TGACCCAACC CGCAAGGGAG GGAGCCGTCG AAGGTGGGAC CGATAACTGG GGTGAAGTCG TAACAAGGTA GCCGTATCGG
MO32-c               TGACCCAACC CGCAAGGGAG GGAGCCGTCG AAGGTGGGAC CGATAACTGG GGTGAAGTCG TAACAAGGTA GCCGTATCGG
MO32-d               TGACCCAACC CGCAAGGGAG GGAGCCGTCG AAGGTGGGAC CGATAACTGG GGTGAAGTCG TAACAAGGTA GCCGTATCGG
MSK.5.24             TGACCCAACC CGCAAGGGAG GGAGCCGTCG AAGGTGGGAC CGATAACTGG GGTGAAGTCG TAACAAGGTA GCCGTATCGG
NB2A-7-D5            TGACCCAACC CGCAAGGGAG GGAGCCGTCG AAGGTGGGAC CGATAACTGG GGTGAAGTCG TAACAAGGTA GCCGTATCGG
NTI82-a              TGACCCAACC CGCAAGGGAG GGAGCCGTCG AAGGTGGGAC CGATAACTGG GGTGAAGTCG TAACAAGGTA GCCGTATCGG
NTI82-b              TGACCCAACC CGCAAGGGAG GGAGCCGTCG AAGGTGGGAC CGATAACTGG GGTGAAGTCG TAACAAGGTA GCCGTATCGG
NTI82-c              TGACCCAACC CGCAAGGGAG GGAGCCGTCG AAGGTGGGAC CGATAACTGG GGTGAAGTCG TAACAAGGTA GCCGTATCGG
NTI82-d              TGACCCAACC CGCAAGGGAG GGAGCCGTCG AAGGTGGGAC CGATAACTGG GGTGAAGTCG TAACAAGGTA GCCGTATCGG
Q4-a                 TGACCCAACC CGTATGGGAG GGAGCCGTCG AAGGTGGGAC CGATAACTGG GGTGAAGTCG TAACAAGGTA GCCGTATCGG
Q4-b                 TGACCCAACC CGTATGGGAG GGAGCCGTCG AAGGTGGGAC CGATAACTGG GGTGAAGTCG TAACAAGGTA GCCGTATCGG
Q4-c                 TGACCCAACC CGTATGGGAG GGAGCCGTCG AAGGTGGGAC CGATAACTGG GGTGAAGTCG TAACAAGGTA GCCGTATCGG
Q4-d                 TGACCCAACC CGTATGGGAG GGAGCCGTCG AAGGTGGGAC CGATAACTGG GGTGAAGTCG TAACAAGGTA GCCGTATCGG
S076-a               TGACCCAACC CGCAAGGGAG GGAGCCGTCG AAGGTGGGAC CGATAACTGG GGTGAAGTCG TAACAAGGTA GCCGTATCGG
S076-b               TGACCCAACC CGCAAGGGAG GGAGCCGTCG AAGGTGGGAC CGATAACTGG GGTGAAGTCG TAACAAGGTA GCCGTATCGG
S076-c               TGACCCAACC CGCAAGGGAG GGAGCCGTCG AAGGTGGGAC CGATAACTGG GGTGAAGTCG TAACAAGGTA GCCGTATCGG
S076-d               TGACCCAACC CGCAAGGGAG GGAGCCGTCG AAGGTGGGAC CGATAACTGG GGTGAAGTCG TAACAAGGTA GCCGTATCGG
SL.1.22              TGACCCAACC CGCAAGGGAG GGAGCCGTCG AAGGTGGGAC CGATAACTGG GGTGAAGTCG TAACAAGGTA GCCGTATCGG
VE202-05             TGACCCAACC CGCAAGGGAG GGAGCCGTCG AAGGTGGGAC CGATAACTGG GGTGAAGTCG TAACAAGGTA GCCGTATCGG
C.hylemonae          TGACCTAACC G--AAAGGAA GGAGCCGTCG AAGGTGGGAC CGATAACTGG GGTGAAGTCG TAACAAGGTA GCCGTATCGG
```

```
                     1521
35704-a              AAGGTGCGGC TGGATCACCT CCTTT
35704-b              AAGGTGCGGC TGGATCACCT CCTTT
35704-c              AAGGTGCGGC TGGATCACCT CCTTT
35704-d              AAGGTGCGGC TGGATCACCT CCTTT
AM05-22              AAGGTGCGGC TGGATCACCT -----
AM07-30              AAGGTGCGGC TGGATCACCT -----
BL389WT3D-a          AAGGTGCGGC TGGATCACCT CCTTT
BL389WT3D-b          AAGGTGCGGC TGGATCACCT CCTTT
BL389WT3D-c          AAGGTGCGGC TGGATCACCT CCTTT
BL389WT3D-d          AAGGTGCGGC TGGATCACCT CCTTT
CE91-St59-a          AAGGTGCGGC TGGATCACCT CCTTT
CE91-St59-b          AAGGTGCGGC TGGATCACCT CCTTT
CE91-St59-c          AAGGTGCGGC TGGATCACCT CCTTT
CE91-St60-a          AAGGTGCGGC TGGATCACCT CC-TT
CE91-St60-b          AAGGTGCGGC TGGATCACCT CCTTT
CE91-St60-c          AAGGTGCGGC TGGATCACCT CC-TT
FDAARGOS_1227-a      AAGGTGCGGC TGGATCACCT CCTTT
FDAARGOS_1227-b      AAGGTGCGGC TGGATCACCT CCTTT
FDAARGOS_1227-c      AAGGTGCGGC TGGATCACCT CCTTT
FDAARGOS_1227-d      AAGGTGCGGC TGGATCACCT CCTTT
G10-a                AAGGTGCGGC TGGATCACCT CCTTT
G10-b                AAGGTGCGGC TGGATCACCT CCTTT
G10-c                AAGGTGCGGC TGGATCACCT CCTTT
G10-d                AAGGTGCGGC TGGATCACCT CCTTT
GGCC_0168            AAGGTGCGGC TGGATCACCT -----
I10-a                AAGGTGCGGC TGGATCACCT CCTTT
I10-b                AAGGTGCGGC TGGATCACCT CCTTT
I10-c                AAGGTGCGGC TGGATCACCT CCTTT
I10-d                AAGGTGCGGC TGGATCACCT CCTTT
JCM10422-a           AAGGTGCGGC TGGATCACCT CCTTT
JCM10422-b           AAGGTGCGGC TGGATCACCT CCTTT
JCM10422-c           AAGGTGCGGC TGGATCACCT CCTTT
JCM10422-d           AAGGTGCGGC TGGATCACCT CCTTT
MGYG-HGUT-01303-a    AAGGTGCGGC TGGATCACCT -----
MGYG-HGUT-01303-b    AAGGTGCGGC TGGATCACCT -----
MSK.1.16             AAGGTGCGGC TGGATCACCT -----
MSK.1.26             ---------- ---------- -----
S077-U-a             AAGGTGCGGC TGGATCACCT CCTTT
S077-U-b             AAGGTGCGGC TGGATCACCT CCTTT
S077-U-c             AAGGTGCGGC TGGATCACCT CCTTT
S077-U-d             AAGGTGCGGC TGGATCACCT CCTTT
VPI12708-a           AAGGTGCGGC TGGATCACCT -----
VPI12708-b           AAGGTGCGGC TGGATCACCT -----
VPI12708-c           AAGGTGCGGC TGGATCACCT -----
VPI12708-d           AAGGTGCGGC TGGATCACCT -----
DFI.1.130            AAGGTGCGGC TGGATCACCT -----
DFI.1.161            ---------- ---------- -----
DFI.1.162            AAGGTGCGGC TGGATCACCT -----
DFI.1.217            AAGGTGCGGC TGGATCACCT -----
DFI.1.234            AAGGTGCGGC TGGATCACCT -----
DFI.1.60             AAGGTGCGGC TGGATCACCT -----
DFI.4.63             AAGGTGCGGC TGGATCACCT -----
JCM10419-a           AAGGTGCGGC TGGATCACCT CCTTT
JCM10419-b           AAGGTGCGGC TGGATCACCT CCTTT
JCM10419-c           AAGGTGCGGC TGGATCACCT CCTTT
JCM10419-d           AAGGTGCGGC TGGATCACCT CCTTT
JCM10420-a           AAGGTGCGGC TGGATCACCT CCTTT
JCM10420-b           AAGGTGCGGC TGGATCACCT CCTTT
JCM10420-c           AAGGTGCGGC TGGATCACCT CCTTT
JCM10420-d           AAGGTGCGGC TGGATCACCT CCTTT
JCM10423-a           AAGGTGCGGC TGGATCACCT CCTTT
JCM10423-b           AAGGTGCGGC TGGATCACCT CCTTT
JCM10423-c           AAGGTGCGGC TGGATCACCT CCTTT
JCM10423-d           AAGGTGCGGC TGGATCACCT CCTTT
MO32-a               AAGGTGCGGC TGGATCACCT CCTTT
MO32-b               AAGGTGCGGC TGGATCACCT CCTTT
MO32-c               AAGGTGCGGC TGGATCACCT CCTTT
MO32-d               AAGGTGCGGC TGGATCACCT CCTTT
MSK.5.24             AAGGTGCGGC TGGATCACCT -----
NB2A-7-D5            AAGGTGCGGC TGGATCACCT -----
NTI82-a              AAGGTGCGGC TGGATCACCT CCTTT
NTI82-b              AAGGTGCGGC TGGATCACCT CCTTT
NTI82-c              AAGGTGCGGC TGGATCACCT CCTTT
NTI82-d              AAGGTGCGGC TGGATCACCT CCTTT
Q4-a                 AAGGTGCGGC TGGATCACCT CCTTT
Q4-b                 AAGGTGCGGC TGGATCACCT CCTTT
Q4-c                 AAGGTGCGGC TGGATCACCT CCTTT
Q4-d                 AAGGTGCGGC TGGATCACCT CCTTT
S076-a               AAGGTGCGGC TGGATCACCT CCTTT
S076-b               AAGGTGCGGC TGGATCACCT CCTTT
S076-c               AAGGTGCGGC TGGATCACCT CCTTT
S076-d               AAGGTGCGGC TGGATCACCT CCTTT
SL.1.22              AAGGTGCGGC TGGATCACCT -----
VE202-05             AAGGTGCGGC TGGATCACCT -----
C.hylemonae          AAGGTGCGGC TGGATCACCT CCTTT
```
